# A small-molecule oral agonist of the human glucagon-like peptide-1 receptor

**DOI:** 10.1101/2020.09.29.319483

**Authors:** David A. Griffith, David J. Edmonds, Jean-Phillipe Fortin, Amit S. Kalgutkar, J. Brent Kuzmiski, Paula M. Loria, Aditi R. Saxena, Scott W. Bagley, Clare Buckeridge, John M. Curto, David R. Derksen, João M. Dias, Matthew C. Griffor, Seungil Han, V. Margaret Jackson, Margaret S. Landis, Daniel J. Lettiere, Chris Limberakis, Yuhang Liu, Alan M. Mathiowetz, David W. Piotrowski, David A. Price, Roger B. Ruggeri, David A. Tess

**Affiliations:** Pfizer Worldwide Research and Development, Cambridge, MA 02139, USA; Pfizer Worldwide Research and Development, Cambridge, MA 02139, USA, at the time the work was performed; Pfizer Worldwide Research and Development, Groton, CT 06340, USA

## Abstract

Peptide agonists of the glucagon-like peptide-1 receptor (GLP-1R) have revolutionized diabetes therapy, but their use has been limited by the requirement for injection. Here we describe the first effective, orally bioavailable small molecule GLP-1R agonists. A sensitized high-throughput screen identified a series of small molecule GLP-1R agonists that were optimized to promote endogenous GLP-1R signaling with nM potency. These small molecule agonists increased insulin levels in primates but not rodents, which is explained by a cryo-EM structure that revealed a binding pocket requiring primate-specific tryptophan 33. Importantly, oral administration of agonist PF-06882961 to healthy humans produced dose-dependent declines in serum glucose (NCT03309241). This opens the door to a new era of oral small molecule therapies that target the well-validated GLP-1R pathway for metabolic health.

**One Sentence Summary:** PF-06882961 is an orally administered small molecule that activates the GLP-1 receptor to lower blood glucose in humans.

## Main Text

Glucagon-like peptide-1 (GLP-1), a neuroendocrine hormone, is derived from a proglucagon precursor (*1*) and secreted by intestinal enteroendocrine L cells in response to nutrient intake (*2*), predominantly in the form of GLP-1(7-36) amide (henceforth GLP-1) (*3*). Activation of the GLP-1 receptor (GLP-1R) by GLP-1 stimulates insulin release and inhibits glucagon secretion in a glucose-dependent manner (*4*). Also, GLP-1 delays gastric emptying (*5*), increases satiety, suppresses food intake, and reduces weight in humans (*6, 7*). Multiple injectable peptidic GLP-1R agonists are approved for the treatment of Type 2 diabetes mellitus (T2DM) (*8*), including liraglutide which is also approved for the treatment of obesity (*9*). Excitement has grown in this drug class, with several GLP-1R agonists demonstrating benefit in cardiovascular outcomes studies (*10*). However, a drawback of these medicines has been the necessity for administration by subcutaneous injection, which limits patient utilization and may reduce opportunities for fixed-dose combination treatments with other small-molecule drugs. Importantly, patients prefer, and are more likely to adhere to, an oral drug regimen versus an injectable alternative (*11*). An orally administered formulation of the peptidic GLP-1R agonist semaglutide was recently approved for the treatment of T2DM (*12*). This peptidic drug is co-formulated with sodium N-[8-(2-hydroxybenzoyl] amino) caprylate (SNAC), a purported gastric absorption enhancer, to promote oral bioavailability. The dosage must be taken once daily in the fasted state with minimal liquid and at a substantially higher dose than the approved once-weekly injectable formulation (*12, 13*). Thus, we sought to identify a small-molecule GLP-1R agonist that is orally bioavailable using standard formulations, and has the potential to be combined with other oral small molecule therapeutics.

The GLP-1R is a seven-transmembrane-spanning, class B, G protein-coupled receptor (GPCR) (*14*). Class B GPCRs, including GLP-1R, are activated by endogenous peptide hormones, and the development of small-molecule agonists of these receptors has proven particularly challenging (*14*). Significant prior efforts across the pharmaceutical industry have failed to identify potent and efficacious small-molecule agonists of the GLP-1R (*15, 16*). Given the significant therapeutic value of this mechanism, we pursued a novel high-throughput screening strategy that identified a series of small-molecule GLP-1R agonist leads. Optimization of the lead series resulted in potent agonists that activate the GLP-1R in an unprecedented manner. The series includes the clinical development candidate PF-06882961, which we show has robust preclinical efficacy, oral bioavailability, and evidence of glucose-lowering in healthy human participants.

## Results

### Development of a sensitized assay to identify weak GLP-1R agonists

Binding of GLP-1 to its receptor activates the guanine nucleotide-binding (G) alpha stimulatory subunit (Gαs) of the heterotrimeric G protein complex, stimulating adenylate cyclase activity, and thereby increasing intracellular concentrations of cyclic adenosine monophosphate (cAMP) (*17*). Activation of the GLP-1R by GLP-1 also results in recruitment of β-arrestin-1 (βArr1) and β-arrestin-2 (βArr2), which mediate receptor internalization and may serve as scaffolds for G protein-independent signaling (*18, 19*). The human GLP-1R has an unusually high activation barrier and is essentially silent in the absence of an agonist ligand, relative to other class B GPCRs (e.g. the gastric inhibitory polypeptide and parathyroid hormone 1 receptors) (*20*). We hypothesized that a positive allosteric modulator (PAM) could be used as a tool to lower this activation barrier, thereby increasing assay sensitivity and facilitating the detection of weak agonists in a cell-based functional assay.

The PAM 4-(3-(benzyloxy)phenyl)-2-ethylsulfinyl-6-(trifluoromethyl)pyrimidine (BETP, Fig. 1A) had been reported to potentiate GLP-1R-mediated cAMP signaling in response to weak peptidic agonists, including the GLP-1 metabolite GLP-1(9–36)amide (*21*) and the GLP-1R/glucagon-receptor dual agonist oxyntomodulin (*22*). Further work demonstrated that BETP positively modulates GLP-1R function through covalent modification of cysteine 347 located on the third intracellular loop (*23*). We postulated that a BETP-sensitized assay would prove effective for identifying weak agonists and were pleased to observe potentiation of GLP-1R-mediated cAMP signaling in response to the weak non-peptide GLP-1R agonist BOC5 (*24*) (Fig. S1A) in Chinese hamster ovary (CHO) cells stably expressing the human GLP-1R (Fig. 1B). In particular, the positive impact on maximal effect (E_max_) in this assay format significantly improves the chances of identifying weak agonists during high-throughput screening (HTS). Further confidence in the assay design came from testing the effects of BETP on receptor-mediated βArr signaling. Peptide agonist **1** (Fig. S1B), identified during our earlier efforts to design orally available peptidic GLP-1R agonists (*25*), is a full (E_max_) cAMP agonist (Fig. 1C), but a partial βArr agonist (Fig. 1D) at the GLP-1R. BETP improved the potencies of peptide **1** for both pathways and potentiated the E_max_ for βArr recruitment (Fig. 1C and 1D, Table S1). The observed potentiation of both the cAMP and βArr signaling pathways was consistent with our hypothesis that BETP treatment resulted in general receptor sensitization, rather than a pathway-specific signaling amplification.

**Fig. 1.**
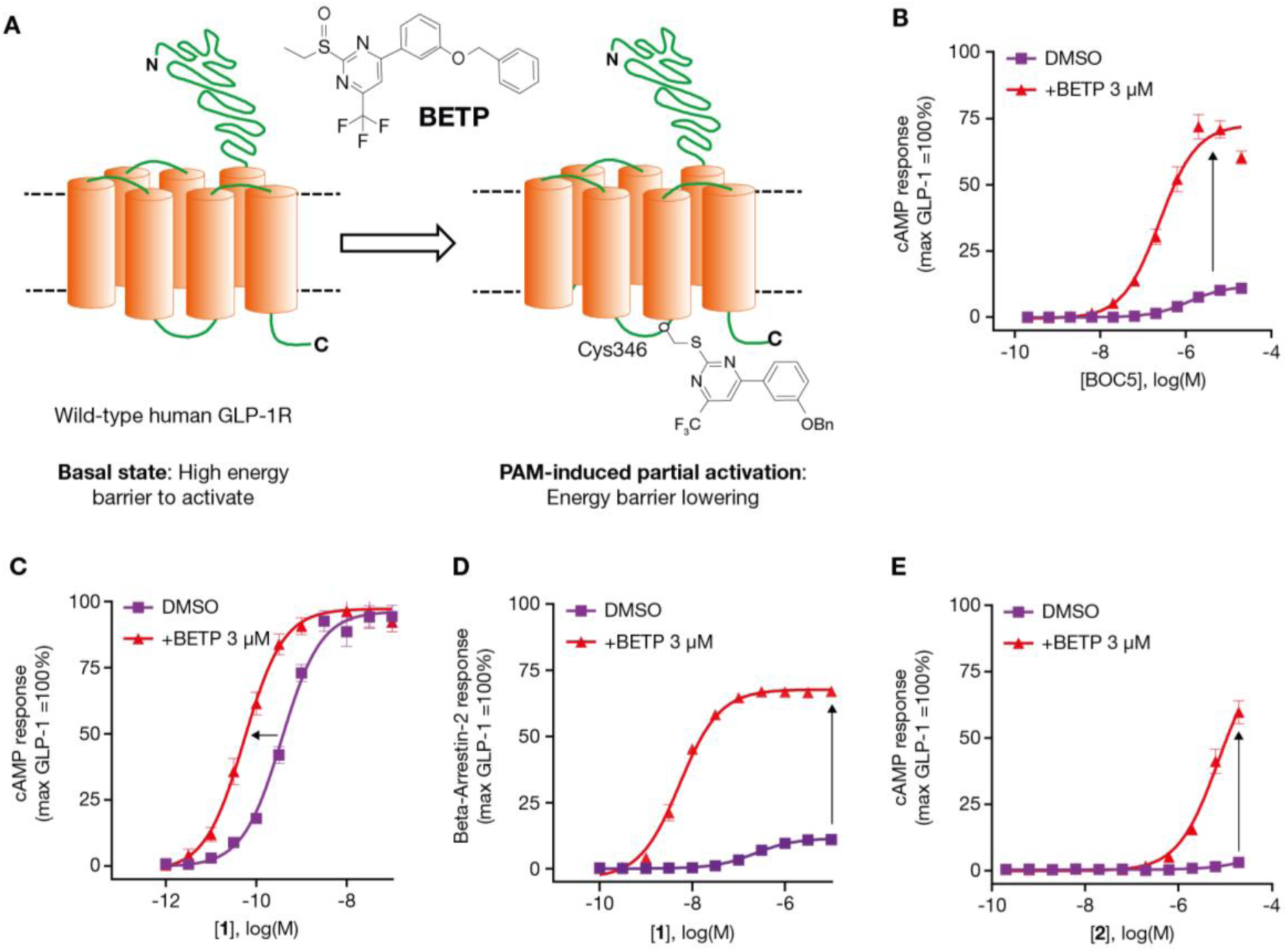
Identification of small-molecule GLP-1R agonists in a CHO-GLP-1R cellular assay in the absence or presence of the positive allosteric modulator BETP. (**A**) Assay concept: Covalent modification of Cys347 in the GLP-1R by BETP lowers receptor activation barrier, enabling the identification of weak agonists. (**B–D**) Validation of the BETP-sensitized screening assay. (**B**) BETP potentiates agonist-induced cAMP production of small molecule (BOC5) (Fig. S1). (**C–D**) BETP potentiates cAMP production (**C**) and β-arrestin recruitment (**D**) by peptide **1** (Fig. S1) at the human GLP-1R. (**E**), cAMP curves of a representative small molecule HTS hit, compound **2** (Fig. 2). Data represent the mean ± SEM. BETP, 4-(3-(benzyloxy)phenyl)-2-ethylsulfinyl-6-(trifluoromethyl)pyrimidine; cAMP, cyclic adenosine monophosphate; DMSO, dimethyl sulfoxide; GLP-1R, glucagon-like peptide-1 receptor; HTS, high-throughput screening; PAM, positive allosteric modulator; SEM, standard error of the mean.

Our BETP-sensitized cAMP screening assay (SA +BETP) was adapted to a single-point format and employed in an HTS of 2.8 million compounds from the Pfizer compound collection. A hit was defined by a threshold of >30% effect (i.e., >30% of the E_max_ of GLP-1) at 10 µM and the screen resulted in a low confirmed hit-rate of 0.013%. A series of pyrimidine derivatives exemplified by **2** emerged from these hits (Fig. 2A). Compound **2** was inactive as a GLP-1R agonist in the unpotentiated cAMP screening assay (SA), which did not include BETP, but demonstrated a ∼70% effect at 20 µM in the presence of BETP (Fig. 1E). This non-traditional screening approach carried the risk that the GLP-1R agonist lead series might remain dependent on the presence of BETP to activate the receptor, and it was unclear at the outset whether we would observe GLP-1R agonism in the absence of BETP. However, as analogs were identified with improved cAMP potency in the BETP-sensitized assay, we also observed a gradual increase in signaling efficacy in the absence of BETP (Fig. 2B) to the point where a dose-response curve and half-maximal effective concentration (EC_50_) could be defined. For weaker agonists, BETP potentiated cAMP potency by ∼100-fold. As potency improved to below ∼100 nM in the absence of BETP, the impact of the PAM gradually diminished (Fig. 2C). This observation is consistent with the relatively minimal effect of BETP on signaling of the potent GLP-1R endogenous agonist GLP-1, compared to the weaker agonist GLP-1(9-36)amide, whose effects are potentiated by BETP (*21*).

**Fig. 2.**
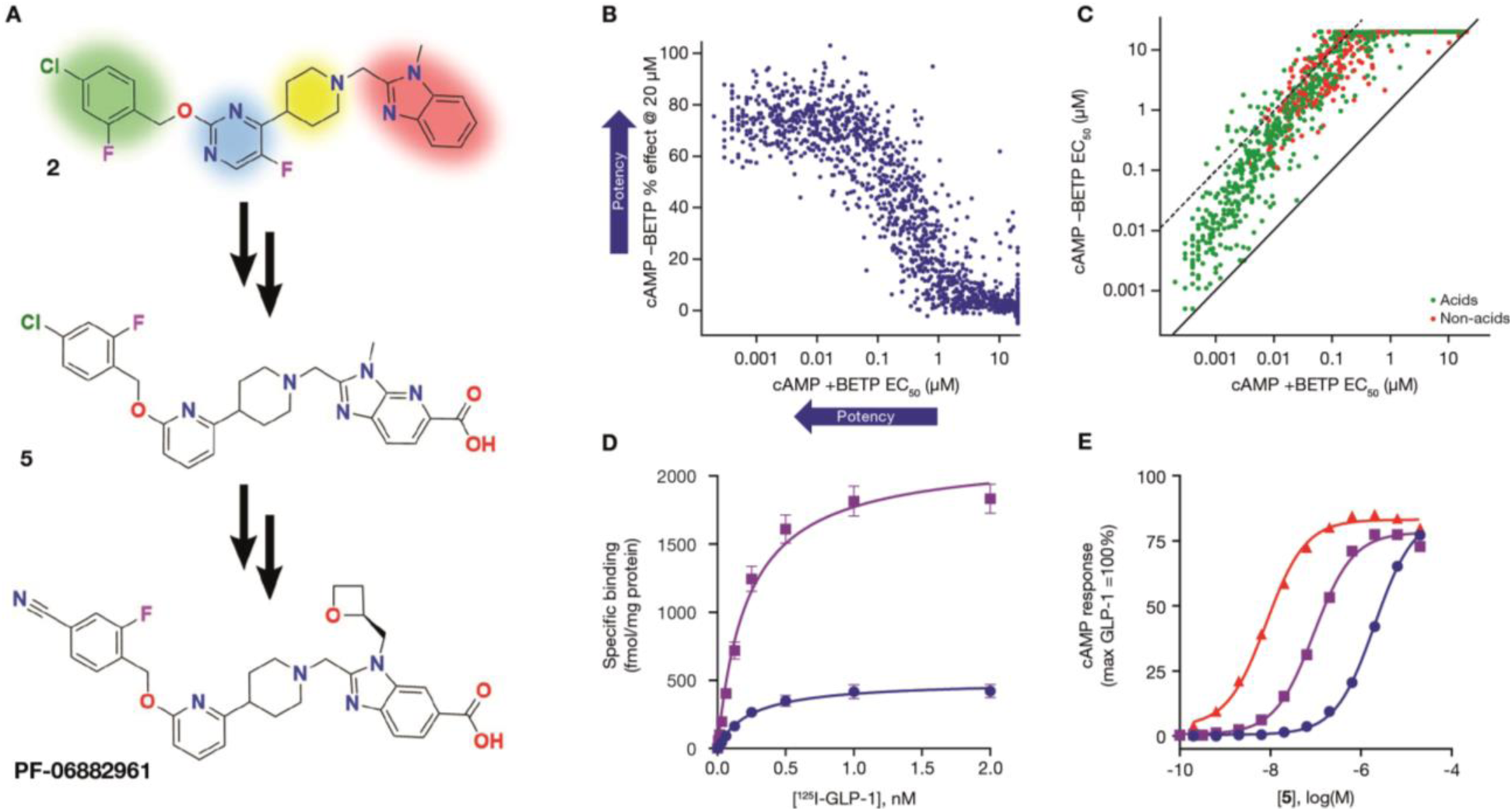
Optimization of small-molecule 2 culminating in the identification of the clinical candidate PF-06882961. **(A)** Structure of small molecule HTS hit **2**, intermediate analog **5**, and clinical candidate PF-06882961. Four structural regions were considered in our efforts to improve GLP-1R agonist activity of **2**: the piperidine ring (yellow), the benzyl ether (green), the 5-fluoro-pyrimidine (blue), and the benzimidazole (red). **(B)** Early evidence of activity in the cAMP assay without BETP. Increased cAMP release (% effect) at 20 µM concentration of test compounds was observed in the absence of BETP as potency (EC_50_) improved in the presence of BETP. **(C)** Small molecule agonist activity independent of BETP sensitization. Increased cAMP potency (EC_50_) was observed in the absence of BETP as potency improved in the presence of BETP (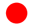, non-acids; 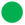, acid-containing analogs). **(D–E)** Candidate selection CHO-GLP-1R cell line with lower GLP-1R expression level confirms the efficacy-driven nature of small molecule **5** agonism at the human GLP-1R. Data represent the mean ± SEM. **(D)** Saturation binding analysis in CHO cells expressing higher (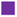, SA) and lower (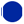, CS) human GLP-1R density. **(E)** Small molecule **5** induced cAMP signaling in CS cell line (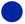), as well as in the SA in the presence (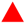) or absence (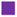) of BETP. BETP, 4-(3-(benzyloxy)phenyl)-2-ethylsulfinyl-6-(trifluoromethyl)pyrimidine; cAMP, cyclic adenosine monophosphate; CHO, Chinese hamster ovary; CS, candidate selection assay; EC_50_, concentration at half maximal effect; GLP-1R, glucagon-like peptide-1 receptor; HTS, high-throughput screening; SA, screening assay; SEM, standard error of the mean.

### Lead series optimization

Our goal was to enhance interactions of the small-molecule agonists with the GLP-1R through lowering the energy barrier to achieve the receptor-bound agonist conformation, and by adjusting polar substituents, with minimal increases in molecular weight. Four structural regions were considered in our quest to improve GLP-1R agonist activity of small molecule **2**: the piperidine ring (yellow), the benzyl ether (green), the 5-fluoro-pyrimidine (blue), and the benzimidazole (red), as highlighted in Fig. 2A. The piperidine ring (yellow) proved optimal in structure-activity relationship (SAR) studies, although other 6-membered rings (e.g., piperazine, cyclohexane) also demonstrated GLP-1R agonism. Likewise, the 4-chloro-2-fluoro-benzyl ether substituent (green) was effective at activating the receptor. Small substituents at the 4-position (e.g., chloro, fluoro, cyano) provided the greatest potency. Significant potency improvements were achieved by replacing the 5-fluoro-pyrimidine (blue) with a pyridyl group. The pyridine appears to optimize the preferred conformation of the pendent benzyl ether through repulsion of the oxygen and nitrogen lone pairs (*26*). Removal of the fluorine likely favors the preferred torsion angle between the aromatic ring and the piperidine. Combining these changes with the incorporation of a more polar 6-aza-benzimidazole led to **3** (EC_50_ = 77 nM in SA + BETP), which was >100-fold more potent than HTS hit **2** (Fig. S2, Table S2) and now active in the cAMP assay without BETP (SA EC_50_ = 2600 nM). It was also encouraging that this compound recruited βArr in the presence of BETP (EC_50_ = 9600 nM, Table S3). However, small-molecule **3** was quite lipophilic (logD_7.4_ = 5.7), resulting in high metabolic intrinsic clearance in human liver microsomes (HLM) (CL_int_ = 130 mL/min/kg, Table S4), which would likely lead to a short pharmacokinetic half-life (t_1/2_) and the requirement for an unacceptably high daily dose in humans. The high lipophilicity was also associated with off-target pharmacology such as inhibition of the human ether-a-go-go-related gene (hERG) ion channel (IC_50_ = 5.6 µM, Table S4). Inhibition of the hERG channel can cause fatal cardiac arrhythmias in humans (*27*).

During our earlier efforts to identify orally available peptides (*25*), we recognized the important role that carboxylic acid substituents played in activating the GLP-1R. Therefore, we sought to incorporate an acid substituent to improve both potency and physiochemical properties. In the absence of structural information for the GLP-1R, the design of acid-containing ligands was driven by SAR and the observation that polarity was better tolerated in the benzimidazole (red) region (Fig. 2A). For example, the introduction of a carboxylic acid-containing substituent at the 7-position of the benzimidazole (red) yielded **4** (Fig. S2 and Table S2). Compound **4** demonstrated comparable potency to **3** (EC_50_ = 4.6 µM; Table S2), but with markedly lower lipophilicity (logD_7.4_ = 2.3) (Table S4), indicating that the acid was likely making a productive interaction (*28*). A carboxylic acid directly attached to the 6-position of the benzimidazole proved optimal towards improvements in potency. For example, **5** (Fig. 2A) was a potent GLP-1R agonist (SA EC_50_ = 95 nM; Table S2) with moderate CL_int_ in HLM and human hepatocyte metabolic stability assays, and excellent selectivity against the hERG channel (>100 µM, Table S4).

During lead optimization, less-sensitive cell assays are valuable for distinguishing the contributions of affinity and efficacy in the cellular response to an agonist (*29*). Therefore, we sought a less-sensitive functional assay with reduced receptor expression levels to enable the optimization of efficacy-driven GLP-1R agonists for clinical development. Considering the lack of robust cellular models for endogenous human GLP-1R, we developed a cell line with a GLP-1R density more comparable with endogenous tissue levels (*30*). The receptor density (Fig. 2D) for the candidate selection (CS) cell line (CS B_max_: 500 ± 28 fmol/mg) was ∼4.3-fold lower than the cell line used for primary screens (SA B_max_: 2200 ± 80 fmol/mg). In the setting of lower GLP-1R levels in this CS cell line, small molecule **5** remained a full agonist (Fig. 2E) but was ∼20-fold less potent (CS EC_50_ = 2.1 µM, Table S2), suggesting that further potency improvements would be required. Optimizing the substituent on the benzimidazole nitrogen proved a fruitful approach to improve potency without detrimentally impacting physiochemical properties, with smaller, more polar groups preferred. For example, a methylene-linked oxetane increased potency ∼100-fold relative to the methyl substituent of **5**, leading to the identification of PF-06882961, which is a full agonist (EC_50_ = 13 nM) in the CS cAMP assay (Fig. 3A, Table S2). PF-06882961 also incorporates a nitrile replacement for the chloride in the benzyl ether region, which served to reduce CL_int_ in HLM as well as in human hepatocytes (Table S4).

**Fig. 3.**
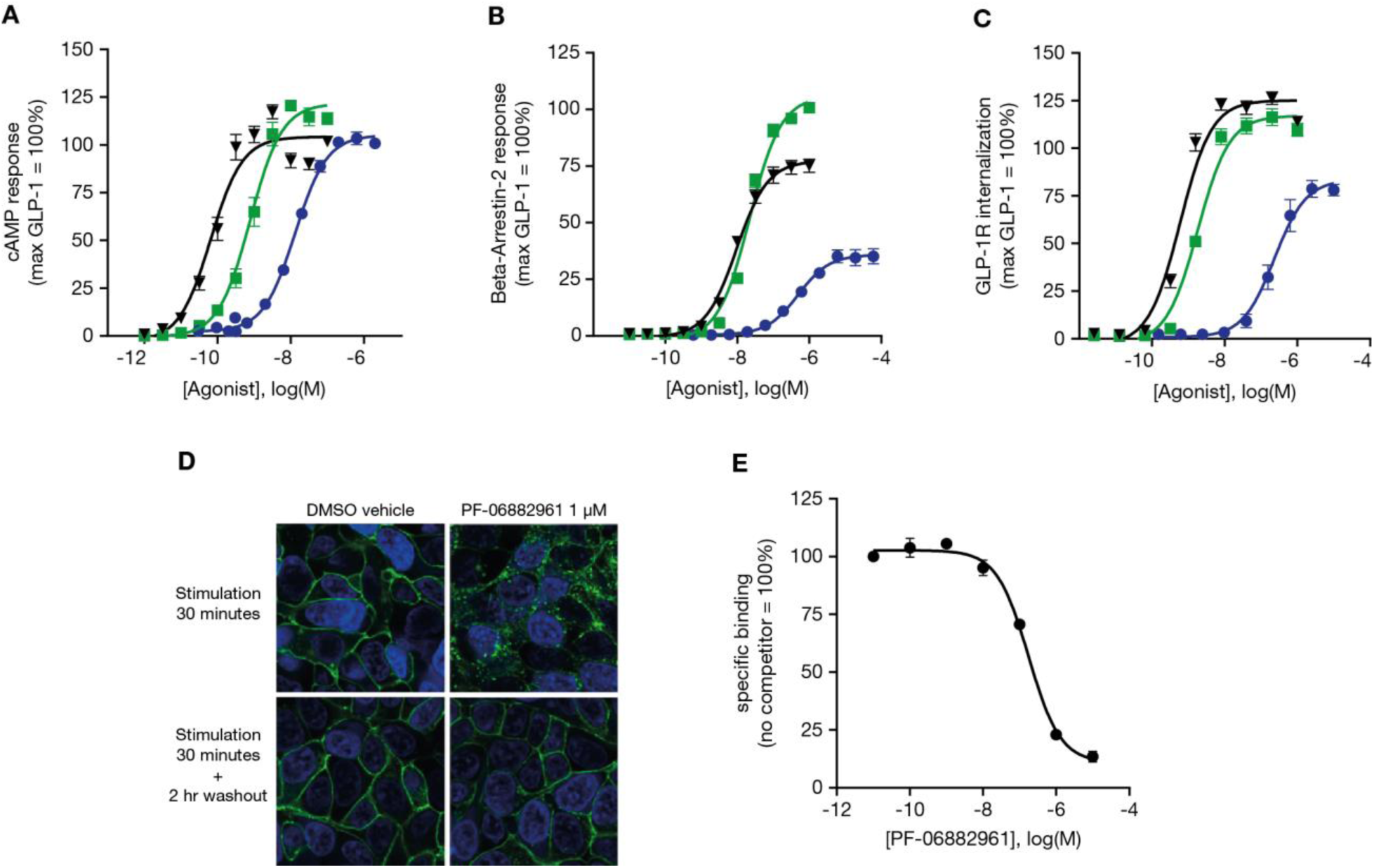
Molecular pharmacology of small molecule GLP-1R agonist PF-06882961. (**A**) Average cAMP curves for exenatide (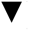), liraglutide (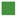), and PF-06882961 (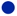) in the candidate selection cell line. (**B**) Average β-arrestin recruitment curves for exenatide (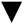), liraglutide (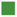), and PF-06882961 (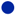). (**C**) GLP-1R agonist driven receptor internalization assessed using FAP-tagged human GLP-1R stably expressed in HEK293 cells. Data represent the mean ± SEM from three independent experiments, each performed in triplicate. (**D**) Assessment of PF-06882961-induced internalization and recycling of a GFP-tagged human GLP-1R (green) in HEK 293 cell construct (blue nuclear staining) using confocal microscopy. (**E**) Competition binding curve for PF-06882961 using the [^3^H]-PF-06883365 probe. Data represent the mean ± SEM. cAMP, cyclic adenosine monophosphate; CHO, Chinese hamster ovary; DMSO, dimethyl sulfoxide; FAP, fluorogen-activated protein; GFP; green fluorescent protein; GLP-1R, glucagon-like peptide-1 receptor; HEK 293, human embryonic kidney 293; SEM, standard error of the mean.

### Molecular pharmacology of PF-06882961

The individual contributions of cAMP signaling and βArr recruitment pathways towards the therapeutic efficacy of marketed GLP-1R agonists remain ill-defined (*18, 19*). Therefore, we sought *in vitro* signaling profiles that were comparable to the marketed peptide agonists during our candidate selection efforts. PF-06882961 agonist activity was assessed at both the cAMP and βArr pathways and compared to the marketed GLP-1R agonists exenatide and liraglutide. The potency of PF-06882961 (CS EC_50_ = 13 nM) on the cAMP pathway was approximately 120- and 14-fold (Fig 3A, Table S2) lower than exenatide (CS EC_50_ = 0.11 nM) and liraglutide (CS EC_50_ = 0.95 nM), respectively. The ability of PF-06882961 and marketed peptides to engage βArr was further assessed using PathHunter^®^ technology (Fig. 3B, Table S5). PF-06882961 was a partial agonist in recruiting βArr2 (EC_50_ = 490 nM, E_max_ = 36%), while exenatide (βArr2 EC_50_ = 9.0 nM, E_max_ = 75%) and liraglutide (βArr2 EC_50_ = 20 nM, E_max_ = 99%) were respectively 54- and 23-fold more potent with greater E_max_ values. βArr1 responses closely mirrored those of βArr2 (Table S5). Calculation of pathway bias (*31*) supports the assertion that, that relative to liraglutide, both PF-06882961 and exenatide have slight (∼5-fold) signaling bias towards the cAMP pathway relative to βArr recruitment (Fig. S3).

βArr recruitment at the GLP-1R leads to internalization of the receptor toward endosomal compartments, which has been proposed to impact receptor desensitization and signaling duration in preclinical models (*18, 19*). As a complementary approach to probing the βArr pathway, we quantified agonist-induced GLP-1R internalization using human embryonic kidney (HEK) 293 cells stably expressing a fluorogen-activated protein (FAP)-tagged version of the human GLP-1R (Fig. 3C). Initial experiments confirmed that PF-06882961 and peptide agonists retain similar rank order potency and full cAMP signaling in this HEK 293 model relative to the CS assay (Fig. S4). Under identical conditions, treatment with PF-06882961 led to FAP-GLP-1R internalization (EC_50_ = 230 nM, E_max_ = 83%). Exenatide (EC_50_ = 0.60 nM, E_max_ = 125%) and liraglutide (EC_50_ = 1.8 nM, E_max_ = 117%) were 380- and 130-fold more potent, respectively, and caused somewhat greater receptor internalization relative to PF-06882961 (Table S5). Pathway bias analysis using parameters derived from this cellular model again further supports that, relative to liraglutide, PF-06882961 has minor (∼3-fold) bias away from internalization (vs. cAMP). Finally, agonist-induced internalization and recycling of a green fluorescent protein (GFP)-tagged human GLP-1R construct expressed in HEK 293 cells was visualized using confocal microscopy (Fig. 3D). Consistent with the FAP-based approach, stimulation with PF-06882961 for 30 minutes triggered marked intracellular accumulation of GFP-GLP-1R, which was reversible following a 2-hour washout period.

To further define the pharmacological profile of PF-06882961, we sought to determine its binding affinity using radioligand binding assays. In competition experiments using [^125^I]-GLP-1 as the radiolabeled probe, the inhibition constant (K_i_) of PF-06882961 for the GLP-1R (K_i_ = 360 nM) (Fig. 3E) was 3900- and 82-fold lower than exenatide (K_i_ = 0.092 nM) and liraglutide (K_i_ = 4.4 nM), respectively (Table S6). However, given that large peptides like GLP-1 interact with both the extracellular and transmembrane domains of the GLP-1R (*32*), and that our small molecules were unlikely to recapitulate this complex binding mode, it was unclear whether the competition binding experiments with radiolabeled GLP-1 were providing an accurate measure of affinity (*25*). Therefore, we developed a novel radiolabeled small-molecule probe, [^3^H]-PF-06883365, which is expected to bind in the same pocket as PF-06882961 (Fig. S5, Table S6). The affinity of PF-06882961 (K_i_ = 80 nM) measured using this new radioligand was 4.5-fold more potent and more consistent with its cAMP potency, whereas the clinical peptides had similar affinities using both radioligands (Table S6).

### Tryptophan 33 in GLP-1R is required for PF-06882961 signaling

Prior to the selection of animal models for *in vivo* pharmacology studies, we characterized the potential for species differences in GLP-1R activation with our small-molecule agonists. PF-06882961 stimulated cAMP accumulation in CHO cells expressing both the human and monkey GLP-1Rs with comparable EC_50_ values (Fig. S6). In contrast, PF-06882961 did not increase cAMP levels in cells expressing the mouse, rat, or rabbit GLP-1R. Consistent with this finding, no improvement in glucose tolerance was observed during an intraperitoneal glucose tolerance test in C57BL6 mice that were administered a subcutaneous dose of PF-06882961 (10 mg/kg) (Fig. 4A). Comparison of the GLP-1R sequences in human and monkey versus other species revealed the notable presence of a tryptophan (W) at position 33 of the primate GLP-1R. In contrast, other species including the mouse, rat, and rabbit GLP-1R contain a serine (S) residue at position 33. Supporting the crucial role of W33, PF-06882961 increased cAMP accumulation in cells expressing the S33W mutant of mouse GLP-1R, whereas it failed to induce signaling at the human W33S receptor (Fig. 4B).

**Fig. 4.**
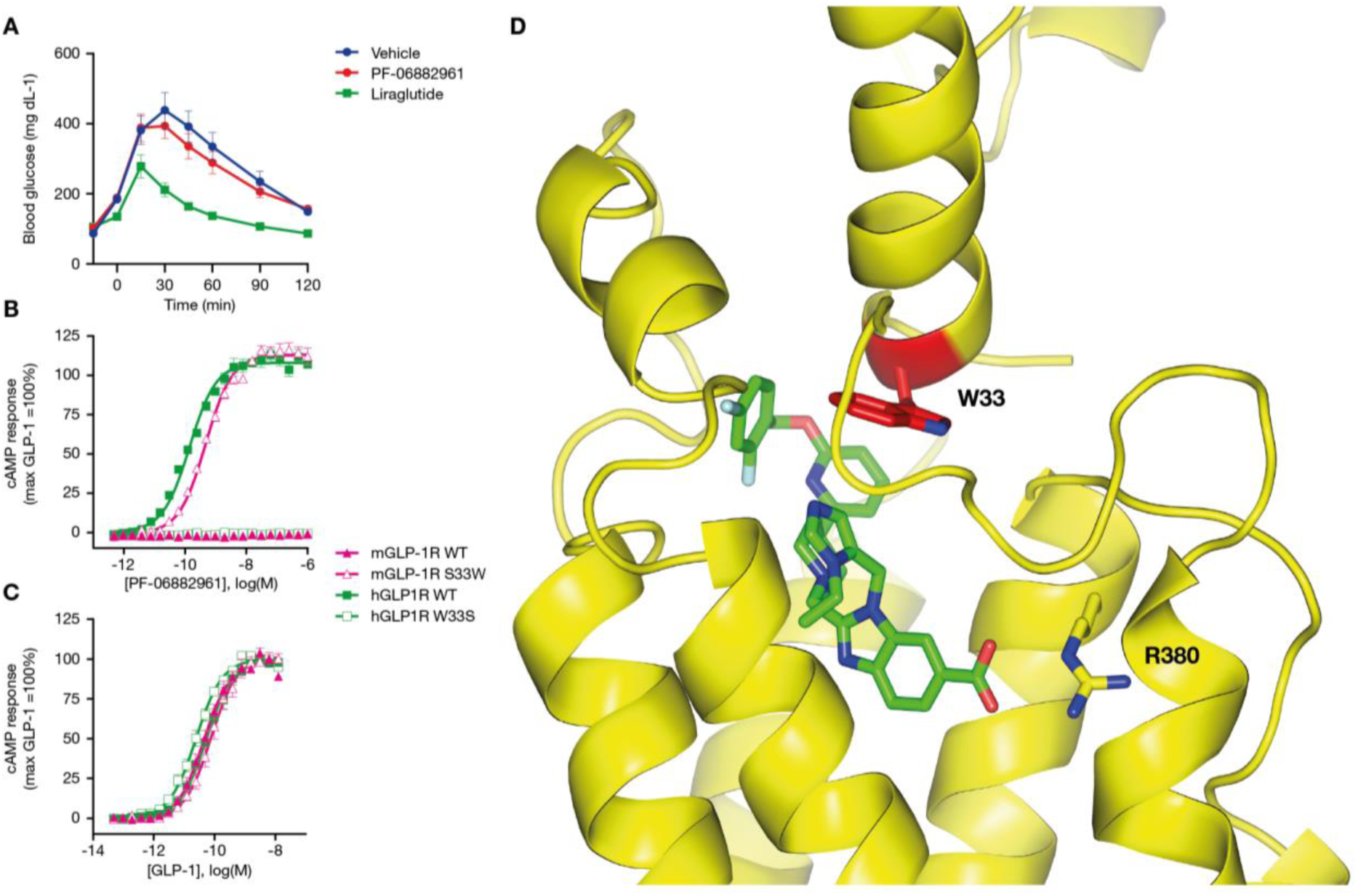
Tryptophan 33 is critical for the function of small-molecule GLP-1R agonists. (**A**) In contrast to liraglutide, PF-06882961 does not reduce glucose AUC during an intraperitoneal glucose tolerance test in C57BL/6 mice. (**B, C**) In contrast to GLP-1, PF-06882961 promotes cAMP production in GLP-1R-expressing cells only when residue 33 is tryptophan (W), not serine (S). (**B**) PF-06882961 signals in CHO cells expressing human-GLP-1R (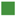), but not the mouse-GLP-1R (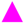). PF-06882961 signaling is restored in CHO cells expressing mouse-GLP-1R S33W (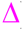) and is negated in human-GLP-1R W33S (□). (**C**) GLP-1 promotes signaling in mouse and human wild-type and mutant constructs. (**D**) Cryo-EM structure of PF-06883365 (green) bound to human GLP-1R. W33 closes the top of the small-molecule binding pocket. Arginine 380 (R380) interacts with the carboxylic substituent of the small molecule agonist. Helix 4 was removed from the figure for clarity. Data represent the mean ± SEM. AUC, area under the curve; cAMP, cyclic adenosine monophosphate; CHO, Chinese hamster ovary; Cryo-EM, cryogenic electron microscope; GLP-1R, glucagon-like peptide-1 receptor; h, human; m, mouse; SEM, standard error of the mean.

In contrast, cAMP signaling in response to GLP-1 was comparable between wild-type and mutant receptors, supporting the hypothesis that altered signaling with PF-06882961 was not due to a marked alteration of surface expression for the mutated constructs (Fig. 4C). The importance of W33 in small-molecule activation of GLP-1R was curious since it is located on the extracellular domain (ECD), distal from the transmembrane domains and connecting loops directly involved in peptide-induced GPCR activation (*20, 32*). Overall, our findings were reminiscent of a previous report describing the 100-fold reduction in the binding affinity of the small-molecule GLP-1R antagonist T-0632 in the W33S mutant of human GLP-1R (*33*). It was postulated that the binding of T-0632 stabilizes a closed, inactive conformation of GLP-1R, which involves W33 (*34*).

Recent cryogenic electron microscope (cryo-EM) structures of the human and rabbit GLP-1Rs (bound to exendin-p5 and GLP-1, respectively) indicate that residue 33 does not interact with peptide agonists but extends towards solvent (Fig. S7) (*32, 35*). Moreover, the species selectivity of a GLP-1R monoclonal antibody (Fab 3F52) was attributed to a binding epitope containing W33 (*36, 37*), which further supports the hypothesis that W33 is solvent-exposed. To better understand the role of W33 in small-molecule agonist binding to GLP-1R, we generated the cryo-EM structure of a PF-06882961 analog (PF-06883365) bound to human GLP-1R (Fig. 4D). In this structure, the GLP-1R ECD has rotated slightly relative to the peptide-bound structures, and W33 has moved ∼14 Å, closing the top of the small-molecule binding pocket. Consistent with the potency gained from introducing the 6-carboxylic acid motif in the benzimidazole region, the cryo-EM structure showed that the carboxylic acid of PF-06883365 is making a productive interaction with an arginine residue at position 380 in the GLP-1R binding pocket.

### PF-06882961 is orally bioavailable and efficacious for lowering glucose and food intake in monkeys

The pharmacokinetics of PF-06882961 were examined in rats and monkeys after intravenous (IV) and oral administration (Fig 5A, Table S7). The oral bioavailability of PF-06882961 in animals was low to moderate and increased in a dose-dependent manner, which was adequate for studying preclinical *in vivo* efficacy and safety of PF-06882961 delivered via the oral route in standard formulations.

**Fig. 5.**
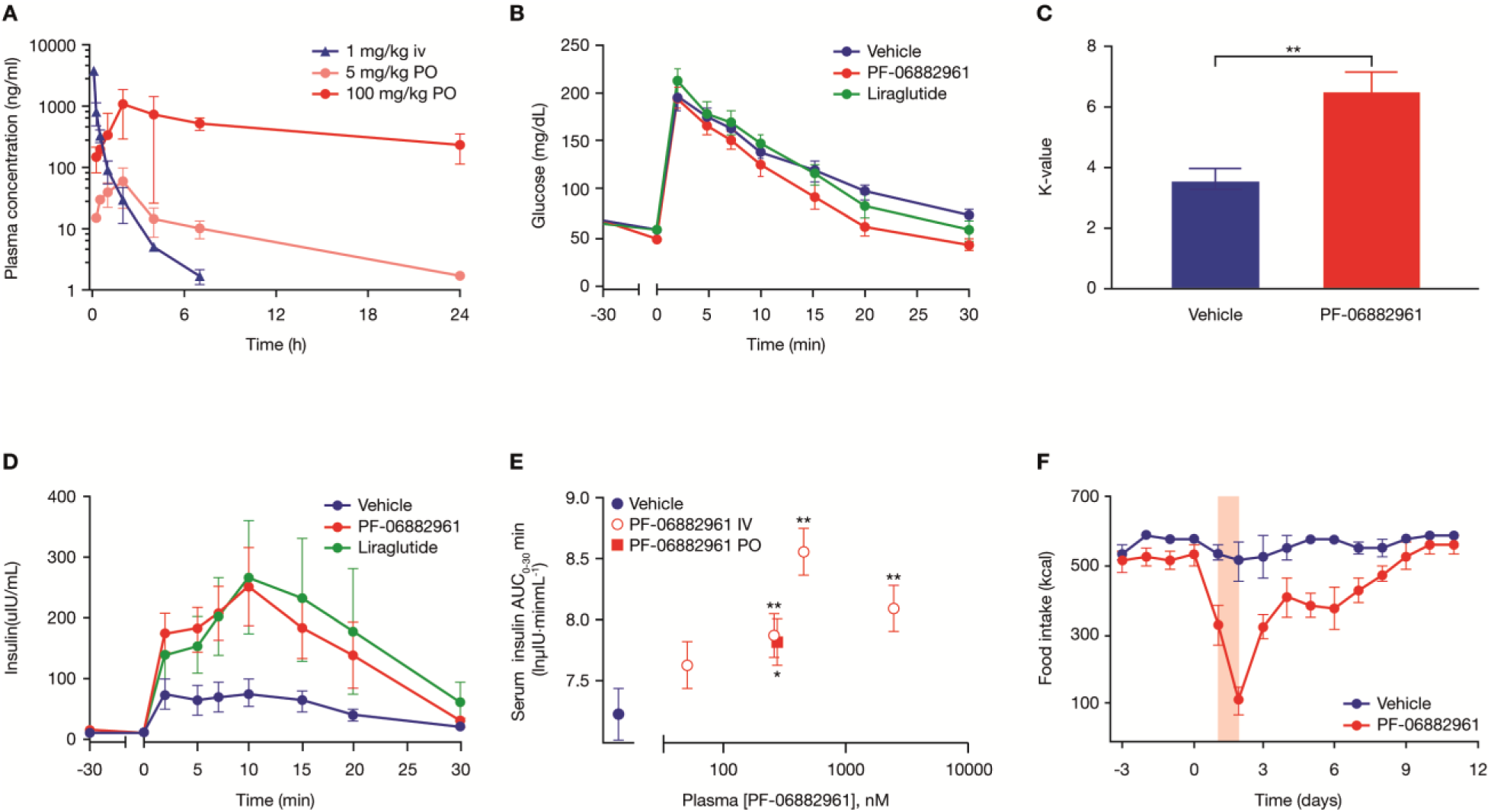
PF-06882961 potentiates glucose-stimulated insulin release and reduces food intake in monkeys. (**A**) PF-06882961 concentrations after IV or PO dosing in monkeys (n=2 each). (**B– E**) PF-06882961 increased the rate of glucose disappearance and enhanced insulin secretion during an IVGTT (250 mg/kg 50% dextrose) in monkeys (n=8 each). Serum glucose (**B**), k-value (**C**), and serum insulin (**D**) during IVGTT when PF-06882961 was IV infused to 3.0 µM (55 nM unbound) serum levels; liraglutide was administered by subcutaneous injection to achieve 58 nM (0.31 nM unbound) serum levels. (**E**) IV and PO administration of PF-06882961 potentiated glucose-stimulated insulin release (AUC_0–30 min_) in an exposure-proportional manner during an IVGTT. **(F**) Food intake in monkeys treated with either vehicle or PF-06882961 (n=6 each). All values are presented as mean ± SEM. *P<0.05 and **P<0.01 AUC, area under the curve; IV, intravenous; IVGTT, intravenous glucose tolerance test; PO, oral; SEM, standard error of the mean.

Since PF-06882961 does not activate the rodent GLP-1R, the therapeutic effects of PF-06882961 on insulin and glucose were examined in an IV glucose tolerance test (IVGTT) in cynomolgus monkeys. Intravenous infusion of PF-06882961 during the IVGTT led to an increase in insulin secretion and the rate of glucose disappearance (K-value) (Fig. 5B–D). Enhancement of glucose-stimulated insulin secretion by PF-06882961 was concentration-dependent and was also observed following oral dosing with similar efficacy when compared to administration by IV infusion (Fig. 5E). Once-daily administration of PF-06882961 for 2 days also inhibited food intake compared to vehicle-treated monkeys (Fig. 5F).

### Oral administration of PF-06882961 shows evidence of glucose-lowering in healthy human study participants

PF-06882961 was selected as a candidate for clinical studies based on its *in vitro* and *in vivo* pharmacologic and disposition profile, including potent agonism of the GLP-1R, preclinical disposition attributes (e.g., low metabolic CL_int_ in human hepatocytes), good safety margins versus the hERG channel (IC_50_ = 4.3 µM, Table S4) and broad panel screening (Table S8), and selectivity versus related class B GPCRs (Table S9). Moreover, adequate safety margins were observed in repeat-dose rat and monkey toxicology studies, which supported advancing PF-06882961 to human clinical studies.

The safety, tolerability, and pharmacokinetics of PF-06882961 were evaluated in a first-in-human, Phase 1, randomized, double-blind, placebo-controlled, single ascending dose study in healthy adult participants. A total of 25 participants in three cohorts were randomized to receive study treatment. Data from the dose-escalation portion of the study (cohorts 1 and 2) are presented in this manuscript; in this portion of the study, 17 participants received tablet formulations of PF-06882961 or matching placebo at single doses ranging from 3 to 300 mg. Following oral administration under fasted conditions, PF-06882961 was generally well-tolerated. There were no serious or severe adverse events (AEs) reported, nor discontinuations due to AEs (Table S10). Most AEs were mild in severity, and a higher proportion of participants reported an AE following administration of PF-06882961 at the 300 mg dose level, compared with other study treatments (Table S10). The most common AEs recorded following administration of PF-06882961 were nausea, vomiting, and decreased appetite, all of which were considered treatment-related by the investigator and consistent with the expected effects of the GLP-1R agonist mechanism. Plasma exposure of PF-06882961, as measured by AUC_inf_ and C_max_, appeared to increase in an approximately dose-proportional manner, with mean t_1/2_ ranging from 4.3 to 6.1 hours (Fig. 6 and Table S11). Median time (T_max_) to maximal concentration (C_max_) values ranged from 2.0 to 6.0 hours post-dose. A tablet formulation of PF-06882961 was also administered under fed conditions at a dose of 100 mg. Assessment of the effect of food on PF-06882961 administration revealed similar exposure (as measured by AUC_inf_) and t_1/2_ when administered in the fed state, compared with the fasted state (Table S11), indicating that PF-06882961 can be dosed in both the fed and fasted states.

**Fig. 6.**
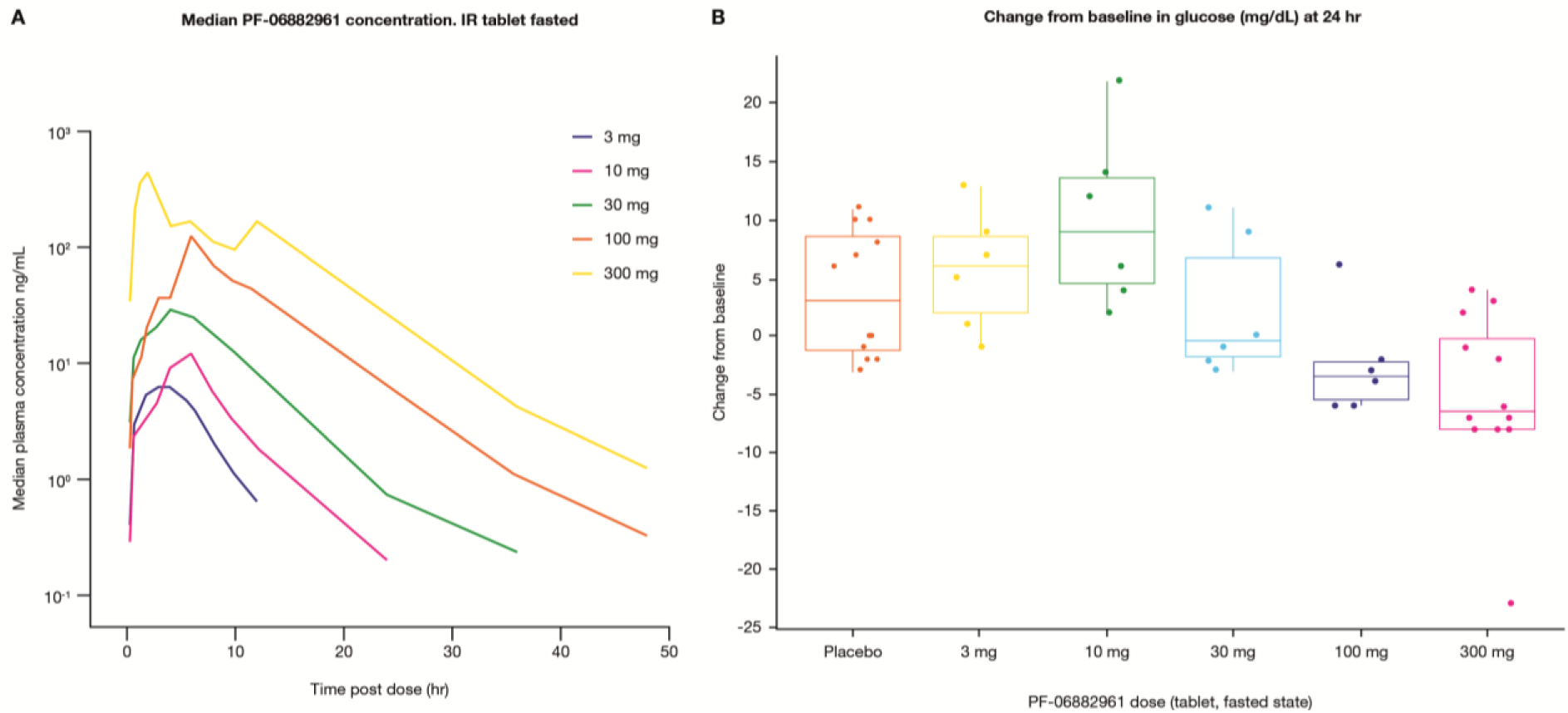
PF-06882961 lowers fasting serum glucose in healthy human volunteers. (**A**) Median plasma PF-06882961 concentration–time profiles after single-dose oral administration of PF-06882961 (3–300 mg) to humans (n = 6/dose, except n = 12 in 300 mg group) in the fasted state. Plasma exposure increased in an approximately dose-proportional manner, as assessed by dose-normalized geometric mean C_max_ and AUC_inf_, with a mean half-life (t_1/2_) ranging from 4.3 to 6.1 hours (Table S11). The median time to maximal concentration (T_max_) values ranged from 2.0 to 6.0 hours post-dose. (**B**) Fasting serum glucose was measured at pre-dose (baseline) and 24 hours post-dose during each dosing period. The subject-level changes from baselines by treatment group for the tablet administrations of PF-06882961 and placebo in the fasted state are presented along with boxplots representing the medians and inter-quartile ranges. In exploratory post-hoc statistical analyses, the 300 mg dose showed a statistical difference from placebo (P < 0.05, using paired t-tests unadjusted for multiple testing). AUC, area under the curve; C_max_, maximum plasma concentration; CI, confidence interval; IR, immediate release; SD, standard deviation; T_max_, time of the first occurrence of C_max_.

The primary and secondary endpoints of the study were safety and pharmacokinetic parameters, and fasting serum glucose was measured pre-dose and post-dose in study participants, all of whom had glucose and glycated hemoglobin (HbA1c) levels within the normal reference range for the laboratory. After the administration of single doses of PF-06882961, exploratory analyses revealed a trend for declining post-dose glucose levels at higher doses, with an apparent dose-related trend observed at 24 hours post dose (Fig. 6B). All post-dose glucose levels remained within the normal reference range for the laboratory, and no adverse events of hypoglycemia were reported.

In summary, we developed a novel sensitized HTS assay that identified a series of small molecule GLP-1R agonists. The series was optimized for pharmacologic potency, safety, and disposition attributes amenable for use in humans. PF-06882961 demonstrates a balanced *in vitro* signaling profile, potentiates glucose-stimulated insulin release, and decreases food intake in monkeys, and is orally available in healthy human participants. To our knowledge, this is the first literature report on glucose-lowering with an oral small-molecule agonist of the GLP-1R in humans.

## Acknowledgments

The authors would like to thank the participants of the FIH study; T. P. Rolph, S. Liras and M. J. Birnbaum for continued support of the oral GLP-1R agonist program; M. E. Flanagan for sharing expertise in medicinal chemistry; G. E. Aspnes, S. B. Coffey, E. L. Conn, A. Dion, M. S. Dowling, G. Ingle, W. Jiao, A. Shavnya, and L. Wei for compound synthesis and route design; J. N. Bradow, J. K. Smith, and X. Wang for analytical support; J. X. Kong, L. Rogers, J. J. Shah, and K. A. Stevens for the execution of *in vitro* activity assays; K. Schildknegt for overseeing radiosynthesis; B. L. Bernardo, S. Joaquim, N. Nammi, and A. H. Smith for execution of *in vivo* testing; D. Gates and K. N. Yip for compound formulation; H. Eng for pharmacokinetics assay execution; K. F. Fennell for protein reagent generation; D. N. Gorman and A. Bannerjee for statistical support; S. Farenden and E. Madigan for coordinating pharmaceutical science activities; M. Popovitz, M. C. Sanford, and S. Uppal for clinical study execution; M. J. Vorko for project management; X. Qiu, J. Pandit and A. H. Varghese for supporting the Pfizer membrane protein CryoEM initiative; and Sosei Heptares for generating a stabilized GLP-1 receptor construct using their proprietary StaR^®^ technology to enable our structural studies. This study was sponsored by Pfizer Inc. Medical writing support, under the direction of the authors, was provided by Eric Comeau, Ph.D., of CMC Connect, McCann Health Medical Communications and was funded by Pfizer Inc., New York, NY, USA, in accordance with Good Publication Practice (GPP3) guidelines (Ann Intern Med 2015;163:461–464).

## Funding

This study was sponsored by Pfizer Inc.

## Author contributions

Formulation or evolution of overarching research goals and aims: DAG, ASK, DJE, JBK, PML, VMJ, JPF

Data curation: JBK, DRD, PML, JPF

Application of statistical and/or mathematical analyses: ARS, JBK, DRD, CB, JPF Acquisition of funding: VMJ

Conducting the research and/or collecting data: DAG, ASK, AMM, CL, SWB, ARS, SH, DJE, MCG, JBK, DRD, JMD, VMJ, JMC, CB, RBR, DJL, JPF, YL, DAT, MSL

Project administration: MSL, DAP

Provision of study resources including animals and patients: VMJ, MSL

Implementation of the relevant software or computer programs: CB

Oversight and leadership of the study planning and execution: DAG, ARS, SH, DJE, JBK, PML, JMD, VMJ, CB, DJL, JPF, MSL

Data verification and validation: ASK, SWB, ARS, DWP, MCG, JBK, DRD, PML, CB, JPF, DAT, MSL

Preparation of the data and graphical elements of the paper: DAG, ASK, CL, DWP, DJE, JBK, DRD, PML, JMD, CB, JPF, YL, DAT

Preparation of the written elements of the paper: DAG, ASK, CL, ARS, DWP, DJE, JBK, DRD, PML, JMD, JPF, YL

Editing and reviewing of the paper: DAG, ASK, AMM, CL, SWB, ARS, DWP, SH, DJE, MCG, JBK, DRD. PML, JMD, CB, RBR, DJL, JMC, JPF, YL, MSL, DAP

## Competing interests

ASK, AMM, MCG, JMD, CB, CL, SWB, DJL, PML, DRD, JMC, JPF, YL, ARS, DAT, DWP, SH, RBR, MSL, and DAG are employees and stockholders of Pfizer Inc. DAP, DJE, JBK and VMJ are stockholders of Pfizer Inc.

## Data and materials availability

Upon request, and subject to certain criteria, conditions, and exceptions (see https://www.pfizer.com/science/clinical-trials/trial-data-and-results for more information), Pfizer will provide access to individual de-identified participant data from Pfizer-sponsored global interventional clinical studies conducted for medicines, vaccines and medical devices (1) for indications that have been approved in the US and/or EU or (2) in programs that have been terminated (i.e., development for all indications has been discontinued). Pfizer will also consider requests for the protocol, data dictionary, and statistical analysis plan. Data may be requested from Pfizer trials 24 months after study completion. The de-identified participant data will be made available to researchers whose proposals meet the research criteria and other conditions, and for which an exception does not apply, via a secure portal. To gain access, data requestors must enter into a data access agreement with Pfizer. Pfizer shares compounds using requests via the ‘compound transfer program’ (see https://www.pfizer.com/science/collaboration/compound-transfer-program).

## Supplementary Materials

Materials and Methods

Figures S1–S7

Tables S1–S11

References (*38-54*)

