## supplemental data for "A small-molecule oral agonist of the human glucagon-like peptide-1 receptor"

**This PDF file includes:**

Materials and Methods
Figs. S1 to S7
Tables S1 to S11
References 38 to 54

**Abbreviations**

AcOH: acetic acid

BINAP: (2,2′-bis(diphenylphosphino)-1,1′-binaphthyl)

Bip: Biphenylalanine

dba: dibenzylidine acetone

DCM: dichloromethane, $CH_2Cl_2$

DCE: 1,2-dichloroethane

DMF: dimethylformamide

DMSO: dimethyl sulfoxide

DPPP: 1,3-Bis(diphenylphosphino)propane

DVB: divinylbenzene

EDTA: ethylenediaminetetraacetic acid

EGTA: ethylene glycol-bis(β-aminoethyl ether)-N,N,N′,N′-tetraacetic acid

ESI-MS: electrospray ionization mass spectrometry

EtOAc: ethyl acetate

EtOH: ethanol

$Et_2O$: diethyl ether

$Et_3N$: triethylamine

Fmoc: Fluorenylmethyl-oxycarbonyl

GFP: green fluorescent protein

HATU: 1-[Bis(dimethylamino)methylene]-1H-1,2,3-triazolo[4,5-b]pyridinium 3-oxide hexafluorophosphate

HBTU**:** *N,N,N′,N′*-Tetramethyl-*O*-(1*H*-benzotriazol-1-yl)uronium hexafluorophosphate

$iPr_2NEt$: *N*,*N*-diisopropylethyl amine, Hunig's base

KOtBu: potassium tert-butoxide

KOTMS: potassium trimethylsilanolate

LiHMDS: lithium bis(trimethylsilyl) amide

mCPBA: 3-chloroperbenzoic acid

MeCN: acetonitrile

MeOH: methanol

MTBE: methyl tertbutyl ether

NaBu: sodium butyrate

NaOtBu: sodium tert-butoxide

NMM: *N*-Methylmorpholine

Nva: norvaline

PE: petroleum ether

PEI: polyethyleneimine

$PhCH_3$: toluene

pTSA: p-toluenesulfonic acid

PyAOP: (7-Azabenzotriazol-1-yloxy) tripyrrolidino-phosphonium hexafluorophosphate

Rink amide MBHA resin: 4-(2′,4′-Dimethoxyphenyl-Fmoc-aminomethyl)-phenoxyacetamido-4-methylbenzhydrylamine resin

RBF: round bottom flask

SPPS: solid phase peptide synthesis

STAB: sodium triacetoxyborohydride

TBD: 1,5,7-triazabicyclo[4.4.0]dec-5-ene

TFA: trifluoroacetic acid

THF: tetrahydrofuran

TIS: *tri*-isopropylsilane

TMSCl: chloro trimethylsilane

Tris: tris(hydroxymethyl)aminomethane

T3P: propylphosphonic anhydride

UV: ultra-violet

### Materials and Methods

Cell Line Generation

The human glucagon-like peptide-1 (GLP-1) receptor (GLP-1R, reference sequence NM_002062) cDNA was amplified by RT-PCR and subcloned into pcDNA3.1 and pcDNA5/FRT/TO (Thermo Fisher, Pittsburgh, PA). CHO-K1 cells (ATCC) were transfected with pcDNA3.1/ GLP-1R and a clonal cell line with high (‘screening cell line’) receptor levels was selected. The pcDNA5/FRT/TO/ GLP-1R plasmid was transfected into the Flp-In™-CHO Cell Line (Thermo Fisher, Pittsburgh, PA) and clonal cell line with low (‘candidate selection cell line’) receptor levels was selected. High and low expression cell lines, selected based on receptor mRNA expression levels (qPCR) and cAMP responses to GLP-1, were utilized for lead identification and structure activity relationship (SAR) efforts, as indicated. A similar procedure was used to generate CHO cells stably expressing the mouse (NM_021332.2) and cynomolgus (NM_001287663.1) GLP-1R.

Compound Syntheses

**General considerations**: Unless otherwise stated, all reactants, reagents and solvents were obtained from commercial sources and used without further purification. Data for $^{1}H$ nuclear magnetic resonance (NMR) spectra are reported relative to residual solvent signals (for $CDCl_3$, δH = 7.27 ppm; for DMSO-$d_6$, δH = 2.50 ppm; for $CD_3OD$, δH = 3.31 ppm) as follows: chemical shift (δ ppm), multiplicity, coupling constant (Hz) and integration. The multiplicities are denoted as follows: s, singlet; d, doublet; t, triplet; q, quartet; spt, septet; m, multiplet; br s, broad singlet. Data for $^{13}C$ NMR spectra are reported in terms of chemical shift (δ ppm) relative to residual solvent signals (for $CDCl_3$, δC = 77.0 ppm; for DMSO-$d_6$, δC = 39.5 ppm; for $CD_3OD$, δC = 49.2 ppm). Flash chromatography was carried out on either a Biotage SP purification system or a Combiflash Companion from Teledyne Isco; Biotage SNAP, KPsil or Redisep Rf silica columns were used. Except where otherwise noted, all reactions were run under an inert atmosphere of nitrogen gas using anhydrous solvents at room temperature (~23 °C). The terms “concentrated” and “evaporated” refer to the removal of solvent at reduced pressure on a rotary evaporator with a water bath temperature not exceeding 60 °C.

### Synthesis of Peptide 1

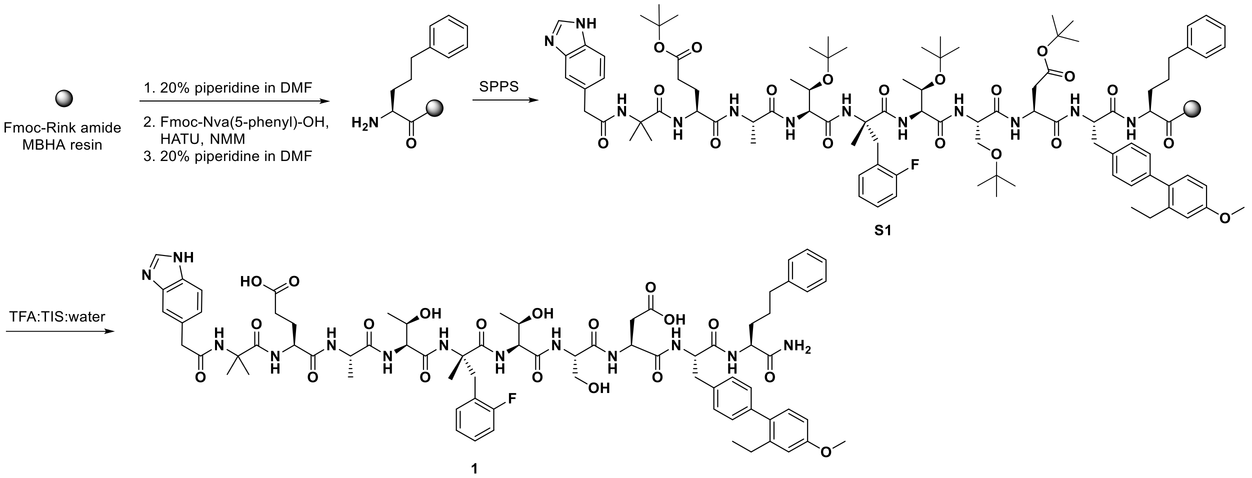


**Sources of unnatural amino acids and 2-(1*H*-benzo[*d*]imidazole-5-yl)acetic acid.** (*S*)-2-(((9*H*-fluoren-9-yl)methoxy)carbonylamino)-5-phenylpentanoic acid (Fmoc-Nva(5-phenyl)-OH) was purchased from PepTech Corporation (USA); (*S*)-*N*-Fmoc-α-methyl-2-fluoro-phenylalanine (Fmoc-[α-Me-Phe(2-F)-OH) was purchased from Nagase & Co. Ltd. (Japan); (*S*)-2-(((9*H*-fluoren-9-yl)methoxy)carbonyl-amino)-3-(2'-ethyl-4'-methoxybiphenyl-4-yl)propanoic acid (Fmoc-Bip(2'-Et,4'-OMe)-OH) was synthesized as previously described (*38*); 2-(1*H*-benzo[*d*]imidazole-5-yl)acetic acid was available from a variety of commercial sources.

**Source of Fmoc-Rink amide MBHA resin and natural Fmoc-amino acids**. Fmoc-Rink amide 4-methylbenzhydrylamine hydrochloride, polymer-bound (MBHA) resin (100-200 mesh, 1% divinyl benzene (DVB) cross linking) was purchased from Tianjin Nankai Hecheng S&T Co., Ltd, and the natural Fmoc-amino acids are readily available from a variety of commercial sources.

**Procedure A** (Standard on-resin amide coupling using 1.9 eq of 1-[bis(dimethylamino)methylene]-1*H*-1,2,3-triazolo[4,5-*b*]pyridinium 3-oxid hexafluorophosphate (HATU) and Fmoc-deprotection). 2 eq of Fmoc-Xaa, 1.9 eq of HATU, and 4 eq of N-methylmorpholine (NMM) in dimethylformamide (DMF) were added to the resin. The reaction was agitated under nitrogen until the amidation went to completion based on the Kaiser ninhydrin test. After the reaction was deemed complete, the suspension was filtered, and the resin was washed with DMF (5 x 3 resin volumes). Then, 20% piperidine in DMF (3 resin volumes) was added to the resin bound peptide. The suspension was kept at room temperature for 0.5 h while a stream of nitrogen was bubbled through it. After 0.5 h, the suspension was filtered, and the resin was washed with DMF (5 x 3 resin volumes).

**Procedure B** (Standard on-resin amide coupling with 2-(1*H*-benzotriazol-1-yl)-1,1,3,3-tetramethyluronium hexafluorophosphate (HBTU) and Fmoc-deprotection): 3 eq of Fmoc-Xaa, 2.9 eq of HBTU, and 6 eq of NMM in DMF (1.2 resin volumes) were added to the resin. The reaction was agitated under nitrogen until the amidation went to completion based on the Kaiser ninhydrin test. After the reaction was deemed complete, the suspension was filtered, and the resin was washed with DMF (5 x 3 resin volumes). Then, 20% piperidine in DMF (3 resin volumes) was added to the resin bound peptide. The suspension was kept at room temperature for 0.5 h while a stream of nitrogen was bubbled through it. After 0.5 h, the suspension was filtered, and the resin was washed with DMF (5 x 3 resin volumes).

**Procedure C** (Standard on-resin amide coupling using 2.9 eq of HATU and Fmoc-deprotection). 3 eq of Fmoc-Xaa, 2.9 eq of HATU, and 6 eq of NMM in DMF were added to the resin. The reaction was agitated under nitrogen until the amidation went to completion based on the Kaiser ninhydrin test. After the reaction was deemed complete, the suspension was filtered, and the resin was washed with DMF (5 x 3 resin volumes). Then, 20% piperidine in DMF (3 resin volumes) was added to the resin bound peptide. The suspension was kept at room temperature for 0.5 h while a stream of nitrogen was bubbled through it. After 0.5 h, the suspension was filtered, and the resin was washed with DMF (5 x 3 resin volumes).

**Procedure D** (Standard amide coupling involving 2-(1*H*-benzo[*d*]imidazole-5-yl)acetic acid). 3 eq of the carboxylic acid, 3 eq of ((7-azabenzotriazol-1-yloxy)tripyrrolidinophosphonium hexafluorophosphate) (PyAOP), 5 eq of N,N-diisopropylethylamine ($iPr_2NEt$) in DMF (1.2 resin volumes) were added to the resin. After the reaction was deemed complete by mass detection, the suspension was filtered, and the resin was washed with DMF (5 x 3 resin volumes).

**Purification of peptide 1**. The peptide was purified using a Waters 4000 system connected to Waters Delta-Pak-C18 reversed phase column, 25 x 200 mm, 15 micron, 100 Å) eluting with a gradient of water (0.1% trifluoroacetic acid (TFA)):acetonitrile (MeCN)/water (4/1, 0.1% TFA) [52:48 to 22:78] over 60 min at a flow rate of 60 mL/min.

**Purity analysis of peptide 1**. The pure peptide was analyzed using a HP1090 system coupled to a Phenomenox C18 reversed phase column (4.6 x 150 mm, 5 micron, 100 Å) eluting with a gradient of water (0.1% TFA):MeCN/water (4/1, 0.1% TFA) [48:52 to 38:62] over 20 minutes, flow rate = 1.0 mL/min].

**Mass spectrometry analysis of peptide 1**. A Thermo-LCQ Advantage system was used to collect data based on electrospray ionization (ESI).

Fmoc-Rink amide MBHA resin (0.48 g, 0.15 mmol) was swelled in DMF for 2 h, and then filtered. 20% piperidine in DMF (3 resin volumes) was added into the reaction vessel to deliver a suspension. While a stream of nitrogen was bubbled through it, the suspension was kept at room temperature for 30 min to achieve complete Fmoc cleavage. Then, the suspension was filtered, and the resin was washed with DMF (2 x 5 mL). Fmoc-(*S*)-2-amino-5-phenylpentanoic acid (0.125 g, 0.3 mmol), HATU (0.108 g, 0.285 mmol), NMM (0.066 mL, 0.6 mmol) and DMF (~1.2 resin volumes) were added to the resin. The amidation was carried out under a nitrogen atmosphere and deemed complete via the Kaiser ninhydrin test. Upon completion, the suspension was filtered, and the resin was washed with DMF (2 x 5 mL) to deliver Fmoc-(*S*)-2-amino-5-phenylpentanoyl Rink amide resin. 20% piperidine in DMF (3 resin volumes) was added to the resin bound peptide, and the suspension was kept at room temperature for 0.5 h while a stream of nitrogen was bubbled through it. After 0.5 h, the suspension was filtered, and the resin was washed with DMF (5 x 3 resin volumes) to furnish the following (*S*)-2-amino-5-phenylpentanoyl Rink amide resin.

The following nine Fmoc-amino acids and 2-(1*H*-benzo[*d*]imidazole-5-yl)acetic acid were sequentially coupled to the peptidyl resin using the stated procedures: Fmoc-Bip(2'-Et,4'-OMe)-OH (procedure A), Fmoc-Asp(O*t*-Bu)-OH (procedure B), Fmoc-Ser(*t*-Bu)-OH (procedure B), Fmoc-Thr(*t*-Bu)-OH (procedure B), Fmoc-[α-Me-Phe(2-F)]-OH (procedure A), Fmoc-Thr(*t*-Bu)-OH (procedure C), Fmoc-Ala-OH (procedure C), Fmoc-Glu(O*t*-Bu)-OH (procedure C), Fmoc-Aib-OH (procedure C), and 2-(1*H*-benzo[*d*]imidazole-5-yl)acetic acid (procedure D). After the last amide coupling, the peptidyl resin **A** was washed with DMF (3 × 10 mL), dichloromethane (DCM) (3 × 10 mL) and MeOH (3 × 10 mL). The resin was dried *in vacuo* overnight. To the peptidyl resin (0.76 g) was added a mixture of TFA:triisopropylsilane(TIS):water (95:2.5:2.5, 10 mL). The mixture was shaken for 3 h and filtered. Ether (80 mL) was then added to the filtrate to give a precipitate. The mixture was centrifuged, and the supernatant was decanted. The resulting solid peptide pellet was washed with ether (3 x 30 mL) and dried *in vacuo* overnight to give crude peptide **1** (0.248 g). The crude peptide was purified by reversed phase high performance liquid chromatography (HPLC) as described in the purification section. Like fractions were combined and lyophilized to deliver 35 mg (15% overall yield) of peptide **1** (TFA salt) as a white solid. UV purity (220 nm) = 95.8 % (retention time = 10.19 min, see purity analysis section for conditions); MS (ES) mass-to-charge ratio *(m/z)* 1500.9 $(M+H)^+$, 751.3 $(M/2+H)^+$.

**Synthesis of Compound 2**

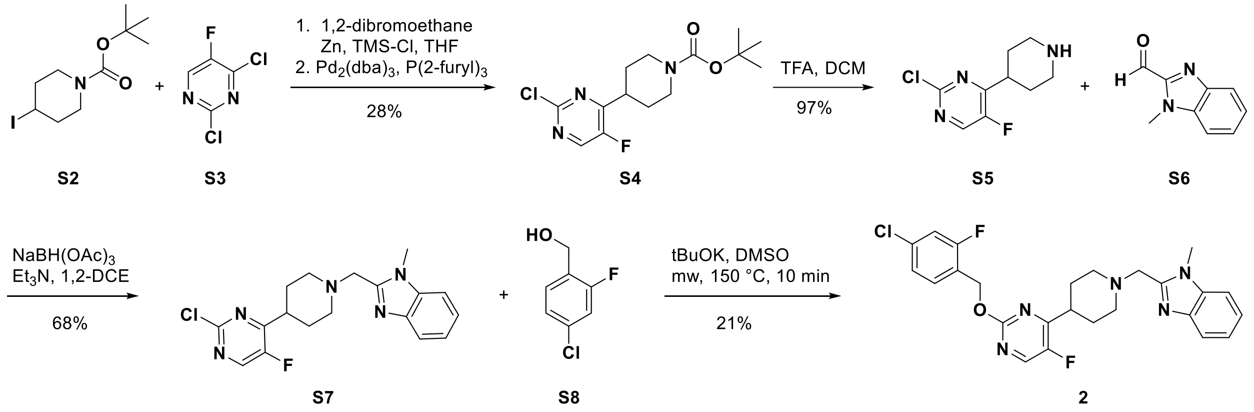


*4-(2-Chloro-5-fluoro-pyrimidin-4-yl)-piperidine-1-carboxylic acid tert-butyl ester (**S4**).* A suspension of zinc dust (63.2 g, 0.970 mol) in tetrahydrofuran (THF) (200 mL) was heated to 35 °C, and then 1,2-dibromoethane (17.4 g, 92.8 mmol) was added and allowed to exotherm (65 °C). Once the reaction mixture cooled to 30 °C, trimethylchlorosilane (TMSCl) (8.90 g, 81.9 mmol) was added (slight exotherm). After 30 min, 4-iodo-piperidine-1-carboxylic acid *tert*-butyl ester (**S2**, 229.1 g, 0.736 mol) in THF (450 mL) was added dropwise and then stirred for 30 min. Then, *tri*-2-furylphosphine (10 g, 43 mmol) and tris(dibenzylideneacetone)dipalladium(0) ($Pd_2(dba)_3$) (11.2 g, 12.2 mmol) were added. After 10 min, 2,4-dichloro-5-fluoro-pyrimidine (**S3**, 102.4 g, 0.613 mol) in THF (250 mL) was added in one portion, and the reaction mixture was heated to 70 °C for 3 h. After cooling to room temperature, the reaction was diluted with ethyl acetate (EtOAc) (0.5 L) and water (0.5 L), filtered through Celite and transferred to a separatory funnel. The organic layer was separated, dried ($Na_2SO_4$), filtered, and concentrated under reduced pressure. The resulting residue was purified by flash chromatography (silica gel) eluting with a gradient of heptanes:EtOAc (70:30 to 50:50) to deliver 55 g (28%) of 4-(2-chloro-5-fluoro-pyrimidin-4-yl)-piperidine-1-carboxylic acid *tert*-butyl ester (**S4**) as a white solid. $^1$H NMR (400 MHz, $CDCl_3$) δ 8.38 (d, $J_{H\text{-}F}$ = 1.2 Hz, 1H), 4.28 (br s, 2H), 3.20–3.13 (m, 1H), 2.85 (br s, 2H), 1.91–1.79 (m, 4H), 1.48 (s, 9H); MS (ES) *m/z* 260.1 (M-56 + H)$^+$, 262.0 (M-56 + H)$^+$.

*2-Chloro-5-fluoro-4-(piperidin-4-yl)pyrimidine (**S5**, TFA salt)*. To a solution of *tert*-butyl 4-(2-chloro-5-fluoropyrimidin-4-yl)piperidine-1-carboxylate (**S4**, 480 mg, 1.52 mmol) in DCM (3 mL) was added TFA (1.05 mL). The mixture was stirred at room temperature for 1 h. The mixture was concentrated under reduced pressure to deliver 489 mg (97%) 2-chloro-5-fluoro-4-(piperidin-4-yl)pyrimidine (**S5**, TFA salt) as a brown oil which was used in the next step without further purification. $^1$H NMR (400 MHz, $CD_3OD$) δ 8.63 (d, $J_{H\text{-}F}$ = 1.2 Hz, 1H), 3.57–3.46 (m, 3H), 3.25–3.18 (m, 2H), 2.15–2.09 (m, 4H); $^{19}$F (376 MHz, $CD_3OD$) δ -77.0, -144.0; MS (atmospheric pressure (AP)) ionization *m/z* 216.1 (M+H)$^+$, 218.1 (M+H)$^+$.

*2-((4-(2-Chloro-5-fluoropyrimidin-4-yl)piperidin-1-yl)methyl)-1-methyl-1H-benzo[d]imidazole (**S7**)*. A solution of the TFA salt of 2-chloro-5-fluoro-4-(piperidin-4-yl)pyrimidine (**S5**, 486 mg, 1.47 mmol) in 1,2-dichloroethane (DCE) (3 mL) was added 1-methyl-1*H*-benzo[*d*]imidazole-2-carbaldehyde (**S6**. 237 mg, 1.48 mmol), $Et_3N$ (1.19 g, 11.8 mmol) and $MgSO_4$ (192 mg, 1.60 mmol). The mixture was stirred at room temperature for 1 h, then sodium triacetoxyborohydride ($NaBH(OAc)_3$) (470 mg, 2.22 mmol) was added. The mixture was stirred at 30 °C for 16 h. The mixture was diluted with DCM (20 mL) and washed with 1N NaOH (13 mL). The layers were separated, and the organic layer was washed with brine, dried ($Na_2SO_4$), filtered, and concentrated under reduced pressure. The resulting residue was purified by flash chromatography (silica gel) eluting with a gradient of petroleum ether (PE):EtOAc (100:0 to 35:65) to afford 360 mg (68%) of 2-((4-(2-chloro-5-fluoropyrimidin-4-yl)piperidin-1-yl)methyl)-1-methyl-1*H*-benzo[*d*]imidazole (**S7**) as a gum. $^1$H NMR (400 MHz, $CDCl_3$) δ 8.36 (s, 1H), 7.76 (d, *J* = 7.8 Hz, 1H), 7.39–7.30 (m, 3H), 3.93 (s, 3H), 3.90 (s, 2H), 3.09–3.02 (m, 3H), 2.36–2.30 (m, 2H), 2.04–1.94 (m, 2H), 1.84–1.81 (m, 2H); MS (ES) *m/z* 359.8 (M+H)$^+$, 361.9 (M+H)$^+$.

*2-((4-(2-((4-Chloro-2-fluorobenzyl)oxy)-5-fluoropyrimidin-4-yl)piperidin-1-yl)methyl)-1-methyl-1H-benzo[d]-imidazole (**2**).* To a solution of (4-chloro-2-fluorophenyl)methanol (**S8**, 64 mg, 0.40 mmol) in dimethyl sulfoxide (DMSO) (1 mL) was added 2-((4-(2-chloro-5-fluoropyrimidin-4-yl)piperidin-1-yl)methyl)-1-methyl-1*H*-benzo[*d*]imidazole (**S7**, 120 mg, 0.333 mmol) and *t*-BuOK (35 mg, 0.31 mmol). The reaction mixture was irradiated in a microwave at 100 °C for 10 min. The mixture was then purified by preparatory HPLC to deliver 50 mg (21%) of the bis-TFA salt of 2-((4-(2-((4-chloro-2-fluorobenzyl)oxy)-5-fluoropyrimidin-4-yl)piperidin-1-yl)methyl)-1-methyl-1*H*-benzo[*d*]-imidazole (**2**) as a gum. $^1$H NMR (400 MHz, $CD_3OD$) δ 8.45 (d, $J_{H\text{-}F}$ = 1.6 Hz, 1H), 7.78–7.73 (m, 2H), 7.56–7.46 (m, 3H), 7.27–7.22 (m, 2H), 5.46 (s, 2H),

4.55 (s, 2H), 4.02 (s, 3H), 3.60–3.57 (m, 2H), 3.36–3.32 (m, 1H), 3.10–3.04 (m, 2H), 2.27–2.16 (m, 2H), 2.08–2.04 (m, 2H); $^{13}$C NMR (100 MHz, $CD_3OD$, $^1$H and $^{19}$F decoupled) δ 162.7, 162.6, 162.4, 162.0, 159.1, 154.1, 149.0, 147.8, 136.4, 136.3, 132.9, 126.3, 126.0, 125.9, 124.1, 118.1, 117.3, 116.2, 112.6, 64.3, 54.6, 53.1, 36.4, 31.9, 29.2; $^{13}$C NMR (100 MHz, $CD_3OD$, $^1$H decoupled, just $J_{sp^2 C(ipso)\text{-}F}$ reported) δ 162.2 (d, *J* = 250.9 Hz), 153.9 (d, *J* = 251.6 Hz); $^{19}$F NMR (376 MHz, $CD_3OD$) δ -77.3, -117.3, -152.5; HRMS (ESI) *m/z* [M + H]$^+$ for $C_{25}H_{25}ClF_2N_5O$ calcd 484.1710, found 484.1693.

Preparatory HPLC method: Column: YMC-Actus Triart C18 (150 x 30 mm, 5 um); Gradient: water (0.1% TFA):MeCN (65:35 to 45:55 over 12 min); Flow rate: 30 mL/min.

Preparatory HPLC delivered 50 mg (21%) of the bis-TFA salt of 2-((4-(2-((4-chloro-2-fluorobenzyl)oxy)-5-fluoropyrimidin-4-yl)piperidin-1-yl)methyl)-1-methyl-1*H*-benzo[*d*]-imidazole **(2)** as a gum.

**Synthesis of Compound 3**

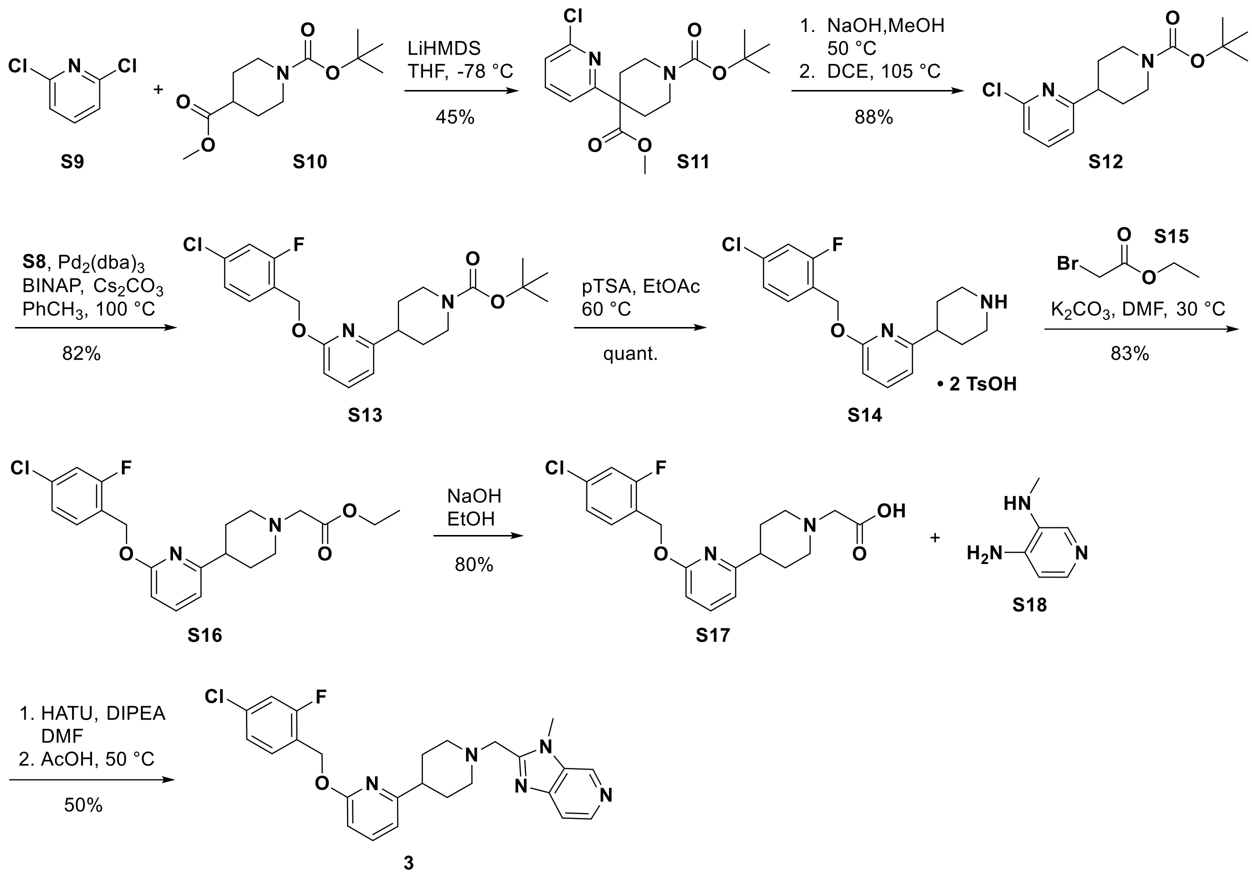


*1-(tert-Butyl) 4-methyl 4-(6-chloropyridin-2-yl)piperidine-1,4-dicarboxylate (**S11**)*. To a stirred solution of 1-(*tert*-butyl)-4-methyl piperdine-1,4-dicarboxylate (**S10**, 1.68 mL, 6.2 mmol) and 2,6-dichloropyridine (**S9**, 0.82 g, 5.5 mmol) in $PhCH_3$ (5.5 mL) was added lithium bis(trimethylsilyl)amide (LiHMDS) in THF (1M, 7.2 mL). After 15 h, the solution was acidified with HCl in water (0.5N, 15 mL). The aqueous phase was extracted with EtOAc (2 x 25 mL), the combined organic layers were dried over anhydrous $Na_2SO_4$, filtered, and evaporated under reduced pressure to give a colorless oil. The crude material was purified using column chromatography eluting with 20% EtOAc in heptane to obtain 1-(*tert*-butyl) 4-methyl 4-(6-chloropyridin-2-yl)piperidine-1,4-dicarboxylate (**S11**) as a colorless oil (0.87 g, 45%). $^1$H NMR (600 MHz, $CDCl_3$) δ 7.62 (t, *J* = 7.9 Hz, 1H), 7.21 (d, *J* = 7.6 Hz, 2H), 3.83 (br s, 2H), 3.71 (s, 3H), 3.14 (br s, 2H), 2.41 (d, *J* = 13.5 Hz, 2H), 2.08 (ddd, *J* = 13.6, 10.4, 3.5 Hz, 2H), 1.45 (s, 9H). MS (ES) *m/z* 255.2 $(M\text{-}Boc+H)^+$.

*tert-Butyl 4-(6-chloropyridin-2-yl)piperidine-1-carboxylate (**S12**).* To a stirred solution of 1-(*tert*-butyl) 4-methyl 4-(6-chloropyridin-2-yl)piperidine-1,4-dicarboxylate (**S11**, 0.87 g, 2.4 mmol) in MeOH (10 mL) was added NaOH in water (2 N, 6 mL) at 60 °C. After 5 h, the solution was allowed to cool to room temperature and acidified with HCl in water (1 N, 50 mL). The aqueous phase was extracted with EtOAc (3 x 75 mL), the combined organic layers were dried over anhydrous $Na_2SO_4$, filtered and evaporated under reduced pressure to give a white solid. The crude material was dissolved in DCE (24 mL) and stirred at reflux. After 2 h, the mixture was concentrated, treated with 25% water in MeOH (4 mL). After 4 h, the resultant solid was filtered and dried under reduced pressure to obtain *tert*-butyl 4-(6-chloropyridin-2-yl)piperidine-1-carboxylate (**S12**) as a white solid (0.64 g, 88%). $^1$H NMR (600 MHz, $CDCl_3$) δ 7.58 (t, *J* = 7.6 Hz, 1H), 7.17 (d, *J* = 8.2 Hz, 1H), 7.06 (d, *J* = 7.6 Hz, 1H), 4.25 (br s, 2H), 2.93–2.66 (m, 3H), 1.91 (d, *J* = 12.9 Hz, 2H), 1.69 (qd, *J* = 12.5, 4.1 Hz, 2H), 1.47 (s, 9H). MS (ES) *m/z* 241.2 $(M\text{-}tBu)^+$.

*4-(6-((4-Chloro-2-fluorobenzyl)oxy)pyridin-2-yl)piperidine-1-carboxylate (**S13**).* A reaction vessel equipped with a reflux condenser was charged with *tert*-butyl 4-(6-chloropyridin-2-yl)piperidine-1-carboxylate (**S12**, 6.5 g, 21.9 mmol), (4-chloro-2-fluorophenyl)methanol (3.5 g, 21.9 mmol), $Pd_2(dba)_3$ (1.0 g, 1.1 mmol), 2,2′-bis(diphenylphosphino)-1,1′-binaphthyl (BINAP) (1.4 g, 2.2 mmol) and cesium carbonate ($Cs_2CO_3$) (14.3 g, 43.8 mmol). Toluene (73 mL) was added and the mixture was heated to 100 °C. After 16 h, the mixture was allowed to cool to room temperature, filtered through Celite with EtOAc (100 mL) and concentrated under reduced pressure. The crude material was purified using column chromatography eluting with 10% EtOAc in PE to obtain *tert*-butyl 4-(6-((4-chloro-2-fluorobenzyl)oxy)pyridin-2-yl)piperidine-1-carboxylate (**S13**) as a yellow oil (7.6 g, 82%). $^1$H NMR (400 MHz, DMSO-$d_6$) δ 7.63 (t, *J* = 7.6 Hz, 1H), 7.54 (t, *J* = 8.2 Hz, 1H), 7.43 (dd, *J* = 1.8, 10.0 Hz, 1H), 7.29 (dd, *J* = 1.6, 8.2 Hz, 1H), 6.87 (d, *J* = 7.0 Hz, 1H), 6.68 (d, *J* = 8.2 Hz, 1H), 5.36 (s, 2H), 4.03 (br d, *J* = 11.7 Hz, 2H), 2.7–2.9 (m, 3H), 1.76 (br d, *J* = 10.9 Hz, 2H), 1.55 (dq, *J* = 4.3, 12.5 Hz, 2H), 1.42 (s, 9H); HRMS (ESI-Orbitrap) *m/z* calcd for $C_{22}H_{27}ClFN_2O_3$ $[M + H]^+$ 421.1694, found 421.1689.

*2-((4-Chloro-2-fluorobenzyl)oxy)-6-(piperidin-4-yl)pyridine (**S14**).* To a stirred solution of *tert*-butyl 4-(6-((4-chloro-2-fluorobenzyl)oxy)pyridin-2-yl)piperidine-1-carboxylate (**S13**, 50 g, 120 mmol) in EtOAc (700 mL) was added p-toluenesulfonic acid (*p*TSA) monohydrate (59 g, 309 mmol). The mixture was heated to 60 °C. After 30 min, the solution was allowed to cool to room temperature, the resultant solid was slurried for 16 h, filtered and dried under reduced pressure to obtain 2-((4-chloro-2-fluorobenzyl)oxy)-6-(piperidin-4-yl)pyridine (**S14**) as a white solid (81.2 g, quant). $^1$H NMR (DMSO-$d_6$) δ: 11.75 (br s, 2H), 8.55 (br. s., 1H), 8.28 (d, *J* = 6.5 Hz, 1H), 7.68 (t, *J* = 7.6 Hz, 1H), 7.60 (t, *J* = 8.2 Hz, 1H), 7.48 (d, *J* = 8.2 Hz, 4H), 7.32 (d, *J* = 8.2 Hz, 1H), 7.12 (d, *J* = 7.6 Hz, 4H), 6.89 (d, *J* = 7.6 Hz, 1H), 6.74 (d, *J* = 8.2 Hz, 1H), 5.38 (s, 2H), 3.37 (d, *J* = 12.3 Hz, 2H), 3.09–2.98 (m, 2H), 2.96–2.87 (m, 1H), 2.29 (s, 6H), 2.01–1.96 (m, 2H), 1.94–1.80 (m, 2H); MS (ES) *m/z* 322.4 $(M+H)^+$.

*2-(4-(6-((4-Chloro-2-fluorobenzyl)oxy)pyridin-2-yl)piperidin-1-yl)acetic acid (**S16**).* To solution of 2-((4-chloro-2-fluorobenzyl)oxy)-6-(piperidin-4-yl)pyridine (**S14**, 70.0 g, 218 mmol) and $K_2CO_3$ (118 g, 853 mmol) in DMF (800 mL) was added ethyl 2-bromoacetate (**S15**, 39.9 g, 239 mmol) portion-wise. The mixture was stirred at 30 °C for 1 h. The mixture was diluted with water (500 mL) and extracted with EtOAc (3 x 400 mL). The combined organic layers were dried ($Na_2SO_4$), filtered and concentrated under reduced pressure. The crude product was purified by flash chromatography (silica gel) eluting with PE:EtOAc (10:1) to afford 2-(4-(6-((4-chloro-2-fluorobenzyl)oxy)pyridin-2-yl)piperidin-1-yl)acetic acid (**S16**) as a yellow oil (74 g, 83%). $^1$H NMR (400 MHz, $CDCl_3$) δ 7.53–7.44 (m, 2H), 7.15–7.10 (m, 2H), 6.75 (d, *J* = 7.4 Hz, 1H), 6.61 (d, *J* = 8.2 Hz, 1H), 5.42 (s, 2H), 4.22 (q, *J* = 7.0 Hz, 2H), 3.29 (s, 2H), 3.13–3.06 (m, 2H), 2.64–2.58 (m, 1H), 2.35 (br s, 2H), 2.08–1.90 (m, 4H), 1.30 (t, *J* = 7.0 Hz, 3H); MS (ES) *m/z* $(M+H)^+$, 406.8.

*2-(4-(6-((4-Chloro-2-fluorobenzyl)oxy)pyridin-2-yl)piperidin-1-yl)acetic acid (**S17**).* To a solution of **S16** (73 g, 0.179 mmol) in EtOH (270 mL) was added 5 N NaOH (156 mL, 780 mmol). The solution was stirred at 25 °C for 2 h. Then, the mixture was acidified to pH ~3.5 with 1 M HCl. The resulting precipitate was collected by filtration, washed with water, and dried *in vacuo* to afford 2-(4-(6-((4-chloro-2-fluorobenzyl)oxy)pyridin-2-yl)piperidin-1-yl)acetic acid (**S17**) as a pale yellow solid (54 g, 80%). $^1$H NMR (400 MHz, DMSO-$d_6$) δ 10.60 (br s, 1H), 7.70–7.62 (m, 2H), 7.47 (dd, *J* = 1.5, 9.7 Hz, 1H), 7.33 (dd, *J* = 1.5, 8.2 Hz, 1H), 6.92 (d, *J* = 7.4 Hz, 1H), 6.73 (d, *J* = 8.2 Hz, 1H), 5.40 (s, 2H), 4.13 (s, 2H), 3.61–3.58 (m, 2H), 3.27–3.21 (m, 2H), 2.92–2.87 (m, 1H), 2.19–2.03 (m, 4H); MS (AP) *m/z* $(M+H)^+$, 379.2 and 381.1.

*2-((4-(6-((4-Chloro-2-fluorobenzyl)oxy)pyridin-2-yl)piperidin-1-yl)methyl)-3-methyl-3H-imidazo[4,5-c]pyridine (**3**).* To a solution of 2-(4-(6-((4-chloro-2-fluorobenzyl)oxy)pyridin-2-yl)piperidin-1-yl)acetic acid (**S17**, 150 mg, 0.397 mmol) in DMF (3 mL) was added HATU (226 mg, 0.594 mmol). The mixture was stirred at 25 °C for 20 min. Then, $N^3$-methylpyridine-3,4-diamine (73.1 mg, 0.594 mmol) and N,N-diisopropylethylamine (*i*$Pr_2$NEt) (154 mg, 1.19 mmol) were added, and the mixture was stirred at 25 °C for 16 h. The mixture was diluted with EtOAc (20 mL) and washed with water (15 mL). The separated aqueous layer was extracted with EtOAc (2 x 20 mL). The combined organic layers were washed with saturated brine (2 x 15 mL), dried ($Na_2SO_4$), filtered and concentrated under reduced pressure to give 200 mg of a brown oil. AcOH (3 mL) was added, and the resulting solution was heated at 50 °C for 2 h. The mixture was diluted with DCM (20 mL) and washed with saturated aqueous $NaHCO_3$ (25 mL). The separated organic layer was dried ($Na_2SO_4$), filtered and concentrated under reduced pressure. The resulting residue was purified by preparatory HPLC to deliver 2-((4-(6-((4-chloro-2-fluorobenzyl)oxy)pyridin-2-yl)piperidin-1-yl)methyl)-3-methyl-3*H*-imidazo[4,5-*c*]pyridine (**3**, partial formate salt, 0.6 eq) as a white solid (96 mg, 50%). $^1$H NMR (400 MHz, $CD_3OD$) δ 8.93 (s, 1H), 8.37 (d, $J$ = 5.9 Hz, 1H), 8.21 (br s, 0.6H), 7.69 (d, $J$ = 5.5 Hz, 1H), 7.58 (dd, $J$ = 7.4, 8.2 Hz, 1H), 7.49 (app t, $J$ = 8 Hz , 1H), 7.22–7.16 (m, 2H), 6.83 (d, $J$ = 7.4 Hz, 1H), 6.64 (d, $J$ = 8.2 Hz, 1H), 5.40 (s, 2H), 4.09 (s, 3H), 4.01 (s, 2H), 3.10–3.07 (m, 2H), 2.71–2.63 (m, 1H), 2.44–2.37 (m, 2H), 1.92–1.87 (m, 4H); $^{13}$C NMR (100 MHz, $CD_3OD$) δ 164.1, 163.7, 162.2 (d, $J_{C-F}$ = 250.2 Hz), 157.5, 148.6, 141.9, 140.8, 135.6, 135.6 (d, $J_{C-F}$ =10.3 Hz), 134.1, 132.5 (d, $J_{C-F}$ = 5.1 Hz), 125.7 (d, $J_{C-F}$ = 3.7 Hz), 125.6 (d, $J_{C-F}$ = 14.7 Hz), 117 (d, $J_{C-F}$ =24.9 Hz), 115.6, 115.2, 109.5, 61.6 (d, $J_{C-F}$ = 4.4 Hz), 55.7, 55.3, 44.5, 32.6, 31.6; $^{19}$F NMR (376 MHz, $CD_3OD$) δ -117.6; HRMS (ESI-Orbitrap) *m/z* calcd for $C_{25}H_{26}ClFN_5O$ $[M + H]^+$ 466.1800, found 466.1786.

**Synthesis of Compound 4**

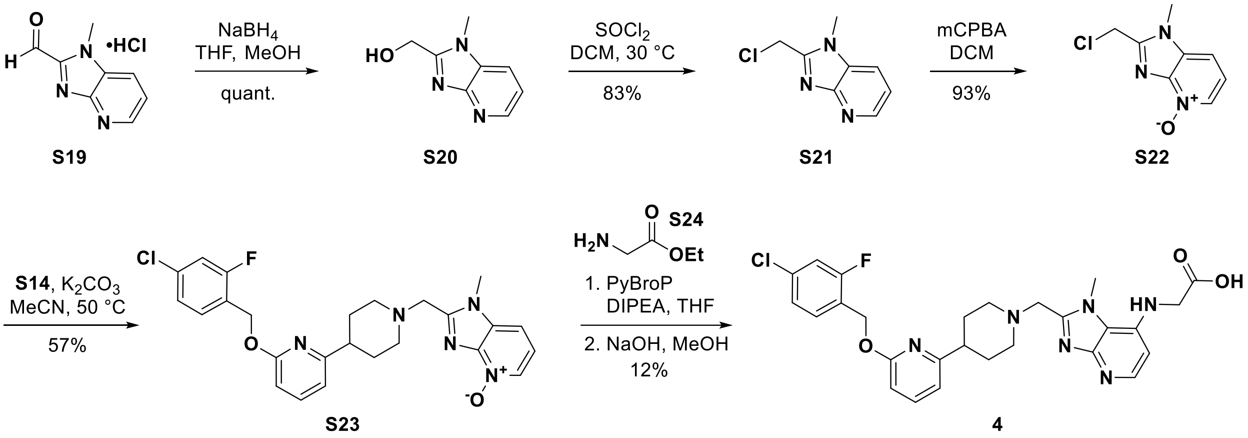


*(1-Methyl-1H-imidazo[4,5-b]pyridin-2-yl)methanol (**S20**)*. To a solution of 1-methyl-1*H*-imidazo[4,5-*b*]pyridine-2-carbaldehyde hydrochloride salt (**S19**, 2.5 g, 12.75 mmol) in mixture of anhydrous THF (20 mL) and MeOH (20 mL) was added sodium borohydride ($NaBH_4$) (1.93 g, 51.0 mmol), portion-wise. The resulting slurry was stirred at 25 °C for 2 h. The mixture was quenched by slow addition of 4 M HCl/dioxane (20 mL) in MeOH (20 mL). After quench, the tan slurry was allowed to stir at 25 °C for 1 h. The mixture was concentrated under reduced pressure to yield (1-methyl-1*H*-imidazo[4,5-*b*]pyridin-2-yl)methanol (**S20**) as a tan solid (2.05 g, quant.). $^1$H NMR (600 MHz, $CDCl_3$) δ 8.45 (d, $J$ = 4.4 Hz, 1H), 7.51 (d, $J$ = 7.9 Hz, 1H), 7.12 (dd, $J$ = 7.9, 4.7 Hz, 1H), 4.99 (s, 2H), 3.84 (s, 3H). MS (ES) *m/z* 164.7 $(M+H)^+$.

*2-(Chloromethyl)-1-methyl-1H-imidazo[4,5-b]pyridine (**S21**)*. To a suspension of (1-methyl-1*H*-imidazo[4,5-*b*]pyridin-2-yl)methanol (**S20**, 1.74 g, 10.7 mmol) in DCM ( 30 mL) was added thionyl chloride ($SOCl_2$) (3.9 mL, 54 mmol). The resulting reaction mixture was stirred at 30 °C for 24 h. The reaction mixture was concentrated under reduced pressure. The resulting brown solid was basified to pH 9 by slow addition of saturated aqueous $NaHCO_3$ and the mixture was extracted with DCM (3 x 70 mL). The combined organic phases were dried over $Na_2SO_4$, filtered and concentrated to yield 2-(chloromethyl)-1-methyl-1*H*-imidazo[4,5-*b*]pyridine (**S21**) as a yellow solid (1.61 g, 83%). $^1$H NMR (600 MHz, $CDCl_3$) δ 8.60 (d, *J* = 4.4 Hz, 1H), 7.75–7.67 (d, *J* = 0.79 Hz, 1H), 7.29–7.25 (m, 1H), 4.90 (s, 2H), 3.92 (s, 3H). MS (ES) *m/z* 182.0 $(M+H)^+$.

*2-(Chloromethyl)-1-methyl-1H-imidazo[4,5-b]pyridine 4-oxide (**S22**)*. To a solution of 2-(chloromethyl)-1-methyl-1*H*-imidazo[4,5-*b*]pyridine (**S21**, 1.38 g, 7.6 mmol) in DCM (20 mL) was added m-chloroperoxybenzoic acid (mCPBA) (3.41 g, 15.2 mmol, 77%). The resulting mixture was stirred at 25 °C under $N_2$ for 4 h. The reaction was treated with 4 M HCl/dioxane (1.9 mL, 7.7 mmol) in EtOAc (30 mL). The mixture was allowed to stir for 3 h to granulate. The solid was isolated by filtration, washed with heptane/$Et_2O$ to deliver 2-(chloromethyl)-1-methyl-1*H*-imidazo[4,5-*b*]pyridine 4-oxide (**S22**) as a pale yellow solid (1.39 g, 93% yield). $^1$H NMR (600 MHz, $CD_3OD$) δ 8.80 (d, *J* = 6.5 Hz, 1H), 8.54 (d, *J* = 8.2 Hz, 1H), 7.75 (t, *J* = 7.3 Hz, 1H), 5.12 (s, 2H), 4.11 (s, 3H). MS (ES) *m/z* 198.5 $(M+H)^+$.

*2-((4-(6-((4-Chloro-2-fluorobenzyl)oxy)pyridin-2-yl)piperidin-1-yl)methyl)-1-methyl-1H-imidazo[4,5-b]pyridine 4-oxide (**S23**)*. To solution of 2-((4-chloro-2-fluorobenzyl)oxy)-6-(piperidin-4-yl)pyridine trifluoroacetate (**S14-TFA salt**, 1.28 g, 3.99 mmol) and 2-(chloromethyl)-1-methyl-1*H*-imidazo[4,5-*b*]pyridine 4-oxide (**S22**, 933 mg, 3.99 mmol) in MeCN (20 mL) was added $K_2CO_3$ (1.65 g, 12 mmol) in one portion. The mixture was stirred at 50 °C for 2 h. The mixture was cooled to room temperature and diluted with DCM (20 mL). The precipitate was removed by filtration and rinsed with DCM (40 mL). The filtrate was concentrated under reduced pressure. The crude product was purified by flash chromatography (silica gel) eluting with MeOH:DCM (0 – 15% gradient) to afford 2-((4-(6-((4-chloro-2-fluorobenzyl)oxy)pyridin-2-yl)piperidin-1-yl)methyl)-1-methyl-1*H*-imidazo[4,5-*b*]pyridine 4-oxide (**S23**) a light brown glass (1.04 g, 57%). $^1$H NMR (400 MHz, $CD_3CN$) δ 8.14–7.97 (d, *J* = 6.2 Hz 1H), 7.57 (t, *J* = 7.6 Hz, 1H), 7.51 (t, *J* = 8.2 Hz, 1H), 7.44 (d, *J* = 8.2 Hz, 1H), 7.27–7.12 (m, 3H), 6.81 (d, *J* = 7.4 Hz, 1H), 6.61 (d, *J* = 8.2 Hz, 1H), 5.40 (s, 2H), 3.91 (s, 3H), 3.84 (s, 2H), 2.96 (br d, *J* = 11.7 Hz, 2H), 2.61 (tt, *J* = 10.5, 5.3 Hz, 1H), 2.25 (br d, *J* = 3.9 Hz, 2H), 2.10 (s, 1H), 1.86–1.72 (m, 4H). MS (ES) *m/z* 482.4 $(M+H)^+$.

*(2-((4-(6-((4-Chloro-2-fluorobenzyl)oxy)pyridin-2-yl)piperidin-1-yl)methyl)-1-methyl-1H-imidazo[4,5-b]pyridin-7-yl)glycine (**4**)*. To a solution of 2-((4-(6-((4-chloro-2-fluorobenzyl)oxy)pyridin-2-yl)piperidin-1-yl)methyl)-1-methyl-1*H*-imidazo[4,5-*b*]pyridine 4-

oxide (**S23**, 41 mg, 0.09 mmol) in THF (2 mL) was added $iPr_2NEt$ (0.06 mL, 0.594 mmol) and glycine ethyl ester (15 mg, 0.11 mmol) followed by bromo-tris-pyrrolidino-phosphonium hexafluorophosphate (PyBrop) (52 mg, 0.11 mmol). The mixture was stirred at 25 °C overnight. The mixture was diluted with water (30 mL) and extracted with DCM (3 x 60 mL). The combined organic layers were washed with saturated brine (2 x 30 mL), dried over $Na_2SO_4$, filtered and concentrated under reduced pressure to give 60 mg of a brown oil. MeOH (3 mL) was added followed by 1N NaOH (0.26 mL, 0.26 mmol), and the resulting solution was stirred at 25 °C overnight. The reaction mixture was concentrated, diluted with water (5 mL) and acidified to pH 3–4 with 1N HCl at 0 °C. The resulting precipitate was collected by filtration, rinsed with water (5 mL), and dried by vacuum to yield 35 mg solid. The solid was purified by preparatory HPLC to deliver 3.5 mg (12%) of (2-((4-(6-((4-chloro-2-fluorobenzyl)oxy)pyridin-2-yl)piperidin-1-yl)methyl)-1-methyl-1*H*-imidazo[4,5-*b*]pyridin-7-yl)glycine (***4***) as a gum as the mono-TFA salt. $^1$H NMR (400 MHz, $CD_3OD$) δ 8.16 (d, *J* = 7.0 Hz, 1H), 7.66 (t, *J* = 7.8 Hz, 1H), 7.51 (t, *J* = 8.2 Hz, 1H), 7.29–7.17 (m, 2H), 6.93 (d, *J* = 7.4 Hz, 1H), 6.79 (d, *J* = 7.0 Hz, 1H), 6.74 (d, *J* = 8.2 Hz, 1H), 5.44 (s, 2H), 4.87 (s, 2H), 4.37 (s, 2H), 4.22 (s, 3H), 3.93 (br s, 2H), 3.53–3.36 (m, 2H), 3.14–2.94 (m, 1H), 2.39–2.12 (m, 4H). $^{13}$C NMR (100 MHz, $CD_3OD$) δ 171.8, 164.2, 162.5 (q, $J_{C\text{-}F}$ = 34.8 Hz), 161.9 (d, $J_{C\text{-}F}$ = 250.2 Hz), 160.90, 150.1, 149.8, 147.5, 141.1, 137.6, 135.6 (d, $J_{C\text{-}F}$ =10.3 Hz), 132.2 (d, $J_{C\text{-}F}$ = 5.1 Hz), 125.6 (d, $J_{C\text{-}F}$ = 3.7 Hz), 125.1 (d, $J_{C\text{-}F}$ = 14.7Hz), 118.6, 118.2 (q, $J_{C\text{-}F}$ = 292.0 Hz), 116.9 (d, $J_{C\text{-}F}$ =24.9 Hz), 115.9, 110.2, 101.9, 61.5, (d, $J_{C\text{-}F}$ = 4.4 Hz), 54.9, 52.3, 45.2, 41.5, 34.1, 29.8; $^{19}$F NMR (376 MHz, $CD_3OD$) δ -77.0, -117.7. HRMS (ESI-Orbitrap) *m/z* calcd for $C_{27}H_{29}ClFN_6O_3$ $[M + H]^+$ 539.1968, found 539.1949.

Preparatory HPLC method: Column: Sunfire C18 (19x100 mm, 5 um); Gradient: water (0.225% trifluoroacetic acid):MeCN (95 : 60); Flow rate: 23.75 mL/min.

**Synthesis of Compound 5**

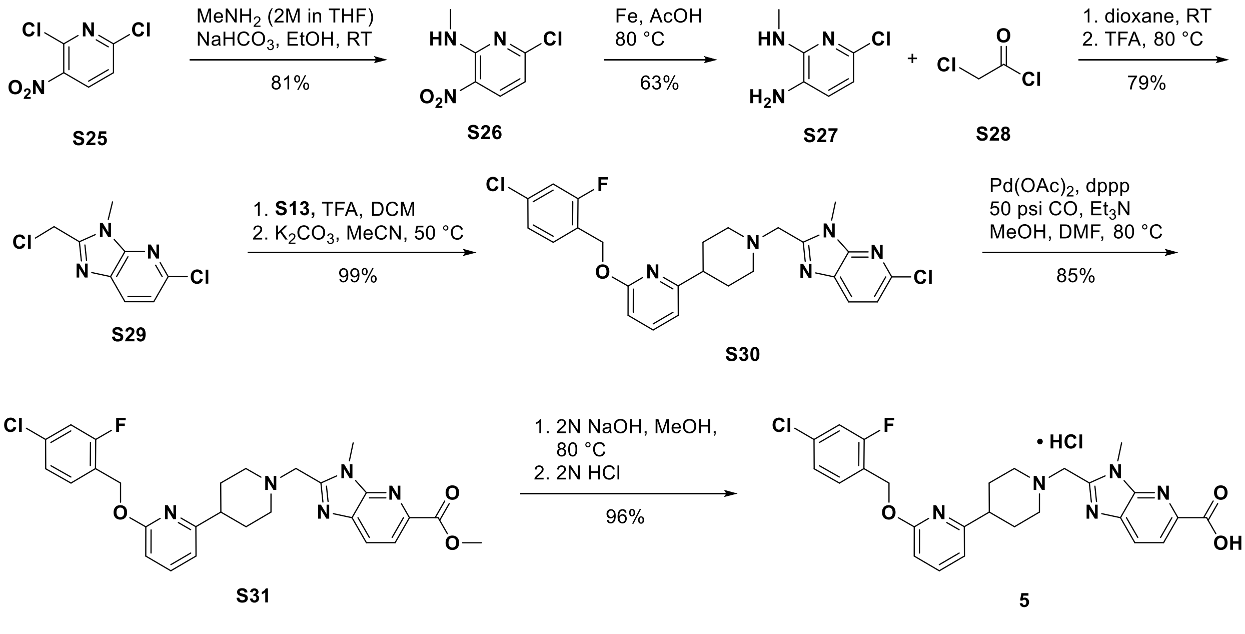

*6-Chloro-N-methyl-3-nitropyridin-2-amine (**S26**)*. To a suspension of 2,6-dichloro-3-nitropyridine (200 g, 1.04 mol) and $Na_2CO_3$ (132 g, 1.24 mol) in EtOH (1 L) was added 2.0 M methylamine ($MeNH_2$) in THF (622 mL,1.24 mol) dropwise at 0 °C via syringe. After the addition, the reaction mixture was stirred at 18 °C for 6 h. The yellow mixture was filtered and the filtrate was concentrated to give a yellow solid. The crude product was purified by flash chromatography (PE/EtOAc 0–5%) to afford 6-chloro-*N*-methyl-3-nitropyridin-2-amine (**S26**, 158 g, 81.3% yield) as a yellow solid. $^1H$ NMR (400 MHz, DMSO-$d_6$) δ 8.72 (br s, 1H), 8.41 (d, $J$ = 8.5 Hz, 1H), 6.76 (d, $J$ = 8.5 Hz, 1H), 3.00 (d, $J$ = 5.0 Hz, 3H); MS (ES) *m/z* 188.0 $(M+H)^+$.

*3-Amino-6-chloro-2-methylaminopyridine (**S27**)*. To a mixture of 6-chloro-*N*-methyl-3-nitropyridin-2-amine (**S26**, 15.8 g, 84.2 mmol) in AcOH (100 mL) was added iron powder (15.4 g, 276 mmol). The yellow mixture was stirred at 80 °C for 3 h. The reaction was cooled to room temperature and filtered. The filter cake was washed with EtOAc (2 x 100). The filtrate was concentrated and the crude product was purified by flash chromatography (120 g silica gel, 50% EtOAc/PE) to afford 3-amino-6-chloro-2-methylaminopyridine (**S27**, 8.40 g, 63.3% yield) as a brown solid. $^1H$ NMR (400 MHz, $CDCl_3$) δ 6.80 (d, $J$ = 7.7 Hz, 1H), 6.50 (d, $J$ = 7.5 Hz, 1H), 3.39 (br s, 2H), 3.01 (s, 3H); MS (ES) *m/z* 157.6 $(M+H)^+$.

*5-Chloro-2-(chloromethyl)-3-methyl-3H-imidazo[4,5-b]pyridine (**S29**)*. To a solution of 3-amino-6-chloro-2-methylaminopyridine (**S27**, 50.0 g, 317 mmol) in dioxane (1.2 L) was added chloroacetyl chloride (**S28**, 55.5 mL, 698 mmol) and the mixture was stirred at 15 °C for 50 min. The brown mixture was concentrated to give a brown solid which was taken up in TFA (1.2 L) and stirred at 80 °C for 60 h. The mixture was concentrated to give a brown oil. The oil was diluted with EtOAc (1 L) and neutralized with saturated aqueous sodium bicarbonate ($NaHCO_3$). When $CO_2$ evolution subsided, the layers were separated and the aqueous layer was extracted with EtOAc (200 mL). The organic extracts were combined, dried over $Na_2SO_4$, filtered and concentrated. The crude product was purified by flash chromatography (10–25% EtOAc/PE gradient) to afford 5-chloro-2-(chloromethyl)-3-methyl-3*H*-imidazo[4,5-*b*]pyridine (**S29**, 61.0 g, 79.3%) as a yellow solid. $^1H$ NMR (400 MHz, DMSO-$d_6$) δ 8.13 (d, $J$ = 8.3 Hz, 1H), 7.37 (d, $J$ = 8.5 Hz, 1H), 5.11 (s, 2H), 3.84 (s, 3H). MS (ES) *m/z* 180.9 $(M-Cl)^+$.

*5-Chloro-2-((4-(6-((4-chloro-2-fluorobenzyl)oxy)pyridin-2-yl)piperidin-1-yl)methyl)-3-methyl-3H-imidazo[4,5-b]pyridine (**S30**)*. To a yellow solution of *tert*-butyl 4-(6-((4-chloro-2-fluorobenzyl)oxy)pyridin-2-yl)piperidine-1-carboxylate (**S13**, 49.0 g, 119 mmol) in DCM (80 mL) was added TFA (80 mL) and the resultant yellow solution was stirred at 20 °C for 2 h. The yellow solution was concentrated and the resultant yellow oil (**S14, TFA salt)** was taken up in MeCN (300 mL) and treated with 5-chloro-2-(chloromethyl)-3-methyl-3*H*-imidazo[4,5-*b*]pyridine (**S29**, 25.9 g, 120 mmol) and $K_2CO_3$ (98.5 g, 713 mmol). The mixture was stirred at 50 °C for 16 h. The yellow mixture was poured into water (300 mL) and extracted with EtOAc (3 x 500 mL). The combined organic layers were washed with brine (500 mL), dried over $MgSO_4$, filtered and concentrated. The crude product was purified by flash chromatography (MeOH, DCM 0–5% gradient) to afford to afford 5-chloro-2-((4-(6-((4-chloro-2-fluorobenzyl)oxy)pyridin-2-yl)piperidin-1-yl)methyl)-3-methyl-3*H*-imidazo[4,5-*b*]pyridine (**S30**, 59.0 g, 99%) as a yellow solid. $^1$H NMR (400 MHz, $CDCl_3$) δ 7.93 (d, *J* = 8.0 Hz, 1H), 7.50 (t, *J* = 7.8 Hz, 1H), 7.44 (t, *J* = 8.3 Hz, 1H), 7.22 (d, *J* = 8.0 Hz, 1H), 7.1–7.1 (m, 2H), 6.73 (d, *J* = 7.0 Hz, 1H), 6.61 (d, *J* = 8.0 Hz, 1H), 5.41 (s, 2H), 3.99 (s, 3H), 3.85 (s, 2H), 2.98 (br d, *J* = 11.0 Hz, 2H), 2.7–2.5 (m, 1H), 2.4–2.2 (m, 2H), 2.0–1.8 (m, 4H); MS(ES) *m/z* 501.3 $(M+H)^+$.

*Methyl 2-((4-(6-((4-chloro-2-fluorobenzyl)oxy)pyridin-2-yl)piperidin-1-yl)methyl)-3-methyl-3H-imidazo[4,5-b]pyridine-5-carboxylate (**S31**).* A yellow solution of 5-chloro-2-((4-(6-((4-chloro-2-fluorobenzyl)oxy)pyridin-2-yl)piperidin-1-yl)methyl)-3-methyl-3*H*-imidazo[4,5-*b*]pyridine (**S30**, 59.0 g, 118 mmol), 1,3-bis(diphenylphosphino)propane (dppp) (6.80 g, 16.5 mmol), $Pd(OAc)_2$ (3.65 g, 16.3 mmol) and $Et_3N$ (125 g, 1240 mmol) in MeOH (800 mL) and DMF (100 mL) was stirred at 80 °C under a CO atmosphere of 50 psi for 16 h. The orange solution was concentrated to a brown oil. The brown oil was diluted with EtOAc (300 mL) and washed with water (200 mL). The organic layer was washed with brine (2 x 200 mL), dried over $MgSO_4$, filtered and concentrated. The crude product was combined with an identical 11 g scale reaction and the combined material was purified by flash chromatography (50–100% EtOAc/PE gradient) to afford methyl 2-((4-(6-((4-chloro-2-fluorobenzyl)oxy)pyridin-2-yl)piperidin-1-yl)methyl)-3-methyl-3*H*-imidazo[4,5-*b*]pyridine-5-carboxylate (**S31**, 62.6 g, 85%) as a pale yellow solid. $^1$H NMR (400 MHz, $CDCl_3$) δ 8.13 (d, *J* = 8.0 Hz, 1H), 8.07 (d, *J* = 8.0 Hz, 1H), 7.50 (t, *J* = 8.0 Hz, 1H), 7.44 (t, *J* = 8.0 Hz, 1H), 7.13–7.10 (m, 2H), 6.73 (d, *J* = 7.5 Hz, 1H), 6.61 (d, *J* = 8.0 Hz, 1H), 5.41 (s, 2H), 4.09 (s, 3H), 4.04 (s, 3H), 3.91 (s, 2H), 2.99 (br d, *J* = 11.0 Hz, 2H), 2.7–2.6 (m, 1H), 2.32 (dt, *J* = 3.0, 11.5 Hz, 2H), 2.0–1.8 (m, 4H); MS(ES) *m/z* 524.3 $(M+H)^+$.

*2-((4-(6-((4-Chloro-2-fluorobenzyl)oxy)pyridin-2-yl)piperidin-1-yl)methyl)-3-methyl-3H-imidazo[4,5-b]pyridine-5-carboxylic acid hydrochloride (**5**).* 2-((4-(6-((4-chloro-2-fluorobenzyl)oxy)pyridin-2-yl)piperidin-1-yl)methyl)-3-methyl-3*H*-imidazo[4,5-*b*]pyridine-5-carboxylate (**S31**, 57.0 g, 108.8 mmol) was suspended in MeOH (1 L) and treated with 2N NaOH (218 mL). The slurry was stirred for 5 min at room temperature and heated to 85 °C for 3 h. The mixture was filtered through Celite and the clear filtrate reheated to 70 °C. The reaction was acidified with 2N HCl (272 mL) and then allowed to cool to room temperature. The slurry of the white solid was allowed to stir for 18 h at room temperature. The solids were collected by filtration to deliver the desired compound 2-((4-(6-((4-chloro-2-fluorobenzyl)oxy)pyridin-2-yl)piperidin-1-yl)methyl)-3-methyl-3*H*-imidazo[4,5-*b*]pyridine-5-carboxylic acid hydrochloride (**5**, 57.12 g, 96%) as an ivory white solid. $^1$H NMR (400 MHz, DMSO-$d_6$) δ 13.20 (br s, 1H), 11.07 (br s, 1H), 8.27 (d, *J* = 8.6 Hz, 1H), 8.07 (d, *J* = 8.2 Hz, 1H), 7.78–7.57 (m, 2H), 7.47 (dd, *J* = 10.2, 2.0 Hz, 1H), 7.32 (dd, *J* = 8.2, 2.0 Hz, 1H), 6.92 (d, *J* = 6.2 Hz, 1H), 6.74 (d, *J* = 8.2 Hz, 1H), 5.40 (s, 2H), 4.84 (br s, 2H), 3.97 (s, 3H), 3.86 (br s, 2H), 3.37 (br s, 2H), 2.93 (br s, 1H), 2.36–1.85 (m, 4H); $^{13}$C NMR (101 MHz, DMSO-$d_6$, $^1$H and $^{19}$F decoupled) δ 166.2, 162.0, 160.5, 160.3, 150.3,

147.3, 142.8, 140.0, 136.1, 133.6, 132.6, 127.3, 124.7, 123.4, 119.9, 116.0, 114.6, 108.7, 60.2, 52.5, 50.7, 29.3, 28.1; $^{13}$C NMR (101 MHz, DMSO-$d_6$, $^1$H decoupled, just $J_{sp^2 C\text{-}F}$ reported) δ 160.5 (d, $J$ = 249.5 Hz), 133.6 (d, $J$ = 10.6 Hz), 124.7 (d, $J$ = 2.9 Hz), 123.4 (d, $J$ = 15.4 Hz), 116.0 (d, $J$ = 25.1 Hz); $^{19}$F NMR (376 MHz, DMSO-$d_6$,) δ -115.0 (br t, $J$ = 9.0 Hz, 1F); MS(ES) *m/z* 510.2 $(M+H)^+$; HRMS (ESI) m/z $[M + H]^+$ for $C_{26}H_{25}ClFN_5O_3$ calcd 510.17, found 510.1698.

**Synthesis of PF-06883365**

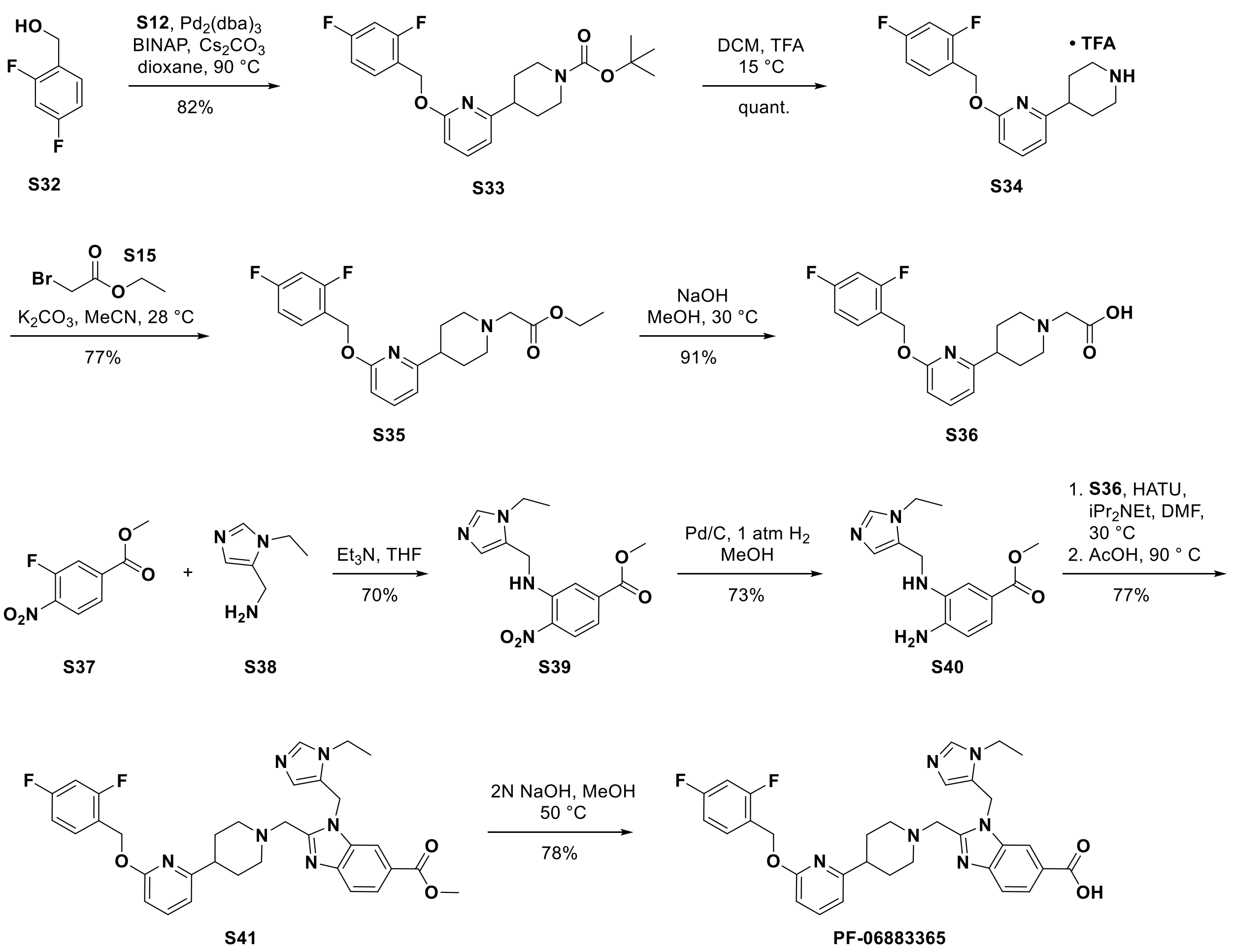


*4-(6-((2,4-Difluorobenzyl)oxy)pyridin-2-yl)piperidine-1-carboxylate (**S33**)*. A suspension of *tert*-butyl 4-(6-chloropyridin-2-yl)piperidine-1-carboxylate (**S12**, 1.98 g, 6.67 mmol), 2,4-difluorobenzyl alcohol (**S32**, 1.25 g, 8.67 mmol), $Pd(OAc)_2$ (105 mg, 0.47 mmol), BINAP (498 mg, 0.80 mmol) and $Cs_2CO_3$ (4.35 g, 13.4 mmol) in dioxane (10 mL) was degassed 3 times with $N_2$ and stirred at 90 °C for 16 h. The brown mixture was diluted with DCM (30 mL) and filtered. The crude product was purified by flash chromatography (20:1 PE/EtOAc, 24 g silica gel) to deliver *tert*-butyl 4-(6-((2,4-difluorobenzyl)oxy)pyridin-2-yl)piperidine-1-carboxylate (**S33**, 2.20 g, 82 %) as a colorless oil. $^1$H NMR (400 MHz, $CDCl_3$) δ 7.5–7.4 (m, 2H), 6.9–6.8 (m, 2H), 6.73 (d, $J$ = 7.5 Hz, 1H), 6.61 (d, $J$ = 7.9 Hz, 1H), 5.40 (s, 2H), 4.23 (br s, 2H), 2.84 (br t, $J$ = 12.1 Hz,

2H), 2.73 (tt, $J$ = 3.6, 11.8 Hz, 1H), 1.9–1.8 (m, 2H), 1.8-1.7 (m, 2H), 1.50 (s, 9H); MS(ES) *m/z* 405.1 $(M+H)^+$.

*2-((2,4-Difluorobenzyl)oxy)-6-(piperidin-4-yl)pyridine (**S34**)*. To a clear solution of *tert*-butyl 4-(6-((2,4-difluorobenzyl)oxy)pyridin-2-yl)piperidine-1-carboxylate (**S33**, 1.9 g, 4.70 mmol) in DCM (20 mL) was added TFA (5 mL). The reaction mixture was stirred at 15 °C for 2 h. The reaction mixture was concentrated to deliver 2-((2,4-difluorobenzyl)oxy)-6-(piperidin-4-yl)pyridine (**S34**, 2.0 g, quant.) as a yellow oil. $^1$H NMR (400 MHz, $CD_3OD$) δ 7.69–7.60 (m, 1H), 7.58–7.51 (m, 1H), 7.02–6.92 (m, 2H), 6.89 (d, $J$ = 7.0 Hz, 1H), 6.71 (d, $J$ = 8.5 Hz, 1H), 5.42 (s, 2H), 3.50 (td, $J$ = 3.0, 12.5 Hz, 2H), 3.14 (dt, $J$ = 3.8, 12.4 Hz, 2H), 3.05–2.93 (m, 1H), 2.19–1.96 (m, 4H); MS(ES) m/z 304.9 $(M+H)^+$.

*Ethyl 2-(4-(6-((2,4-difluorobenzyl)oxy)pyridin-2-yl)piperidin-1-yl)acetate (**S35**)*. $K_2CO_3$ (10.5 g, 76.2 mmol) was added to a stirred solution of 2-((2,4-difluorobenzyl)oxy)-6-(piperidin-4-yl)pyridine (**S34**, 8.11 g, 15.2 mmol) in MeCN (60 mL). The mixture was treated with ethyl bromoacetate (**S15**, 2.80 g, 16.8 mmol). The white suspension was stirred at 28 °C for 16 h. The reaction mixture was treated with water (25 mL) and extracted with EtOAc (3 x 50 mL). The combined organic extracts were dried over $Na_2SO_4$, filtered, and concentrated. The crude product was purified by flash chromatography (10-25% EtOAc/PE gradient, 80 g silica) to deliver ethyl 2-(4-(6-((2,4-difluorobenzyl)oxy)pyridin-2-yl)piperidin-1-yl)acetate (**S35**, 4.59 g, 77%) as a colorless oil. $^1$H NMR (400 MHz, $CDCl_3$) δ 7.54–7.45 (m, 2H), 6.91–6.80 (m, 2H), 6.75 (d, $J$ = 7.5 Hz, 1H), 6.60 (d, $J$ = 8.0 Hz, 1H), 5.41 (s, 2H), 4.22 (q, $J$ = 7.0 Hz, 2H), 3.27 (s, 2H), 3.08 (br d, $J$ = 11.5 Hz, 2H), 2.60 (tt, $J$ = 4.0, 11.5 Hz, 1H), 2.34 (dt, $J$ = 2.5, 11.5 Hz, 2H), 2.07–1.93 (m, 2H), 1.93–1.86 (m, 2H), 1.30 (t, $J$ = 7.0 Hz, 3H).

*2-(4-(6-((2,4-Difluorobenzyl)oxy)pyridin-2-yl)piperidin-1-yl)acetic acid (**S36**)*. To a solution of ethyl 2-(4-(6-((2,4-difluorobenzyl)oxy)pyridin-2-yl)piperidin-1-yl)acetate (**S35**, 1.99 g, 5.10 mmol) in MeOH (5.8 mL) and THF (2.9 mL) was added 3M NaOH (2.6 mL, 7.8 mmol). The reaction was stirred at 30 °C for 16 h. The mixture was neutralized (pH = 7) with 1N HCl and the resultant solution was extracted with 10:1 DCM:MeOH (4 x 10 mL). The combined organic extracts were dried over $MgSO_4$, filtered and concentrated to deliver 2-(4-(6-((2,4-difluorobenzyl)oxy)pyridin-2-yl)piperidin-1-yl)acetic acid (**S36**, 1.68 g, 91%) as a yellow solid. $^1$H NMR (400 MHz, $CD_3OD$) δ 7.63 (dd, $J$ = 7.3, 8.3 Hz, 1H), 7.6–7.5 (m, 1H), 7.0–6.9 (m, 2H), 6.89 (d, $J$ = 7.0 Hz, 1H), 6.70 (d, $J$ = 8.0 Hz, 1H), 5.43 (s, 2H), 3.8–3.7 (m, 2H), 3.67 (s, 2H), 3.2–3.1 (m, 2H), 3.0–2.9 (m, 1H), 2.3–2.1 (m, 4H); MS(ES) *m/z* 363.1 $(M+H)^+$.

*Methyl 3-(((1-ethyl-1H-imidazol-5-yl)methyl)amino)-4-nitrobenzoate (**S39**)*. To a pale yellow solution of methyl 3-fluoro-4-nitrobenzoate (**S37**, 1.59 g, 7.99 mmol) in THF (50 mL) was added (1-ethyl-1*H*-imidazol-5-yl)methanamine (**S38**, 1.00 g, 7.99 mmol), followed by $Et_3N$ (1.62 g, 16.0 mmol). The resultant orange suspension was stirred at 25 °C for 16 h. The reaction mixture was concentrated in vacuo and the crude product was purified by flash chromatography (40 g silica gel, eluting with DCM/MeOH = 50:1 to 10:1) to obtain methyl 3-(((1-ethyl-1*H*-imidazol-5-yl)methyl)amino)-4-nitrobenzoate (**S39**) as yellow oil (1.36 g, 70%). $^1$H NMR (400 MHz, $CD_3OD$) δ 8.23 (d, *J* = 8.5 Hz, 1H), 7.76 (d, *J* = 1.5 Hz, 1H), 7.70 (d, *J* = 1.0 Hz, 1H), 7.28 (dd, *J* = 1.8, 8.8 Hz, 1H), 7.00 (s, 1H), 4.69 (s, 2H), 4.13 (q, *J* = 7.5 Hz, 2H), 3.92 (s, 3H), 1.43 (t, *J* = 7.3 Hz, 3H); MS(ES) m/z 305.2 $(M+H)^+$.

*Methyl 4-amino-3-(((1-ethyl-1H-imidazol-5-yl)methyl)amino)benzoate (**S40**).* A black suspension of methyl 3-(((1-ethyl-1*H*-imidazol-5-yl)methyl)amino)-4-nitrobenzoate (**S39**, 1.36 g, 3.1 mmol) and 10%Pd/C (500 mg) in MeOH (50 mL) was stirred at 25 °C for 1 hour under 1 atm $H_2$. The black suspension was filtered through a pad of celite and the filtrate concentrated to afford methyl 4-amino-3-(((1-ethyl-1*H*-imidazol-5-yl)methyl)amino)benzoate (**S40**) as a grey solid (630 mg, 73%). $^1$H NMR (400 MHz, $CD_3OD$) δ 7.67 (d, *J* = 1.0 Hz, 1H), 7.38 - 7.31 (m, 2H), 6.97 (s, 1H), 6.67 (d, *J* = 8.1 Hz, 1H), 4.35 (s, 2H), 4.12 (q, *J* = 7.2 Hz, 2H), 3.81 (s, 3H), 1.44 (t, *J* = 7.3 Hz, 3H); MS(ES) m/z 275.2 $(M+H)^+$.

*Methyl 2-((4-(6-((2,4-difluorobenzyl)oxy)pyridin-2-yl)piperidin-1-yl)methyl)-1-((1-ethyl-1H-imidazol-5-yl)methyl)-1H-benzo[d]imidazole-6-carboxylate (**S41**).* A dark yellow solution of 2-(4-(6-((2,4-difluorobenzyl)oxy)pyridin-2-yl)piperidin-1-yl)acetic acid (**S36**, 2.38 g, 6.57 mmol), and HATU (3.38 g, 8.89 mmol) in DMF (20 mL) was stirred at 30 °C for 3 h. Then a solution of methyl 4-amino-3-(((1-ethyl-1*H*-imidazol-5-yl)methyl)amino)benzoate (**S40**, 1.47 g, 5.36 mmol) in DMF (6 mL) was added, followed by $iPr_2NEt$ (3.7 mL, 21 mmol). The mixture was stirred at 30 °C for 15 h. The mixture was concentrated under high vacuum. The brown oily residue was washed with saturated aqueous $NH_4Cl$ (40 mL) and then saturated aqueous $NaHCO_3$. The aqueous layer was extracted with DCM/MeOH (10:1, 3 x 50 mL). The combined organic layers were dried over $MgSO_4$, filtered and concentrated. The amide intermediate was taken up in AcOH (80 mL) and the mixture was stirred at 90 °C for 12 h. The reaction mixture was concentrated under high vacuum. The dark yellow oil was neutralized with saturated aqueous $K_2CO_3$ and the mixture was extracted with DCM/MeOH (10:1, 4 x 40 mL). The combined organic layers were dried over $Na_2SO_4$, filtered and concentrated. The crude product was purified by flash chromatography (80 g silica) to deliver methyl 2-((4-(6-((2,4-difluorobenzyl)oxy)pyridin-2-yl)piperidin-1-yl)methyl)-1-((1-ethyl-1*H*-imidazol-5-yl)methyl)-1*H*-benzo[*d*]imidazole-6-carboxylate (**S41**, 2.47 g, 76.6%) as a yellow gum. $^1$H NMR (400 MHz, $CD_3OD$) δ 8.13 (s, 1H), 7.98 (d, *J* = 8.5 Hz, 1H), 7.8–7.7 (m, 2H), 7.6–7.5 (m, 2H), 7.0–6.9 (m, 2H), 6.76 (d, *J* = 7.0 Hz, 1H), 6.60 (d, *J* = 8.0 Hz, 1H), 6.57 (s, 1H), 5.81 (s, 2H), 5.38 (s, 2H), 4.09 (q, *J* = 7.0 Hz, 2H), 3.90 (s, 3H), 3.87 (s, 2H), 2.93 (br d, *J* = 11.0 Hz, 2H), 2.7–2.5 (m, 1H), 2.24 (br t, *J* = 10.8 Hz, 2H), 1.9–1.8 (m, 2H), 1.7–1.6 (m, 2H), 1.28 (t, *J* = 7.3 Hz, 3H); MS(ES) *m/z* 623.2 $(M+H)^+$.

*2-((4-(6-((2,4-difluorobenzyl)oxy)pyridin-2-yl)piperidin-1-yl)methyl)-1-((1-ethyl-1H-imidazol-5-yl)methyl)-1H-benzo[d]imidazole-6-carboxylic acid (**PF-06883365**).* A solution of methyl 2-((4-(6-((2,4-difluorobenzyl)oxy)pyridin-2-yl)piperidin-1-yl)methyl)-1-((1-ethyl-1*H*-imidazol-5-yl)methyl)-1*H*-benzo[*d*]imidazole-6-carboxylate (**S41**, 1.11 g, 1.85 mmol) in MeOH (7.0mL) was treated with aqueous 2M NaOH (2.3 mL, 4.6 mmol). The resulting yellow turbid mixture was stirred at 50 °C for 1.5 h. The reaction was acidified to pH ~ 6–7 with 1N aqueous HCl. The mixture was extracted with DCM/MeOH (10:1, 3 x 10 mL). The combined organic layers were dried over $Na_2SO_4$, filtered and concentrated. The crude product was purified by flash chromatography (12 g silica gel, MeOH/DCM: 0 to 20%) to deliver 2-((4-(6-((2,4-difluorobenzyl)oxy)pyridin-2-yl)piperidin-1-yl)methyl)-1-((1-ethyl-1*H*-imidazol-5-yl)methyl)-1*H*-benzo[*d*]imidazole-6-carboxylic acid (**PF-06883365**, 840 mg, 78%) as a light yellow foamy solid. $^1$H NMR (400 MHz, DMSO-$d_6$) δ 12.79 (br s, 1H), 8.08 (d, $J$ = 0.8 Hz, 1H), 7.82 (dd, $J$ = 1.6, 8.6 Hz, 1H), 7.68 (d, $J$ = 8.6 Hz, 1H), 7.7–7.5 (m, 3H), 7.3–7.2 (m, 1H), 7.09 (dt, $J$ = 2.0, 8.6 Hz, 1H), 6.81 (d, $J$ = 7.4 Hz, 1H), 6.65 (d, $J$ = 8.2 Hz, 1H), 6.46 (s, 1H), 5.75 (s, 2H), 5.34 (s, 2H), 4.00 (q, $J$ = 7.3 Hz, 2H), 3.82 (s, 2H), 2.89 (br d, $J$ = 10.9 Hz, 2H), 2.6–2.5 (m, 1H), 2.16 (br t, $J$ = 10.9 Hz, 2H), 1.74 (br d, $J$ = 11.3 Hz, 2H), 1.6–1.5 (m, 2H), 1.15 (t, $J$ = 7.2 Hz, 3H); $^{13}$C NMR (101 MHz, DMSO-$d_6$, $^1$H and $^{19}$F decoupled) δ 167.7, 162.3, 162.1, 161.9, 160.7, 154.3, 145.2, 139.7, 137.6, 135.4, 132.2, 127.0, 126.5, 124.9, 122.9, 120.8, 118.7, 114.2, 112.5, 111.4, 108.1, 103.9, 60.0, 55.3, 53.4, 42.8, 38.6, 31.1, 16.0; $^{13}$C NMR (101 MHz, DMSO-$d_6$, $^1$H decoupled, just $J_{sp^2\text{C-F}}$ reported) δ 162.1 (dd, $J$ = 12.5, 246.6 Hz), 160.7 (dd, $J$ = 12.5, 248.5 Hz), 132.2 (dd, $J$ = 5.8, 10.6 Hz), 120.8 (m), 111.4 (dd, $J$ = 21.2, 3.9 Hz), 103.9 (t, $J$ = 25.5 Hz), 60.0 (d, $J$ = 2.9 Hz); $^{19}$F NMR (376 MHz, DMSO-$d_6$,) δ -110.1 (quin, $J$ = 7.9 Hz, 1F), -113.7 (q, $J$ = 8.3 Hz, 1F); HRMS (ESI) m/z $[M + H]^+$ for $C_{32}H_{33}F_2N_6O_3$ calcd 587.2582, found 587.2566.

**Synthesis of PF-06882961**

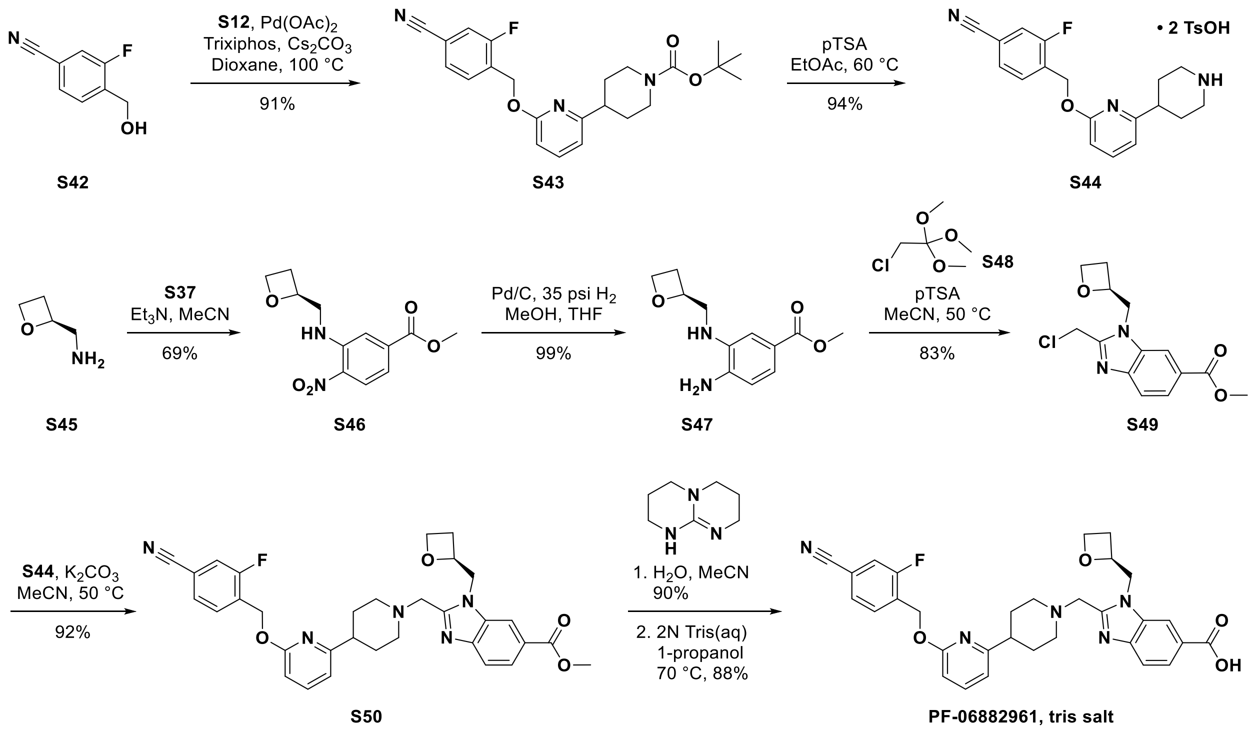

*tert-Butyl 4-(6-((4-cyano-2-fluorobenzyl)oxy)pyridin-2-yl)piperidine-1-carboxylate (**S43**)*. A reaction vessel equipped with a reflux condenser was charged with *tert*-butyl 4-(6-chloropyridin-2-yl)piperidine-1-carboxylate (**S12**, 3.0 g, 10.1 mmol), 3-fluoro-4-(hydroxymethyl)benzonitrile (**S42**, 1.6 g, 10.6 mmol), $Pd(OAc)_2$ (0.11 g, 0.51 mmol), (*S*)-2-(di-*tert*-butylphosphino)-1,1′-binaphthyl (0.20 g, 0.51 mmol) and $Cs_2CO_3$ (6.8 g, 21 mmol). 1,4-Dioxane (51 mL) was added and the mixture was heated to 105 °C. After 2 h, the mixture was allowed to cool to room temperature, filtered through a plug of silica gel and Celite, eluted with 50/50 mixture of EtOAc/heptanes (200 mL) and the filtrate was evaporated under reduced pressure. The crude material was purified using column chromatography eluting with EtOAc in heptane (10:90 to 25:75) to obtain *tert*-butyl 4-(6-((4-cyano-2-fluorobenzyl)oxy)pyridin-2-yl)piperidine-1-carboxylate (**S43**), as an off-white solid (3.8 g, 91%). $^1$H NMR (600 MHz, $CDCl_3$) δ 7.62 (t, *J* = 7.3 Hz, 1H), 7.53 (t, *J* = 7.9 Hz, 1H), 7.44 (d, *J* = 7.6 Hz, 1H), 7.37 (d, *J* = 9.4 Hz, 1H), 6.75 (d, *J* = 7.0 Hz, 1H), 6.65 (d, *J* = 8.2 Hz, 1H), 5.49 (s, 2H), 4.20 (br s, 2H), 2.81 (br s, 2H), 2.70 (tt, *J* = 11.7, 3.5 Hz, 1H), 1.82 (d, *J* = 12.9 Hz, 2H), 1.67 (d, *J* = 11.2 Hz, 2H), 1.49 (s, 9H); MS ($ES^+$): 356.2 $(M\text{-}tBu)^+$.

*3-Fluoro-4-(((6-(piperidin-4-yl)pyridin-2-yl)oxy)methyl)benzonitrile (**S44**)*. To a stirred solution of *tert*-butyl 4-(6-((4-cyano-2-fluorobenzyl)oxy)pyridin-2-yl)piperidine-1-carboxylate (**S43**, 102 g, 242 mmol) in EtOAc (1350 mL) was added pTSA monohydrate (120 g, 630 mmol). The mixture was heated to 60 °C. After 15 min, the solution was allowed to cool to room temperature, the resultant solid was filtered and dried under reduced pressure to obtain 3-fluoro-4-(((6-(piperidin-4-yl)pyridin-2-yl)oxy)methyl)benzonitrile bis-pTSA salt (**S44**, 149 g, 94%) as a white solid. $^1$H NMR (600 MHz, DMSO-$d_6$) δ 8.53 (br s, 1H), 8.26 (br s, 1H), 7.89 (d, *J* = 10.0 Hz, 1H), 7.78-7.67 (m, 3H), 7.48 (d, *J* = 8.2 Hz, 4H), 7.11 (d, *J* = 7.6 Hz, 4H), 6.90 (d, *J* = 7.0 Hz, 1H), 6.79 (d, *J* = 8.2 Hz, 1H), 5.48 (s, 2H), 3.35 (d, *J* = 12.3 Hz, 2H), 3.09-2.96 (m, 2H), 2.96-2.79 (m, 1H), 2.29 (s, 6H), 2.03-1.93 (m, 2H), 1.90-1.77 (m, 2H). MS(ES) *m/z* 312.5 $(M+H)^+$.

*Methyl (S)-4-nitro-3-((oxetan-2-ylmethyl)amino)benzoate (**S46**)*. To a stirred solution of methyl-3-fluoro-4-nitrobenzoate (**S37**, 0.38 g, 4.4 mmol) in MeCN (25 mL) was added (*S*)-oxetan-2-ylmethanamine (**S45**, 1.0 g, 5.2 mmol, (*39*)), followed by $Et_3N$ (1.8 mL, 13.1 mmol). After 18 h, the reaction mixture was concentrated in vacuo, treated with satd. $NH_4Cl$ (15 mL). The aqueous phase was extracted with EtOAc (3 x 25 mL), the combined organic layers were dried over anhydrous $Na_2SO_4$, filtered and evaporated under reduced pressure. The crude material was purified using column chromatography eluting with EtOAc in heptane (0:100 to 40:60) to obtain methyl (*S*)-4-nitro-3-((oxetan-2-ylmethyl)amino)benzoate (**S46**, 0.81 g, 69%) as an orange solid. $^1$H NMR (600 MHz, $CDCl_3$) δ 8.37 (br s, 1H), 8.25 (d, *J* = 8.8 Hz, 1H), 7.64 (s, 1H), 7.28 (d, *J* = 8.8 Hz,1H), 5.22-5.12 (m, 1H), 4.76 (q, *J* = 7.5 Hz, 1H), 4.64 (dt, *J* = 5.9, 9.2 Hz, 1H), 3.95 (s, 3H), 3.64 (t, *J* = 4.3 Hz, 2H), 2.84-2.75 (m, 1H), 2.66-2.57 (m, 1H). MS(ES) *m/z* 242.4 $(M\text{-}24)^+$.

*(S)-4-Amino-3-((oxetan-2-ylmethyl)amino)benzoate (**S47**)*. To a stirred solution of methyl (*S*)-4-nitro-3-((oxetan-2-ylmethyl)amino)benzoate (**S46**, 1.4 g, 5.4 mmol) in MeOH (120 mL) and THF (20 mL) was added Pd/C (10% w/w, 0.42 g, 0.22 mmol). The solution was subject to a hydrogen atmosphere (35 PSI) at room temperature. After 4 h, the solution was filtered through a Celite plug, washed with MeOH (2 x 25 mL) and concentrated under reduced pressure to provide (*S*)-4-amino-3-((oxetan-2-ylmethyl)amino)benzoate (**S47**, 1.3 g, >99%) as a beige oil. $^{1}$H NMR (600 MHz, $CDCl_3$) δ 7.49 (d, 1H), 7.39 (s, 1H), 6.70 (d, *J* = 7.9 Hz, 1H), 5.16-5.08 (m, 1H), 4.76 (q, *J* = 6.2, 9.1 Hz, 1H), 4.62 (q, *J* = 6.2, 9.1 Hz, 1H), 3.87 (br s, 5H), 3.54 (br s, 1H), 3.45 (q, *J* = 5.9, 12.3 Hz, 1H), 3.40-3.34 (m, 1H), 2.83-2.73 (m, 1H), 2.66-2.56 (m, 1H). MS(ES) *m/z* 237.2 $(M+H)^+$.

*(S)-2-(Chloromethyl)-1-(oxetan-2-ylmethyl)-1H-benzo[d]imidazole-6-carboxylate (**S49**)*. To a stirred solution of methyl (*S*)-4-amino-3-((oxetan-2-ylmethyl)amino)benzoate (**S47**, 3.0 g, 12.7 mmol) in THF (51 mL) was added 2-chloro-1,1,1-trimethoxyethane (**S48**, 1.9 mL, 14 mmol) followed by pTSA monohydrate (0.13 g, 0.67 mmol). The mixture was heated to 40 °C. After 1.5 h, the reaction mixture was concentrated in vacuo, and azetroped with 50/50 EtOAc/heptanes (50 mL) to provide an orange solid. The solid was treated with EtOAc (25 mL), heated to 50 °C, diluted with heptane (140 mL) and allowed to cool to room temperature. After 2 h, the mixture was filtered, washed with heptane (2 x 10 mL) and dried to provide (*S*)-2-(chloromethyl)-1-(oxetan-2-ylmethyl)-1*H*-benzo[*d*]imidazole-6-carboxylate (**S49**, 3.1 g, 83%) as an off-white solid. $^{1}$H NMR (600 MHz, $CDCl_3$) δ 8.13 (s, 1H), 8.02 (d, *J* = 8.5 Hz, 1H), 7.81 (d, *J* = 8.5 Hz, 1H), 5.22 (dq, *J* = 2.6, 6.9 Hz, 1H), 5.05 (s, 2H), 4.66-4.60 (m, 2H), 4.57-4.52 (m, 1H), 4.35 (td, *J* = 6.0, 9.1 Hz, 1H), 3.97 (s, 3H), 2.82-2.73 (m, 1H), 2.48-2.39 (m, 1H). MS(AP) *m/z* 295.2 $(M+H)^+$.

*(S)-2-((4-(6-((4-Cyano-2-fluorobenzyl)oxy)pyridin-2-yl)piperidin-1-yl)methyl)-1-(oxetan-2-ylmethyl)-1H-benzo[d]imidazole-6-carboxylate (**S50**)*. To a stirred solution of methyl (*S*)-2-(chloromethyl)-1-(oxetan-2-ylmethyl)-1*H*-benzo[*d*]imidazole-6-carboxylate (**S49**, 1.8 g, 6.1 mmol) in MeCN (35 mL) was added 3-fluoro-4-(((6-(piperidin-4-yl)pyridin-2-yl)oxy)methyl)benzonitrile bis-pTSA salt (**S44**, 4.1 g, 6.1 mmol) and $K_2CO_3$ (4.2 g, 30.4 mmol). The heterogenous mixture was heated to 50 °C. After 2 h, the solution was treated with water (70 mL), allowed to cool to room temperature and stirred for 2 h. The mixture was filtered, washed with water (2 x 30 mL) and dried under reduced pressure to provide (*S*)-2-((4-(6-((4-cyano-2-fluorobenzyl)oxy)pyridin-2-yl)piperidin-1-yl)methyl)-1-(oxetan-2-ylmethyl)-1*H*-benzo[*d*]imidazole-6-carboxylate (**S50**, 3.2 g, 92%) as a white solid. $^{1}$H NMR (600 MHz, $CDCl_3$) δ 8.19 (s, 1H), 7.98 (d, *J* = 8.6 Hz, 1H), 7.76 (d, *J* = 8.6 Hz, 1H), 7.63 (t, *J* = 7.6 Hz, 1H), 7.53 (t, *J* = 7.6 Hz, 1H), 7.45 (d, *J* = 8.2 Hz, 1H), 7.38 (d, *J* = 9.8 Hz, 1H), 6.76 (d, *J* = 7.4 Hz, 1H), 6.65 (d, *J* = 8.2 Hz, 1H), 5.51 (s, 2H), 5.29-5.19 (m, 1H), 4.81-4.68 (m, 2H), 4.67-4.60 (m, 1H), 4.42 (td, *J* = 6.0, 9.1 Hz, 1H), 3.97 (s, 2H), 3.96 (s, 3H), 3.04-2.92 (m, 2H), 2.81-2.69 (m, 1H), 2.67-2.55 (m, 1H), 2.54-2.42 (m, 1H), 2.35-2.21 (m, 2H), 1.93-1.72 (m, 4H). MS(AP) *m/z* 570.5 $(M+H)^+$.

*(S)-2-((4-(6-((4-Cyano-2-fluorobenzyl)oxy)pyridin-2-yl)piperidin-1-yl)methyl)-1-(oxetan-2-ylmethyl)-1H-benzo[d]imidazole-6-carboxylic acid (**PF-06882961**)*. To a stirred solution of methyl (*S*)-2-((4-(6-((4-cyano-2-fluorobenzyl)oxy)pyridin-2-yl)piperidin-1-yl)methyl)-1-(oxetan-2-ylmethyl)-1*H*-benzo[*d*]imidazole-6-carboxylate (**S50**, 4 g, 7 mmol) in MeCN (70 mL) was added a solution of 1,5,7-triazabicyclo[4.4.0]dec-5-ene in water (TBD, 0.97 M, 14.7 mL, 14.4 mmol). After 24 h, the solution was acidified to pH ~4.5 with citric acid in water (1N, 14mL) and diluted with water (50 mL). The aqueous phase was extracted with EtOAc (2 x 75 mL), the combined organic layers were dried over anhydrous $Na_2SO_4$, filtered, and evaporated under reduced pressure to give an off-white solid. The crude material was purified using column chromatography eluting with MeOH/DCM (0:100 to 8:92) to obtain (*S*)-2-((4-(6-((4-cyano-2-fluorobenzyl)oxy)pyridin-2-yl)piperidin-1-yl)methyl)-1-(oxetan-2-ylmethyl)-1*H*-benzo[*d*]imidazole-6-carboxylic acid (**PF-06882961**, 3.65 g, 90%) as a white amorphous solid. $^1$H NMR (400 MHz, DMSO-*d*$_6$) δ 12.75 (br s, 1H), 8.27 (s, 1H), 7.89 (d, *J* = 10.1 Hz, 1H), 7.80 (d, *J* = 9.4 Hz, 1H), 7.72-7.68 (m, 2H), 7.67-7.60 (m, 2H), 6.89 (d, *J* = 7.4 Hz, 1H), 6.72 (d, *J* = 8.2 Hz, 1H), 5.47 (s, 2H), 5.11 (d, *J* = 3.9 Hz, 1H), 4.86-4.74 (m, 1H), 4.72-4.62 (m, 1H), 4.53-4.43 (m, 1H), 4.35-4.32 (m, 1H), 3.95 (d, *J* = 13.7 Hz, 1H), 3.77 (d, *J* = 13.7 Hz, 1H), 2.98 (d, *J* = 10.1 Hz, 1H), 2.84 (d, *J* = 11.7 Hz, 1H), 2.77-2.65 (m, 1H), 2.64-2.53 (m, 1H), 2.45-2.37 (m, 1H), 2.28-2.10 (m, 2H), 1.84-1.57 (m, 4H). MS(ES) *m/z* 556.6 $(M+H)^+$.

*(S)-2-((4-(6-((4-Cyano-2-fluorobenzyl)oxy)pyridin-2-yl)piperidin-1-yl)methyl)-1-(oxetan-2-ylmethyl)-1H-benzo[d]imidazole-6-carboxylic acid (**PF-06882961**, tris salt)*. To a stirred solution of (*S*)-2-((4-(6-((4-cyano-2-fluorobenzyl)oxy)pyridin-2-yl)piperidin-1-yl)methyl)-1-(oxetan-2-ylmethyl)-1*H*-benzo[*d*]imidazole-6-carboxylic acid (**PF-06882961**, 6.5 g, 11.7 mmol) in 1-propanol (275 mL) at 70 °C was added a solution of tris(hydroxymethyl)aminomethane in water (2M, 6.1 mL, 12.2 mmol) drop wise. The homogenous solution was seeded, allowed to cool to room temperature over 2 h and stirred for 15 h. The mixture was filtered, washed with 1-propanol (2 x 30 mL) and dried under reduced pressure to provide (*S*)-2-((4-(6-((4-cyano-2-fluorobenzyl)oxy)pyridin-2-yl)piperidin-1-yl)methyl)-1-(oxetan-2-ylmethyl)-1*H*-benzo[*d*]imidazole-6-carboxylic acid (**PF-06882961**, tris salt, 7.0 g, 88%) as a white crystalline solid. $^1$H NMR (400 MHz, DMSO-*d*$_6$) δ 8.20 (s, 1H), 7.89 (d, *J* = 9.8 Hz, 1H), 7.79 (d, *J* = 8.3 Hz, 1H), 7.70 (br. s., 2H), 7.64 (t, *J* = 7.8 Hz, 1H), 7.55 (d, *J* = 8.3 Hz, 1H), 6.88 (d, *J* = 7.3 Hz, 1H), 6.72 (d, *J* = 8.3 Hz, 1H), 5.47 (s, 2H), 5.15-5.06 (m, 1H), 4.84-4.73 (m, 1H), 4.72-4.59 (m, 1H), 4.53-4.44 (m, 1H), 4.38 (dt, *J* = 6.0, 8.8 Hz, 1H), 3.93 (d, *J* = 13.4 Hz, 1H), 3.76 (d, *J* = 13.4 Hz, 1H), 3.36 (s, 9H), 2.98 (d, *J* = 11.0 Hz, 1H), 2.85 (d, *J* = 11.0 Hz, 1H), 2.77-2.64 (m, 1H), 2.63-2.53 (m, 1H), 2.47-2.40 (m, 1H), 2.26-2.11 (m, 2H), 1.82-1.56 (m, 4H); $^{13}$C NMR (101 MHz, DMSO-*d*$_6$, $^1$H and $^{19}$F decoupled) δ 169.7, 162.4, 161.6, 159.6, 153.6, 143.9, 139.9, 135.6, 131.2, 131.0, 129.7, 128.7, 123.0, 119.2, 117.6, 114.6, 112.3, 112.0, 108.1, 80.6, 67.6, 61.1, 60.0, 59.2, 55.1, 53.7, 53.2, 48.9, 42.8, 31.2, 31.1, 24.5; $^{13}$C NMR (101 MHz, DMSO-d$_6$, $^1$H decoupled, just $J_{sp^2 C\text{-}F}$ reported) δ 159.5 (d, *J* = 248.5 Hz), 133.6 (d, *J* = 4.5 Hz), 131.0 (d, *J* = 14.6 Hz), 128.7 (d,

$J$ = 3.5 Hz), 119.2 (d, $J$ = 25.6 Hz), 117.5 (d, $J$ = 2.5 Hz), 112.0 (d, $J$ = 10 Hz); $^{19}$F NMR (376 MHz, DMSO-$d_6$) δ -115.5. HRMS (ESI) *m/z* $[M + H]^+$ for $C_{31}H_{30}FN_5O_4$ calcd 556.2350, found 556.2334. M.P. = 194 °C.

**Synthesis of Tri-iodo PF-06883365 (S58)**

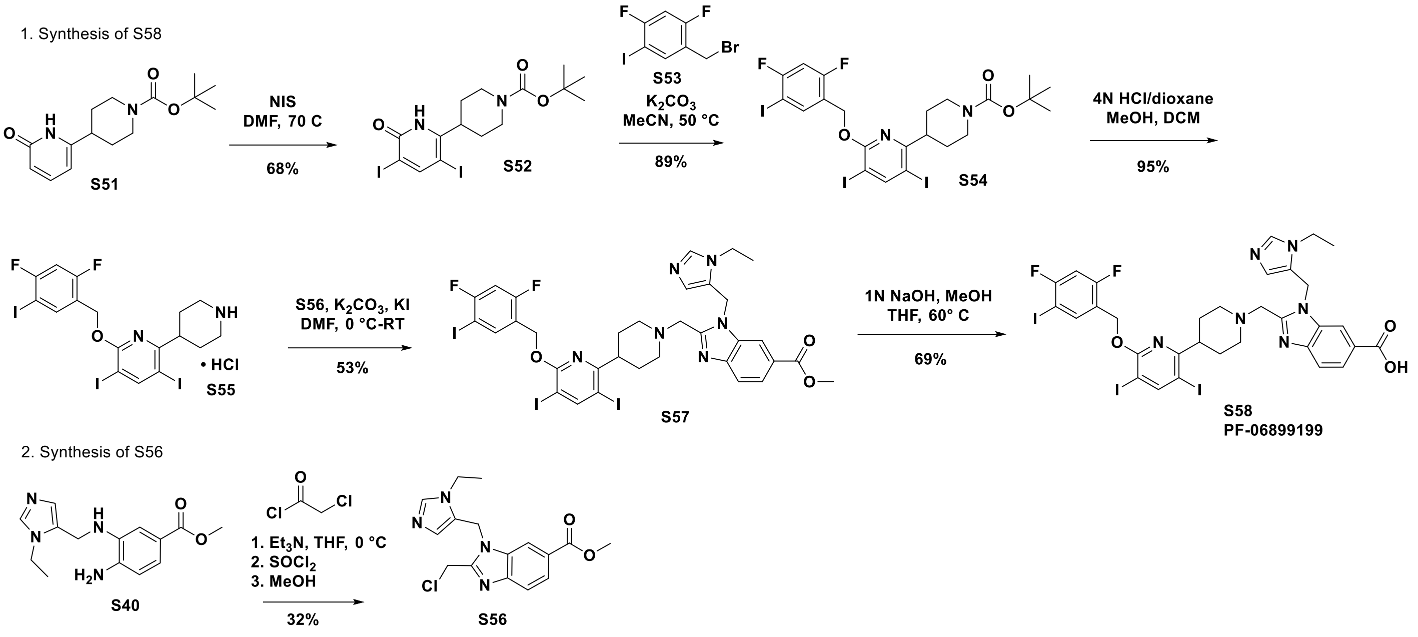


*tert-butyl 4-(6-hydroxy-3,5-diiodopyridin-2-yl)piperidine-1-carboxylate (**S52**).* A solution of tert-butyl 4-(6-hydroxypyridin-2-yl)piperidine-1-carboxylate (**S51**, 470 mg, 1.69 mmol) in DMF (10mL) was treated with N-iodosuccinimide (NIS) (760 mg, 3.38 mmol) and heated to 70 °C. After 6 h, a solution of $Na_2S_2O_3$ (50 mg in 10 mL water) was added and the reaction mixture was stirred at 25 °C for 30 mins. The mixture was extracted with DCM (3 x 10 mL). The combined organic layers were washed with brine (10 mL), dried over $Na_2SO_4$, filtered and concentrated. The crude material was purified using column chromatography eluting with EtOAc in heptane (0:100 to 50:50) to obtain tert-butyl 4-(6-hydroxy-3,5-diiodopyridin-2-yl)piperidine-1-carboxylate (**S52**, 610 mg, 68%) as a white solid. $^1$H NMR (400 MHz, DMSO-$d_6$) δ 11.85 (br s, 1H), 8.28 (s, 1H), 4.08 (br d, $J$ = 9.6 Hz, 2H), 2.96 (br s, 1H), 2.85 - 2.60 (m, 2H), 1.79 (br d, $J$ = 2.6 Hz, 2H), 1.56 (br d, $J$ = 8.3 Hz, 2H), 1.41 (s, 9H); MS(ES) *m/z* 553.0 $(M+H)^+$.

*tert-butyl 4-(6-((2,4-difluoro-5-iodobenzyl)oxy)-3,5-diiodopyridin-2-yl)piperidine-1-carboxylate (**S54**).* To a 25 mL RBF charged with *tert*-butyl 4-(6-hydroxy-3,5-diiodopyridin-2-yl)piperidine-1-carboxylate (**S52**, 570 mg, 1.08 mmol) in MeCN (20 mL) was added $K_2CO_3$ (297 mg, 2.15 mmol) and 2,4-difluoro-5-iodobenzyl bromide (**S53**, 430 mg, 1.29 mmol). The mixture was heated to 50 °C. After 10 h, the reaction mixture was concentrated under reduced pressure, the resultant residue was taken up in EtOAc (20 mL) and washed with water (15 mL), dried over $Na_2SO_4$, filtered and concentrated. The crude material was purified using column chromatography eluting with EtOAc in heptane (0:100 to 30:70) to obtain *tert*-butyl 4-(6-((2,4-difluoro-5-iodobenzyl)oxy)-3,5-diiodopyridin-2-yl)piperidine-1-carboxylate (**S54**, 750 mg, 89%) as a white solid. $^1$H NMR (400 MHz, $CDCl_3$) δ 8.32 (s, 1H), 7.96 (t, *J* = 7.5 Hz, 1H), 6.88 (dd, *J* = 7.9, 9.6 Hz, 1H), 5.37 (s, 2H), 4.26 (br d, *J* = 12.7 Hz, 2H), 3.10 (tt, *J* = 5.2, 10.2 Hz, 1H), 2.92 - 2.75 (m, 2H), 1.81 - 1.68 (m, 4H), 1.50 (s, 9H); MS(ES) *m/z* 727.1 (M+H-*t*-Bu)$^+$.

*2-((2,4-difluoro-5-iodobenzyl)oxy)-3,5-diiodo-6-(piperidin-4-yl)pyridine hydrochloride (**S55**).* To a solution of *tert*-butyl 4-(6-((2,4-difluoro-5-iodobenzyl)oxy)-3,5-diiodopyridin-2-yl)piperidine-1-carboxylate (**S54**, 750 mg, 0.96 mmol) in MeOH (2 mL) and DCM (4 mL) was added HCl in dioxane (4N, 1 mL, 4 mmol) in one portion and the resultant mixture was stirred at 25 °C for 1 h. The reaction mixture was concentrated under reduced pressure. The residue was dissolved in DCM (30 mL) and quenched with saturated aqueous $NaHCO_3$ (15 mL). The mixture was extracted with DCM (5 x 50 mL). The combined extracts were washed with brine (20 mL), dried over $Na_2SO_4$, filtered and concentrated to obtain 2-((2,4-difluoro-5-iodobenzyl)oxy)-3,5-diiodo-6-(piperidin-4-yl)pyridine hydrochloride (**S55**, 620 mg, 95%) as a white solid. $^1$H NMR (400 MHz, DMSO-$d_6$) δ 8.45 (s, 1H), 8.07 (t, *J* = 7.7 Hz, 1H), 7.43 (dd, *J* = 8.3, 10.0 Hz, 1H), 5.39 (s, 2H), 3.09 - 2.90 (m, 3H), 2.65 - 2.52 (m, 2H), 1.75 - 1.56 (m, 4H); MS(ES) *m/z* 682.9 (M+H)$^+$.

*Methyl 2-(chloromethyl)-1-((1-ethyl-1H-imidazol-5-yl)methyl)-1H-benzo[d]imidazole-6-carboxylate (**S56**).* To a solution of methyl 4-amino-3-(((1-ethyl-1*H*-imidazol-5-yl)methyl)amino)benzoate (**S40**, 600 mg, 2.19 mmol) in THF (5 mL) was successively added triethylamine (221 mg, 2.19 mmol) and chloroacetyl chloride (494 mg, 4.37 mmol) at 0 °C and the mixture was stirred for 1 h. The mixture was treated with $SOCl_2$ (1.56 g, 13.1 mmol) and stirring was continued for 15 h at 25 °C. The mixture was quenched by addition of MeOH (3 mL) and concentrated under reduced pressure. The crude material was purified using column chromatography eluting with MeOH in DCM (1:30 to 1:15) to obtain methyl 2-(chloromethyl)-1-((1-ethyl-1*H*-imidazol-5-yl)methyl)-1*H*-benzo[*d*]imidazole-6-carboxylate (**S56**, 230 mg, 32%) as a pale yellow solid. $^1$H NMR (400 MHz, DMSO-$d_6$) δ 9.20 (d, *J* = 1.5 Hz, 1H), 8.29 (d, *J* = 1.0 Hz, 1H), 7.92 (dd, *J* = 1.6, 8.4 Hz, 1H), 7.82 (d, *J* = 8.6 Hz, 1H), 7.13 (d, *J* = 0.7 Hz, 1H), 5.93 (s, 2H), 5.13 (s, 2H), 4.25 (q, *J* = 7.3 Hz, 2H), 3.87 (s, 3H), 1.38 (t, *J* = 7.3 Hz, 3H); MS(ES) *m/z* 333.0 (M+H)$^+$.

*Methyl 2-((4-(6-((2,4-difluoro-5-iodobenzyl)oxy)-3,5-diiodopyridin-2-yl)piperidin-1-yl)methyl)-1-((1-ethyl-1H-imidazol-5-yl)methyl)-1H-benzo[d]imidazole-6-carboxylate (**S57**).* To a solution of 2-((2,4-difluoro-5-iodobenzyl)oxy)-3,5-diiodo-6-(piperidin-4-yl)pyridine hydrochloride (**S55**, 613 mg, 0.90 mmol) in DMF (8 mL) was successively added $K_2CO_3$ (191 mg, 1.38 mmol), KI (115 mg, 0.691 mmol) and methyl 2-(chloromethyl)-1-((1-ethyl-1*H*-imidazol-5-yl)methyl)-1*H*-benzo[*d*]imidazole-6-carboxylate (**S56**, 230 mg, 0.691 mmol) at 0 °C and the mixture was stirred for 16 h at 25 °C. The reaction mixture was concentrated under reduced pressure and the resultant residue was taken up in DCM (20 mL) and washed with water (15 mL), dried over $Na_2SO_4$, filtered and concentrated. The crude material was purified using column chromatography eluting with MeOH in DCM (0:100 to 30:70) to obtain methyl 2-((4-(6-((2,4-difluoro-5-iodobenzyl)oxy)-3,5-diiodopyridin-2-yl)piperidin-1-yl)methyl)-1-((1-ethyl-1*H*-imidazol-5-yl)methyl)-1*H*-benzo[*d*]imidazole-6-carboxylate (**S57**, 360 mg, 53%) as a white solid. $^1$H NMR (400 MHz, $CDCl_3$) δ 8.32 (s, 1H), 8.09 (s, 1H), 8.04 - 7.95 (m, 2H), 7.79 (d, *J* = 8.3 Hz, 1H), 7.53 (br s, 1H), 6.86 (dd, *J* = 7.5, 9.6 Hz, 2H), 5.76 (br s, 2H), 5.40 (s, 2H), 3.95 (s, 3H), 3.88 (br dd, *J* = 6.4, 13.4 Hz, 2H), 3.12 - 2.95 (m, 3H), 2.36 (br d, *J* = 7.0 Hz, 2H), 1.84 (br s, 2H), 1.77 - 1.57 (m, 4H), 1.33 - 1.16 (m, 3H); MS(ES) *m/z* 978.8 $(M+H)^+$.

*2-((4-(6-((2,4-difluoro-5-iodobenzyl)oxy)-3,5-diiodopyridin-2-yl)piperidin-1-yl)methyl)-1-((1-ethyl-1H-imidazol-5-yl)methyl)-1H-benzo[d]imidazole-6-carboxylic acid (**S58**).* To a solution of methyl 2-((4-(6-((2,4-difluoro-5-iodobenzyl)oxy)-3,5-diiodopyridin-2-yl)piperidin-1-yl)methyl)-1-((1-ethyl-1*H*-imidazol-5-yl)methyl)-1*H*-benzo[*d*]imidazole-6-carboxylate (**S57**, 360 mg, 0.37 mmol) in THF (6 mL) and MeOH (3 mL) at 25 °C was added aqueous NaOH (1N, 1.1 mL, 1.1 mmol). The reaction mixture was heated to 60 °C and stirred for 2 h. The reaction was cooled to 25 °C and the solvent was removed under reduced pressure. Water (20 mL) was added and the mixture was acidified to pH ~2-3 with 1M HCl. The resultant suspension was stirred for 30 mins to granulate. The solids were collected by filtration, rinsed with water and air dried to obtain 2-((4-(6-((2,4-difluoro-5-iodobenzyl)oxy)-3,5-diiodopyridin-2-yl)piperidin-1-yl)methyl)-1-((1-ethyl-1*H*-imidazol-5-yl)methyl)-1*H*-benzo[*d*]imidazole-6-carboxylic acid (**S58 PF-06899199**, 245 mg, 69%) as an off-white solid. $^1$H NMR (600 MHz, DMSO-$d_6$) δ 12.83 - 12.69 (m, 1H), 8.45 (s, 1H), 8.08 (s, 1H), 8.04 (br t, *J* = 7.5 Hz, 1H), 7.82 (br d, *J* = 8.5 Hz, 1H), 7.77 (br s, 1H), 7.70 (br d, *J* = 8.5 Hz, 1H), 7.41 (br t, *J* = 9.1 Hz, 1H), 6.53 (s, 1H), 5.77 (s, 2H), 5.38 (s, 2H), 4.03 (q, *J* = 7.0 Hz, 2H), 3.89 (br s, 2H), 2.98 (br s, 2H), 2.90 (br s, 1H), 2.21 (br s, 2H), 1.65 (br s, 4H), 1.18 (br t, *J* = 7.0 Hz, 3H); $^{13}$C NMR (101 MHz, DMSO-$d_6$, $^1$H and $^{19}$F decoupled) δ 167.5, 161.3, 160.8, 160.1, 156.5, 145.1, 140.0, 136.4, 134.9, 128.9, 125.6, 123.6, 122.3, 120.5, 119.1, 112.8, 104.7, 86.7, 79.3, 76.5, 61.5, 52.6, 45.3, 41.3, 38.5, 34.0, 29.0, 25.1, 14.9, 8.4; $^{19}$F NMR (376 MHz, DMSO-$d_6$) δ -91.0, -113.4; HRMS (ESI) *m/z* $[M + H]^+$ for $C_{32}H_{29}F_2I_3N_6O_3$ calcd 964.9476, found 964.9484.

### NMR Spectra

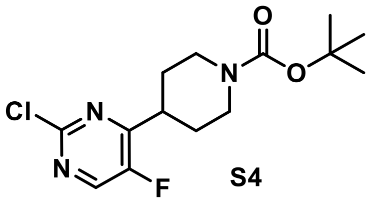

S4

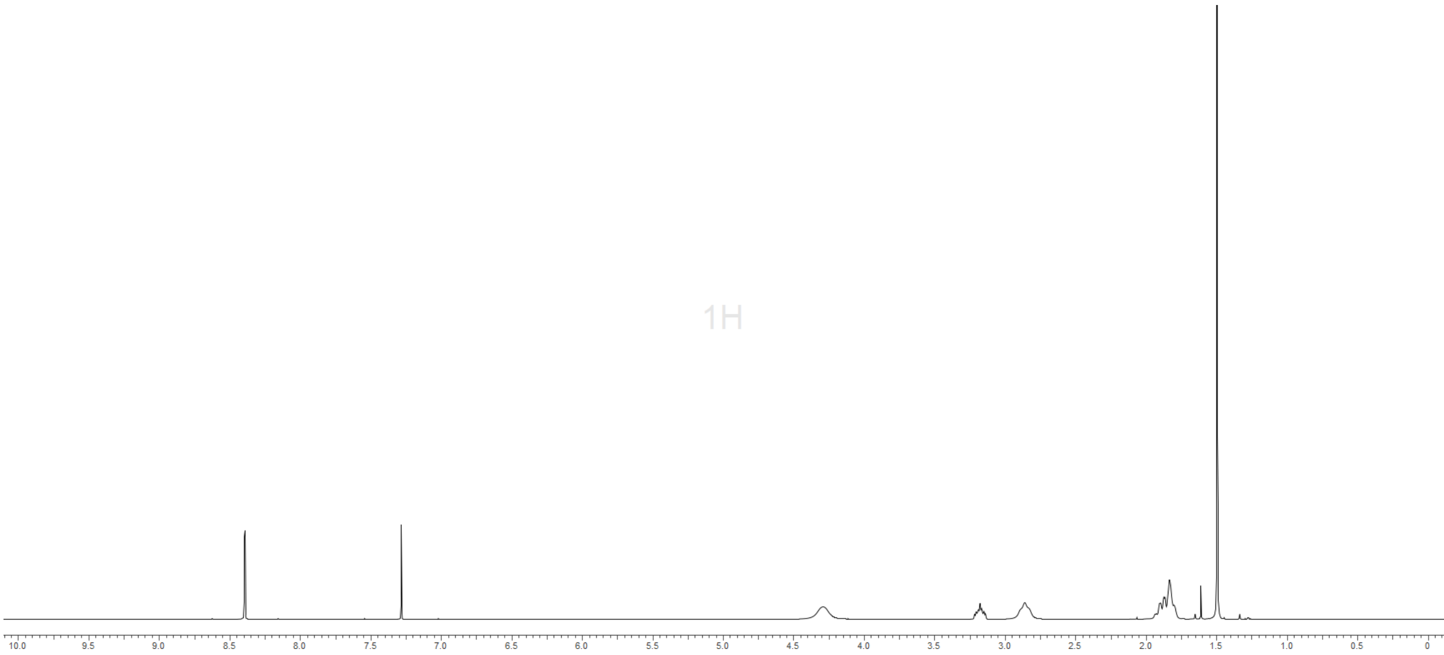

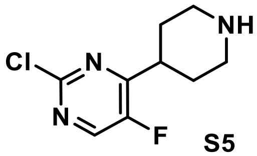

Cl
N
NH
N
F


S5

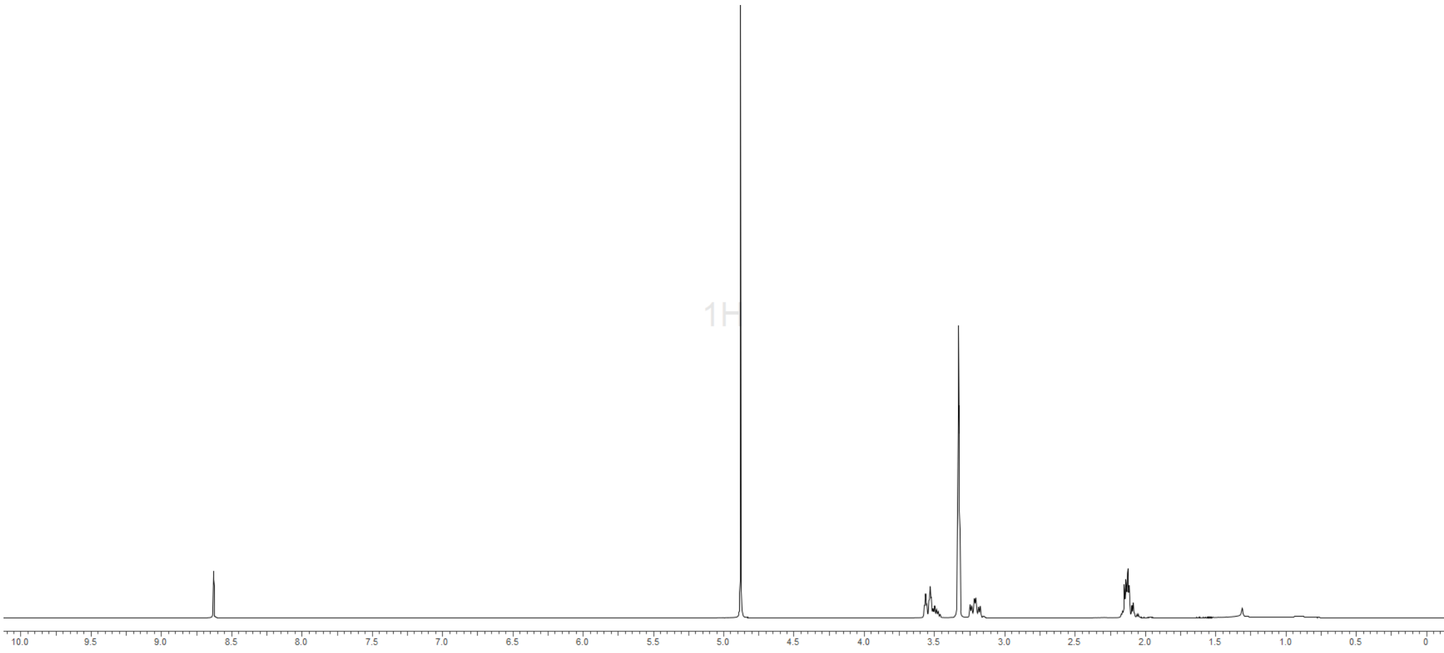

1H
10.0 9.5 9.0 8.5 8.0 7.5 7.0 6.5 6.0 5.5 5.0 4.5 4.0 3.5 3.0 2.5 2.0 1.5 1.0 0.5 0


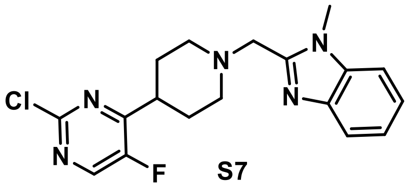

Cl
N
N
N
N
N
F


S7

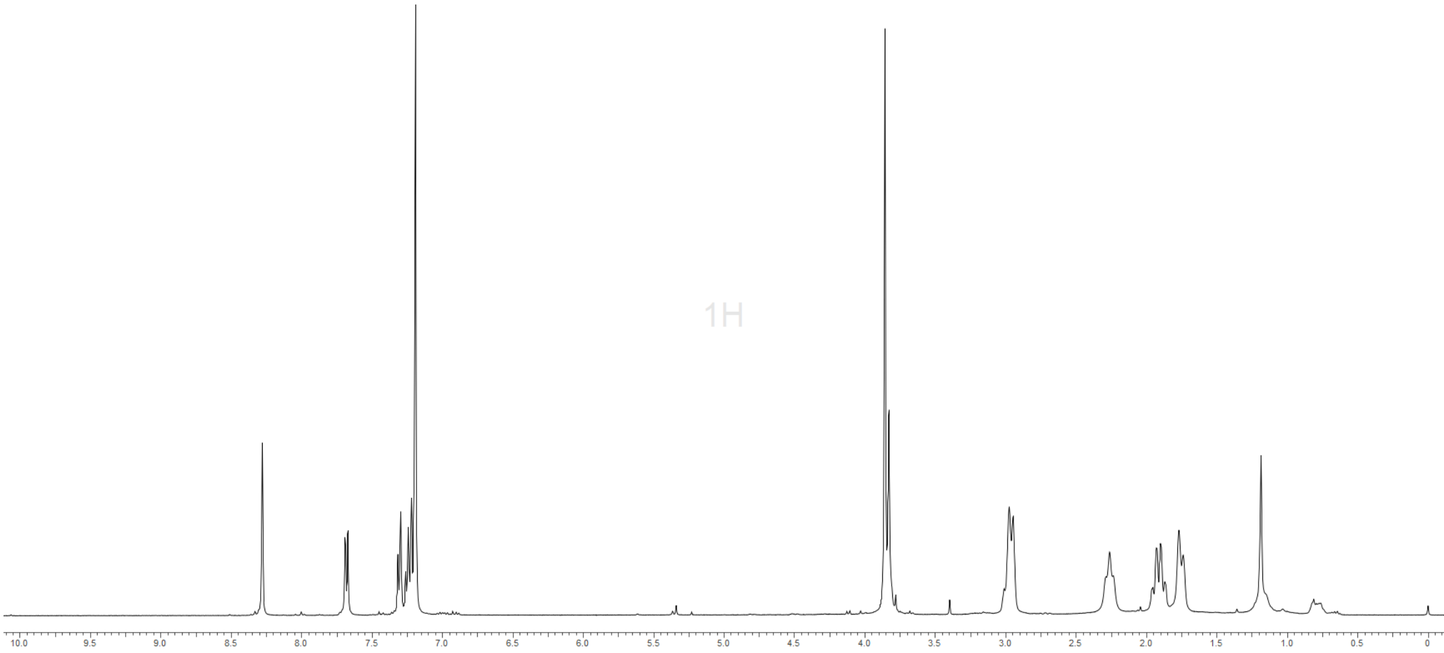

1H
10.0 9.5 9.0 8.5 8.0 7.5 7.0 6.5 6.0 5.5 5.0 4.5 4.0 3.5 3.0 2.5 2.0 1.5 1.0 0.5 0

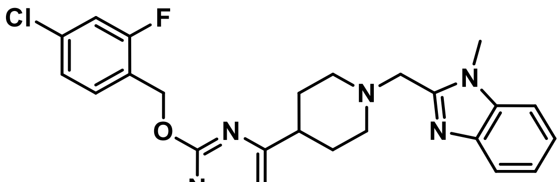

**2**

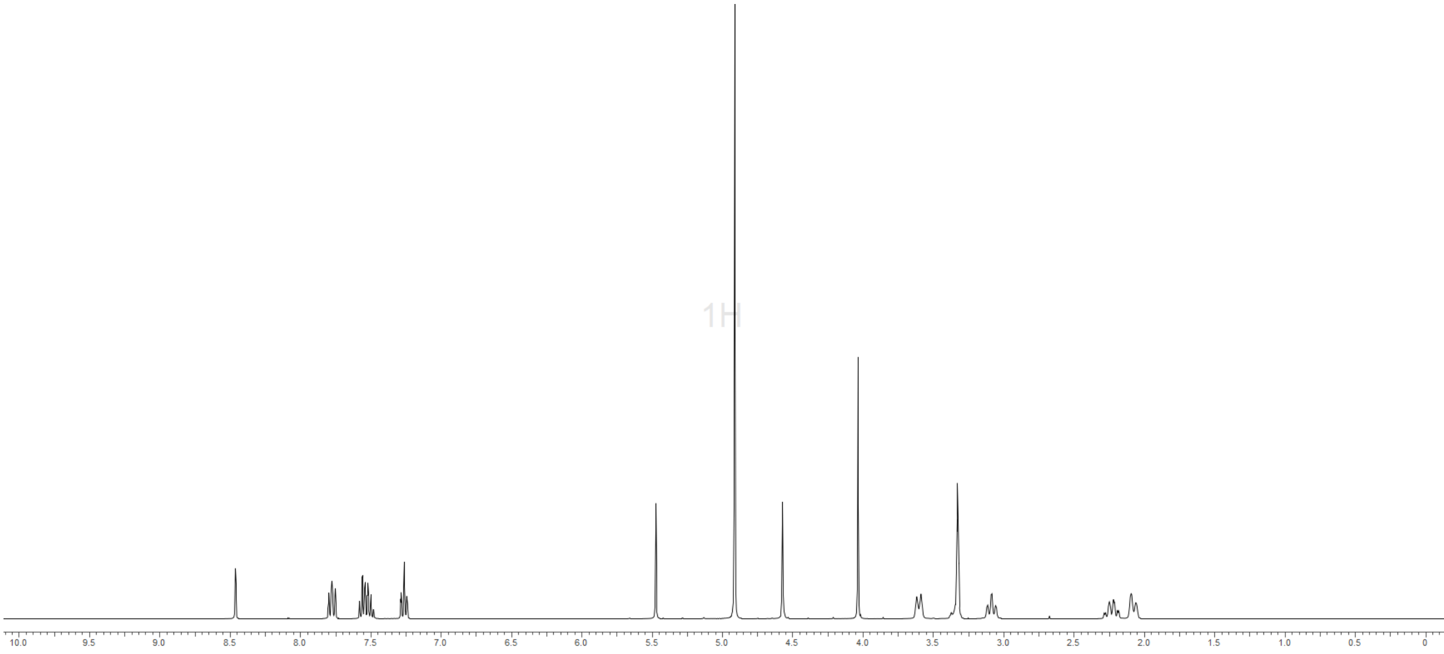


($^{1}H$ and $^{19}F$ decoupled)

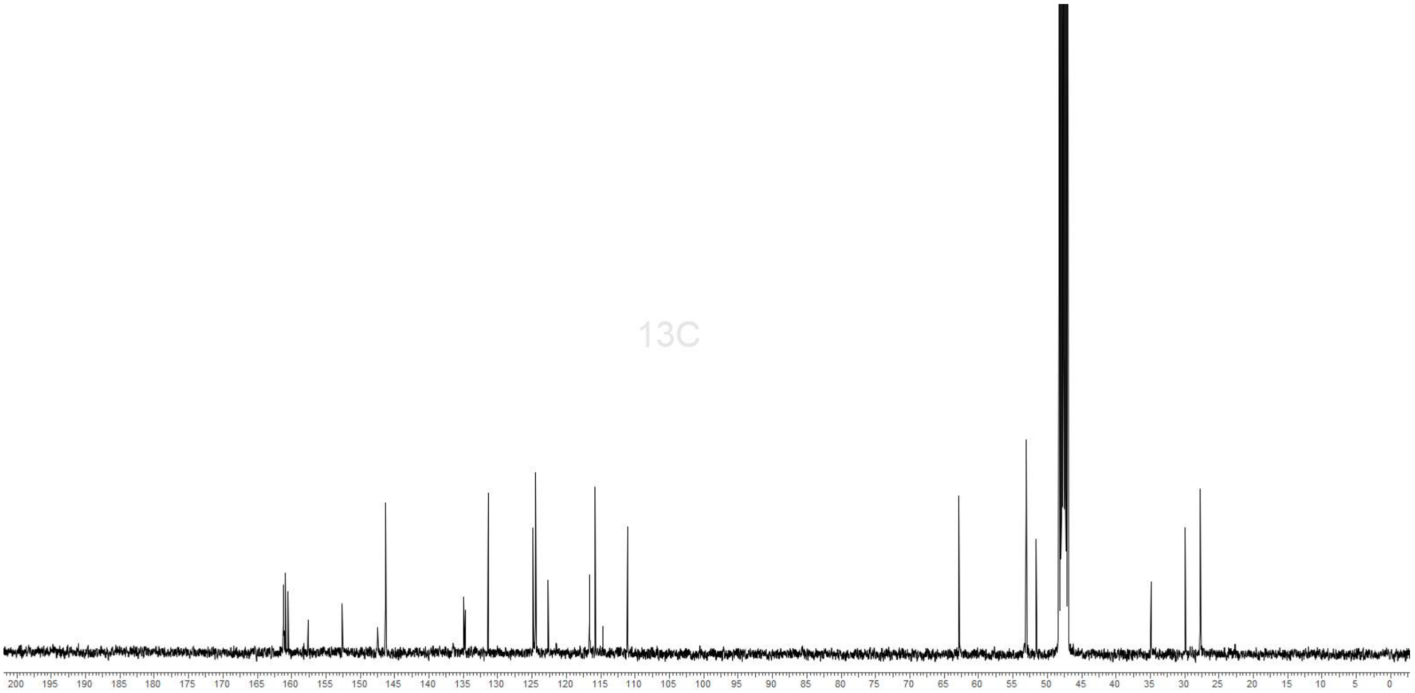

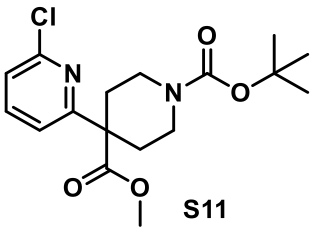

S11

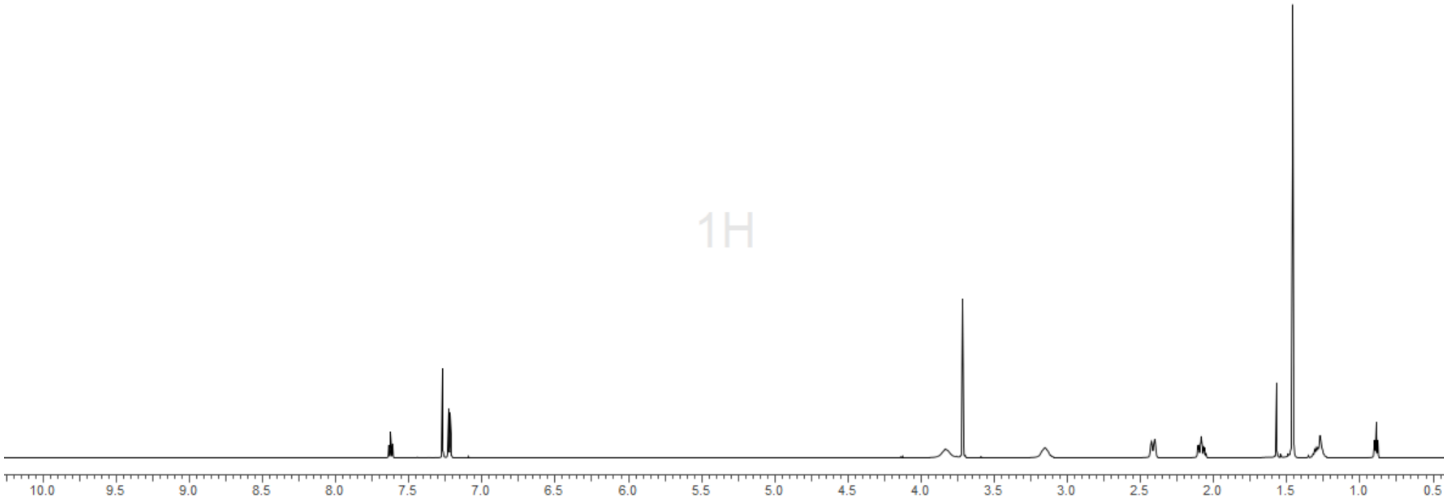


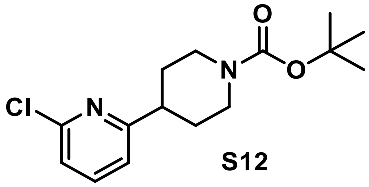

S12

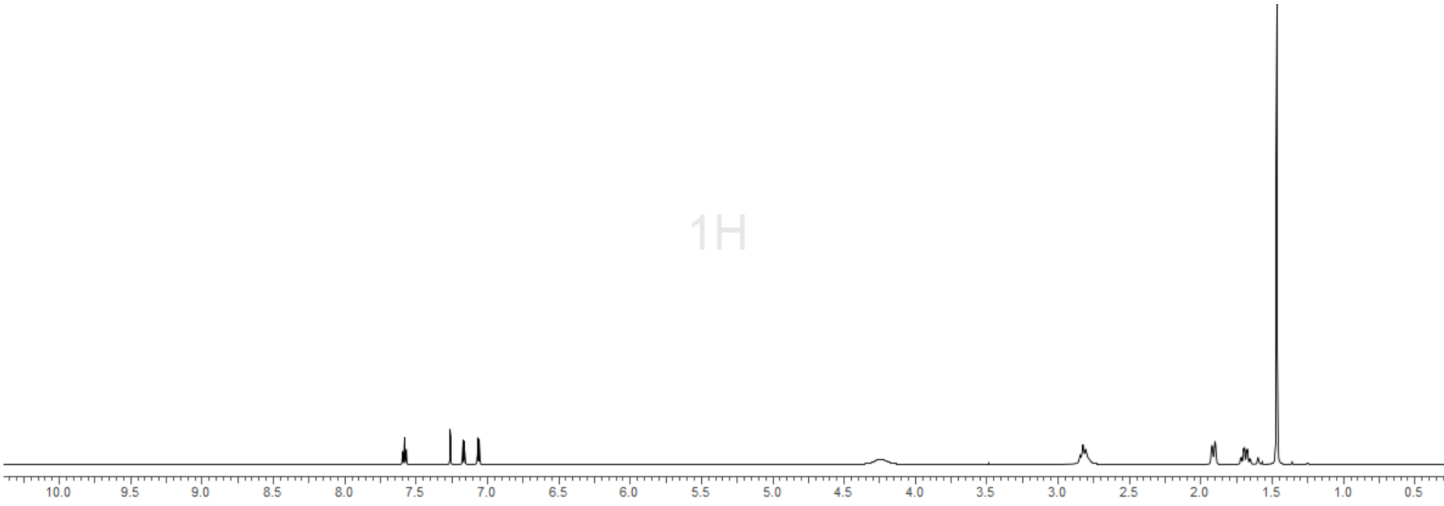

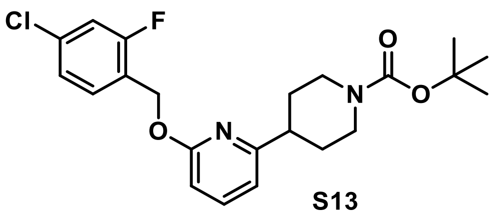
Cl
F
O
N
O
O
N

**S13**

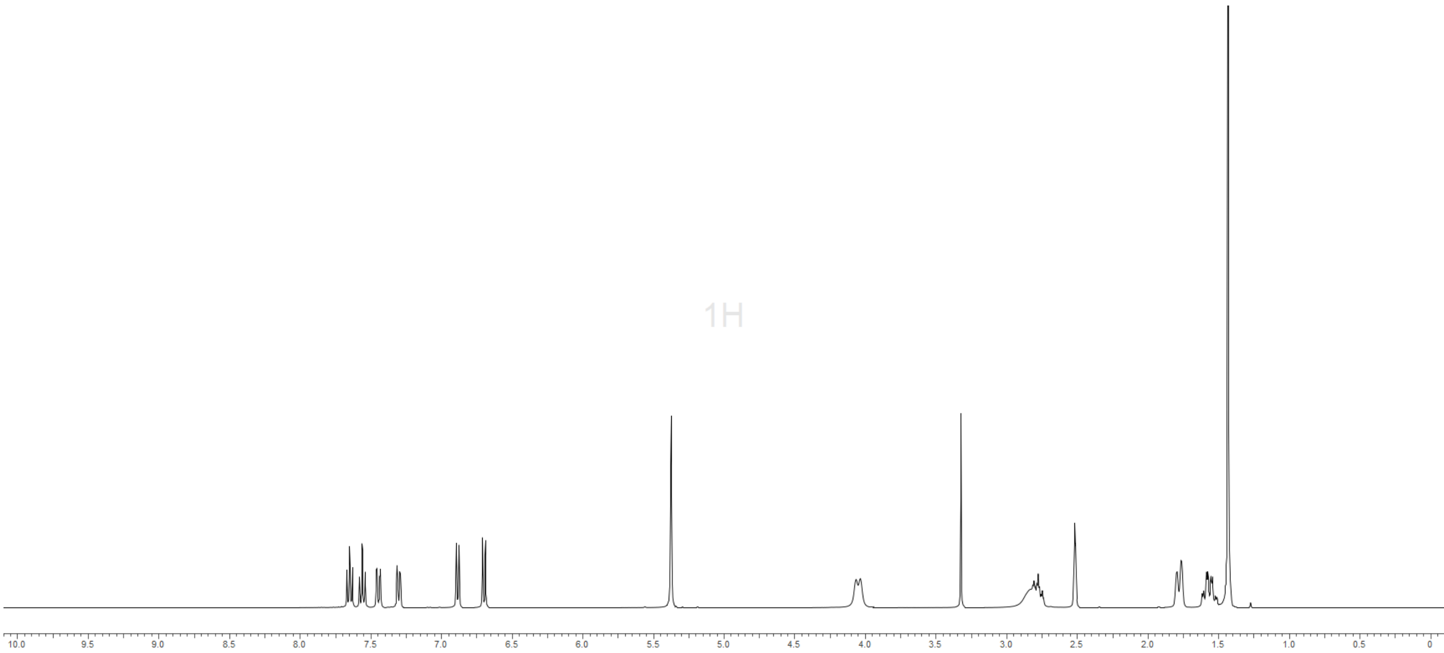
1H
10.0 9.5 9.0 8.5 8.0 7.5 7.0 6.5 6.0 5.5 5.0 4.5 4.0 3.5 3.0 2.5 2.0 1.5 1.0 0.5 0

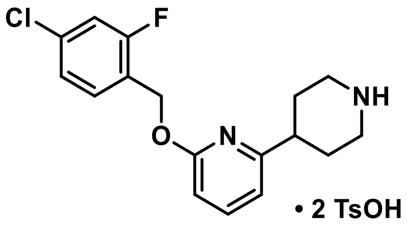
Cl
F
O
N
NH
• 2 TsOH

S14

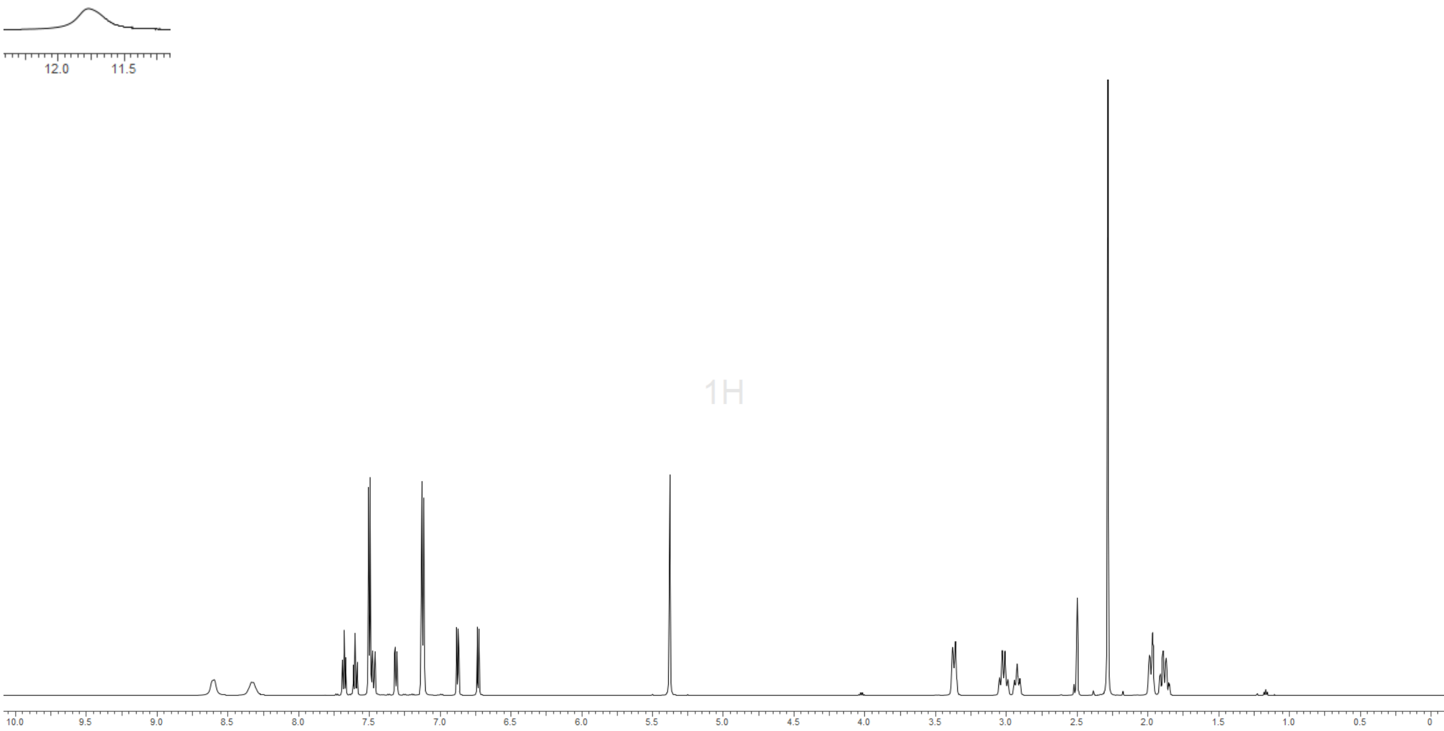
12.0
11.5
1H
10.0
9.5
9.0
8.5
8.0
7.5
7.0
6.5
6.0
5.5
5.0
4.5
4.0
3.5
3.0
2.5
2.0
1.5
1.0
0.5
0

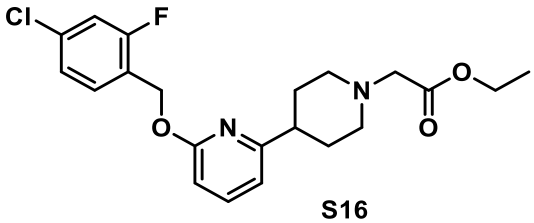
Cl
F
O
N
N
O
O

**S16**

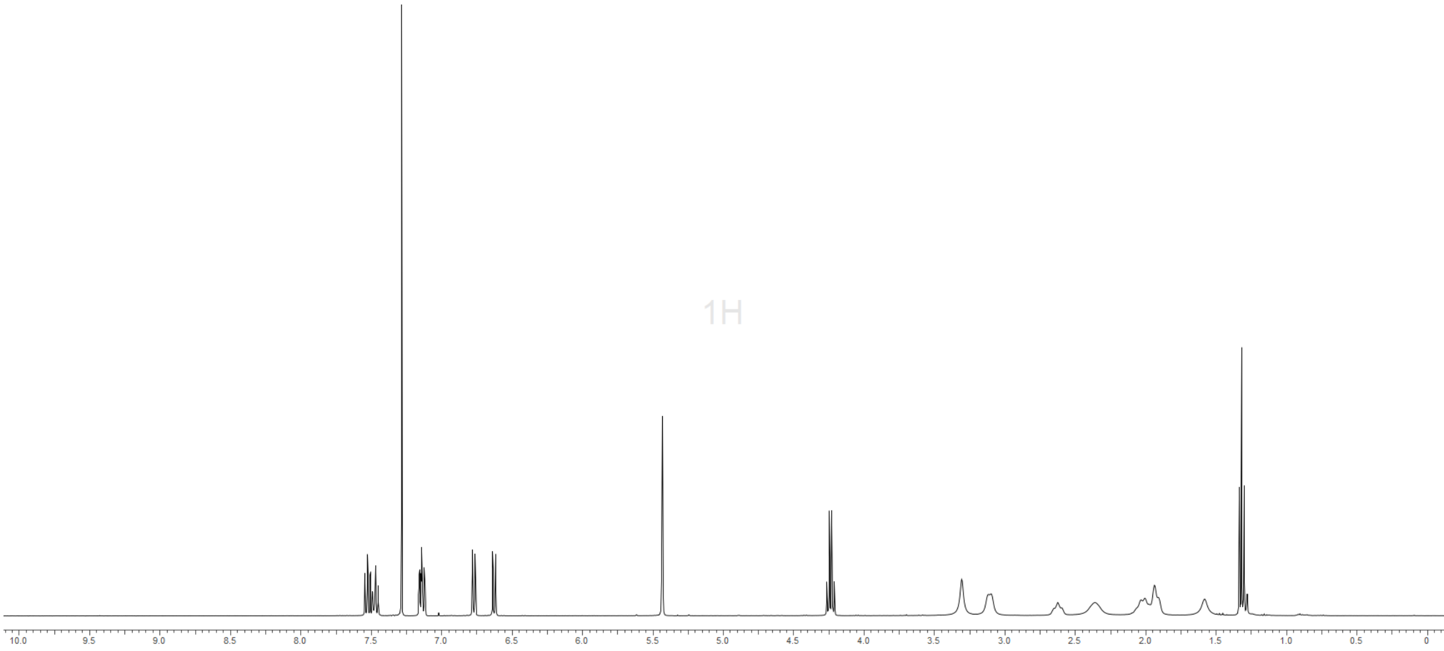
1H
10.0
9.5
9.0
8.5
8.0
7.5
7.0
6.5
6.0
5.5
5.0
4.5
4.0
3.5
3.0
2.5
2.0
1.5
1.0
0.5
0

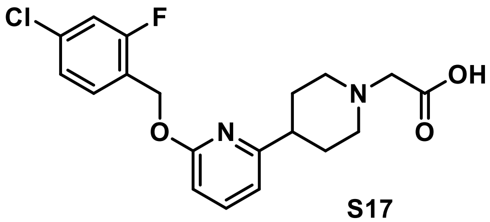
Cl
F
OH
N
O
N
O

S17

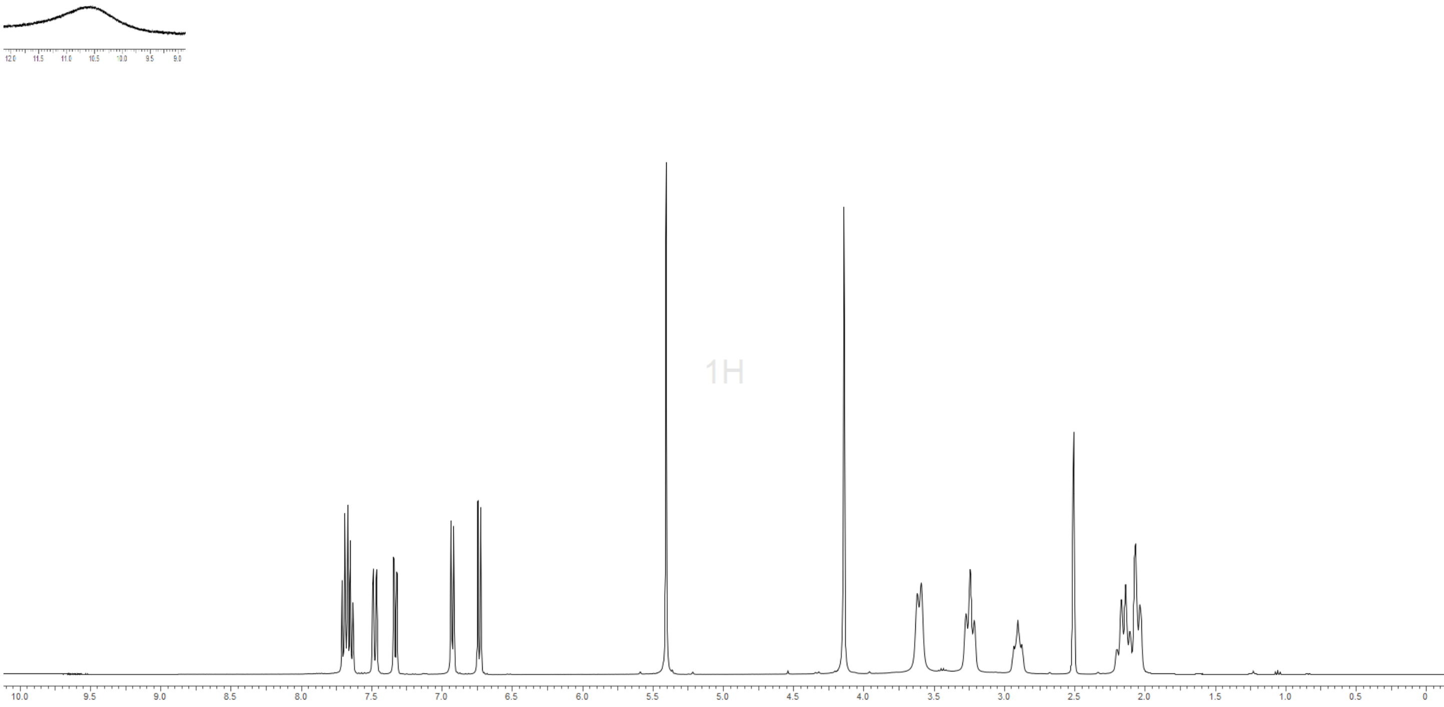
1H
10.0
9.5
9.0
8.5
8.0
7.5
7.0
6.5
6.0
5.5
5.0
4.5
4.0
3.5
3.0
2.5
2.0
1.5
1.0
0.5
0

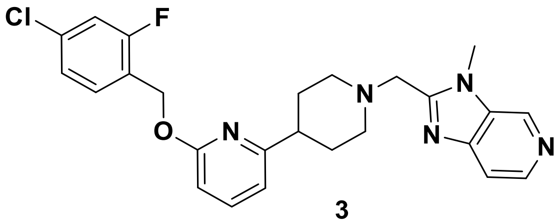


**3**

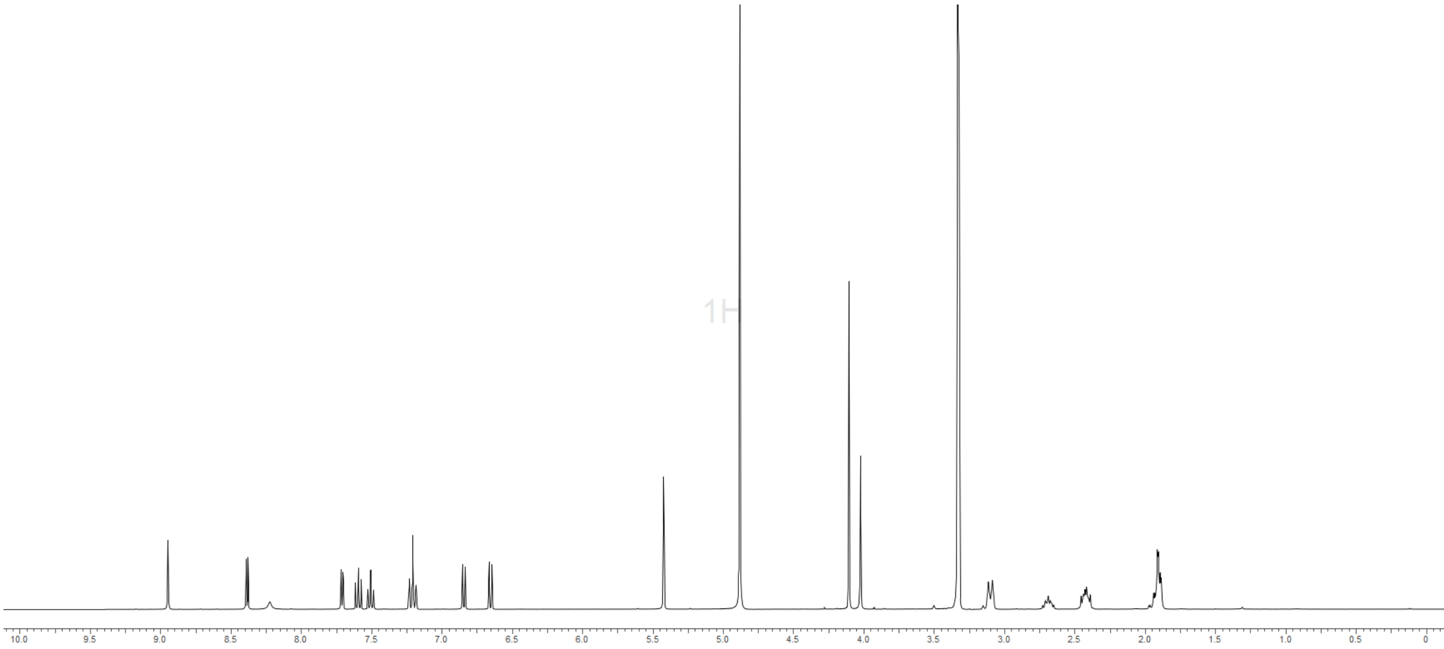


($^{1}$H decoupled)

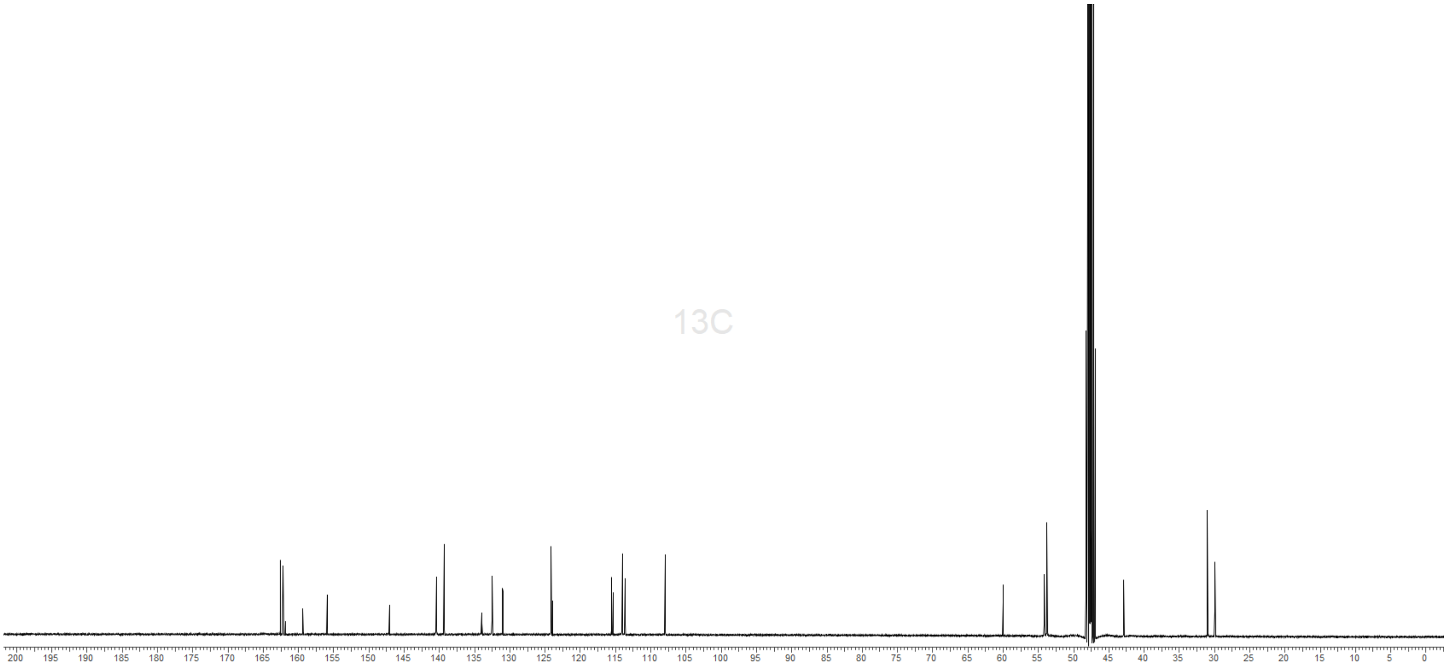

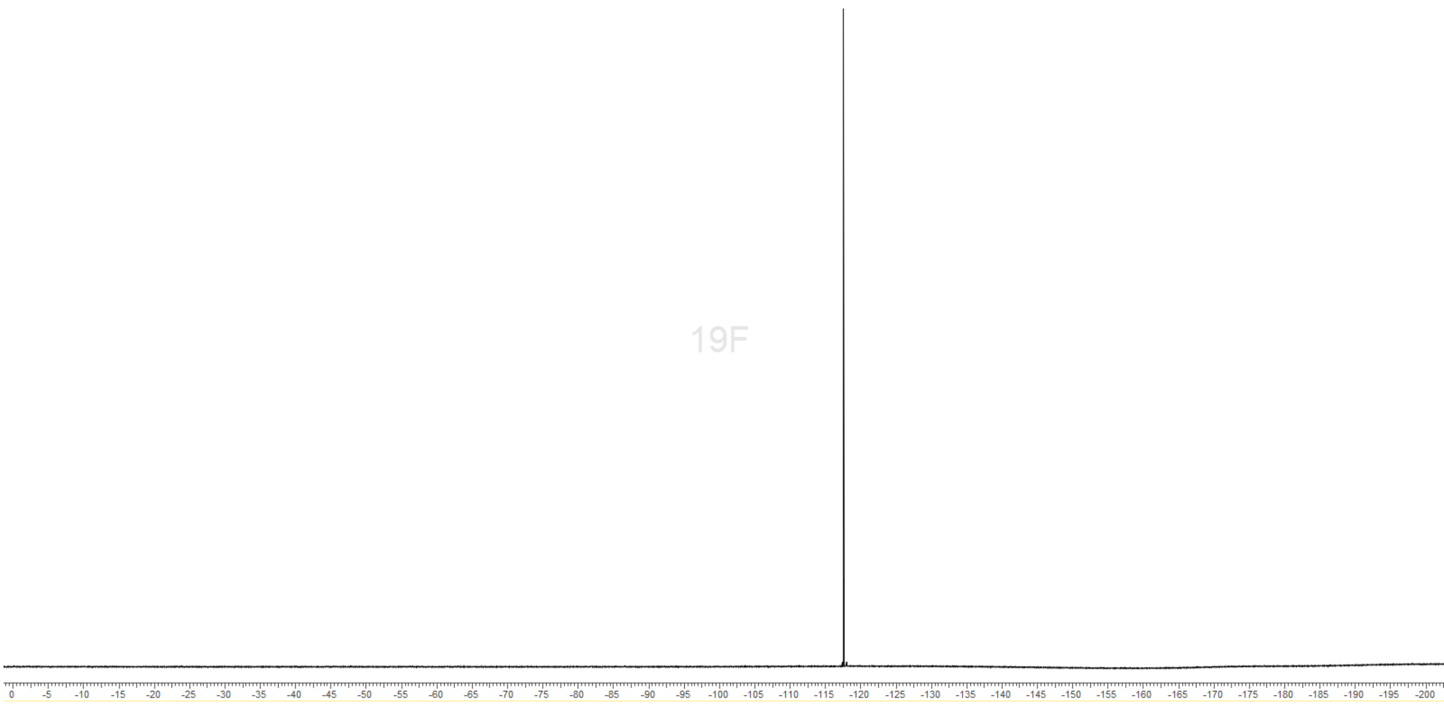
19F
0 -5 -10 -15 -20 -25 -30 -35 -40 -45 -50 -55 -60 -65 -70 -75 -80 -85 -90 -95 -100 -105 -110 -115 -120 -125 -130 -135 -140 -145 -150 -155 -160 -165 -170 -175 -180 -185 -190 -195 -200

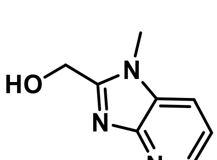
HO
N
N
N

**S20**

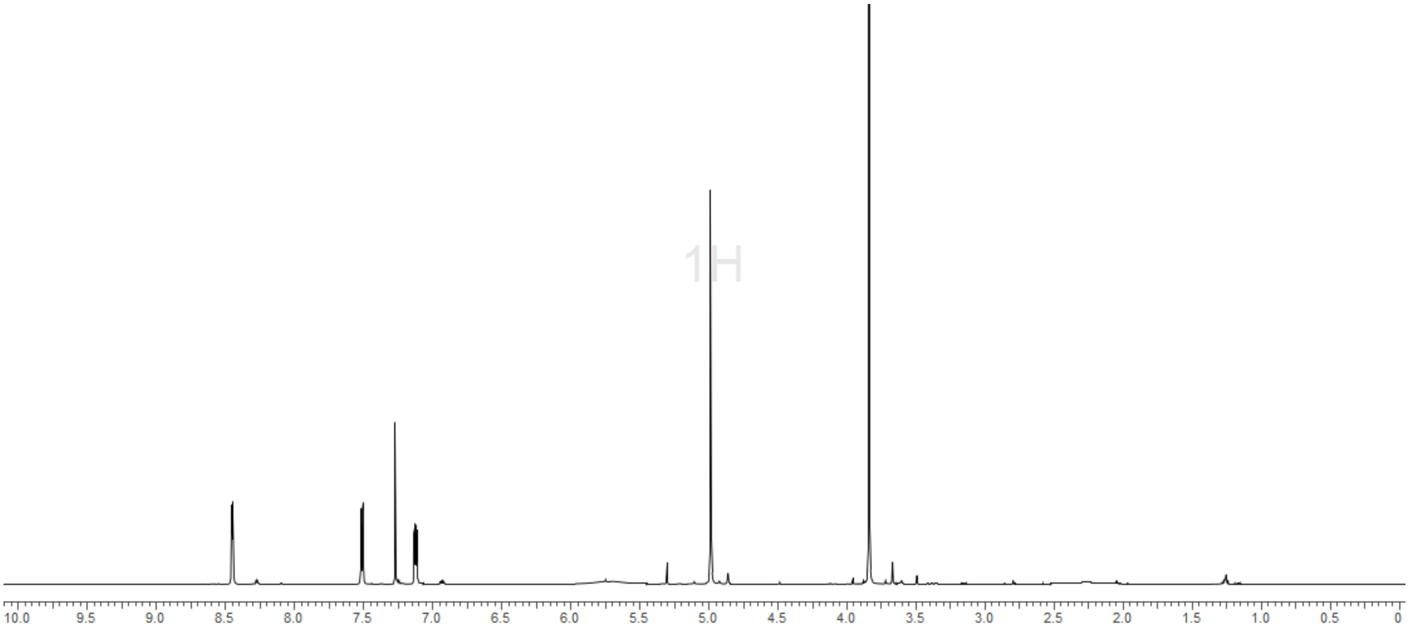
1H
10.0
9.5
9.0
8.5
8.0
7.5
7.0
6.5
6.0
5.5
5.0
4.5
4.0
3.5
3.0
2.5
2.0
1.5
1.0
0.5
0

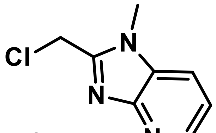
Cl
N
N
N

**S21**

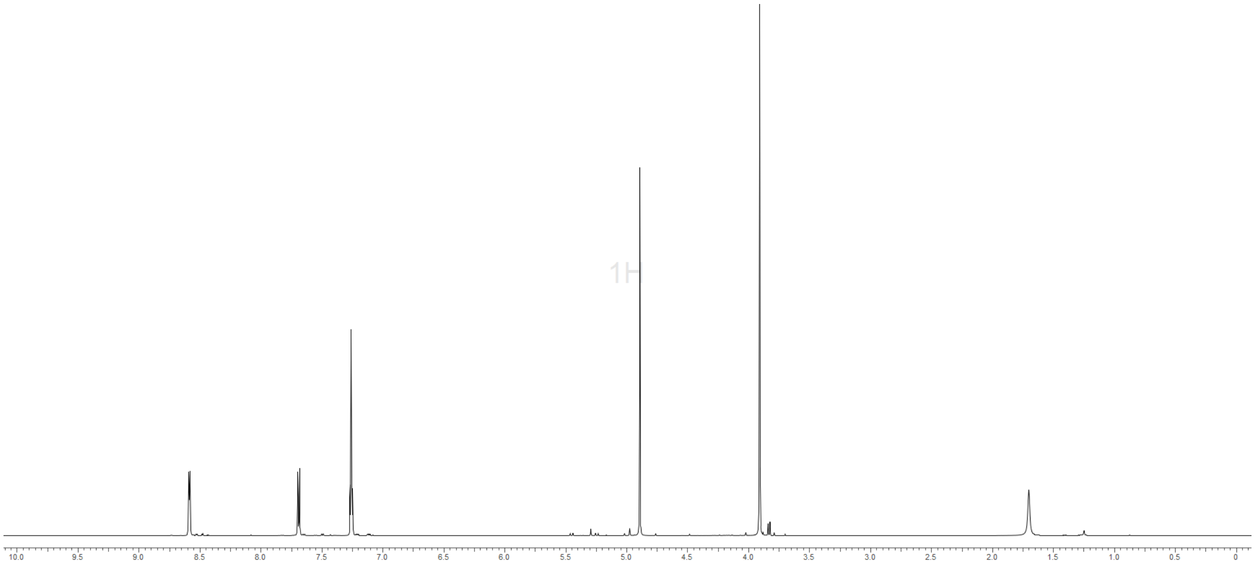
1H

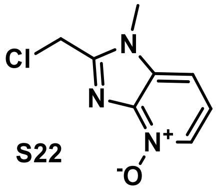
Cl
N
N
N+
-O

S22

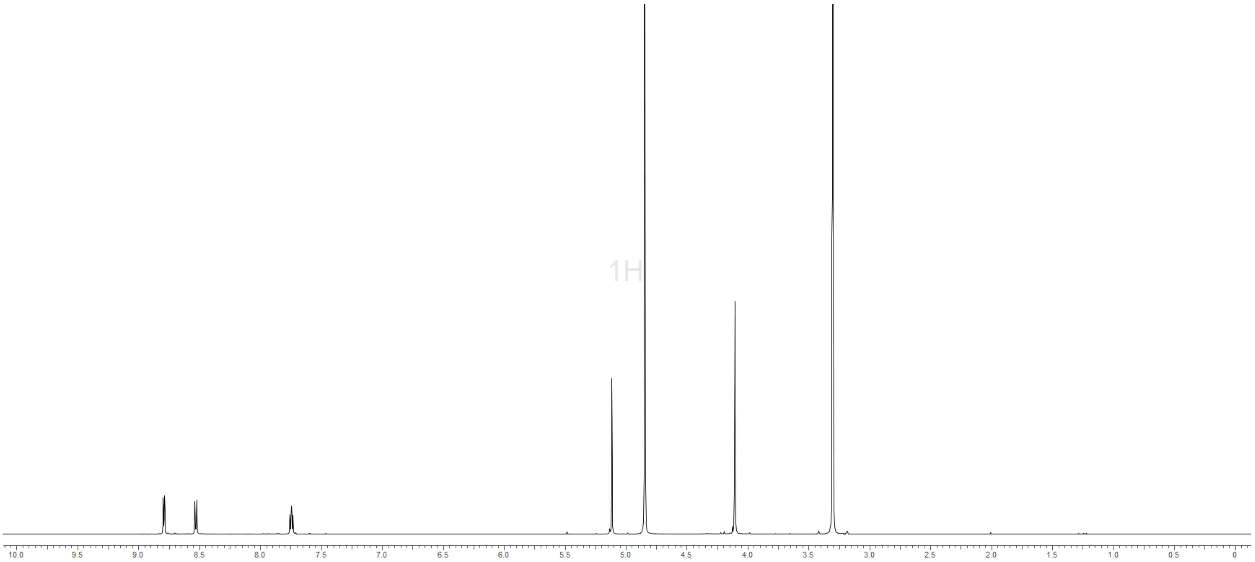
1H

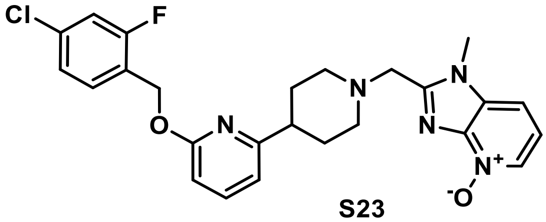
Cl
F
O
N
N
N
N
N+
-O

S23

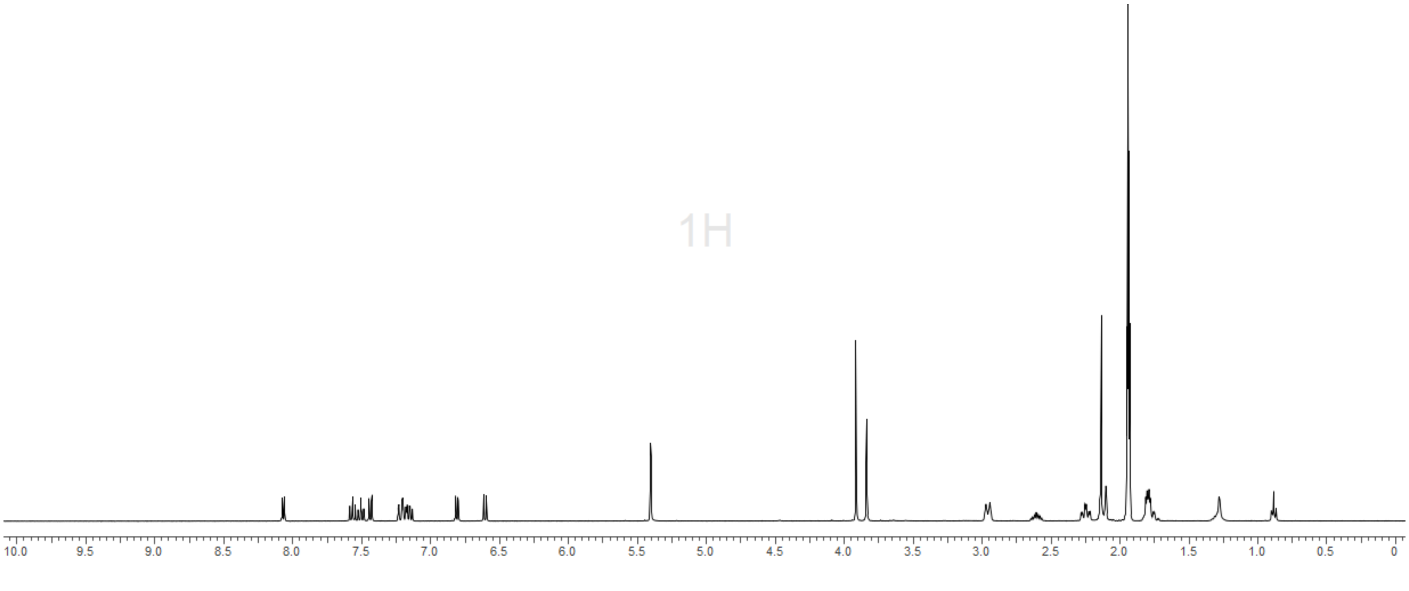
1H
10.0
9.5
9.0
8.5
8.0
7.5
7.0
6.5
6.0
5.5
5.0
4.5
4.0
3.5
3.0
2.5
2.0
1.5
1.0
0.5
0

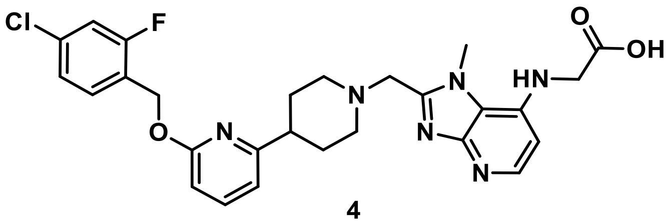

**4**

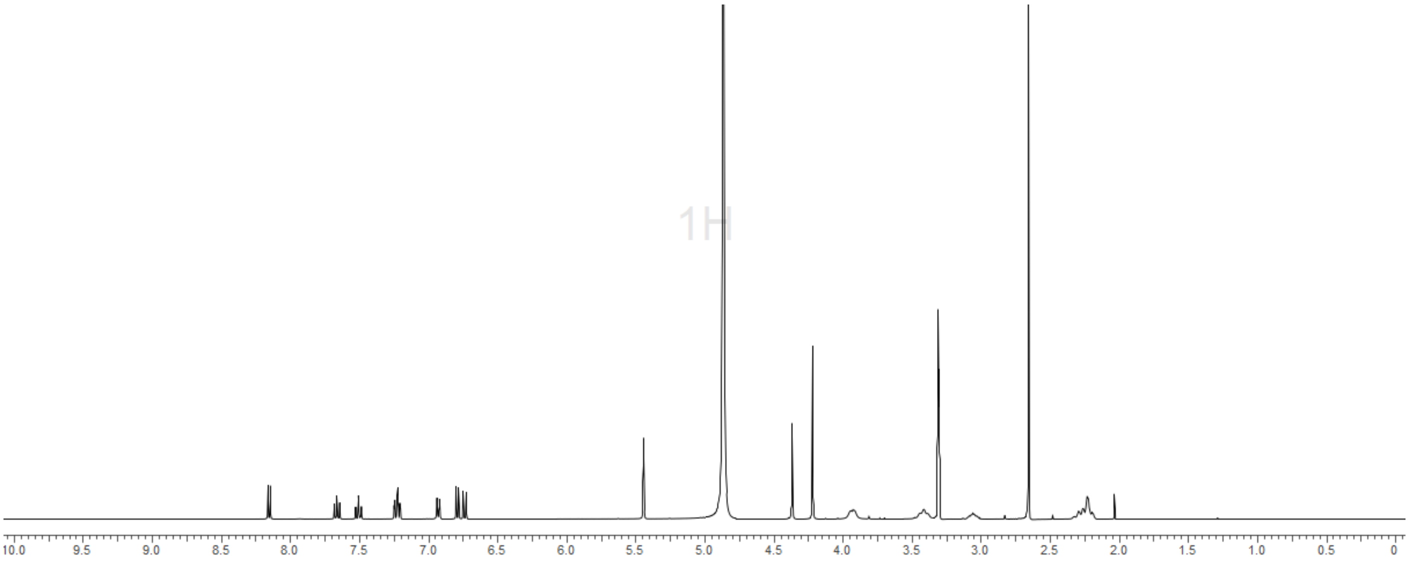


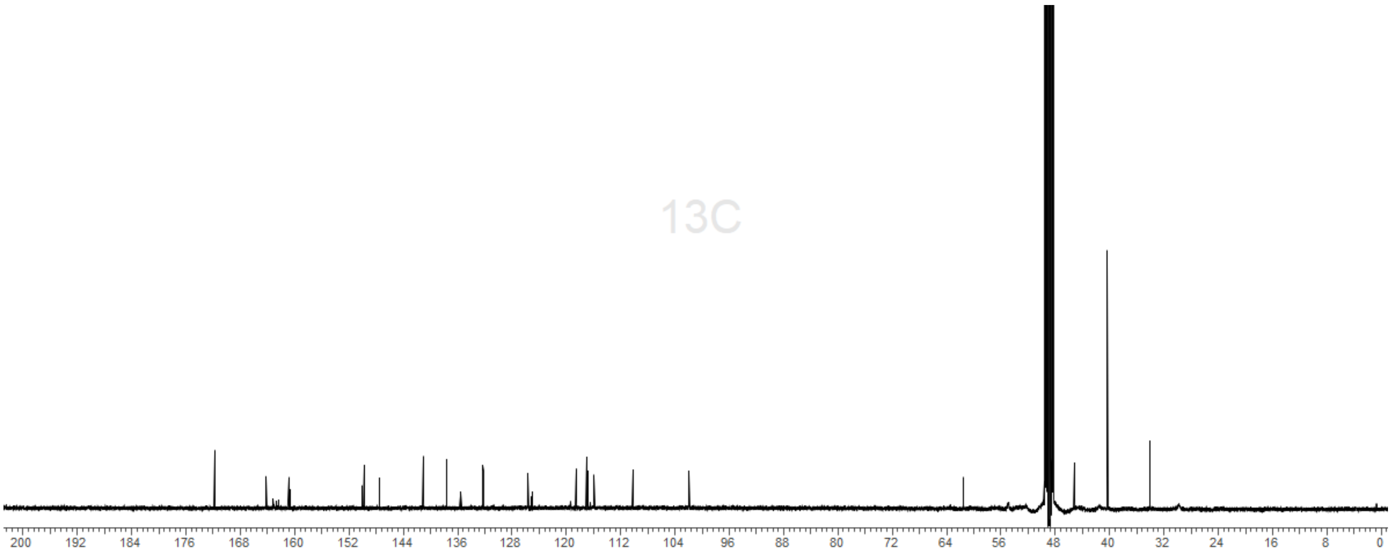

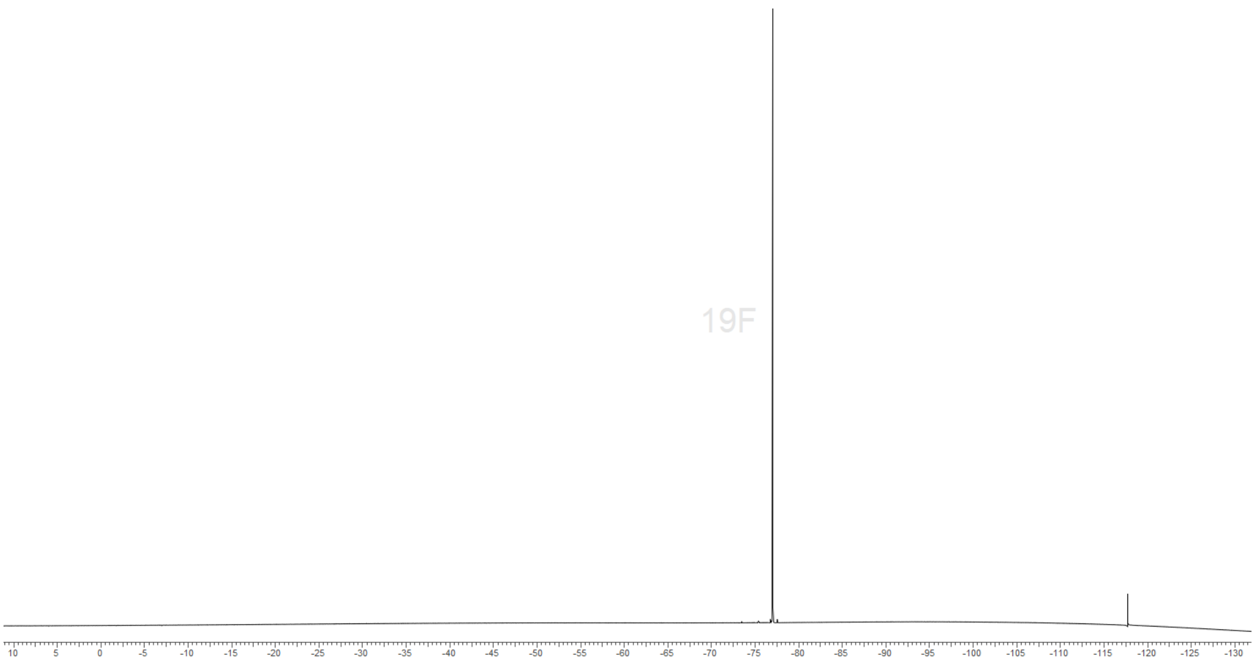
19F
10
5
0
-5
-10
-15
-20
-25
-30
-35
-40
-45
-50
-55
-60
-65
-70
-75
-80
-85
-90
-95
-100
-105
-110
-115
-120
-125
-130

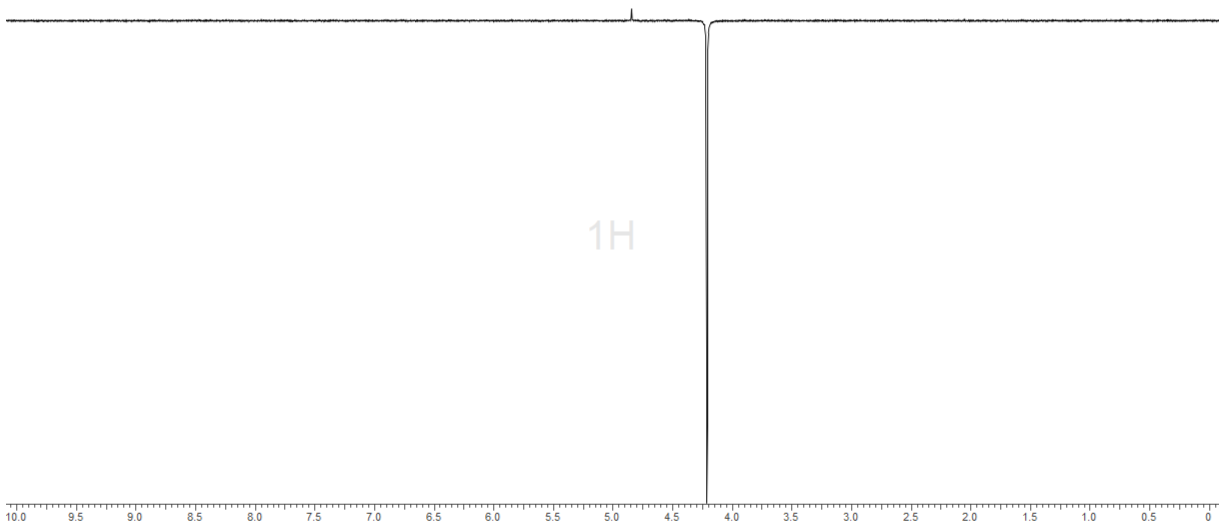
1H
10.0
9.5
9.0
8.5
8.0
7.5
7.0
6.5
6.0
5.5
5.0
4.5
4.0
3.5
3.0
2.5
2.0
1.5
1.0
0.5
0

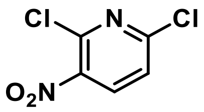
Cl
N
Cl
O2N

**S25**

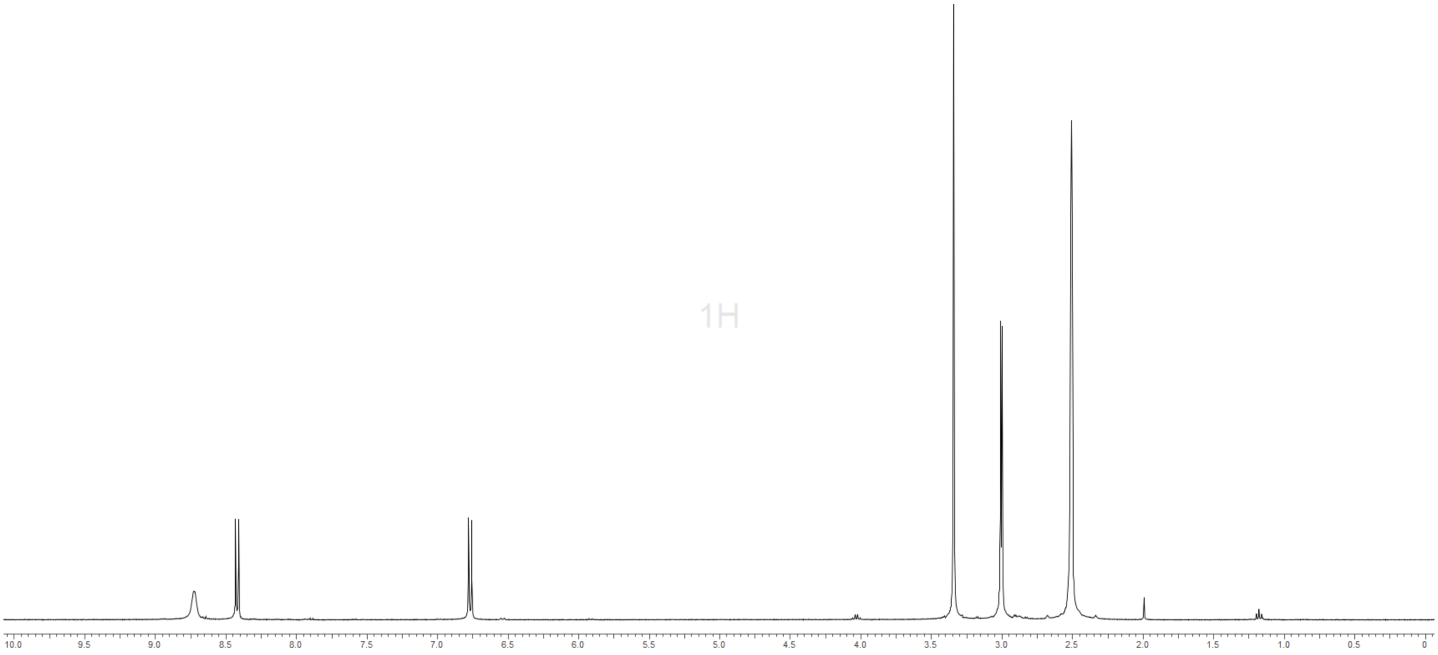
1H
10.0
9.5
9.0
8.5
8.0
7.5
7.0
6.5
6.0
5.5
5.0
4.5
4.0
3.5
3.0
2.5
2.0
1.5
1.0
0.5
0

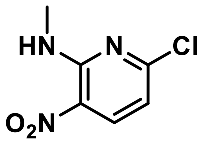

**S26**

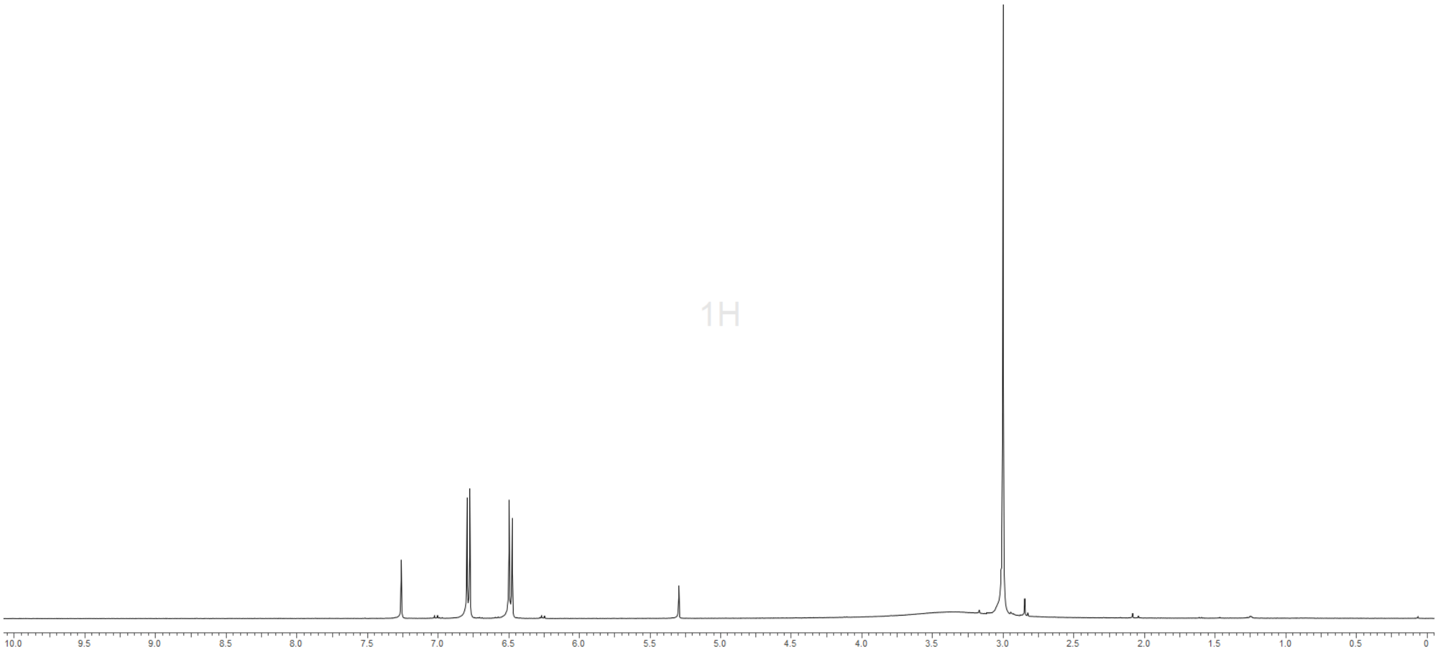

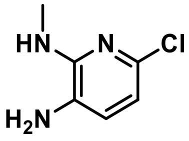

**S27**

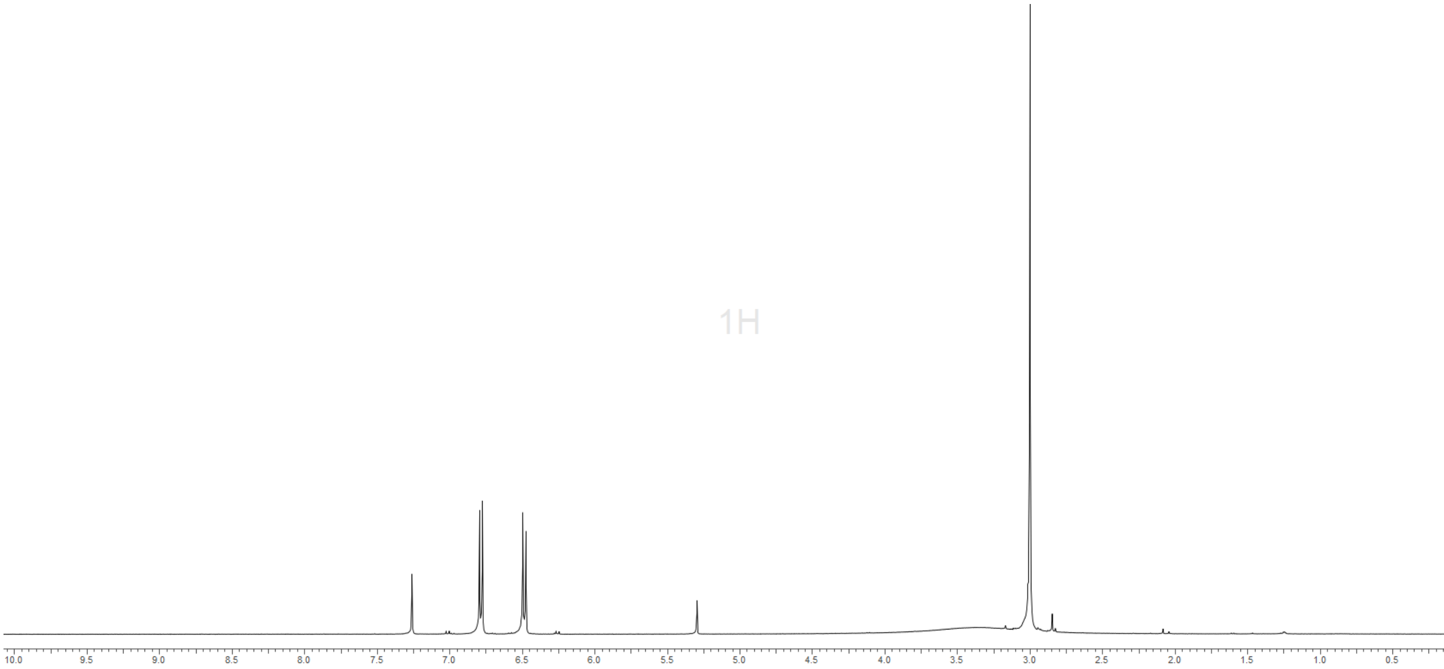

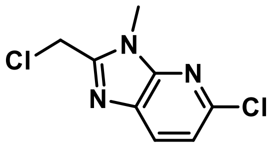
Cl
N
N
N
Cl

**S29**

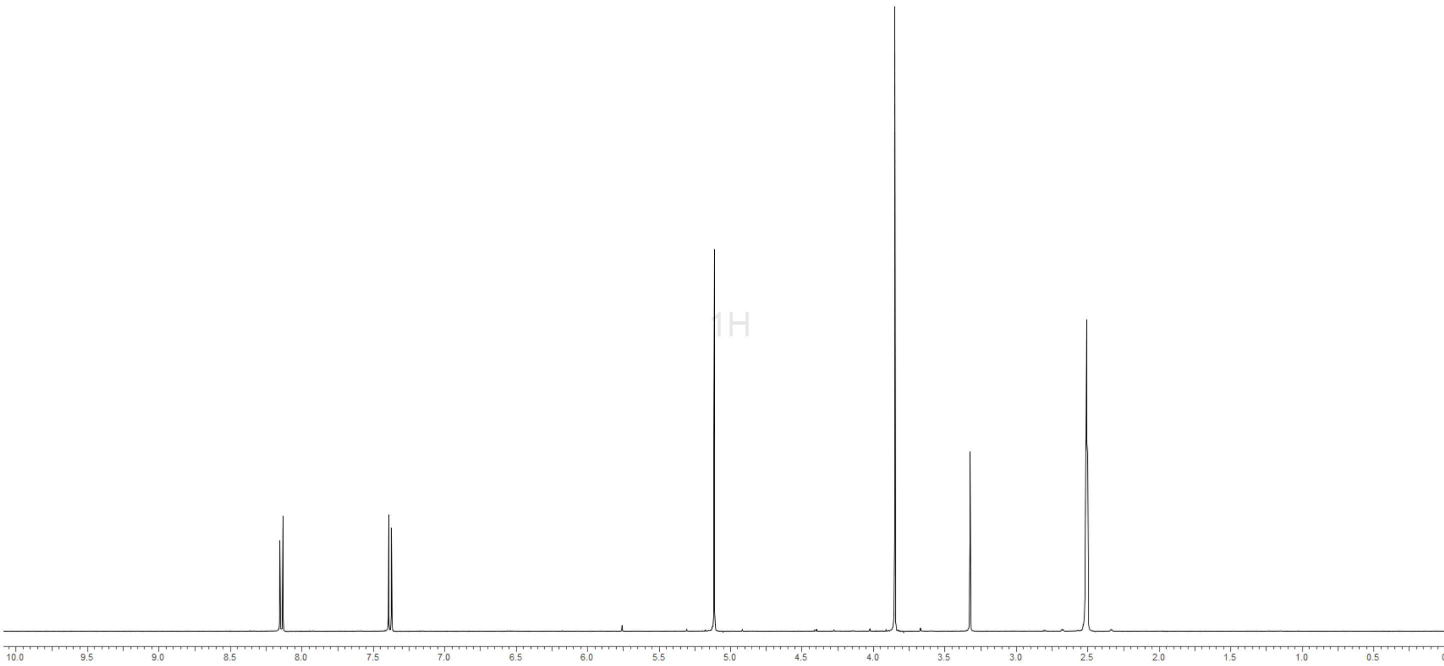
1H
10.0
9.5
9.0
8.5
8.0
7.5
7.0
6.5
6.0
5.5
5.0
4.5
4.0
3.5
3.0
2.5
2.0
1.5
1.0
0.5

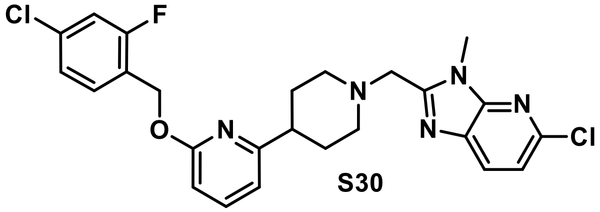

**S30**

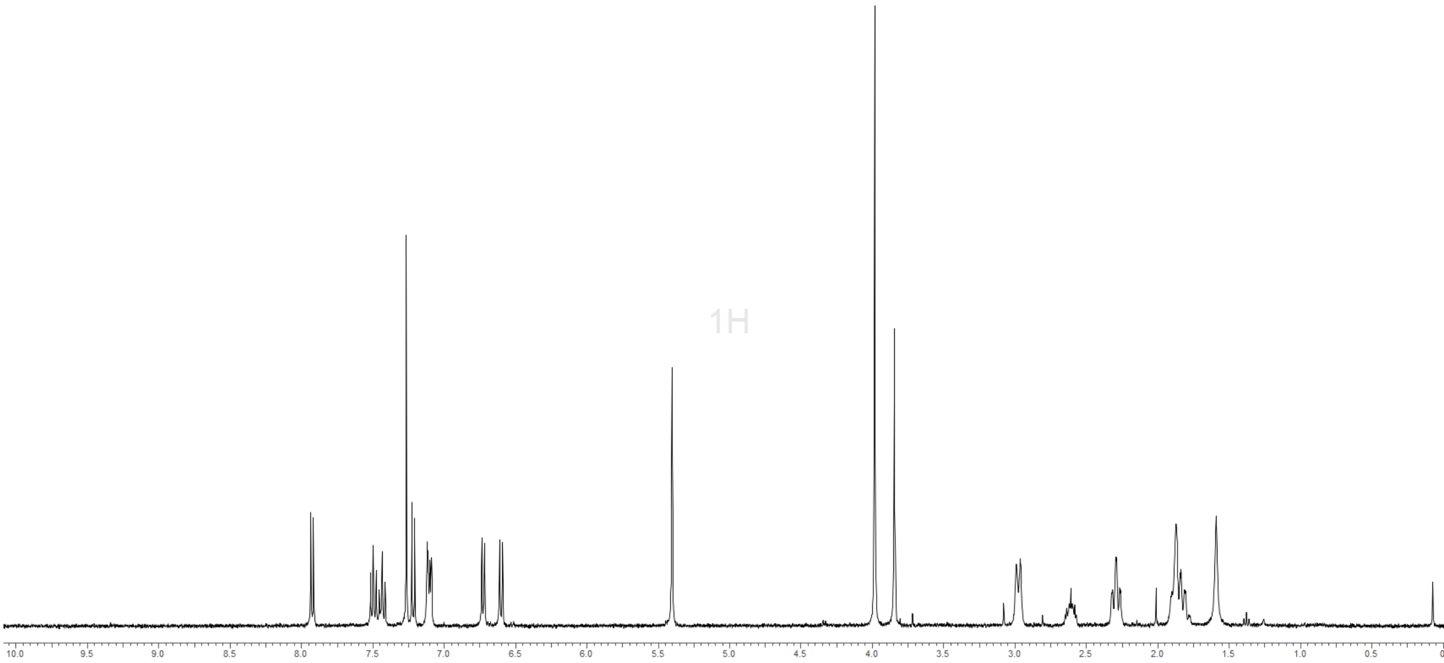

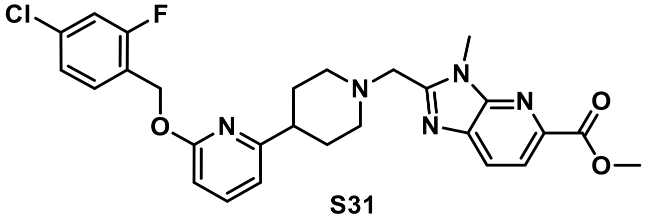
Cl
F
O
N
N
N
N
N
O
O

**S31**

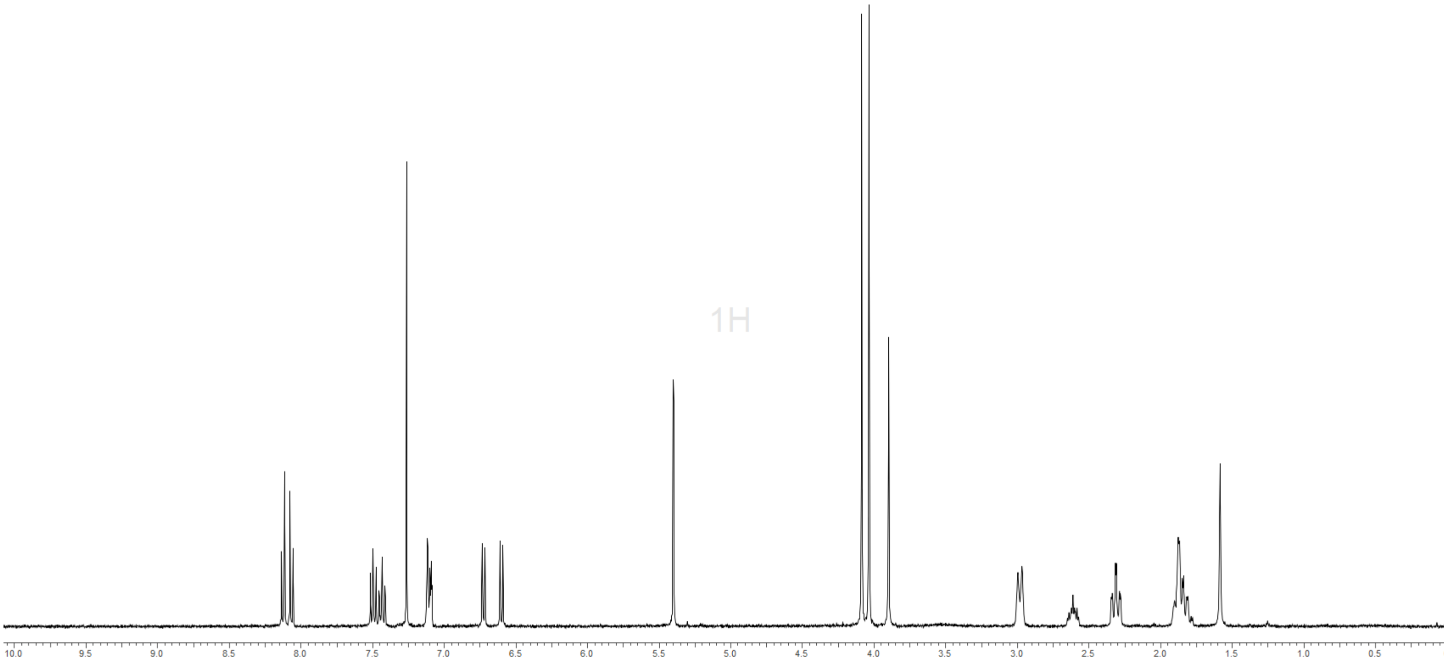
1H
10.0
9.5
9.0
8.5
8.0
7.5
7.0
6.5
6.0
5.5
5.0
4.5
4.0
3.5
3.0
2.5
2.0
1.5
1.0
0.5

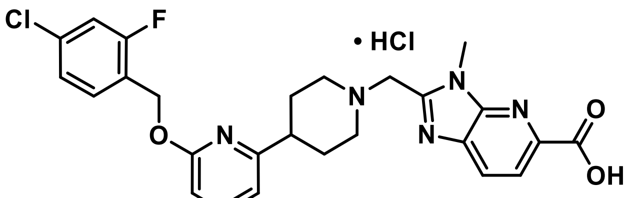


**5**

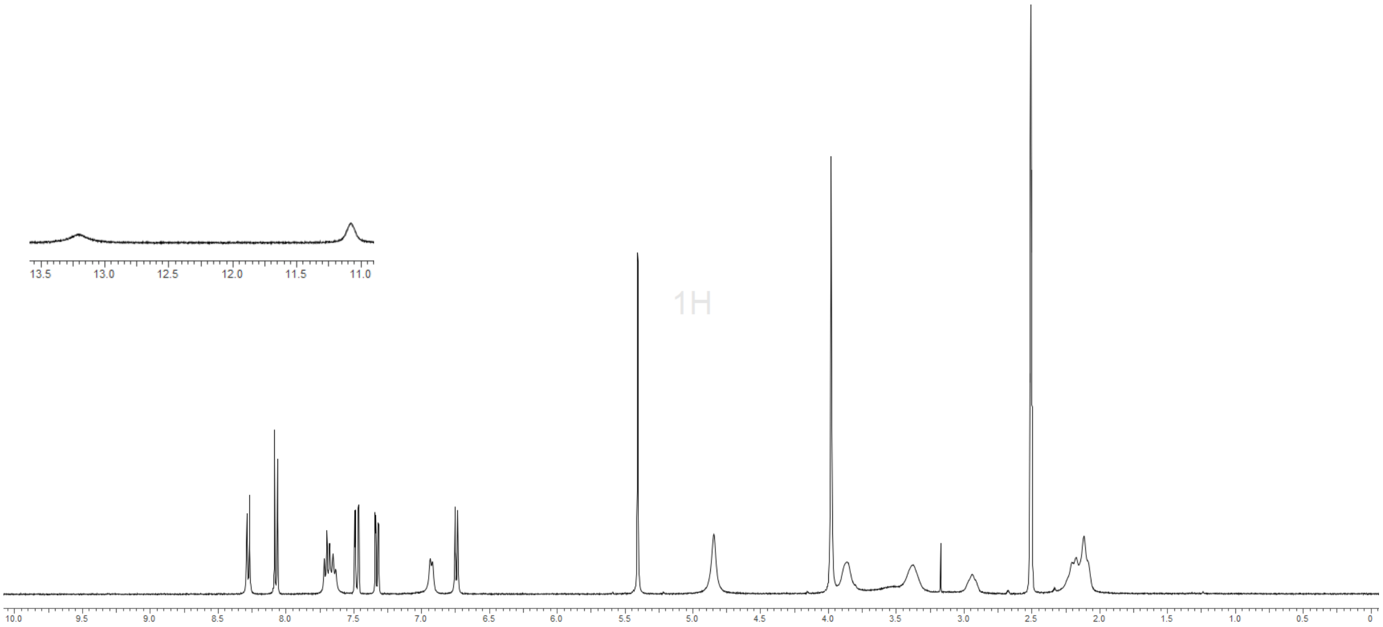


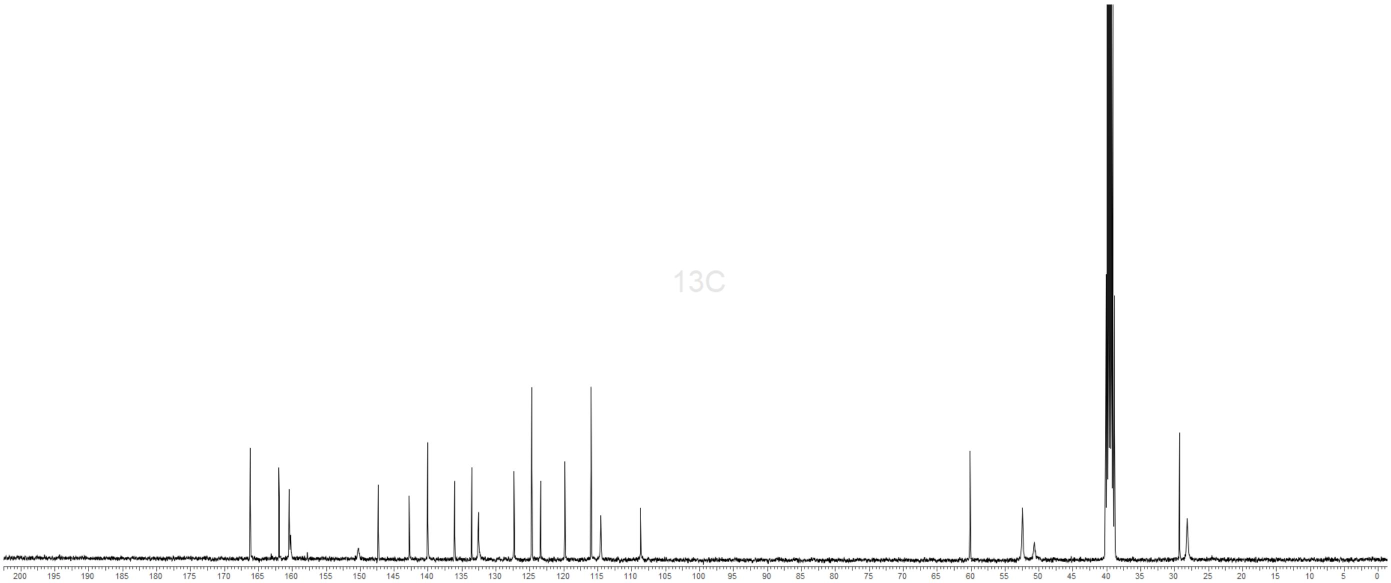

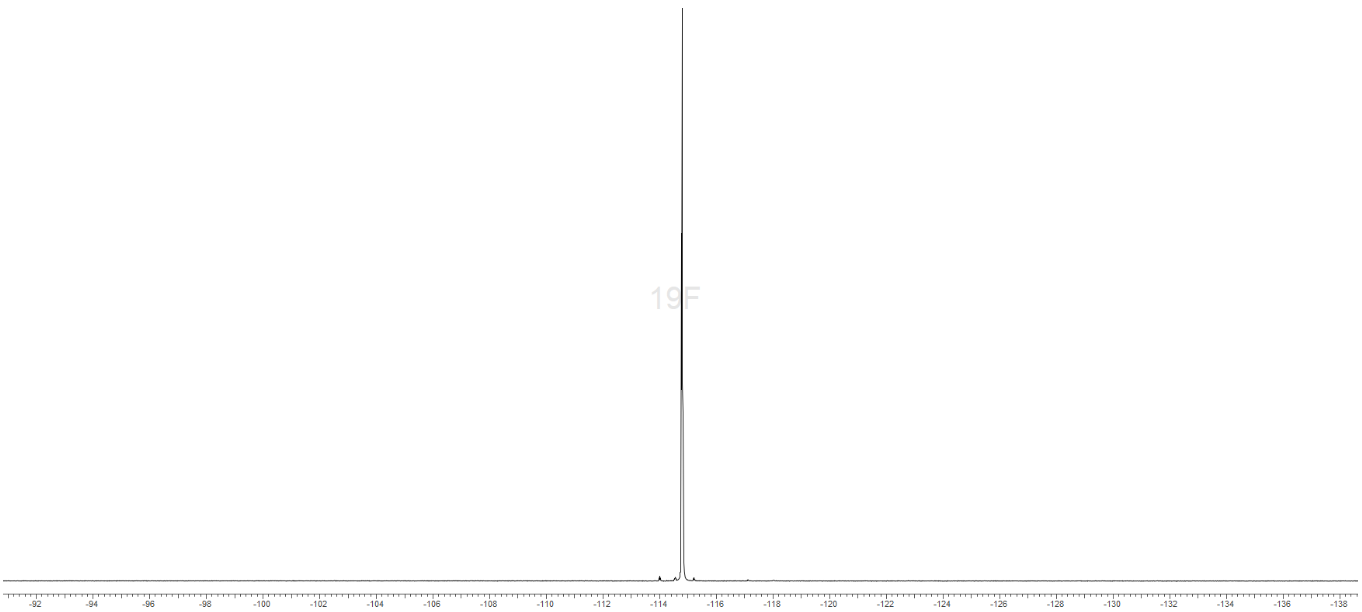
19F
-92
-94
-96
-98
-100
-102
-104
-106
-108
-110
-112
-114
-116
-118
-120
-122
-124
-126
-128
-130
-132
-134
-136
-138

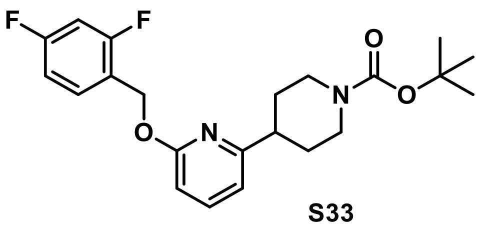
F
F
O
N
N
O
O

**S33**

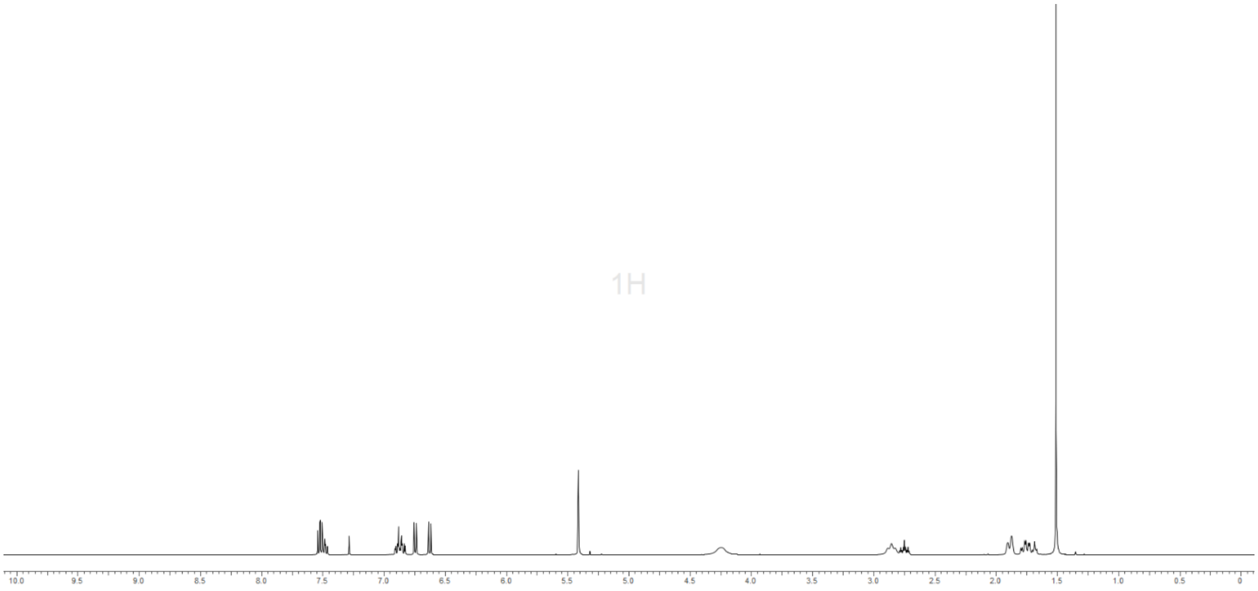
1H
10.0 9.5 9.0 8.5 8.0 7.5 7.0 6.5 6.0 5.5 5.0 4.5 4.0 3.5 3.0 2.5 2.0 1.5 1.0 0.5 0

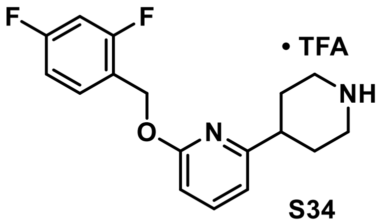
F
F
• TFA
NH
O
N

**S34**

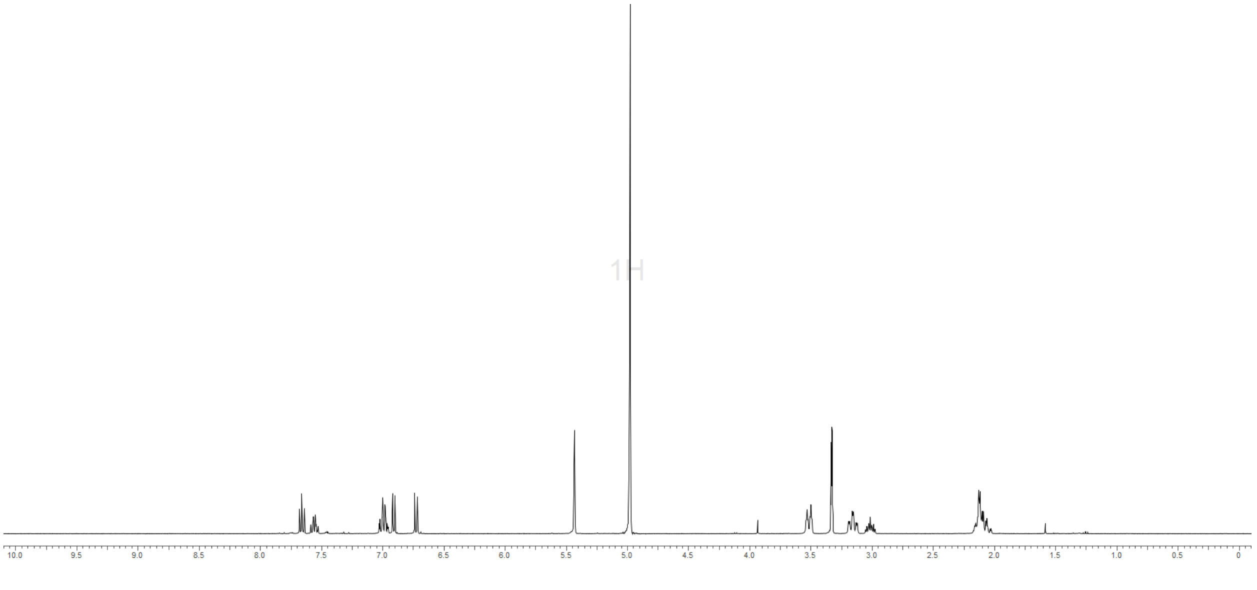
1H
10.0 9.5 9.0 8.5 8.0 7.5 7.0 6.5 6.0 5.5 5.0 4.5 4.0 3.5 3.0 2.5 2.0 1.5 1.0 0.5 0

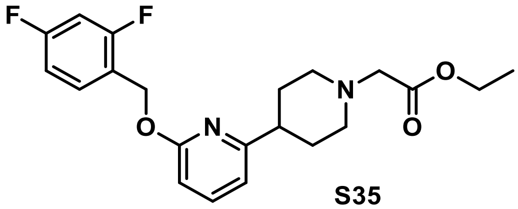

**S35**

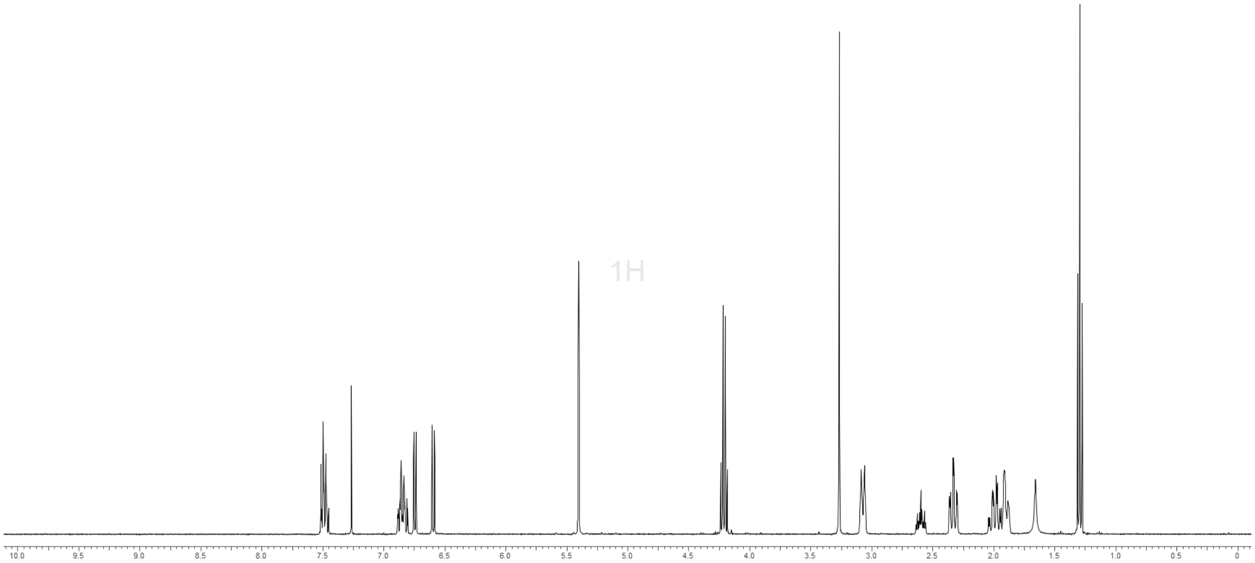

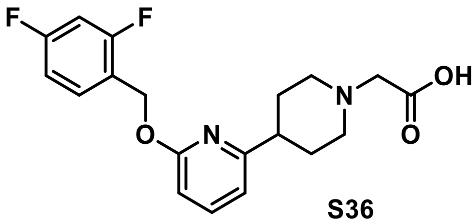
F
F
O
N
N
OH
O

**S36**

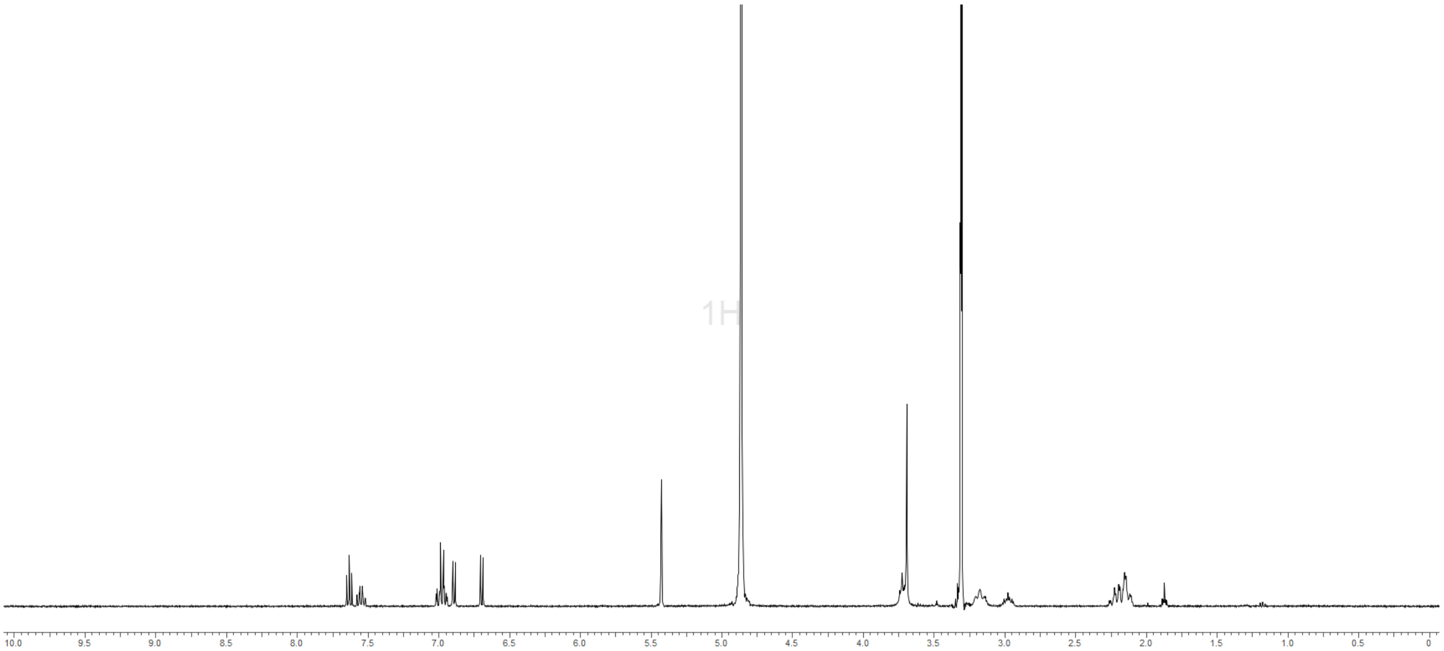
1H
10.0
9.5
9.0
8.5
8.0
7.5
7.0
6.5
6.0
5.5
5.0
4.5
4.0
3.5
3.0
2.5
2.0
1.5
1.0
0.5
0

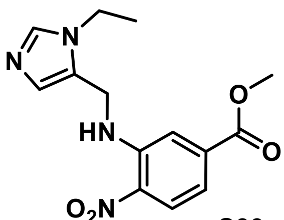
N
N
HN
O
O
$O_2N$

S39

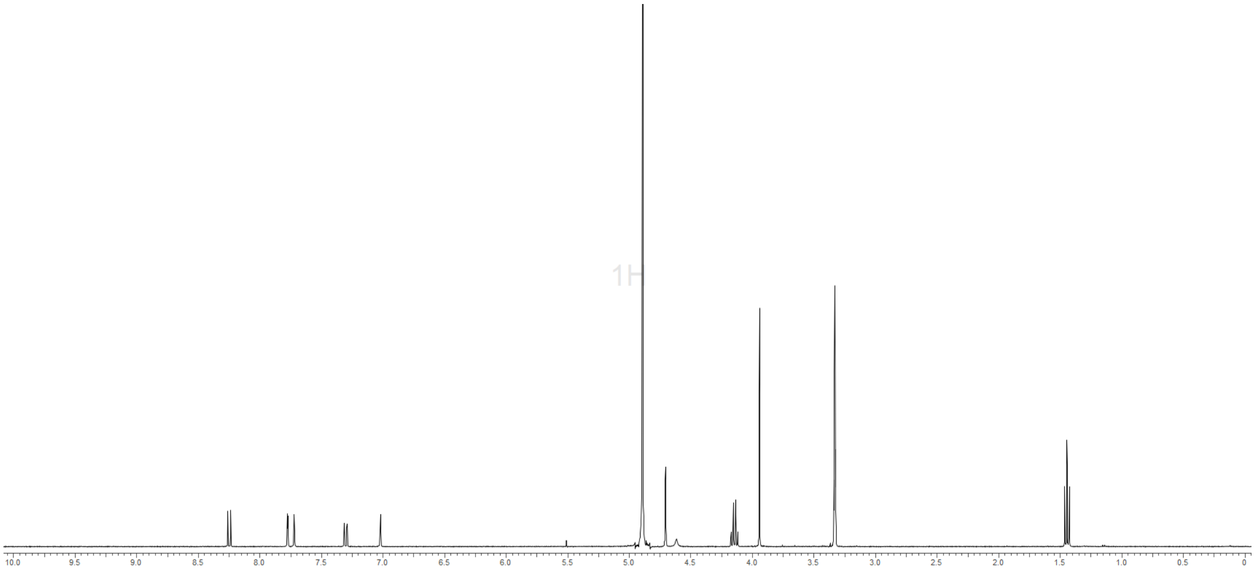
1H
10.0 9.5 9.0 8.5 8.0 7.5 7.0 6.5 6.0 5.5 5.0 4.5 4.0 3.5 3.0 2.5 2.0 1.5 1.0 0.5 0

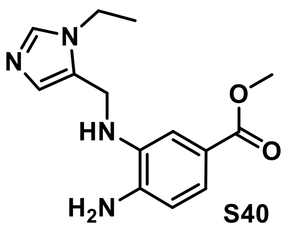
N
N
HN
O
O
$H_2N$

S40

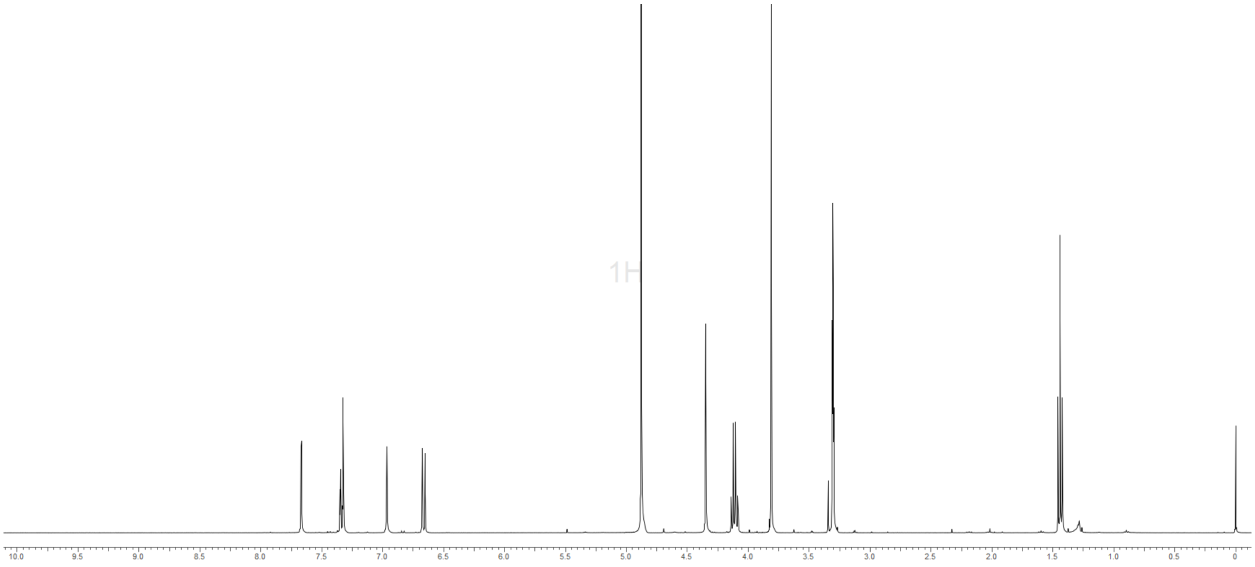
1H
10.0 9.5 9.0 8.5 8.0 7.5 7.0 6.5 6.0 5.5 5.0 4.5 4.0 3.5 3.0 2.5 2.0 1.5 1.0 0.5 0

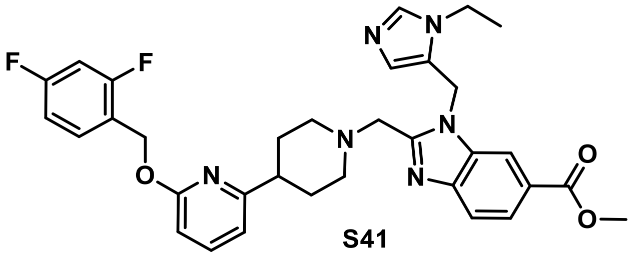

**S41**

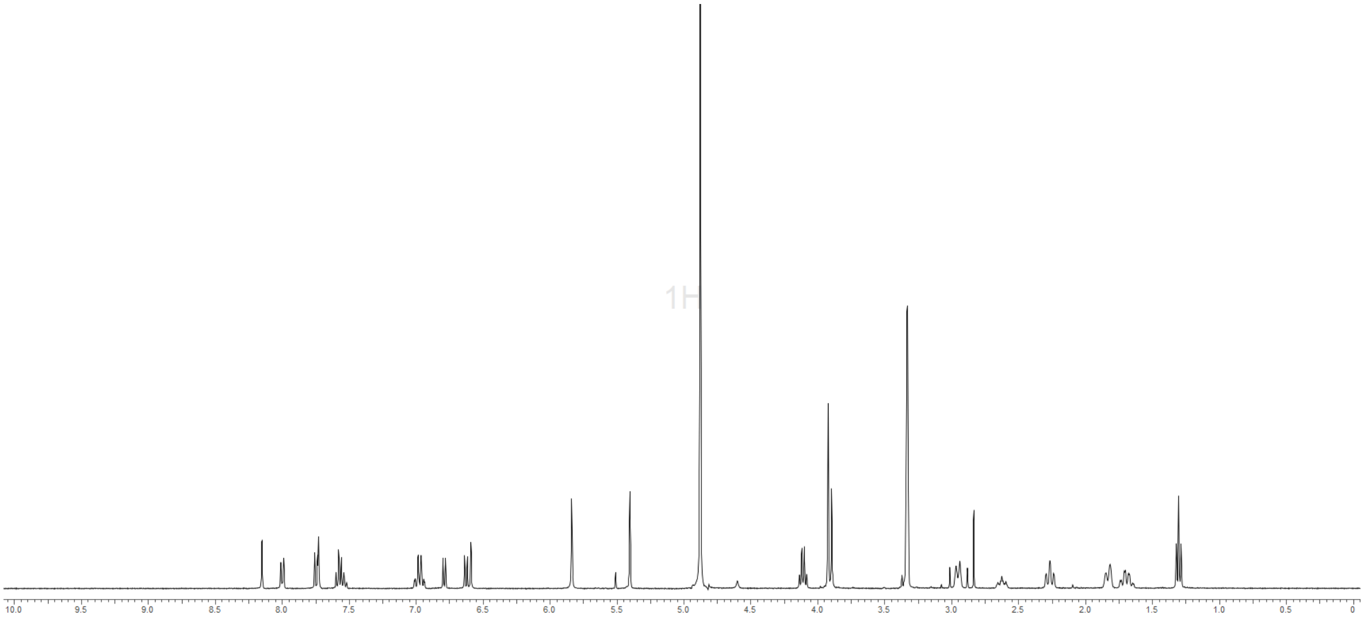


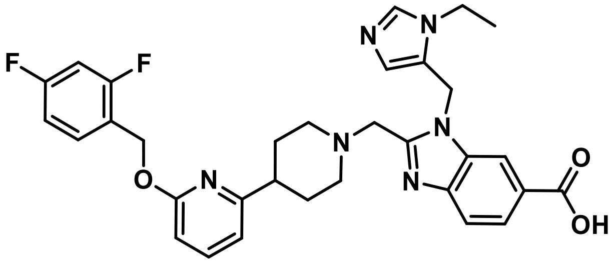

**PF-06883365**

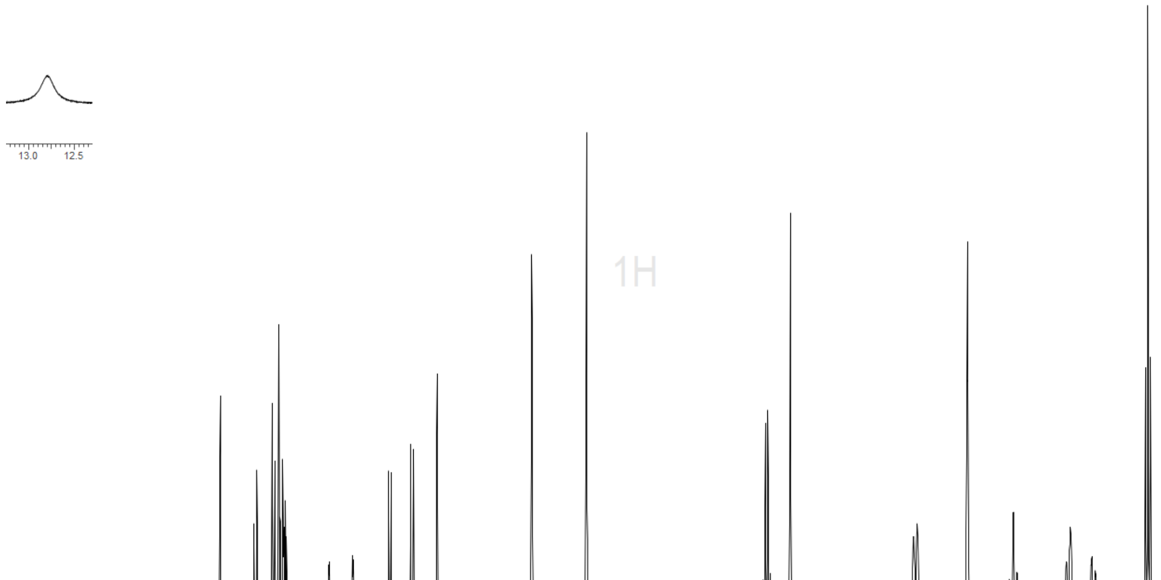

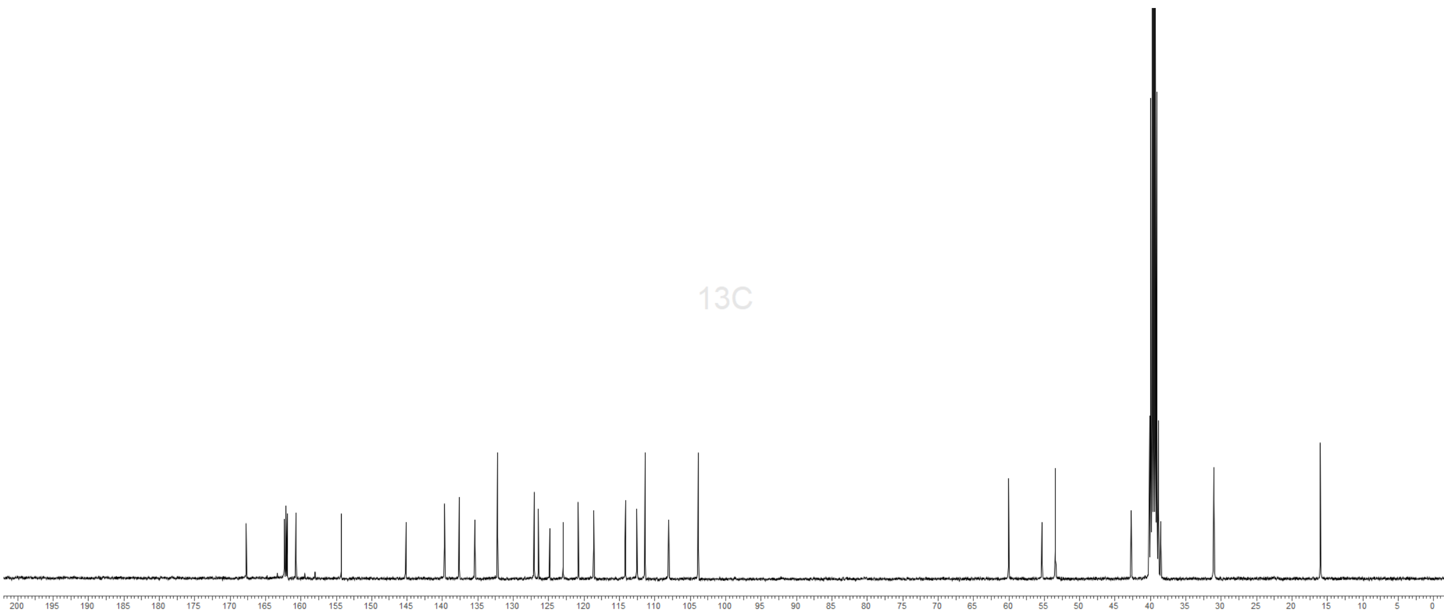
13C
200
195
190
185
180
175
170
165
160
155
150
145
140
135
130
125
120
115
110
105
100
95
90
85
80
75
70
65
60
55
50
45
40
35
30
25
20
15
10
5
0

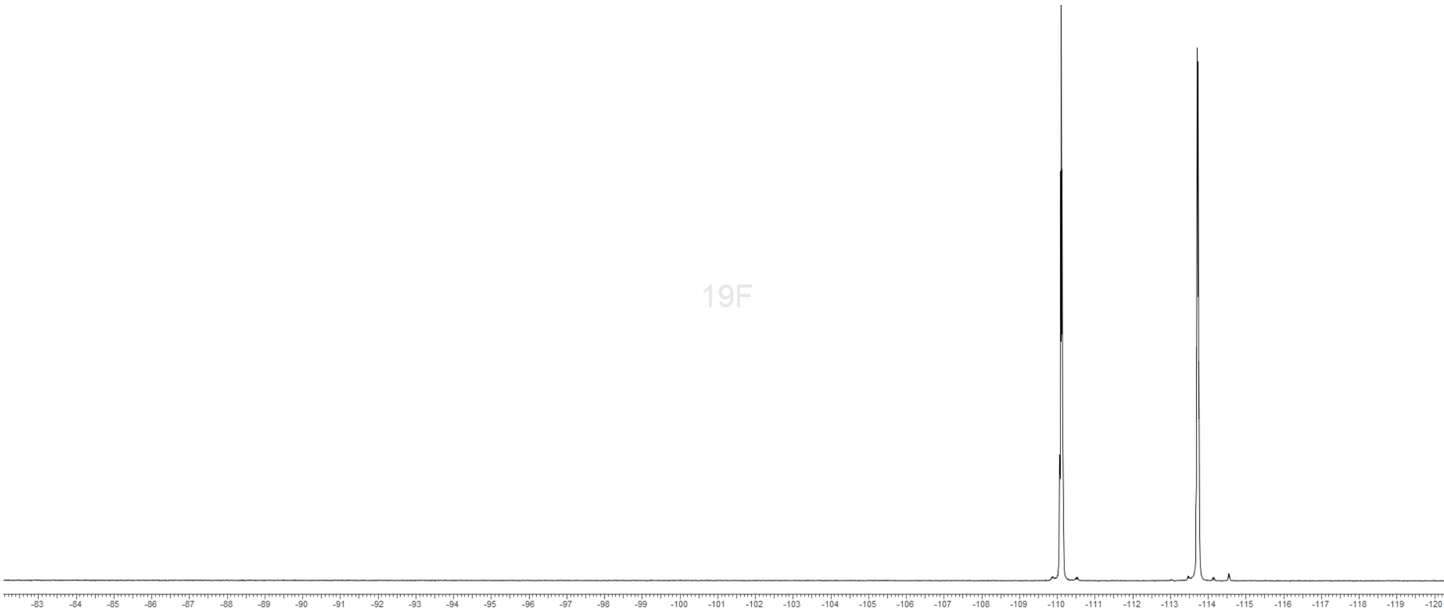
19F
-83
-84
-85
-86
-87
-88
-89
-90
-91
-92
-93
-94
-95
-96
-97
-98
-99
-100
-101
-102
-103
-104
-105
-106
-107
-108
-109
-110
-111
-112
-113
-114
-115
-116
-117
-118
-119
-120

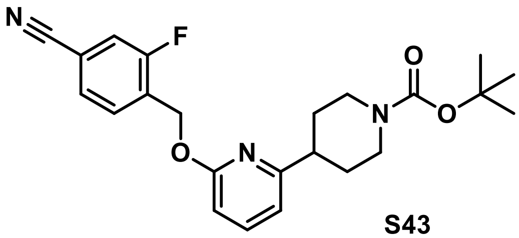

**S43**

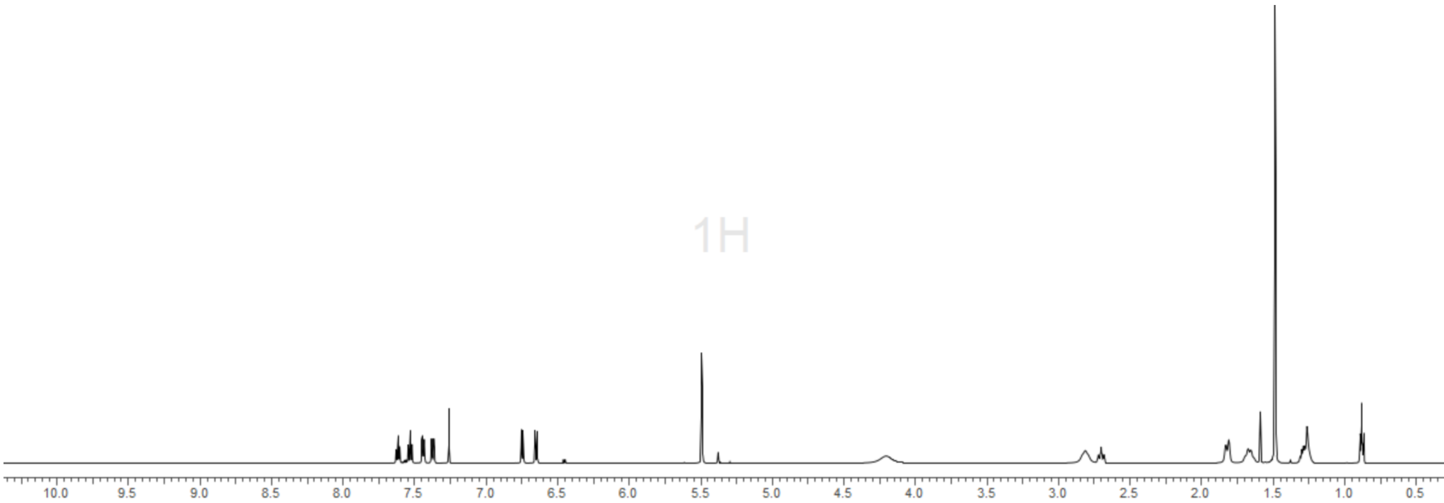


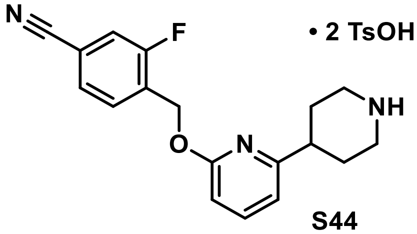

**• 2 TsOH**

**S44**

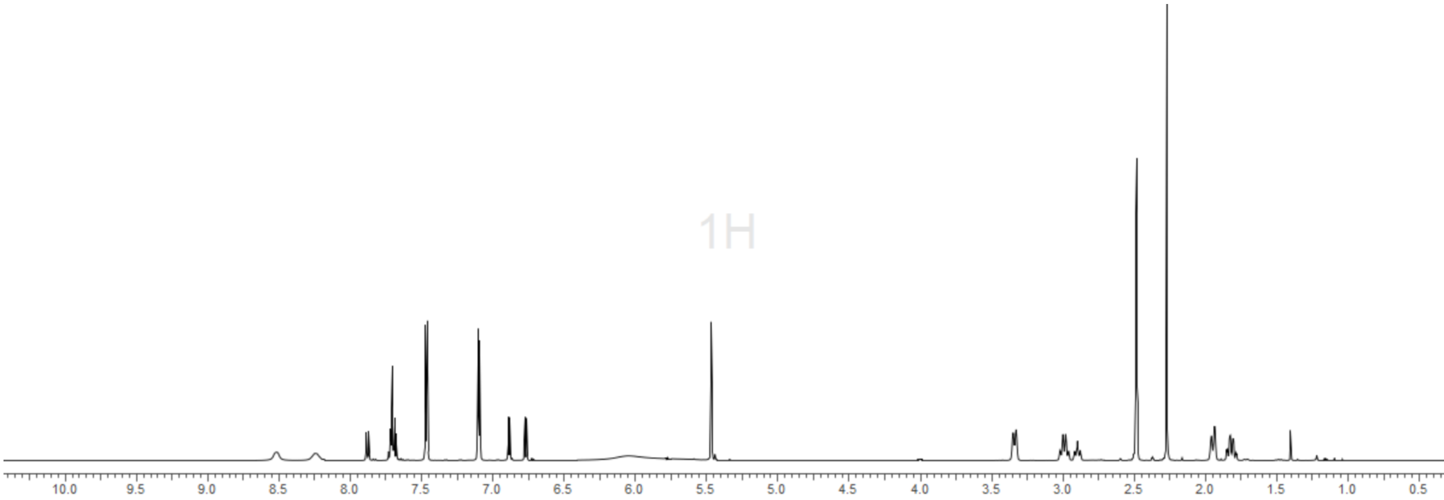

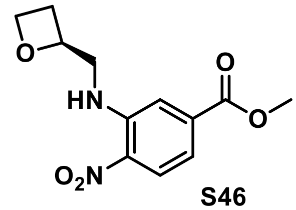
O
HN
O
O
$O_2N$

**S46**

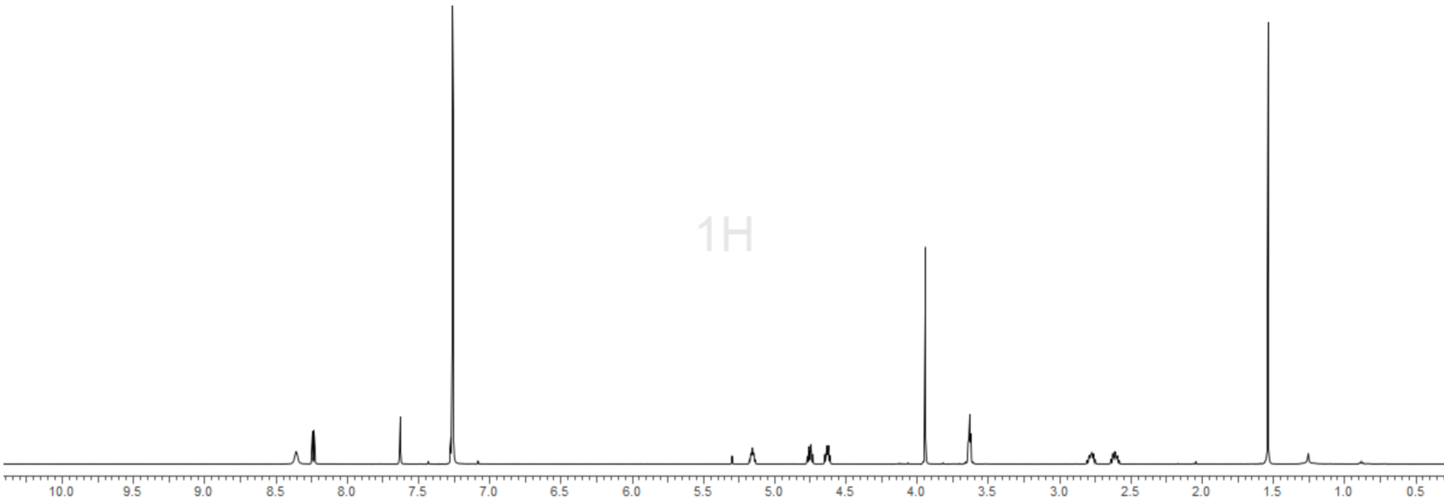
1H
10.0 9.5 9.0 8.5 8.0 7.5 7.0 6.5 6.0 5.5 5.0 4.5 4.0 3.5 3.0 2.5 2.0 1.5 1.0 0.5

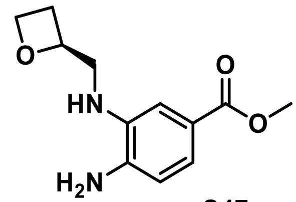
O
HN
O
O
$H_2N$

**S47**

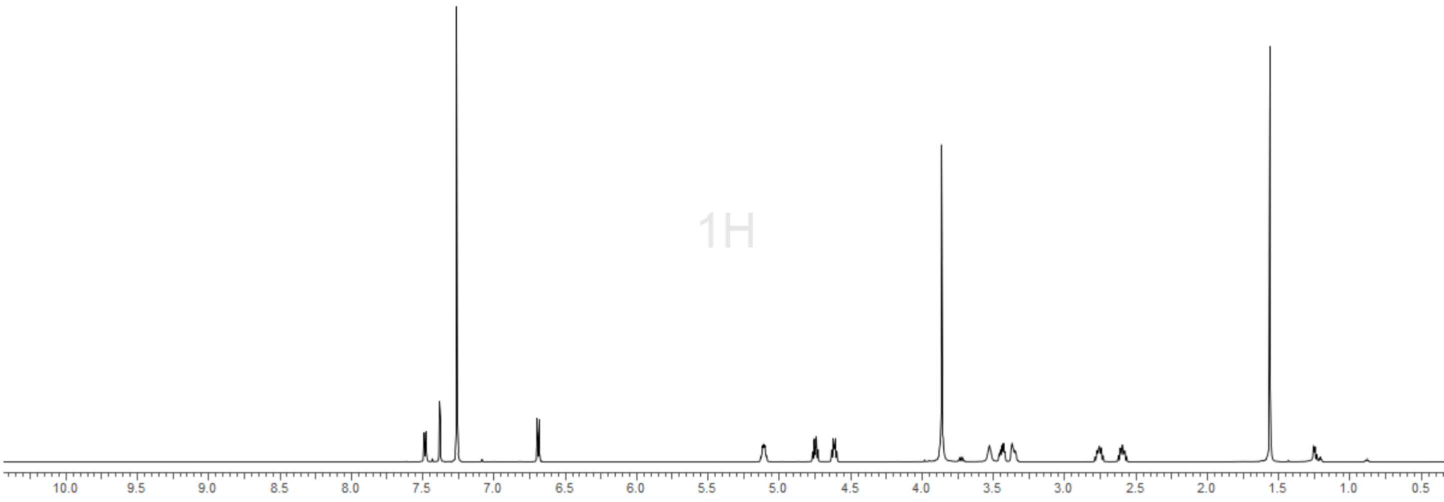
1H
10.0 9.5 9.0 8.5 8.0 7.5 7.0 6.5 6.0 5.5 5.0 4.5 4.0 3.5 3.0 2.5 2.0 1.5 1.0 0.5

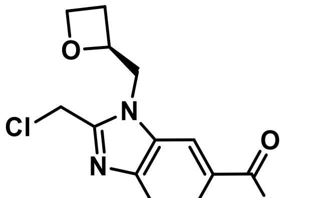

S49

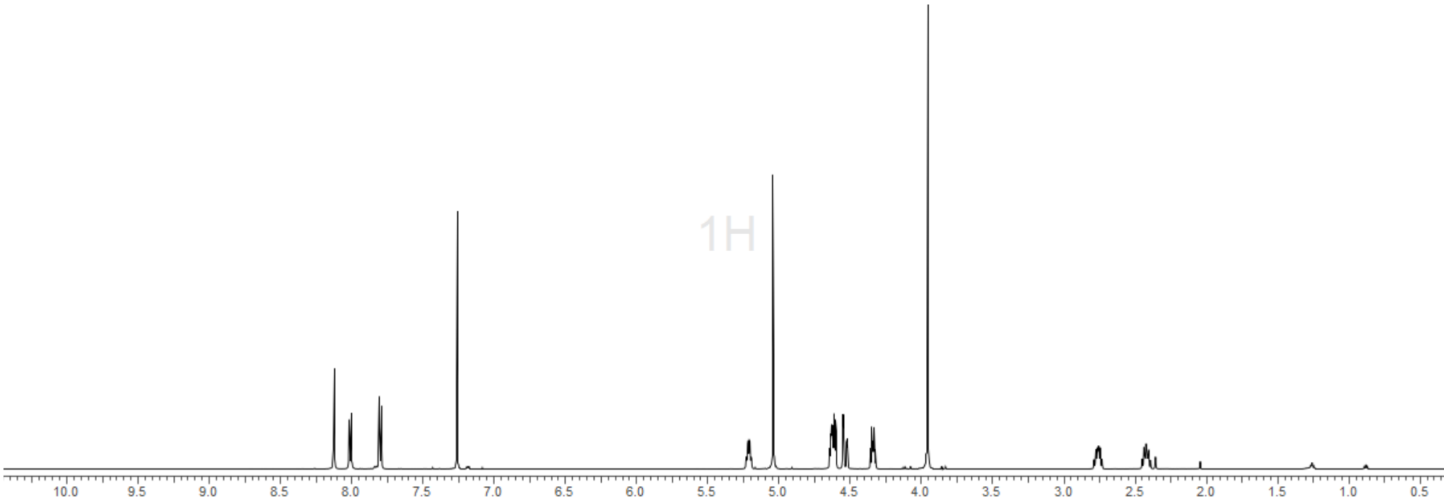


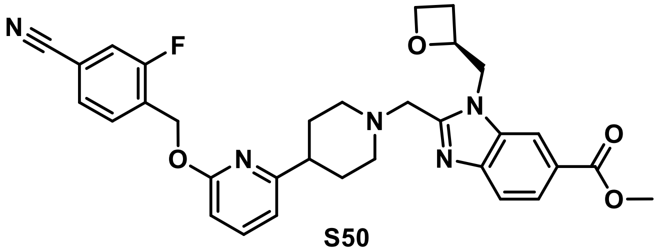

S50

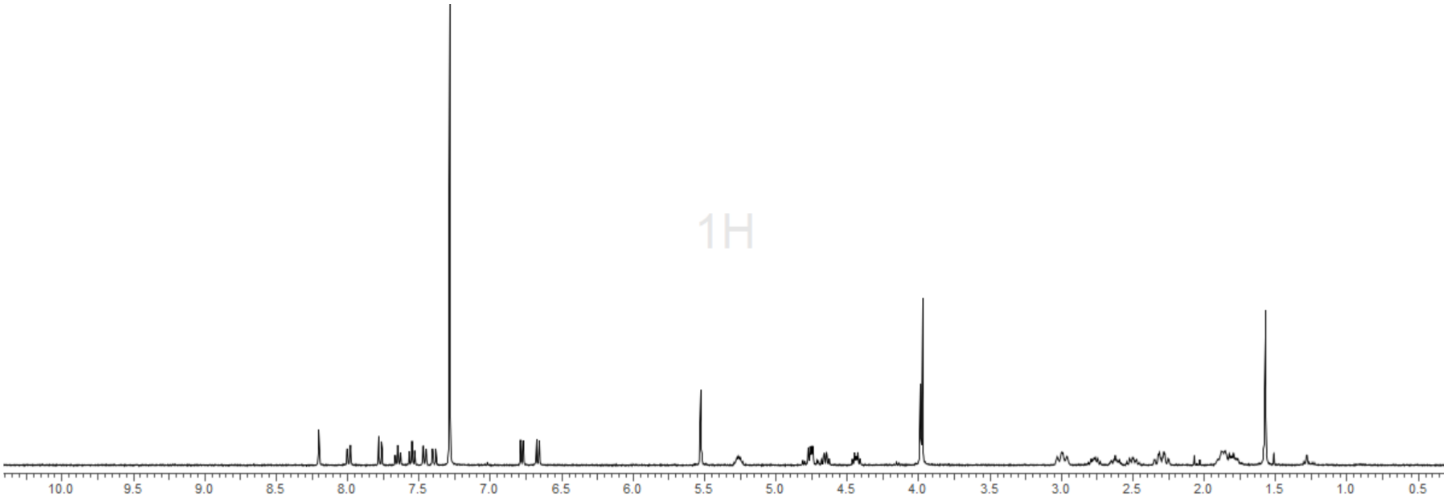

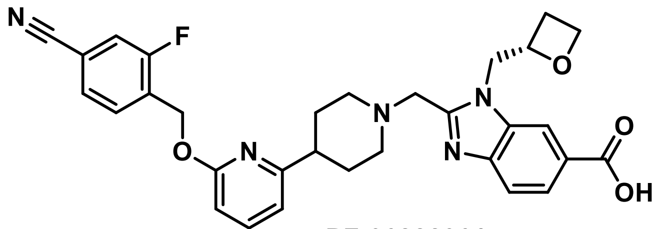
N
F
O
N
N
N
O
O
OH

**PF-06882961**

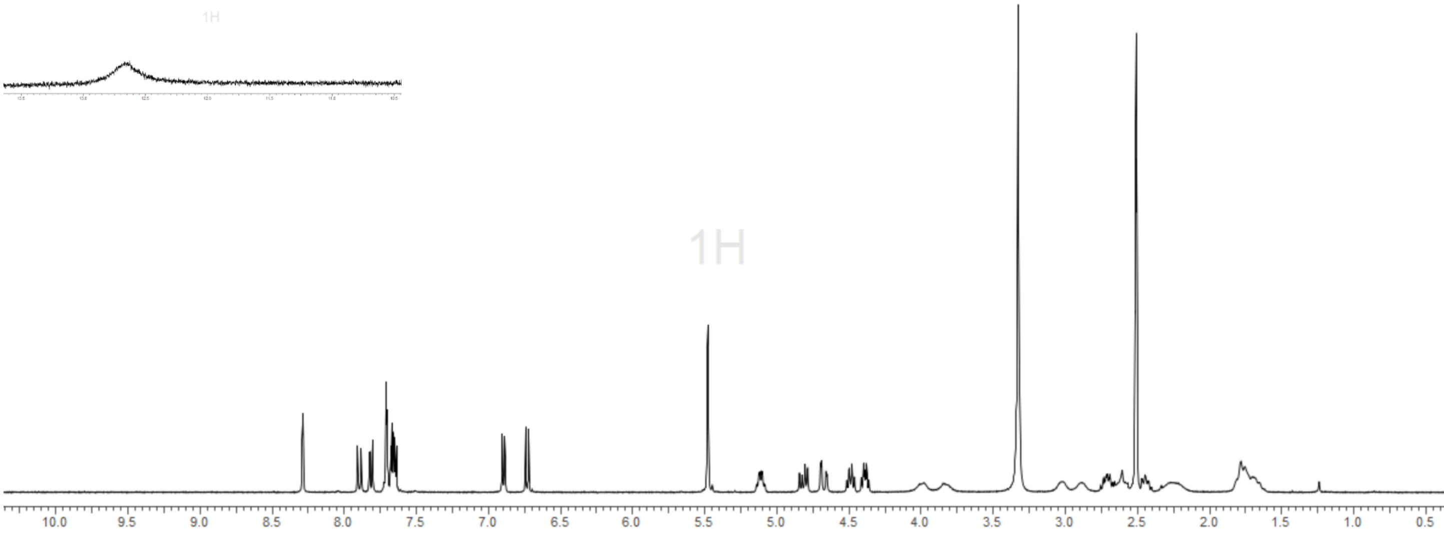
1H
1H
10.0
9.5
9.0
8.5
8.0
7.5
7.0
6.5
6.0
5.5
5.0
4.5
4.0
3.5
3.0
2.5
2.0
1.5
1.0
0.5

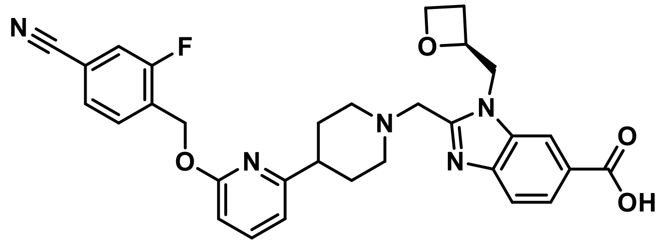


**PF-06882961, tris salt**

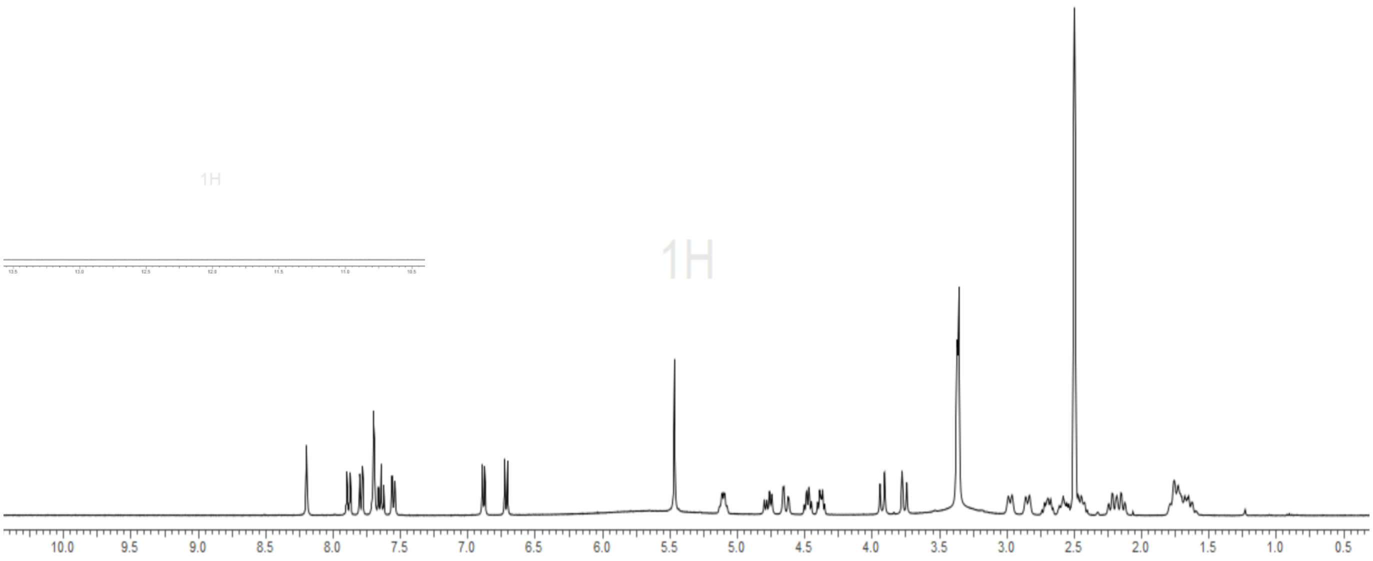


1H/19F decoupled

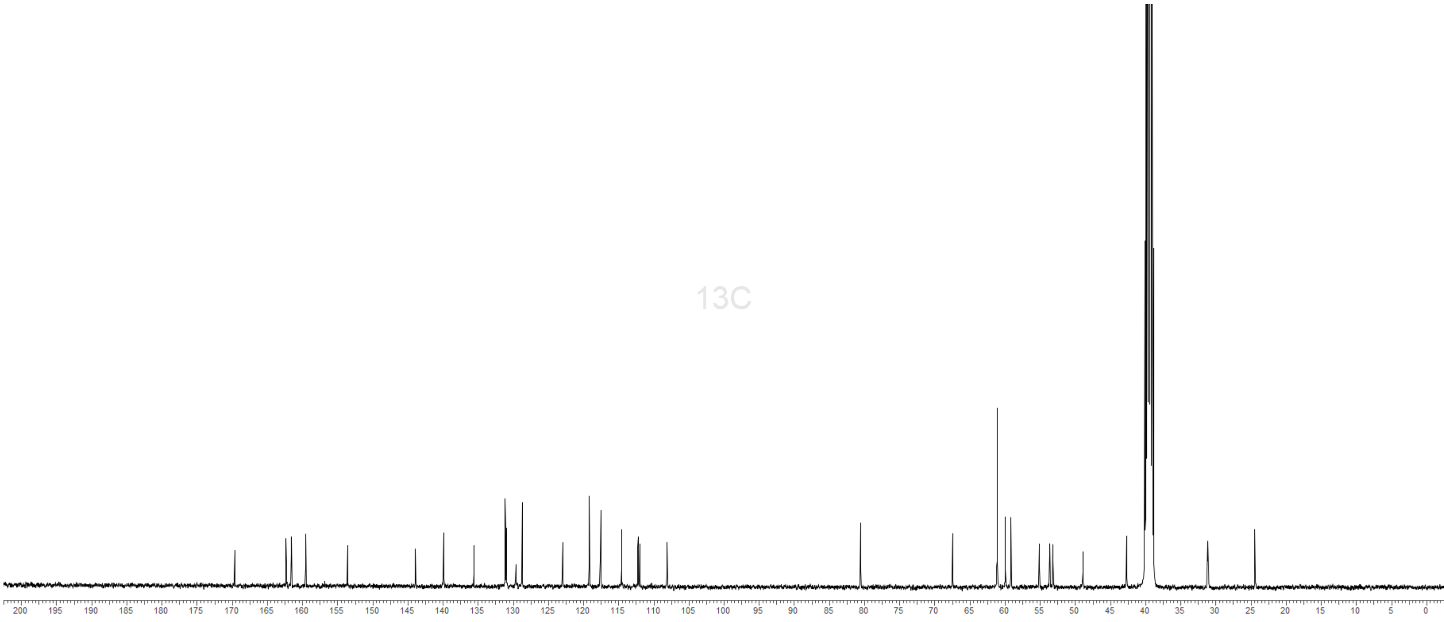

1H decoupled

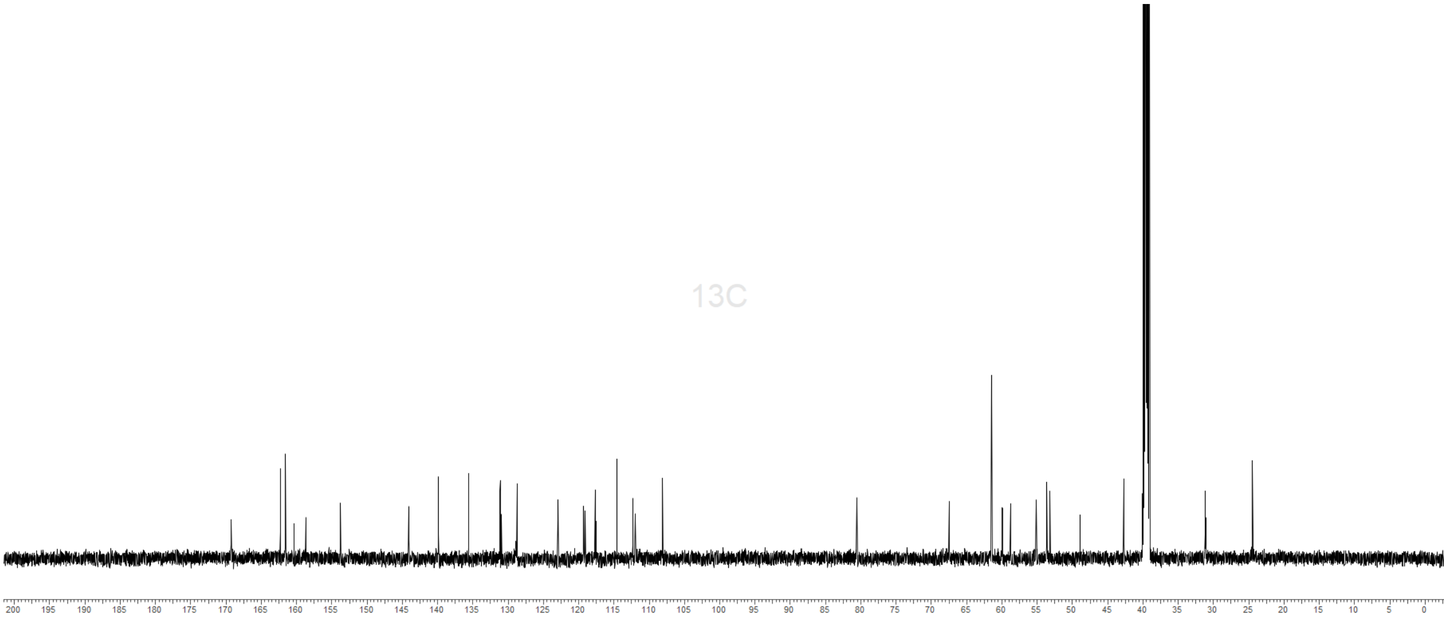


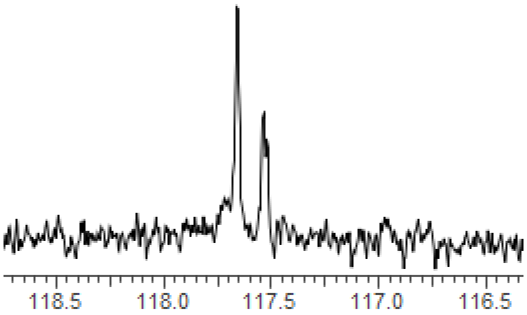


1H decoupled

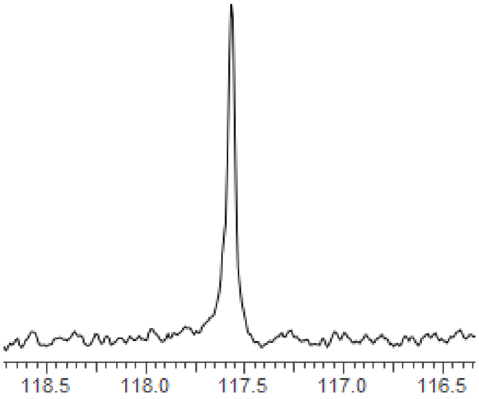


1H/19F decoupled

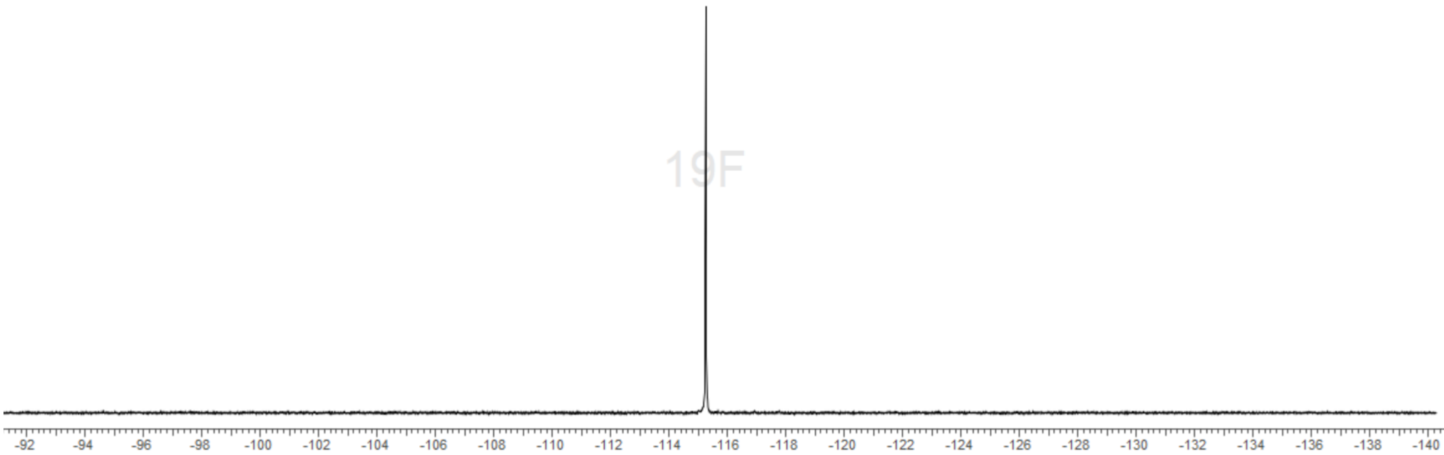

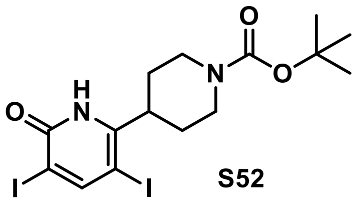

**S52**

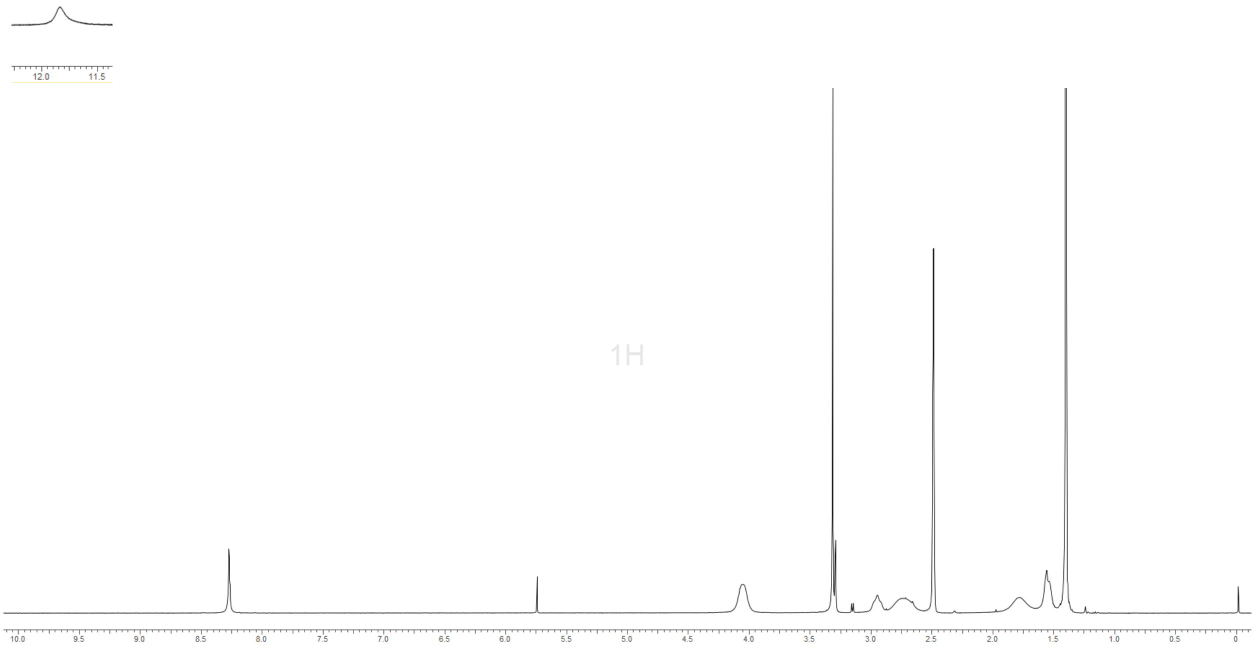

12.0
11.5
1H
10.0
9.5
9.0
8.5
8.0
7.5
7.0
6.5
6.0
5.5
5.0
4.5
4.0
3.5
3.0
2.5
2.0
1.5
1.0
0.5
0

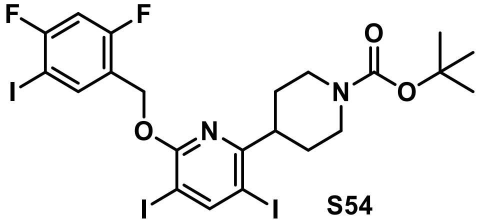


**S54**

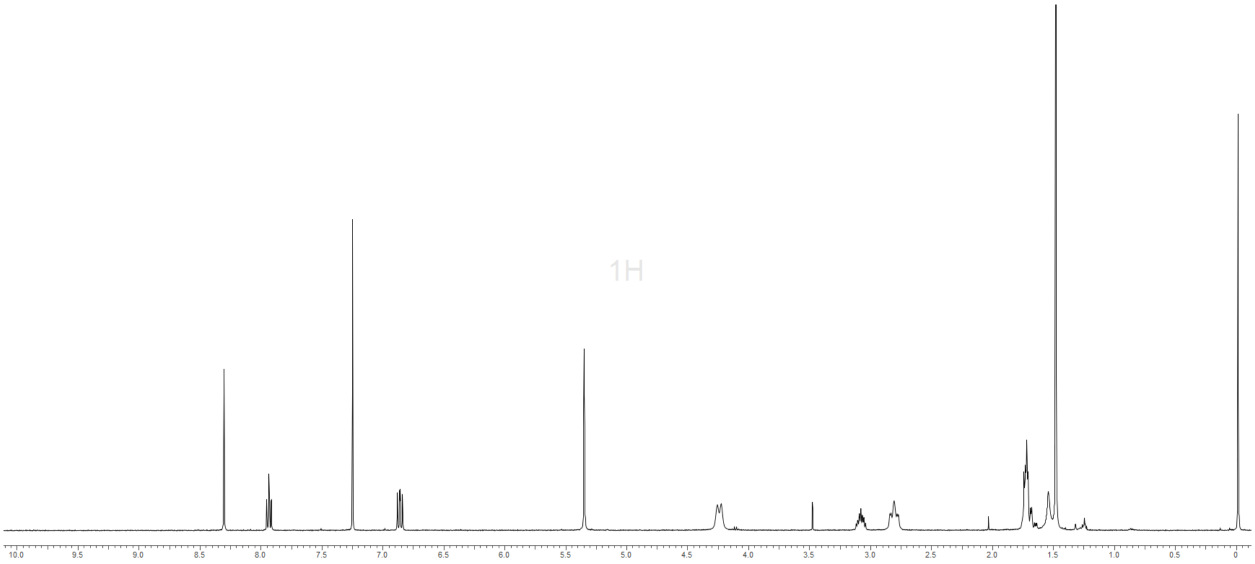


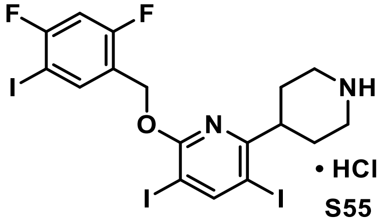


• HCl
**S55**

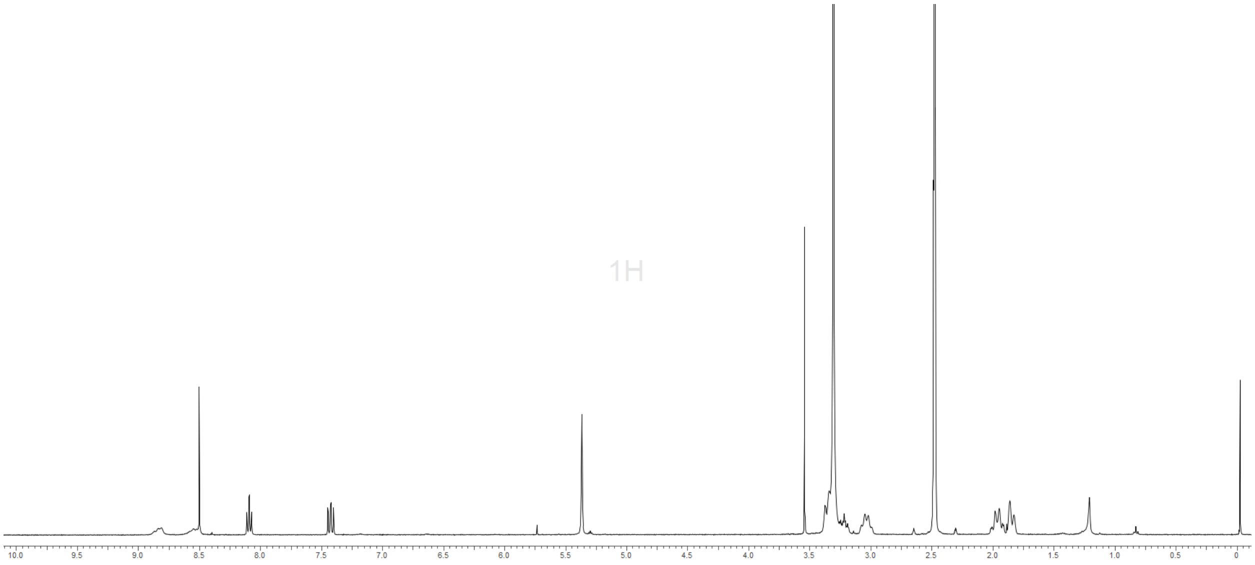

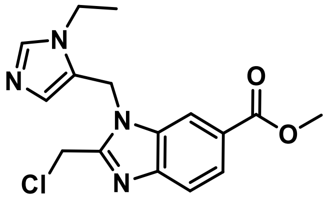

**S56**

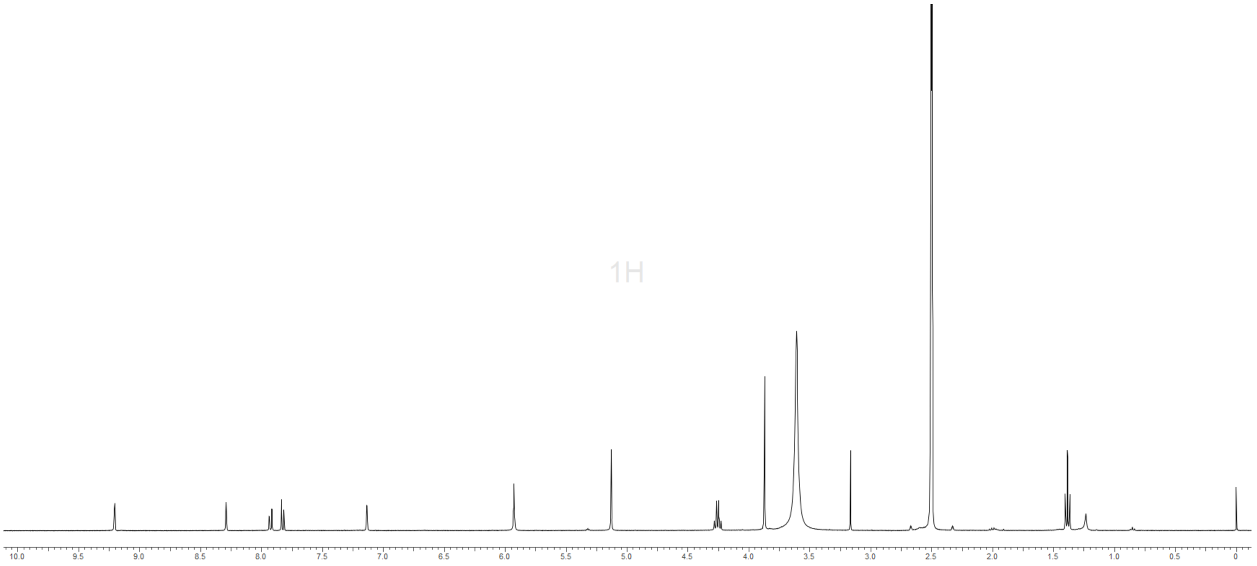

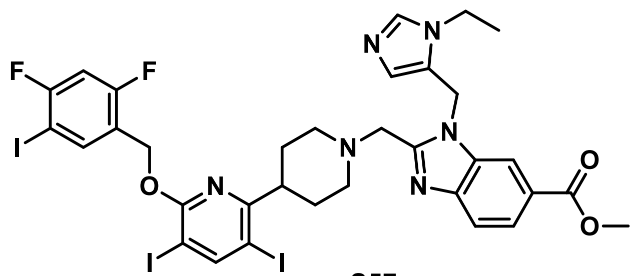

**S57**

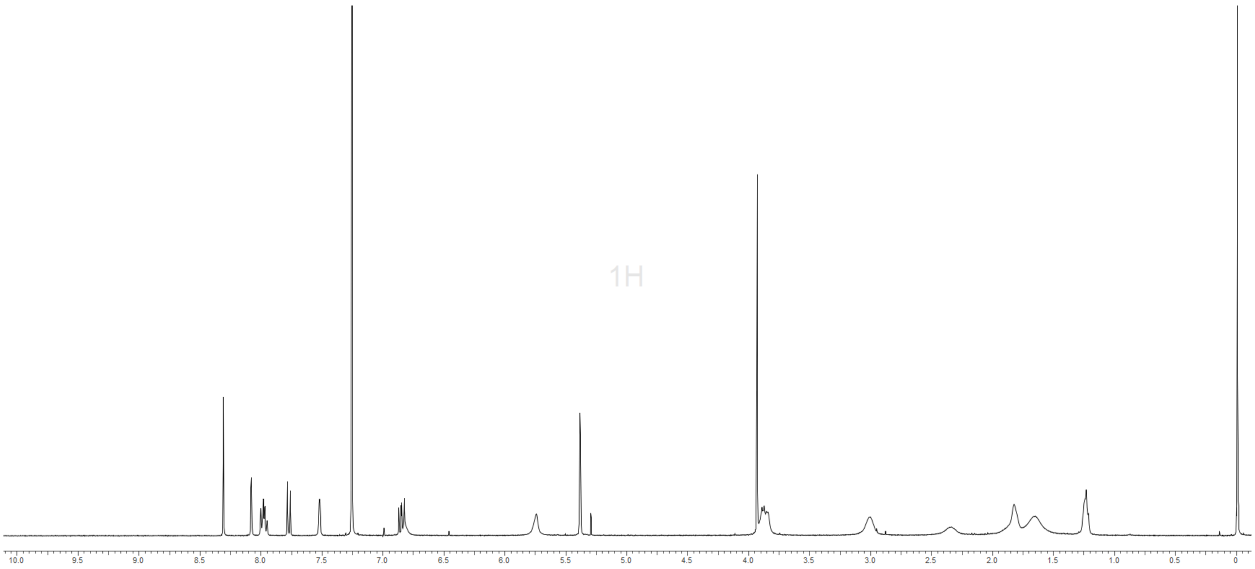


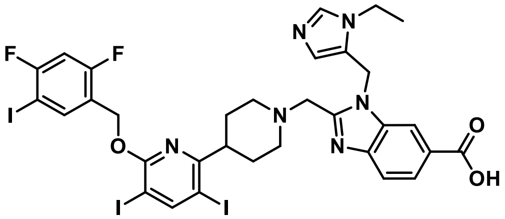

**PF-06899199**

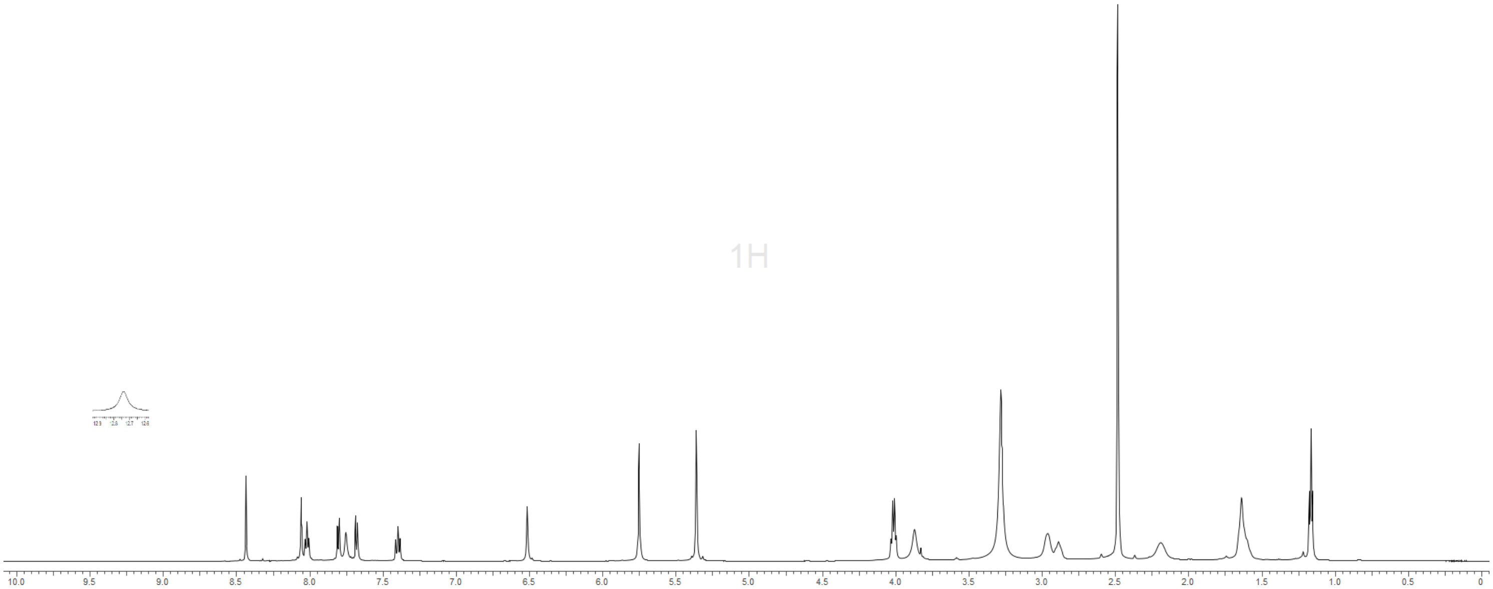

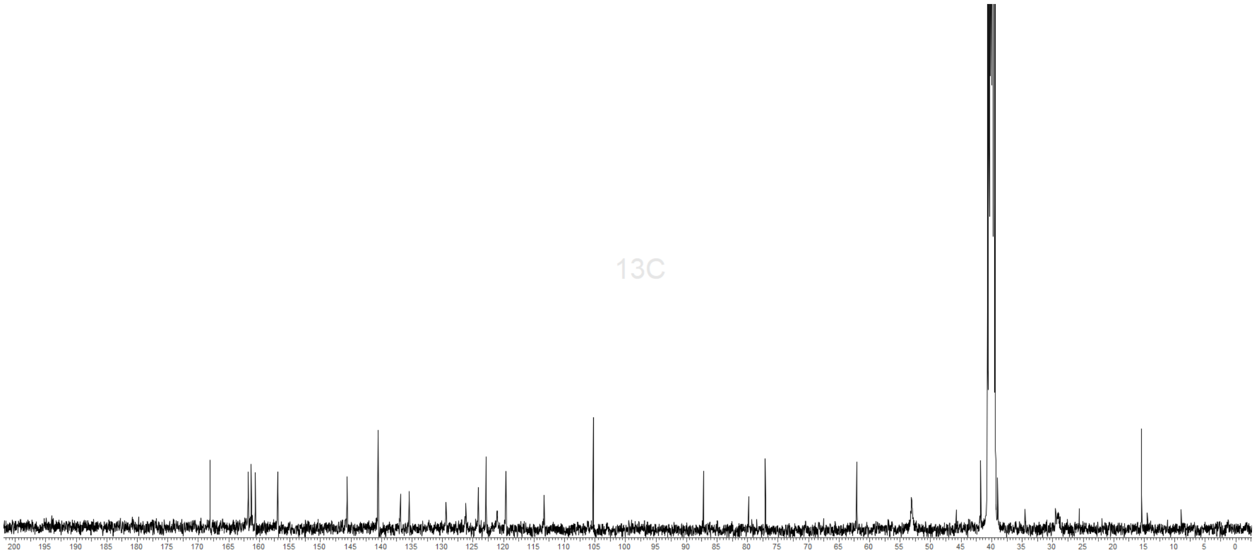
13C

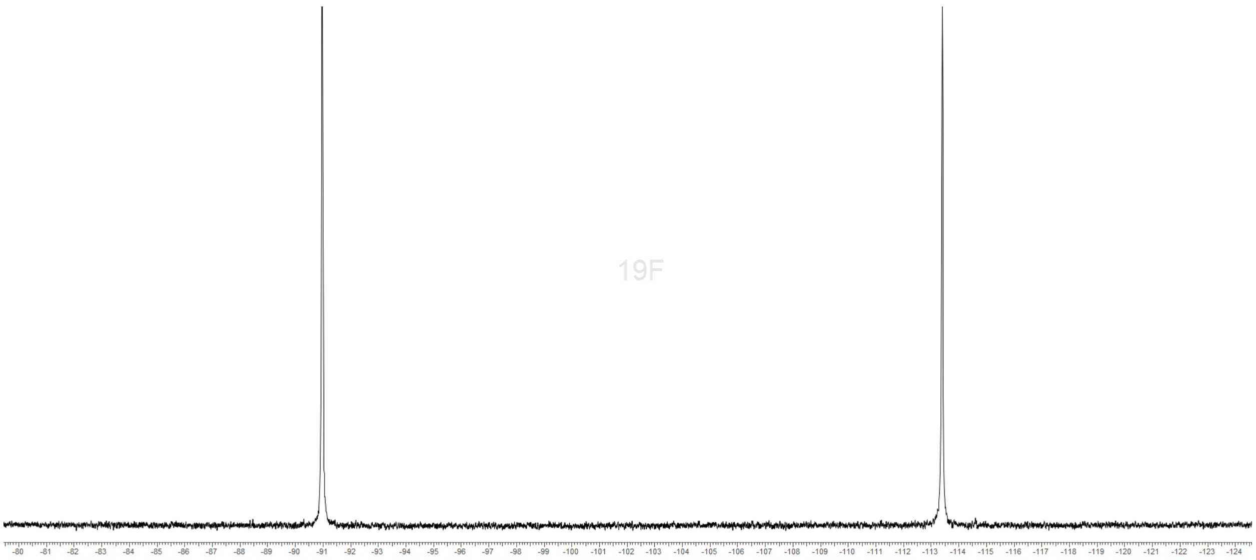
19F

Radioligands

**[$^3$H]-PF-06883365** was prepared by Pharmaron UK (Cardiff, UK) from tri-iodo precursor **S58** and supplied with a specific activity of 86 Ci/mmol and stored at a radiochemical concentration of 1mCi/mL in ethanol/water (95:5). $^{125}$I-TYR-GLP-1(7-36)amide ([$^{125}$I]-GLP-1) was purchased at Perkin Elmer (Cat #NEX308, Billerica, MA)

BETP-sensitized cAMP High Throughput Screening Assay

The high throughput screen (HTS) utilized the CisBio cyclic AMP (cAMP) Homogeneous Time-Resolved Fluorescence (HTRF) assay technology. CHO-K1 cells stably expressing the human GLP-1R (screening cell line) were removed from cryopreservation, thawed, centrifuged (1000 rpm, 5 minutes) and resuspended in complete cell culture media (Dulbecco's Modified Eagle Medium F12, 10% heat inactivated fetal bovine serum, 500 μg/mL Geneticin [G418], 50 units/mL penicillin and 50 μg/mL streptomycin, and 2 mM glutamine). Cells were plated at a density of 1,500 cells/well (20 μL/well) into 384 well assay plates and cultured for 48 hours in a humidified environment at 37 °C with 5% $CO_2$. A library of approximately 2.8 million compounds from the Pfizer Sample Bank was screened in a compressed format, with 14 compounds per well, each at a final assay concentration of 10 μM each. On the day of the assay, compounds were added as 1 μL per well in 384-well polypropylene plates to create intermediate plates. The intermediate plates were then diluted with 100 μL/well of assay buffer (Hanks Balanced Salt Solution [HBSS] with calcium and magnesium, 10 mM HEPES, 0.1% bovine serum albumin, 100 μM IBMX, 3 μM BETP). Positive control wells containing 100 nM GLP-1(7-36)amide (GLP-1; final concentration) and negative control wells containing 1% DMSO (final concentration) were included on all plates. The cell culture media was removed from the cell plates and 20 μL of the diluted compounds was transferred to the cells. Assay plates were then incubated for 30 minutes in a humidified environment at 37 °C with 5% $CO_2$. The production of cAMP was measured using the HI Range cAMP detection kit (Cisbio, Bedford, MA) according to the manufacturer's instructions. The cAMP-d2 working solution was added to the assay plates as 10 μL per well followed by 10 μL anti-cAMP-cryptate working solution. Assay plates were incubated for 1 hour at room temperature and the fluorescence was read using an EnVision plate reader (Perkin Elmer, Chicago, IL) with excitation at 330 nm and emissions of 615 nm and 665 nm. A cAMP standard curve was used to convert the raw data (ratio of the 665 nm/615 nm reads) to cAMP concentrations, as recommended by the manufacturer. The interpolated data (cAMP concentrations) were analyzed using Activity Base (IDBS). The percent effect at each concentration of compound was calculated by Activity Base relative to the amount of cAMP in the positive and negative control wells on each assay plate.

cAMP Production assays for SAR

CHO-K1 cells stably expressing human GLP-1R (either the screening cell line or the candidate selection cell line, as noted in the text) were removed from cryopreservation, thawed, centrifuged (1000 rpm, 5 minutes) and resuspended in complete cell culture media (Dulbecco's

Modified Eagle Medium F12, 10% heat inactivated fetal bovine serum; 500 μg/mL Geneticin, 50 units/mL penicillin, 50 μg/mL streptomycin, and 2 mM glutamine). Cells were plated at a density of 1,600 cells/well (50 μL/well) into Corning 3570 assay plates (Fisher Scientific, Pittsburgh, PA) and cultured for 48 hours in a humidified environment at 37 °C with 5% $CO_2$. On the day of the assay, compounds were serially diluted by half logs in DMSO as an 11-point concentration response and added as 1 μL per well in 384-well polypropylene plates to create intermediate plates. The intermediate plates were then diluted with 100 μL/well of assay buffer (Hanks Balanced Salt Solution with calcium and magnesium; 10 mM HEPES; 0.1% bovine serum albumin; 100 μM IBMX). For BETP-sensitized assays, 3 μM was added to the assay buffer. Positive control wells containing 1 μM GLP-1 (final concentration) and negative control wells containing 1% DMSO (final concentration) were included on all plates. The cell culture media was removed from the cell plates and 20 μL of the diluted compounds was transferred to the cells. Assay plates were then incubated for 30 minutes in a humidified environment at 37 °C with 5% $CO_2$. The production of cAMP was measured using the HI Range cAMP detection kit (Cisbio, Bedford MA) according to the manufacturer's instructions. The cAMP-d2 working solution was added to the assay plates as 10 μL per well followed by 10 μL anti-cAMP-cryptate working solution. Assay plates were incubated for 1 hour at room temperature on the lab bench and the fluorescence was read using an EnVision plate reader (Perkin Elmer, Chicago, IL) with excitation at 330 nm and emissions of 615 nm and 665 nm. A cAMP standard curve was used to convert the raw data (ratio of the 665 nm/615 nm reads) to cAMP concentrations, as recommended by the manufacturer. The interpolated data (cAMP concentrations) were analyzed using Activity Base (IDBS). The percent effect at each concentration of compound was calculated by Activity Base relative to the amount of cAMP in the positive and negative control wells on each assay plate. Compound $EC_{50}$ values were determined using a logistic 4 parameter fit model. The efficacy for each compound was defined by the maximum asymptote of the fitted curve and expressed as a percent of the maximum response produced by the positive control on each plate.

<u>Selectivity assessment at GIPR, GLP-2R and GCGR.</u>

These studies were performed using the Hit Hunter® assay technology (DiscoveRX Corp, Freemont, CA). Hit Hunter cell lines stably expressing human GLP-2R or GCG-R were counted and seeded into white 384-well microplates and incubated at 37 °C for 24 hours. Culture media was then removed and replaced with assay buffer (2:1 HBSS/10mM HEPES : cAMP XS + Ab reagent). Varying concentrations of PF-06882961 (in DMSO) were diluted in assay buffer. To assess agonist activity, cells were stimulated with increasing concentrations of PF-06882961 for 30 minutes at 37 °C. To assess antagonist activity, cells were pretreated with increasing concentrations of PF-06882961 for 30 minutes at 37 °C, followed by stimulation with an $EC_{80}$ concentration of GLP-2 (1.3 nM) or glucagon (0.75 nM) for an additional 30 minutes at 37 °C. Assay signal was then revealed through incubation with the cAMP XS+ ED/ CL lysis cocktail for one hour, followed by incubation with the cAMP XS+ EA reagent for 3 hours at room temperature. Microplates were read following signal generation with a PerkinElmer Envision multi-label plate reader for chemiluminescent signal detection. For agonist mode assays, the percent effect was determined relative to a saturating concentration of the cognate full agonist included on each plate. For all antagonist mode assays, cAMP values were normalized relative to the effect triggered by the $EC_{80}$ of the cognate endogenous agonist in the absence of PF-06882961. Four parameters sigmoidal curve fitting of ligand concentration-response curves was performed using GraphPad (GraphPad Prism software version 5.0, San Diego, CA) and used for calculating the $EC_{50}/IC_{50}$ and $E_{max}$ values.

Plasma membrane isolation

Cells stably expressing the human GLP-1R were grown in T225 cell culture flasks to 90% confluency as described above. The following procedure was performed at 4 °C. Cells were washed once with ice-cold PBS and 15 mL of PBS supplemented with EDTA 5 mM was added to each flask to dissociate cells. Following a 10 minutes incubation period, the cell suspension was transferred into 50 mL Falcon tubes and centrifuged at 1000 g for 10 minutes. The supernatant was discarded, and cell pellets were resuspended in 5 mL of ice-cold homogenization buffer (HB: 20 mM Tris-HCL, pH 7.4 containing 2 mM EDTA, 2 mM EGTA and 1X protease inhibitors [Pierce, Catalog no. A32963]). The cell suspension was then homogenized using a tight-fitting glass Dounce Homogenizer (20-25 strokes). The cell homogenate was transferred into 50 mL Falcon tubes and spun down at 1000 g for 10 minutes. Supernatants were saved, and the remaining cell pellets were homogenized and centrifuged again using the procedure described above. Combined supernatants from both homogenization steps were then centrifuged at 25,000 rpm (Beckman JA-17 rotor, Indianapolis, IN) for 20 minutes. The pellet was washed once using 10 mL of HB and centrifuged again using the same conditions. The membrane pellet was resuspended in an appropriate volume of HB yielding a total protein concentration of approximately 5 mg/mL. Protein concentration was measured using BCA kit (Pierce, Catalog no. 23225).

Radioligand binding assays

Saturation binding analyses using [$^3$H]-PF-06883365 were carried out using a filter-binding format in total volumes of 250 µL per well in 96-well polypropylene microtiter plates. Two hundred µL of plasma membranes (12 µg protein/well) diluted in assay buffer (50 mM HEPES pH 7.4 containing 0.1% BSA, 2 mM $CaCl_2$, 10 mM $MgCl_2$) was added to each assay well. The wells defined as “Total Binding” and “Non-specific Binding” on each plate received either 25 µL of assay buffer or 25 µL of unlabeled PF-06883365 diluted to a final concentration of 10 µM in assay buffer, respectively. Following 10 minutes incubation at room temperature, 11 final concentration of [$^3$H]-PF-06883365 (400, 200, 150, 100, 50, 25, 12.5, 6.25, 3.13, 1.56 nM) diluted in assay buffer were added in quadruplicate to wells designated as specific and non-specific binding. Incubation was then carried out on a shaker over a 6-hour period at room temperature. At the end of the incubation period, binding reactions were harvested (Harvester 96

Mach III, Tomtec, Hamden, CT) onto 96-well glass-fiber filter plates pretreated with 0.3% polyethyleneimine (PEI). Each filter plate was rapidly washed 4 times with washing buffer (ice cold 50 mM Tris, pH 7.4, containing 250 mM NaCl, filtered). The plates were air dried overnight, sealed and counted in a Microbeta counter after 35 μL of Ultima Gold scintillation cocktail (Perkin Elmer, Chicago, IL), was added into each well. The total number of specific binding sites ($B_{max}$) was determined by globally analyzing the total and nonspecific binding at one time using GraphPad (GraphPad Prism software version 5.0, San Diego, CA). Calculated $B_{max}$ values (in CPM) were converted to fmol/mg of protein and normalized as % of maximal specific binding.

For unlabeled PF-06882961, competition binding assays based on [$^3$H]-PF-06883365 were carried out in a filter-binding format in total volumes of 250 μL per well using 96-well polypropylene microtiter plates. Eight concentrations of PF-06882961 (0.001, 0.01, 0.1, 1, 10, 100, 1000 and 10000 nM), diluted in assay buffer (50 mM HEPES pH 7.4 containing 0.1% BSA, 10 mM $MgCl_2$, 2 mM $CaCl_2$), were added at 25 μL to wells on the plates in quadruplicate. Wells defined as "Total Binding" or "Non-specific Binding" received either 25 μL of assay buffer or 25 μL of unlabeled PF-06883365 diluted to a final concentration of 10 μM in assay buffer, respectively. A final concentration of 50 nM [$^3$H]-PF-06883365 (in 25 μL) was then added to each assay well. The binding reactions were initiated with the addition of 200 μL of plasma membranes (12 μg protein/well) derived from a sodium butyrate-treated (6 mM NaBu overnight) stable CHO cell line stably expressing the human GLP-1R (clone G18) into each assay well. Incubation was then carried out on a shaker over a 4-hour period at room temperature. At the end of the incubation period, binding reactions were harvested (Harvester 96 Mach III, Tomtec, Hamden, CT) onto 96-well glass-fiber filter plates pretreated with 0.3% polyethyleneimine (PEI). Each filter plate was rapidly washed 4 times with washing buffer (ice cold 25 mM Tris, pH 7.4, containing 500 mM NaCl, filtered) and then air dried overnight at room temperature. The filter plates were sealed and counted in a Microbeta counter after 35 μL of Ultima Gold scintillation cocktail (Perkin Elmer, Chicago, IL), was added into each well. Specific radioligand binding was calculated as total binding (total counts per minute, CPM) for each well minus the averaged non-specific binding measured in the presence of the excess of PF-06883365. The ability of small molecules to bind the human GLP-1R was calculated as the inhibitor concentration required for 50% inhibition ($IC_{50}$ values) of specific binding. Four parameters sigmoidal curve fitting of binding competition curves was performed using GraphPad (GraphPad Prism software version 5.0, San Diego, CA) and used for calculating the $IC_{50}$ values. Using the Cheng-Prusoff equation, $IC_{50}$ were converted to Ki values using a Kd value (38 nM) determined from saturation binding experiments (*40*).

For unlabeled PF-06883365, Liraglutide and Exenatide, competition binding assays based on [$^3$H]-PF-06883365 were carried out in a filter-binding format in total volumes of 100 μL per well using 96-well polypropylene microtiter plates. Compounds were serially diluted as half log,

10-point dose responses in 100% DMSO. 1 μL of the serially diluted compound was spotted into the appropriate wells of the 96 well polypropylene microtiter assay plate. The compound spots were then diluted with 50 μL of assay buffer (50 mM HEPES pH 7.4 containing 0.1% BSA, 10 mM $MgCl_2$, 5 mM $CaCl_2$) containing [$^3$H]-PF-06883365 for a final assay concentration of 8 nM [$^3$H]-PF-06883365. Wells defined as "Total Binding" or "Non-specific Binding" contained final well concentration of 1% DMSO or 20 μM unlabeled PF-06883365 respectively. The binding reactions were initiated with the addition of 50 μL of plasma membranes (10 μg protein/well). The plasma membrane was derived from a sodium butyrate-treated (6 mM NaBu overnight) CHO cell line stably expressing the human GLP-1R (clone G18). Incubation was then carried out on a shaker over a 6 hour time period at room temperature. At the end of the incubation period, binding reactions were harvested using a Packard 96 well Harvester onto 96-well GF/C filter plates (Perkin Elmer) pretreated with 0.3% polyethyleneimine (PEI). Each filter plate was rapidly washed 4 times with wash buffer (ice cold 50 mM HEPES, pH 7.4, containing 250 mM NaCl, 0.1% BSA) and then air dried overnight at room temperature. The next day, 50 μL of Ultima Gold scintillation cocktail (Perkin Elmer, Chicago, IL) was added to each well, the plates were top sealed, and then counted in a Microbeta counter (Perkin Elmer). Specific radioligand binding was calculated as total binding (total counts per minute, CPM) for each well minus the averaged non-specific binding measured in the presence of the excess of PF-06883365. The ability of compounds to bind the human GLP-1R was calculated as the inhibitory concentration required for 50% inhibition ($IC_{50}$ values) of specific binding. Data were normalized using plate total binding and non-specific binding values, and four parameters sigmoidal curve fitting of binding competition curves was performed using ABASE data analysis software (IDBS)) and used for calculating the $IC_{50}$ values. Using the Cheng-Prusoff equation, $IC_{50}$ values were converted to Ki values based on $K_d$ values (80 nM) obtained from saturation binding analysis (*40*).

Saturation binding analyses based on [$^{125}$I]-GLP-1 was also carried out using membrane derived from CHO cell lines stably expressing the hGLP1-R (screening cell line and candidate selection cell line). Total volumes of of 100 µL per well in 96-well polypropylene microtiter plates were used in a filter binding assay format. Fifty µL of plasma membranes (1.5 µg and 3.5 µg protein/well for the screening assay and candidate selection cell lines, respectively) diluted in assay buffer (50 mM HEPES pH 7.4 containing 0.2% BSA, 10 mM $MgCl_2$, 5 mM $CaCl_2$) was added to each assay well. The wells defined as "Total Binding" and "Non-specific Binding" on each plate received either 25 μL of assay buffer or 25 μL of unlabeled Exendin-4 diluted to a final concentration of 300 nM in assay buffer, respectively. Following 10 minutes incubation at room temperature, 9 final concentrations of [$^{125}$I]-GLP-1 (0.015, 0.031, 0.061, 0.125, 0.25, 0.5, 1 and 2 nM) diluted in assay buffer were added in triplicate to wells designated as specific and non-specific binding. Incubation was then carried out on a shaker over a 3-hour period at room temperature. At the end of the incubation period, binding reactions were harvested as described above and each filter plate was rapidly washed 10 times with washing buffer (ice cold 50 mM

Tris, pH 7.4, containing 250 mM NaCl, filtered). The plates were air dried overnight and counted in a Microbeta counter as described above. The total number of specific binding sites ($B_{max}$) was determined by globally analyzing the total and nonspecific binding at one time using GraphPad (GraphPad Prism software version 5.0, San Diego, CA). Calculated $B_{max}$ values (in CPM) were converted to fmol/mg of protein.

Competition binding assays based on [$^{125}$I]-GLP-1 (7-36) were carried out in a 384 well SPA binding assay format in total volumes of 100 μL per well. Compounds or peptides were serially diluted as half log, 11-point dose responses in 100% DMSO. 1 μL of the serially diluted compound was spotted into a 384 well intermediate assay plate, and 100 μL of assay buffer (50 mM HEPES pH 7.4 containing 0.2% Fatty Acid Free BSA, 0.01% Pluronic F127, and 5% glycerol) was added to the compound intermediate plate. Wells defined as "Total Binding" or "Non-specific Binding" contained a final well assay concentration of 1% DMSO 1 μM Exendin-4 respectively. After thorough mixing, 25 μL of compound in assay buffer was transferred from the compound intermediate assay plate to a Matrix 4322 white clear bottom assay plate. 25 μL of WGA PVT SPA beads were added to the Matrix 4322 assay plate for a final assay amount of 100 ug/well, followed by the addition of 25 μL of [$^{125}$I]-GLP-1 for a final assay concentration of 250 pM. Binding reactions were initiated with the addition of 25 μL of plasma membranes (2 μg/well) derived from a CHO cell line stably expressing the human GLP-1R (clone G18) into each assay. Incubation was then carried out on a shaker over a 30 minutes time period at room temperature, followed by a 10 hour incubation in the dark. The assay plate was then read on a Trilux Microbeta using a normalized protocol with a 1 minute/well read. The ability of small molecules and peptides to bind the human GLP-1R was calculated as the inhibitory concentration required for 50% inhibition ($IC_{50}$ values) of specific binding. Data were normalized using plate total binding and non-specific binding values, and four parameters sigmoidal curve fitting of binding competition curves was performed using ABASE data analysis software (IDBS) and used for calculating the $IC_{50}$ values. Using the Cheng-Prusoff equation, $IC_{50}$ values were converted to Ki values based on a $K_d$ value (0.3 nM) obtained from saturation binding analysis (*40*).

<u>β-Arrestin Recruitment Assay (Assay Media)</u>

β-Arrestin recruitment at the human GLP-1R was evaluated using DiscoverX's Enzyme Fragment Complementation (EFC) technology as described by the manufacturer (DiscoverX, Freemont, CA). Cryopreserved Path Hunter GLP-1 β-arrestin 1 or β-arrestin 2 expressing cells were thawed, washed, and resuspended in complete growth media (HAMs/F-12 media containing 10% FBS heat inactivated, 1X Pen/Strep, 1X glutamine, 300 μg/mL hygromycin and 800 μg/mL geneticin). Following overnight incubation at 37 °C in a humidified environment (5% $CO_2$), cells were dissociated and re-plated in white 384-well plates at 5000 cells/well in 18 μL assay media overnight (HAMs/F-12 media containing 10% FBS heat inactivated, 1X Pen/Strep, 1X glutamine).

Compounds were serially diluted as 11-point, half log dose responses in 100% DMSO, and 1 μL of the serially diluted compounds was transferred to a compound source plate. Prior to addition to cells, the 1 μL compound spots were diluted in assay media +/- BETP. 2 μL of a 20X compound dose response series was transferred from the compound source plate to the appropriate wells in the assay plate. The final concentration of BETP-containing wells was 10 μM BETP. Plates were then incubated with compound for 90 minutes at 37 °C in a humidified environment (5% $CO_2$). Following the 90 minute incubation, 10 μL of Path Hunter Detection Reagent was added per well and assay plates were incubated for 60 minutes at room temperature. Chemiluminescence activity was then measured using an Envision multi-label plate reader.

The raw data were analyzed using Activity Base (IDBS). The percent effect at each compound concentration was calculated in Activity Base, relative to the positive and negative control wells on each assay plate. The negative control wells contained a final assay concentration of 1% DMSO, while the positive control wells contained a final assay concentration of 1 μM GLP-1. Compound $EC_{50}$ values were determined using a logistic 4 parameter fit model. The efficacy for each compound was defined by the maximum asymptote of the fitted curve and expressed as a percent of the maximum response produced by the plated positive control.

<u>β-arrestin Recruitment Assay (assay buffer with 0.1% BSA)</u>

β-arrestin recruitment at the human GLP-1R was evaluated using DiscoverX's Enzyme Fragment Complementation (EFC) technology as described by the manufacturer (Eurofins DiscoverX, Freemont, CA). Cryopreserved Path Hunter GLP-1 β-arrestin 1 or β-arrestin 2 cells (DiscoverX, Freemont, CA) were thawed, washed, resuspended in complete growth media (HAMs/F-12 media containing 10% FBS heat inactivated, 1X Pen/Strep, 1X glutamine, 300 μg/mL hygromycin and 800 μg/mL geneticin). Following overnight incubation at 37 °C in a humidified environment (95% $O_2$, 5% $CO_2$) cells were dissociated and re-plated in white 384-well plates at 5000 cells/well in 20 μL assay media overnight (HAMs/F-12 media containing 10% FBS heat inactivated, 1X Pen/Strep, 1X glutamine).

Compounds were serially diluted as 11-point, half log dose responses in 100% DMSO, and 1 μL of the serially diluted compounds was transferred to a compound source plate. The 1 μL compound spots were then diluted in assay buffer (HBSS [+ calcium/magnesium] containing 20 mM HEPES, 0.1% BSA). 20 μL of the compound dose response series was transferred from the compound source plate to the appropriate wells in the assay plate. Plates were then incubated with compound for 90 minutes at 37 °C in a humidified environment (5% $CO_2$). Following the 90 minute incubation, 10 μL of Path Hunter Detection Reagent was added per well and assay plates were incubated for 60 minutes at room temperature. Chemiluminescence activity was then measured using an Envision multi-label plate reader.

The raw data were analyzed using Activity Base (IDBS). The percent effect at each compound concentration was calculated in Activity Base, relative to the positive and negative control wells on each assay plate. The negative control wells contained a final assay concentration of 1% DMSO, while the positive control wells contained a final assay concentration of 1 μM GLP-1. Compound $EC_{50}$ values were determined using a logistic 4 parameter fit model. The efficacy for each compound was defined by the maximum asymptote of the fitted curve and expressed as a percent of the maximum response produced by the plated positive control.

Bias calculations

Bias factor calculations were performed as described by Kenakin (2017) as follows: For each ligand and respective response, individual experimental curves were used to calculate $\log(E_{max}/EC_{50})$. The difference in $\log(E_{max}/EC_{50})$ between cAMP and either βArr1, βArr2 or internalization, $\Delta\log(E_{max}/EC_{50})$, was then calculated for each ligand. Finally, the differences between the $\Delta\log(max/EC_{50})$ values for the reference ligand (Liraglutide) and test ligand were calculated to give a $\Delta\Delta\log(max/EC_{50})$ values, the antilog of which is the bias factor. Bias factors were calculated separately for assays performed using i) HEK 293 expressing FAP-GLP1R (cAMP, internalization) and ii) CHO cells assays (CS cAMP, βArr1, βArr2), and are displayed using a web of bias.

Confocal Microscopy

HEK293 cells stably expressing hGLP-1R fused to green fluorescent protein (GFP) (400,000 cells per well) were cultured onto 6 well plates for 24 hours and stimulated with PF-06882961 for 30 minutes. An agonist concentration of 1 μM shown to induce maximal internalization was chosen for these studies. In selected wells, cells were washed three times with PBS containing 0.1 % BSA and incubated at 37 °C for an additional 2 hours to assess reversibility of the endocytosis process. Cells were then fixed with 4% paraformaldehyde (Alfa Aesar, Catalog no. 43368-9M) for 15 minutes at room temperature followed by three washes with PBS containing 0.1 % BSA. Nuclei were then stained by with Hoescht 33342 (Invitrogen, catalog no. H3570) and imaged with a confocal microscope (Zeiss LSM 780, White Plains, NY) using a W-Plan Apochromat 63x/1.0 NA lens. For quantification of receptor internalization, raw images were analyzed in CellProfiler Analyst, which analyses a range of features on each cell (e.g. intensity, morphology, granularity, and puncta counts per cell) (*41*, *42*).

Functional Assessment of GLP-1R Mutants

GLP-1R expression constructs for mutant analysis were made by amplifying the human (NP_002053.3), murine (NP_067307.2), and cynomolgus monkey (XM_005553112) GLP-1R codon optimized template DNA by PCR with primers designed to incorporate restriction sites followed by restriction endonuclease digestion and ligation into pcDNA3.1 (Thermo Fisher, Pittsburgh, PA). Site-specific mutagenesis was performed using QuikChange mutagenesis kit (Agilent, Santa Clara CA).

For transient transfection, Expi293 cells (Thermo Fisher, Pittsburgh, PA) were seeded in 30 mL of Expi293 Expression Media (Thermo Fisher, Pittsburgh, PA) at $2x10^6$ cells/mL and cultured in a 125 mL shake flask at 125 rpm at 37 °C, 5% $CO_2$ for 16-18 hours. Cells were then suspended at $7.5x10^7$ cells in 25.5 mL of Expi293 Expression Media in a 125 mL shake flask. Plasmid DNA for each respective GLP-1R mutant (30 μg) was added to 1.5 mL OptiMEM1 Reduced Serum Media (Thermo Fisher, Pittsburgh, PA) and mixed gently. Expifectamine (81 μL, Thermo Fisher, Pittsburgh, PA) was added to 1.5 mL of OptiMEM1 Reduced Serum Media. Both the plasmid mix and transfection reagent mix were incubated independently for 5 minutes at room temperature. Expifectamine/media solution was added to the DNA/media solution, mixed gently, and incubated for 20 minutes at room temperature. The 3 mL of DNA/transfection reagent mix was added to the cells, and the cell flask was shaken 125 rpm at 37 °C and 5% $CO_2$. After 16-18 hours, 150 μL of Enhancer 1 (Thermo Fisher, Pittsburgh, PA) and 1.5 mL of Enhancer 2 (Thermo Fisher, Pittsburgh, PA) were added to the cells for a final culture volume of 30 mL. After another 26-28 hours, transfected cells were harvested and frozen at a concentration of 8 million viable cells per vial in 1 mL freezing media (60% Expi293 Expression Media, 30% Heat Inactivated FBS and 10% DMSO). Cell vials were frozen slowly in a cryo-container placed in a -80 °C freezer overnight, and then transferred to a liquid nitrogen tank for storage in the vapor phase until the day of the assay.

On the day of the assay, one vial of transfected cells was rapidly thawed in a 37 °C water bath and the cells were rinsed with Dulbecco's phosphate-buffered saline (DPBS, Thermo Fisher, Pittsburgh, PA). Each cell sample was centrifuged at 300 x g for 5 minutes, the supernatant was aspirated, and the cells were resuspended in assay buffer consisting of HBSS (+ calcium and magnesium), 0.1% BSA, 20 mM HEPES and 100 μM IBMX. The stock cell concentrations were prepared at either 500,000 viable cells/mL or 250,000 viable cells/mL depending on the transiently transfected construct.

Compounds were prepared as 10 mM stocks in 100% DMSO and stored in a nitrogen environment. Peptides were prepared as 1 mM stocks in 100% DMSO and also stored in a nitrogen environment. Compounds and peptides were serially diluted 1:2 in 100% DMSO as a 22-point concentration response. The serially diluted compound or peptide was added as 1 μL per well in 384-well polypropylene plates to create intermediate plates. Intermediate plates were then diluted with 49 μL/well of assay buffer. A GLP-1 dose response curve was included on each plate, and negative control wells containing 1% DMSO (final concentration) were included on all plates. After thorough mixing, 10 μL of 2X serially diluted compound or peptide in assay buffer was added to a Corning 384 well 3570 assay plate, followed by the addition of 10 μL of cell solution. The final concentration of compound ranged from 0.476 pM –1 μM, while the final concentration of peptide in the assay was either 0.476 pM – 1 μM or 0.048 pM –100 nM depending on the GLP-1R construct being screened. The viable cell concentration was adjusted to deliver either 2,500 or 5,000 cells/well in 10 μL of assay buffer, depending on the GLP-1R construct being screened.

Working solutions of cAMP-d2 and anti-cAMP-cryptate from the HI Range cAMP detection kit (Cisbio, Bedford MA) were prepared according to the manufacturer’s instructions. The cAMP-d2 working solution was added to the assay plates as 10 μL per well to stop the reaction and then 10 μL anti-cAMP-cryptate working solution was added to the plates. Assay plates were incubated for 1 hour at room temperature then the fluorescence was read using an EnVision plate reader (Perkin Elmer, Chicago, IL) with excitation at 330 nm and emissions of 615 nm and 665 nm. A cAMP standard curve was generated using the cAMP stock solution provided in the Cisbio kit which was then used to convert the raw data (ratio of the 665 nm/615 nm reads) to cAMP concentrations. The percent effect at each concentration of compound or peptide was calculated relative to the positive and negative controls in the assay. The cAMP level determined for the GLP-1 maximum asymptote derived from the dose response on each plate was set as the positive control, while the negative control wells on each assay plate contained only DMSO. Compound $EC_{50}$ values were determined using a logistic 4 parameter fit model. The efficacy for each compound was defined by the maximum asymptote of the fitted curve and expressed as a percent of the maximum response produced by the positive controls on each plate.

Cryo-EM Structure Determination

The GLP-1R protein was purified in the presence of the ligand PF-06883365, a close analog of PF-06882961, following the protocols described previously for GLP-1R structures (*35, 43*). The CryoEM protein sample was vitrified on a graphene oxide (GO)-covered quantifoil EM grid. Aliquots of 3.5 µL sample solution with protein concentrations of ~0.075 mg/mL were applied to the grid and blotted using Vitrobot (FEI, Thermo Fisher Scientific, MA, USA) for 4 seconds with -1 offset setting before freezing. The grids were loaded into a Titan Krios (FEI, Thermo Fisher Scientific, MA, USA) microscope and data were collected at a nominal magnification of 29,000X. A movie of 20 frames with a 40 electron dose/ $Å^2$ on the sample were recorded with a K2 direct electron camera with super resolution mode and resulted the unbinned pixel size as 0.848 Å.

A dataset with approximately 19K movies were motion corrected by the MotionCor2 software (*44*). Contrast Transfer Function parameters were determined with the Gctf software (*43*). Projection images with a frequency cutoff at 30 Å of the electron density map from calcitonin receptor (*45*) were used as reference for automatic particle picking by Gautomatch (*46*) and resulted with more than 6 million particle coordinates. 2D classification with Relion2 (*47*) was performed to clean the particle dataset. Classes with protein secondary structural features were kept and subjected to further 3D classification with 8 reference classes. One class that had the most structural details (with ~120K particles) was selected and further processed with an auto-refinement step. This resulted in a density map with 4.0 Å global resolution. Local resolution of the core transmembrane domain of the receptor was further improved to ~3.3 Å by focused refinement. Visual inspection and model building was done with the Coot program (*48*). A published cryoEM GLP-1R structure (PDB:5VAI) (*35*) and the crystal structure of the ECD (PDB:3IOL) (*49*) were used as starting models for the protein components, which were manually adjusted based on the density features. Several rounds of real space refinements were performed with phenix.real_space_refine (*50*) followed by manual local adjusting. Pymol (*51*) was used for visualization and figure generation.

Intraperitoneal Glucose Tolerance Test (IPGTT) in C57BL/6 Mice

All experiments involving animals were conducted in our AAALAC-accredited facilities and were reviewed and approved by Pfizer Institutional Animal Care and Use Committee. Male C57BL/6N mice (22.8–29.2 g body weight; Taconic Biosciences, Hudson, NY) were singly housed and fed standard laboratory rodent chow (LabDiet, PicoLab®Rodent Diet 20, St. Louis, MO). Mice were maintained on a 12-hour light, 12-hour dark cycle and received food and water *ad libitum*. Animals were acclimated to the laboratory environment for a minimum of 7 days prior to study start and acclimated to subcutaneous (SC) vehicle injections for a minimum of 3 days prior to initiation of the IPGTT. On the day prior to the IPGTT, mice were weighed and randomized into treatment groups. In the afternoon prior to the study day, food was removed and mice were fasted overnight. The next morning, mice were administered with vehicle [2% Tween 80:98% (5% (w/v) dextrose in water) (v/v)], liraglutide (0.3 mg/kg, Victoza® pen, Novo Nordisk, Bagsvaerd, Denmark) or PF-06882961 (10 mg/kg) in vehicle via SC injection. Fifteen minutes after vehicle, liraglutide or PF-06882961 in vehicle injection, mice were administered an intraperitoneal (IP) injection of 40% (w/v) dextrose at 2 g/kg. Blood glucose was measured via tail nick using a handheld glucometer (Accu-Chek Aviva, Roche Diabetes Care Inc., Indianapolis, IN) at 16 and 1 minutes prior to and 15, 30, 45, 60, 90 and 120 minutes after administration of dextrose. Blood was collected 1 minute prior to dextrose administration via tail nick and at the end of the study via cardiac stick for analysis of PF-06882961 levels in plasma. Blood samples (approximately 50 µL) were collected, placed into dipotassium EDTA tubes, centrifuged (Eppendorf Centrifuge 5417R, Hamburg, DE) and stored at -80 ºC until analysis of plasma PF-06882961 concentrations. The primary outcome measure for the IPGTT was the serum glucose area under the serum concentration-time curve from $t$ = 0 to 120 minutes ($AUC_{0\text{-}120\ min}$). Statistical analysis of the data was completed using SAS (version 9.4, SAS Institute Inc, Cary, NC) and plotted using GraphPad Prism (version 6.03, GraphPad Software Inc, La Jolla, CA).

Animal Pharmacokinetics Studies

Jugular vein/carotid artery double cannulated male Wistar-Han rats (~250 g), obtained from Charles River Laboratories (Wilmington, MA) and male cynomolgus monkeys (~7 kg) were used for these studies. Animals were fasted overnight and through the duration of the study (1.0 or 2.0 h), whereas access to water was provided *ad libitum*. PF-06882961 was administered intravenously (IV) as a solution (1 mg/mL) in [5% polyethylene glycol 400: 95% (12% (w/v) sulfobutyl-β-cyclodextrin in water), (v/v)] or [10% DMSO: 50% polyethylene glycol 400: 40% water, (v/v/v)] via the tail vein in rats ($n$=4) or femoral vein in monkeys ($n$=2) at a dose of 1.0 mg/kg in a dosing volume of 1 mL/kg. Serial blood samples were collected before dosing and 0.083, 0.25, 0.5, 1.0, 2.0, 4.0, 7.0, and 24 hours after dosing. Urine samples (0–7.0 and 7.0–24

h) were also collected after IV administration to rats and monkeys. The crystalline 2-amino-2-hydroxymethyl-propane-1,3-diol (tris) salt form of PF-06882961 was also administered by oral (PO) gavage to rats (5 and 100 mg/kg at 10 mL/kg) and monkeys (5 mg/kg (5.0 mL/kg) and 100 mg/kg (10 mL/kg) as a homogeneous suspension in [2% Tween 80: 98% (0.5% (w/v) methylcellulose A4M in distilled water), (v/v)]. Blood samples were taken prior to PO administration, then serial samples were collected at 0.083, 0.25, 0.5, 1, 2, 4, 7, and 24 hours after dosing. Blood samples from the pharmacokinetic studies were centrifuged to generate plasma. All plasma samples were kept frozen until analysis. For rat and monkey samples, aliquots of plasma or urine (20-50 μL) were transferred to 96-well blocks, and acetonitrile (150-200 μL) containing verapamil (monkeys) or terfenadine (rat) as internal standard was added to each well. Supernatant was dried under nitrogen and reconstituted with 100 μL water without evaporation. Following extraction, the samples were then analyzed by liquid chromatography tandem mass spectrometry (LC-MS/MS) and concentrations of PF-06882961 in plasma and urine were determined by interpolation from a standard curve.

LC-MS/MS Analysis for Quantitation of PF-06882961

Concentrations of PF-06882961 were determined on an AB Sciex model API 400 or API 5500 LC-MS/MS triple quadrupole mass spectrometer (AB Sciex, Framingham, MA). Analytes were chromatographically separated using a Shimadzu LC-20AD (Shimadzu Scientific Instruments, MD) pump. A CTC PAL autosampler was programmed to inject 10 μL of matrix on Acquity UPLC HSS T3 1.8 μm 2.1 x 50 mm (monkey) (Waters, Milford, MA) or Kinetex 2.6 μ 100 Å (30 mm x 3.00 mm) (rat) (Phenomenex®, Torrance, CA) columns using a mobile phase consisting of water with 0.1% formic acid (solvent A) and acetonitrile with 0.1% formic acid (solvent B) at a flow rate of 0.6–0.7 mL/min. Ionization was conducted in the positive ion mode at the ionspray interface temperature of 400 °C, using nitrogen for nebulizing and heating gas. The ion spray voltage was 5.0 kV and the declustering potential was optimized at 41 eV. PF-06882961 was detected using electrospray ionization in the multiple reaction monitoring mode monitoring for *m/z* transition 556.3→324.2. Corresponding internal standards verapamil and tolbutamide were detected in the multiple reaction monitoring mode monitoring for *m/z* transition 455.2→165.4 and 472.4→436.4, respectively. Analyst® software (SCIEX, Redwood City, CA) was used to measure peak areas, and peak area ratios of analyte to internal standard were calculated. A calibration curve was constructed from the peak area ratios with a weighted linear ($1/x^2$) regression using Watson Laboratory Information Management Systems (LIMS™) software (Thermo Fisher Scientific, Waltham, MA). PF-06882961 standards were fit by least-squares regression of their areas to a weighted linear equation, from which the unknown concentrations were calculated. The dynamic range of the assay was 1.0-5000 ng/mL. Assay performance was monitored by the inclusion of quality control samples with acceptance criteria of ± 30% target values.

Determination of Pharmacokinetic Parameters

Pharmacokinetic parameters in animals were determined using noncompartmental analysis (Watson v.7.4, Thermo Scientific, Waltham, MA). Maximum plasma concentrations ($C_{max}$) of PF-06882961 in plasma after PO dosing in rats and monkeys were determined directly from the experimental data, with $T_{max}$ defined as the time of first occurrence of $C_{max}$. The area under the plasma concentration-time curve from $t$ = 0 to 24 h ($AUC_{0\text{-}24}$) and $t$ = 0 to infinity ($AUC_{0\text{-}\infty}$) was estimated using the linear trapezoidal rule. Systemic plasma clearance ($CL_p$) was calculated as the intravenous dose divided by $AUC_{0–\infty}^{IV}$. The terminal rate constant ($k_{el}$) was calculated by a linear regression of the log-linear concentration-time curve, and the terminal elimination $t_{1/2}$ was calculated as 0.693 divided by $k_{el}$. Apparent steady state distribution volume ($Vd_{ss}$) in animals were determined as the IV or PO dose divided by the product of $AUC_{0–\infty}$ and $k_{el}$. The absolute bioavailability (F) of the PO doses in animals was calculated by using the following equation: $F = AUC_{0–\infty}^{PO}/AUC_{0–\infty}^{IV} \times dose^{IV}/dose^{PO}$. Percentage of unchanged PF-06882961 excreted in urine over 24 h ($A_{e,urine,(0\text{-}24h)}$) was calculated using the following equation: amount (in mg) of PF-06882961 in urine over the 24 h interval post dose/actual amount of PF-06882961 dose administered (mg) x 100%. The renal clearance ($CL_{renal}$) was derived as the ratio of amount of PF-06882961 in urine (in mg) over the 24 h interval post dose/$AUC_{0\text{-}24}$.

Cynomolgus Monkey Intravenous Glucose Tolerance Test (IVGTT)

These studies were conducted in accordance with the current guidelines for animal welfare (National Research Council Guide for the Care and Use of Laboratory Animals, 2011; Animal Welfare Act [AWA], 1966, as amended in 1970, 1976, 1985, and 1990, and the AWA implementing regulations in Title 9, Code of Federal Regulations, Chapter 1, Subchapter A, Parts 1-3). The procedures used in this study have been reviewed and approved by the Institutional Animal Care and Use Committee (IACUC) (Animal Use Procedure No. GTN-2011-00178).

Male cynomolgus macaques (7.5-11.5 kg body weight) were singly housed in stainless steel cages and fed a diet consisting of Certified Hi Fiber Primate Diet (Catalog No 5K91, LabDiet©, St Louis, MO) supplemented with vegetables and fruits. Each animal was identified by an individual tattoo and system generated identification number and cross-referenced to an assigned study animal number. Monkeys were acclimated to the laboratory environment for at least 30 days prior to study and acclimated to the restraining chairs for a period of up to 4 hours. The laboratory environment was maintained at a constant temperature between 19 ºC and 25 ºC, humidity between 30%–70% and a 12-hour light 12-hour dark cycle.

Two studies were performed with each study consisting of a Latin square cross over design that was balanced and uniform within sequences. In each study, 8 animals were randomized to one of the sequences. Experiments were performed with 4 animals tested each day with each animal receiving a different treatment. A washout period of at least 72 hours was provided after each treatment.

Study 1 Treatments

| Treatment | Bolus | Infusion |
|---|---|---|
| Vehicle (IV) | 0.3 mL/kg, IV | 1.0 mL/kg/hr, IV |
| liraglutide | 0.03 mg/kg, SC | |
| PF-06882961 (Low IV Dose) | 0.0121 mg/kg, IV | 0.102 mg/kg/hr, IV |
| PF-06882961 (High IV Dose) | 0.121 mg/kg, IV | 1.02 mg/kg/hr, IV |

Study 2 Treatments

| Treatment | Bolus | Infusion |
|---|---|---|
| Vehicle (Oral and IV) | 5.0 mL/kg, PO and<br>0.3 mL/kg, IV | ----<br>1.0 mL/kg/hr, IV |
| PF-06882961 (X-Low IV Dose) | 0.00303 mg/kg, IV | 0.0255 mg/kg/hr, IV |
| PF-06882961 (Med IV Dose) | 0.0303 mg/kg, IV | 0.255 mg/kg/hr, IV |
| PF-06882961 (Oral Dose) | 100 mg/kg, PO | ---- |

In study 1, animals receiving liraglutide will also be given a SC injection of liraglutide the afternoon prior to the IVGTT, all others will receive a SC injection of [1.35% propylene glycol: 98.65% Dulbecco® phosphate buffered saline (without calcium chloride and magnesium chloride), (v/v)] vehicle the afternoon prior to the IVGTT. Animals were fasted overnight prior to the IVGTT. On the morning of each IVGTT at approximately 6:00 AM, each monkey was briefly placed in a quick-release primate chair restraint and administered a SC injection of liraglutide or [1.35% propylene glycol: 98.65% Dulbecco® phosphate buffered saline (without calcium chloride and magnesium chloride), (v/v)] vehicle. A blood sample (1 mL) was collected, placed into dipotassium EDTA tubes on ice, centrifuged, and stored at -20 ºC until analysis for plasma concentration of PF-06882961 to ensure there was no carry-over from a previous study sequence. Following the blood sample, the monkeys were returned to their home cage. Approximately 1 hour later, the monkeys were placed in a chair restraint (Plas Labs Inc., Model 515 SASR, Rhesus Chair with custom modifications, Lansing, MI) and a venous catheter inserted for IV infusion. A baseline blood sample was withdrawn and the IV bolus and IV infusion of PF-06882961 in vehicle or vehicle alone [5% polyethylene glycol 400: 95% (12% (w/v) sulfobutylether-β-cyclodextrin (SBECD) in deionized water) (v/v)], pH adjusted to pH 7.0-7.4, were initiated. The IVGTT was initiated 30 minutes after the start of the IV drug infusion. Blood samples were collected at 30, 15 and 0 minutes prior to administration of the glucose dose and 30 minutes post glucose dose. Samples were placed into dipotassium EDTA tubes on ice, centrifuged and stored at -20 ºC until analysis for plasma concentration of PF 06882961. Plasma concentrations of PF-06882961 were determined by the Pfizer Department of Pharmacokinetics, Dynamics and Metabolism.

In study 2, which included oral dosing, monkeys were administered either PF-06882961 in vehicle or vehicle alone [2% Tween 80: 98% (0.5% [w/v] methylcellulose A4M in deionized water), (v/v), containing 1.5 molar equivalents of NaOH, pH 6.0-7.0], via oral gavage at approximately 6:00 AM. Oral gavage of vehicle in monkeys in the IV PF-06882961 or vehicle control sequence was to control for the dosing activity itself. Following the dose, monkeys were returned to their home cage. Approximately 1 hour post oral dose, the monkeys were placed in a chair restraint and a venous catheter was inserted for IV infusion. A baseline blood sample was withdrawn, and the IV bolus and IV infusion of PF-06882961 in vehicle or vehicle alone [5% polyethylene glycol 400: 95% (12% sulfobutylether-β-cyclodextrin (SBECD) in deionized water), (v/v), pH adjusted to pH 7.0-7.4], was initiated. The IVGTT was initiated 30 minutes after the start of the IV drug infusion.

For the IVGTT, glucose was administered by IV bolus at a dose of 250 mg/kg using a sterile 50% (w/v) dextrose in deionized water solution at a dose volume of 0.5 mL/kg. The glucose bolus was injected over approximately 20 seconds. Blood samples (1 mL) were collected from each animal on each day of the IVGTT at 15 and 0 minutes prior to administration of the glucose and at 2, 5, 7, 10, 15, 20 and 30 minutes post administration of the glucose dose. Samples were collected via syringe from catheters, placed into serum separator tubes with a clot activator on ice, centrifuged (Jouan GR412 Centrifuge, Thermo Fisher Scientific, Waltham, MA) and stored at 4 °C until analysis on a clinical analyzer for glucose (Siemens Advia 1800 Chemistry Analyzer, Berlin, Germany) and insulin (Siemens Advia Centaur XP Immunoassay System, Berlin, Germany).

The primary outcome measure for the IVGTT was the insulin area under the serum concentration-time curve from $t = 0$ to 30 minutes ($AUC_{0\text{-}30\ min}$). For analysis, the insulin $AUC_{0\text{-}30\ min}$ endpoint was transformed with the natural logarithm (ln) so that it followed a normal distribution. The insulin $AUC_{0\text{-}30\ min}$ was analyzed with a linear crossover model including fixed effect terms for period, treatment, carry over and a random animal effect. The model also included covariates for age, weight and baseline insulin. The plasma concentration of PF-06882961 for each individual monkey during the IVGTT at each dose was determined by taking the mean of the concentration at the start of the IVGTT (0 min) and at the end of the IVGTT (30 min). The mean plasma concentration of PF-06882961 for each dose group was determined by calculating the geometric mean of the individual animal plasma concentrations of PF-06882961 during the IVGTT. The rate of glucose clearance from serum (K-value) was calculated from the 5- and 20-minute serum glucose concentrations. Statistical analysis of the data was completed using SAS (version 9.4, SAS Institute Inc, Cary, NC) and plotted using GraphPad Prism (version 6.03, GraphPad Software Inc, La Jolla, CA).

Cynomolgus Monkey Food Intake

Male cynomolgus macaques (7.3 – 11.0 kg body weight) were singly housed in stainless steel cages and fed a diet consisting of Certified Hi-Fiber Primate Diet (~72.9 kcal/biscuit, Catalog No. 5K91, LabDiet®, St Louis, MO) supplemented with apple and peanuts. Monkeys were fed on a daily schedule that included: 5 biscuits at 7:30 AM, 3 biscuits + ½ apple at 10:30 AM and 3 peanuts at 12:00 PM. An assessment of the portion consumed was recorded just before the 10:30 AM feed and again at 2:00 PM when any remaining food was removed. Animals did not have access to food overnight. Water was available *ad libitum*. Twelve cynomolgus monkeys were randomized to one of two groups (1:1), stratified by body weight and baseline food intake. Food intake was monitored for at least 10 days prior to dosing (baseline), during 2 days of once-daily injections of PF-06882961 in vehicle or vehicle alone, and until food intake returned to baseline levels. Monkeys were acclimated to vehicle injections for at least 5 days prior to PF-06882961. Each morning at approximately 6:30 AM, monkeys were briefly restrained at the front of their home cages for SC administration of 26 mgA/mL PF-06882961 (2.9 mg/kg) in vehicle or vehicle alone. Each monkey received a dose volume of 0.11 mL/kg PF-06882961. The vehicle used in this study was [2% Tween 80: 98% (5% (w/v) dextrose in deionized water), (v/v), pH adjusted to pH 7.6 (+/- 0.1)]. Monkeys were acclimated to the restraint and vehicle injections for 4 days prior to the start of PF-06882961 dosing. At 2:00 PM after completion of the food assessment on the second day of dosing, blood samples were collected via syringe from a venous catheter and placed into 0.5 mL dipotassium EDTA blood collection vials (SAI Infusion Technologies Micro 500, Catalog No M500-E, Lake Villa, IL) on ice. Blood samples were centrifuged, and the resulting plasma was aliquoted and stored at -80 ºC. Plasma concentrations of PF-06882961 were determined using the LC-MS/MS method previously described.

Food intake during treatment was analyzed as a least squares mean change from baseline using a repeated measures model including fixed effects terms for treatment, day, random animal effect, age and body weight as covariates. Statistical analysis of the data was completed using SAS (version 9.4, SAS Institute Inc, Cary, NC) and plotted using GraphPad Prism (version 6.03, GraphPad Software Inc, La Jolla, CA).

Study Design for First-in-Human Study in Healthy Adult Participants

This phase 1 study [NCT03309241] was conducted at the sponsor's Clinical Research Unit in New Haven, CT, USA, from October 17, 2017 to February 21, 2018. The protocol was approved by the study center's institutional review board located in New Haven, CT, USA. All participants provided informed consent before screening. The study was conducted in compliance with ethical principles of the Declaration of Helsinki and International Council for Harmonization Good Clinical Practice guidelines. All local regulatory requirements were followed.

The design for this first-in-human study was investigator- and participant-blinded, Sponsor-open, randomized, single ascending oral dose, 4-period cross-over in 2 interleaving cohorts with placebo substitution. The interleaving design enabled both within- and between-participant assessments. Data from Cohorts 1 & 2, in which participants received a tablet formulation of PF-06882961 or matching placebo with water, following a ≥10-hour overnight fast or following a high-fat breakfast, are presented in this manuscript. Based on emerging data from Cohorts 1 and 2, the Sponsor elected to enroll Cohort 3, and participants were randomized to receive PF-

06882961 or placebo administered as split doses in the fed state in a 4-period cross-over design to better characterize safety, tolerability and pharmacokinetics of PF-06882961.

Participants were required to be healthy adults aged 18–55 years (females were of nonchildbearing potential), with a body mass index of 17.5–30.5 kg/m$^2$ and a body weight of >50 kg. Exclusion criteria included history of clinically significant hematologic, renal, endocrine, pulmonary, gastrointestinal, cardiovascular, hepatic, psychiatric, neurologic, or allergic disease and conditions affecting drug absorption.  Randomization was performed using a sponsor-provided randomization schedule.

A total of 25 participants were randomized. Each participant received up to 4 single oral doses of PF-06882961 and up to 2 placebo doses.  In a given participant, there was an interval of at least 7 days between consecutive study periods to allow for washout of PF-0688296 and review safety and pharmacokinetics data from each dose level before decisions were made on the next dose. Participants who discontinued for non-safety related reasons prior to completion of the study may have been replaced at the discretion of the principal investigator and Sponsor. The starting dose of 3 mg was derived from nonclinical information on pharmacokinetics and metabolism of PF-06882961. Blood samples for safety (including fasting serum glucose) and pharmacokinetic assessments were collected at intervals ≤48 hours post dose.

All observed and self-reported adverse events (AEs), any clinically significant changes in physical examination results, and abnormal test results were recorded. Laboratory evaluations included hematology, chemistry, and urinalysis. Twelve-lead electrocardiogram (ECG) and vital signs were monitored throughout.

A sample size of 8 participants per cohort was chosen to minimize the first exposure in humans of a new chemical entity while providing safety information for each dose. The safety analysis set comprised all participants receiving ≥1 dose of study drug. AEs, ECGs, blood pressure, pulse rate, and safety laboratory data were reviewed on an ongoing basis. Safety data were summarized descriptively. No formal inferential statistics were applied to plasma pharmacokinetic data, and pharmacokinetic parameters were summarized descriptively by dose.

Plasma samples were analyzed for PF-06882961 concentrations at Syneos Health (formerly inVentiv Health Clinical Lab Inc, located in Princeton, New Jersey, US) using a validated sensitive and specific LC-MS/MS method in compliance with the Sponsor SOPs. Plasma specimens were required to be stored at approximately -80 °C until analysis and assayed within the 150 days of established stability data generated during validation.  Calibration standard responses were linear over the range of 0.100 to 100 ng/mL using a weighted (L/concentration$^2$) linear least squares regression. Those samples with concentrations above the upper limits of quantification were adequately diluted into calibration range. The lower limit of quantification (LLOQ) for PF-06882961 was 0.100 ng/mL. Clinical specimens with plasma PF-06882961 concentrations below the LLOQ were reported as "<0.100 ng/mL". The between-day assay accuracy, expressed as percent relative error (%RE), for quality control (QC) concentrations, ranged from -4.00% to 7.00% for the low, medium, high, and diluted QC samples.  Assay precision, expressed as the between-day percent coefficient of variation (%CV) of the mean estimated concentrations of QC samples was ≤8.13% for low (0.300 ng/mL), medium (35.0 ng/mL), high (75.0 ng/mL), and diluted (1500 ng/mL) concentrations.

Fasting serum glucose was measured pre-dose and 24 hours post-dose in study participants who received single doses of PF-06882961 ranging from 3 mg to 300 mg or placebo. To permit comparison of the effect of PF-06882961 on fasting glucose versus placebo, only participants who received both placebo and doses of PF-06882961 were included in the analysis. Serum glucose samples were analyzed using the photometric module of a random access chemistry and immunochemistry analyzer (Roche Cobas 6000) in compliance with sponsor SOPs at the Pfizer Clinical Research Unit in New Haven, CT. Samples were analyzed immediately according to manufacturer's instructions.

**Supplementary figures and tables**

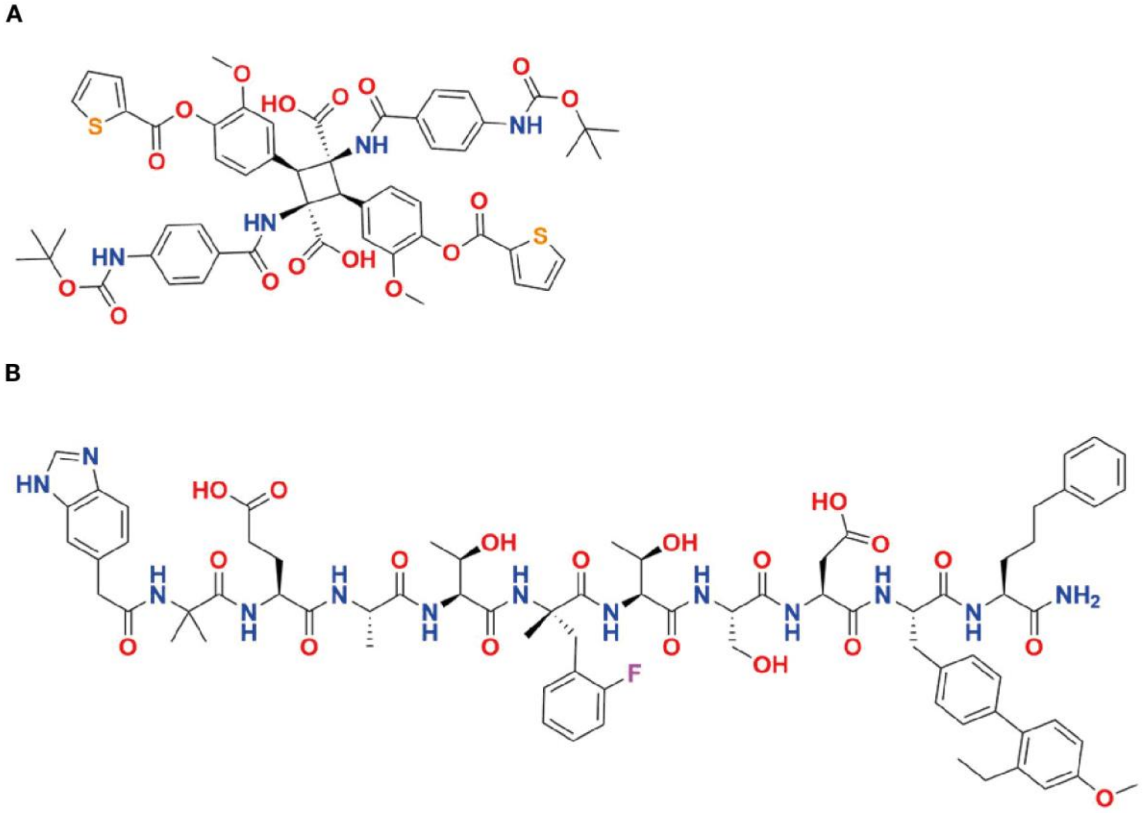


Fig. S1.

**Structures of reference compounds.** (**A**) Structure of **Boc-5**. (**B**) Structure of peptide **1**.

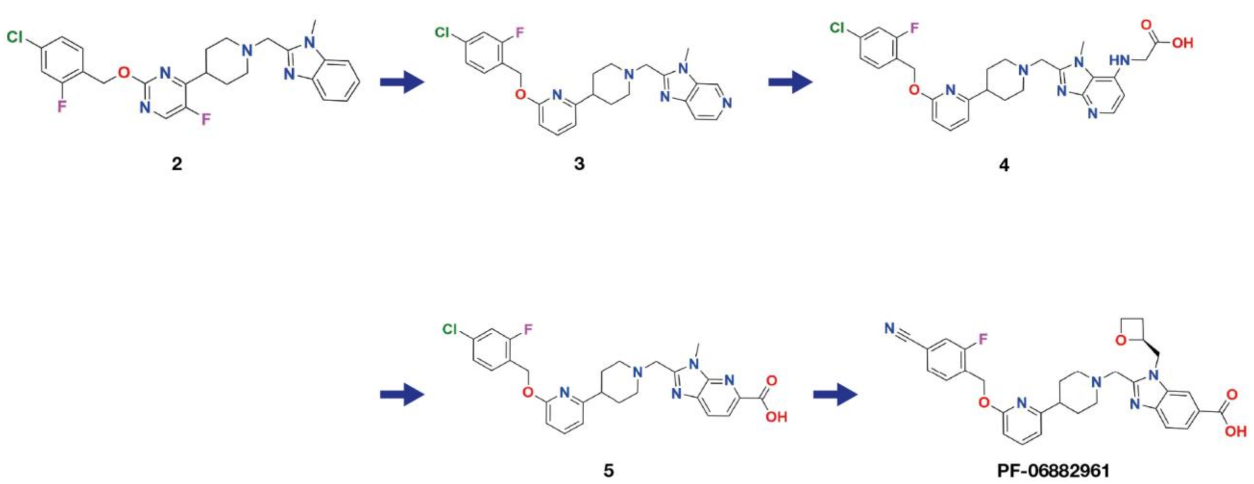


Fig. S2.

**Key compounds in the progression of the small molecule HTS hit 2 to clinical candidate (PF-06882961).**

HTS, high throughput screening

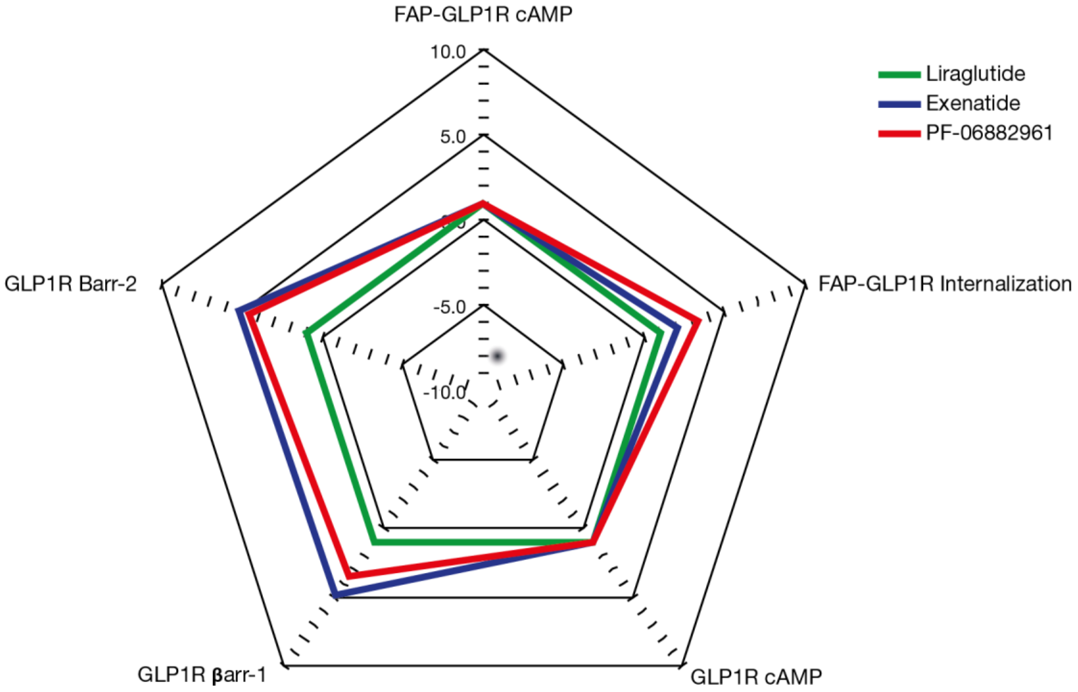


Fig. S3.

**Webs of bias for exenatide and PF-06882961, relative to liraglutide.** The log($E_{max}$/$EC_{50}$) ratio extracted from standard concentration-response data is used to calculate bias factors through normalization of the log($E_{max}$/$EC_{50}$) ratio to a reference ligand (Liraglutide) and reference pathway (cAMP accumulation); for further details, see Methods.

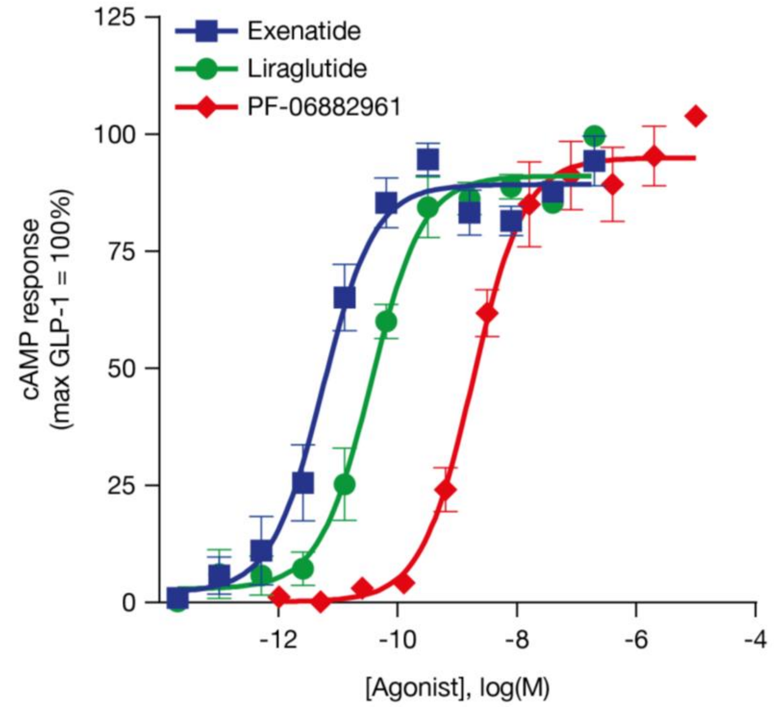


Fig. S4.

**Functional activity of PF-06882961 at FAP-tagged human GLP-1R stably expressed in HEK293 cells, as assessed using cAMP**. Data represent the mean ± SEM. from 3 independent experiments, each performed in triplicate.

cAMP, cyclic adenosine monophosphate; FAP, fluorogen-activated protein; GLP-1, glucagon-like peptide-1 (7-36) amide; GLP-1R, glucagon-like peptide-1 receptor; SEM, standard error of the mean

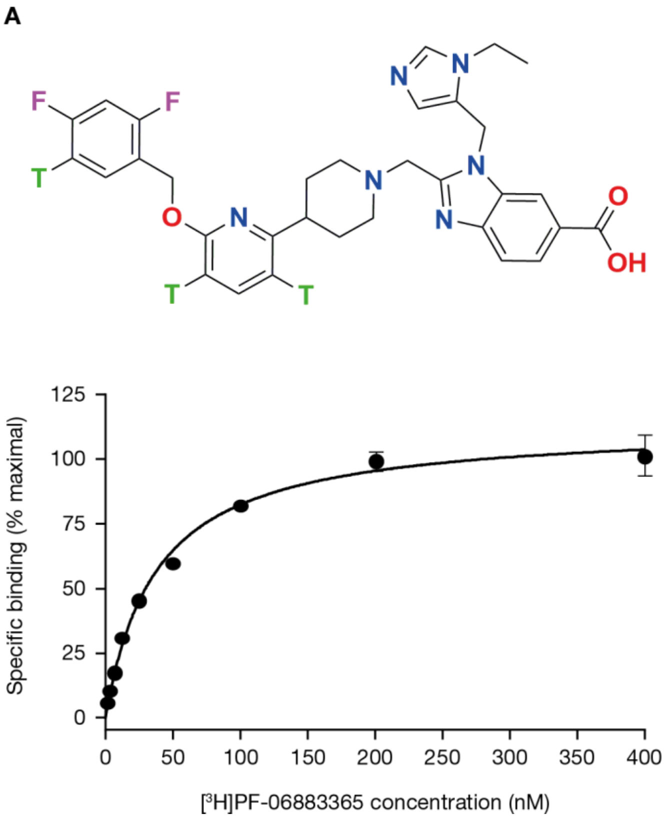


Fig. S5.

**Binding affinity of PF-06882961, as evaluated using a competition binding assay based on radiolabeled small molecule agonist [$^{3}$H]PF-06883365.** (**A**) Structure of the small molecule agonist radioligand [$^{3}$H]PF-06883365. (**B**) Saturation binding analysis for [$^{3}$H]PF-06883365 showed that [$^{3}$H]PF-06883365 binds plasma membranes from CHO cells stably expressing a high density of hGLP-1R with an averaged Kd of 38 nM and Bmax of 5470 fmol/mg.

A

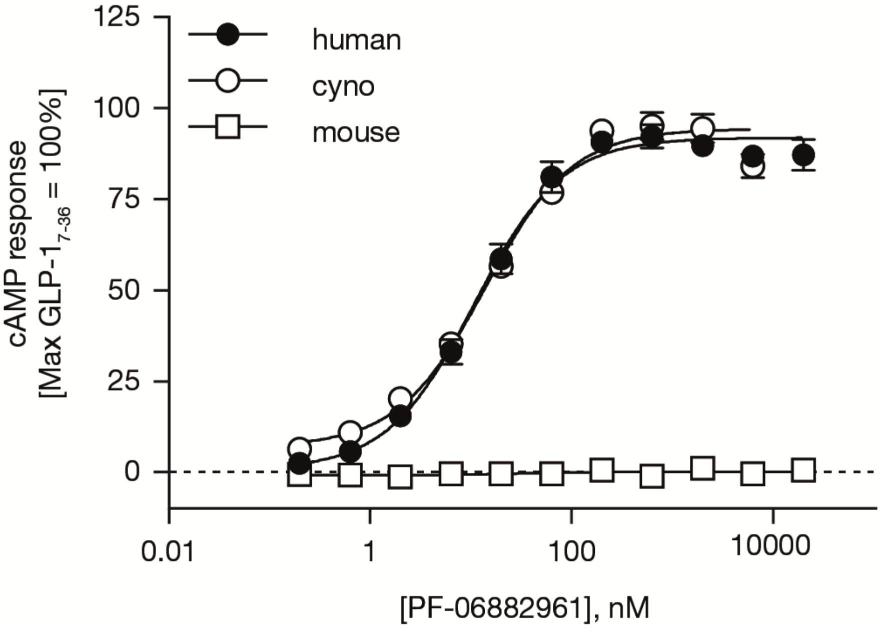


B

| Species | $EC_{50}$ (nM), free (95% CI) | $E_{max}$ (%) (95% CI) | n |
|---|---|---|---|
| Cynomolgus macaque | 4.4 (2.3-7.5) | 95 (80.6-109.3) | 5 |
| Rat | >20,000 | N.M. | 3 |
| Rabbit | >20,000 | N.M. | 3 |
| Mouse | >20,000 | N.M. | 3 |

Fig. S6.

**Functional activity of PF-06882961 at GLP-1R stably expressed by CHO cells, as assessed using cAMP.** (A) cAMP accumulation in CHO cells expressing either the human, cynomolgus macaque or mouse GLP-1. Data represent the mean ± SEM. (B) $EC_{50}$ values of PF-06882961 at the cynomolgus macaque, rat, rabbit, and mouse GLP-1R stably expressed in CHO cells. The $EC_{50}$ value value is expressed as the geometric mean with 95% confidence interval while the efficacy value is reported as the arithmetic mean with standard deviation for the number of replicates indicated. cAMP = 3'-5'-Cyclic adenosine monophosphate; CHO, Chinese hamster ovary; CI = 95% Confidence interval; $EC_{50}$ = Concentration required for 50% effect; GLP-1, glucagon-like peptide-1 (7-36) amide; GLP-1R = Glucagon-like peptide-1 receptor; n = Number of determinations; N.M. = Non-measureable; nM = Nanomolar; SD = Standard deviation; SEM, standard error of the mean.

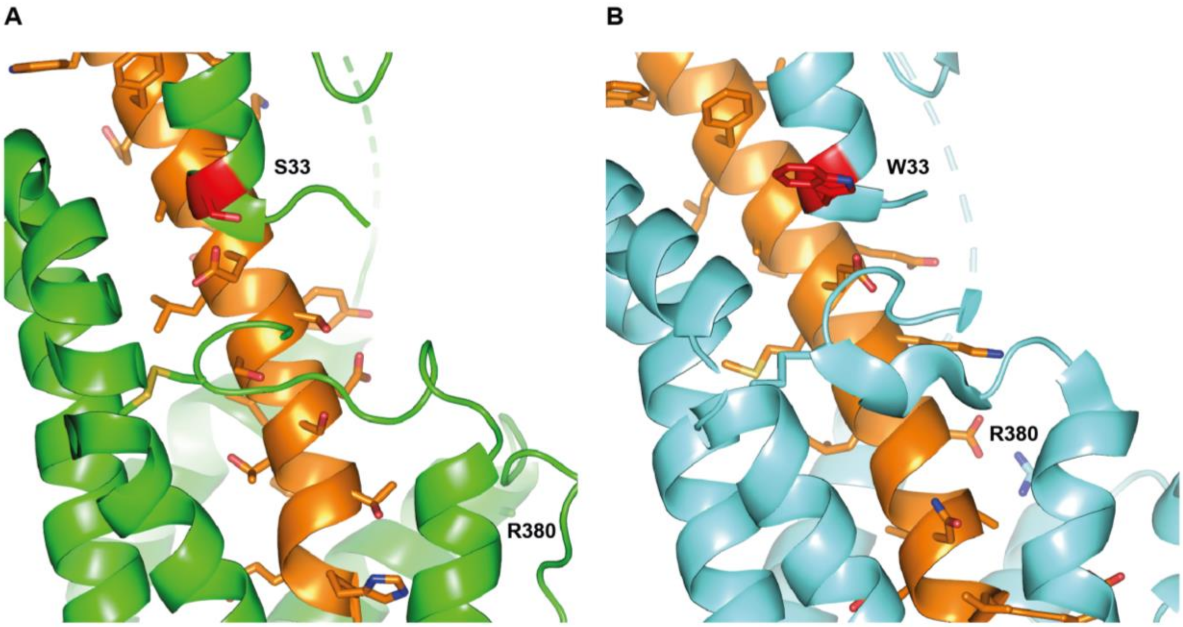


<u>Fig. S7.</u>

**Cryo-EM structures of peptides bound to GLP-1R reveal residue 33 extending towards solvent.** Cryo-EM structures of (A) GLP-1 bound to rabbit GLP-1R (PDB=5VAI) and (B) exendin-P5 bound to human GLP-1R (PDB=6BJ3); ECD residue 33 extends towards solvent in both structures.

PDB, Protein Data Base; S, serine; W, tryptophan.

Table S1.

Summary of BETP-sensitized and non-sensitized agonist data for BOC-5 and peptide **1**.[a]

| Compound | cAMP + BETP[b] | | | cAMP – BETP[b] | | | β-Arr 2 + BETP[c] | | | β-Arr 2 – BETP[c] | | |
|---|---|---|---|---|---|---|---|---|---|---|---|---|
| | $EC_{50}$ (nM) | Efficacy (%) | N | $EC_{50}$ (nM) | Efficacy (%) | N | $EC_{50}$ (nM) | Efficacy (%) | N | $EC_{50}$ (nM) | Efficacy (%) | N |
| BOC5 | 270 (190-380) | 74 ± 15 | 6 | >20000 | n.a. | 2 | n.t. | | | n.t. | | |
| **1** | 0.052 (0.022-0.12) | 98 ± 11 | 3 | 0.37 (0.23-0.60) | 95 ± 9.4 | 4 | 5.6 (2-16) | 67 ± 4.2 | 3 | 210 (160-290) | 11 ± 1.1 | 4 |

[a]The $EC_{50}$ value is expressed as the geometric mean with 95% CI while the efficacy value is reported as the arithmetic mean ± SD from the number of replicates indicated, each performed in duplicate. Efficacy data are presented relative to the response of 1 µM GLP-1 (100%). [b]cAMP production measured in the SA ± BETP. [c]β-Arr recruitment measured in DiscoveRx cell-line; conducted in Assay Media.

β-Arr, β-arrestin; BETP, 4-(3-(benzyloxy)phenyl)-2-ethylsulfinyl-6-(trifluoromethyl)pyrimidine; cAMP, cyclic adenosine monophosphate; CI, confidence interval; GLP-1, glucagon-like peptide-1; N, number of replicates; n.a., not applicable; n.t., not tested; SA, screening assay; SD, standard deviation

Table S2.

Agonist-mediated cAMP release in the BETP-sensitized SA, non-sensitized SA, and CS cell lines.[a]

| Compound | SA cAMP + BETP[b] | | | SA cAMP – BETP[b] | | | CS cAMP[c] | | |
|---|---|---|---|---|---|---|---|---|---|
| | $EC_{50}$ (nM) | Efficacy (%) | N | $EC_{50}$ (nM) | Efficacy (%) | N | $EC_{50}$ (nM) | Efficacy (%) | N |
| **2** | >9900 (5500-18000) | 96 ± 11 | 8 | >20000 | n.a. | 7 | >20000 | n.a. | 2 |
| **3** | 77 (62-95) | 86 ± 12 | 43 | 2600 (2100-3100) | 79 ± 14 | 47 | >20000 | n.a. | 4 |
| **4** | 44 (23-85) | 90 ± 15 | 6 | 4600 (3300-6300) | 100 ± 1.6 | 6 | >20000 | n.a. | 3 |
| **5** | 6.7 (5.6-7.9) | 84 ± 10 | 81 | 95 (84-110) | 78 ± 9.5 | 105 | 2100 (1900-2400) | 87 ± 15 | 87 |
| PF-06882961 | 0.71 (0.35-1.4) | 110 ± 6.0 | 3 | 1.1 (0.61-1.9) | 79 ± 6.2 | 5 | 13 (9.9-17) | 110 ± 13 | 22 |
| PF-06883365 | n.t. | | | 0.75 (0.34-1.7) | 81 ± 7.3 | 7 | 8.6 (4-19) | 98 ± 11 | 9 |
| Exenatide | n.t. | | | 0.061 (0.035-0.11) | 91 ± 14 | 6 | 0.11 (0.075-0.17) | 120 ± 15 | 14 |
| Liraglutide | n.t. | | | 0.39 (0.25-0.60) | 94 ± 9.9 | 6 | 0.95 (0.55-1.6) | 120 ± 12 | 12 |

[a]The $EC_{50}$ value is expressed as the geometric mean with 95% CI while the efficacy value is reported as the arithmetic mean ± SD from the number of replicates indicated, each performed in duplicate. Efficacy data are presented relative to the response of 1 µM GLP-1(7-36)amide (100%). [b]cAMP production measured in the SA ± BETP. [c]cAMP production measured in CS cell line ± BETP
BETP, 4-(3-(benzyloxy)phenyl)-2-ethylsulfinyl-6-(trifluoromethyl)pyrimidine; cAMP, cyclic adenosine monophosphate; CI, confidence interval; CS, candidate selection; GLP-1, glucagon-like peptide-1; N, number of replicates; n.a., not applicable; n.t., not tested; SA, screening assay; SD, standard deviation

Table S3.

**Agonist-mediated β-Arr recruitment.** BETP potentiates β-Arr recruitment by weak agonists in DiscoveRx cell line.[a,b]

| Compound | β-Arr 2 + BETP | | | β-Arr 2 – BETP | | |
|---|---|---|---|---|---|---|
| | $EC_{50}$ (nM) | % Effect | N | $EC_{50}$ (nM) | % Effect | N |
| **3** | 9600 (6600-14000) | 100 ± 3.8 | 7 | >30000 | n.a. | 1 |
| **4** | 1500 (1400, 1600) | 23 ± 5.1 | 2 | >30000 | n.a. | 1 |
| **5** | 2800 (740-11000) | 99 ± 11 | 3 | >30000 | n.a. | 3 |
| Liraglutide | n.t. | | | 17 (8.7-34) | 120 ± 26 | 6 |

[a]The $EC_{50}$ value is expressed as the geometric mean with 95% CI while the efficacy value is reported as the arithmetic mean ± SD from the number of replicates indicated, each performed in duplicate. For N = 2, the two replicates are listed. Efficacy data are presented relative to the response of 1 µM GLP-1 (100%).

[b]Conducted in Assay Media.

β-Arr, beta-arrestin; BETP, 4-(3-(benzyloxy)phenyl)-2-ethylsulfinyl-6-(trifluoromethyl)pyrimidine; CI, confidence interval; GLP-1, glucagon-like peptide-1; GLP-1, glucagon-like peptide-1; N, number of replicates; n.a., not applicable; n.t., not tested; SD, standard deviation

Table S4.

In vitro disposition and hERG inhibition with small molecule GLP-1R agonists.

| Compound | logD pH 7.4 | HLM $CL_{int}$ (mL/min/kg)[c] | hHEP $CL_{int}$ (µL/min/million)[c] | hERG[d] $IC_{50}$ (µM) |
|---|---|---|---|---|
| **2** | 5.6[a] | >140 | n.t. | 5.4 |
| **3** | 5.7[a] | 130 | n.t | 5.6 |
| **4** | 2.3[b] | <11 | 37 | >10 |
| **5** | 2.0[b] | 28 | 31 | >100 |
| PF-06882961 | 1.8[b] | <10 | 6.9 | 4.3 |

[a]eLogD was measured at pH 7.4 using the previously described RP-HPLC method (*52*). [b]Shake-flask logD (SFlogD) was measured at pH 7.4 using the previously described shake-flask method (*53*). [c]$CL_{int}$ refers to total intrinsic metabolic clearance obtained from scaling in vitro half-lives of test compounds in human liver microsomes (HLM) or cryopreserved human hepatocytes (hHEP) as previously described (*54*). [d]Inhibition of the hERG channel in a patch-clamp assay.

$CL_{int}$, intrinsic clearance; hHEP, human hepatocytes; HLM, human liver microsomes

Table S5.

**Agonist-mediated β-Arr recruitment and receptor internalization.** Recruitment of β-Arr 1 and 2 by agonists in DiscoveRx cell line and internalization of FAP-tagged GLP-1R in HEK293 cells.[a]

| Compound | β-Arr 1[b] | | | β-Arr 2[b] | | | Internalization | | |
|---|---|---|---|---|---|---|---|---|---|
| | $EC_{50}$ (nM) | % Effect | N | $EC_{50}$ (nM) | % Effect | N | $EC_{50}$ (nM) | % Effect | N |
| PF-06882961 | 760 (530-1100) | 41 ± 0.98 | 3 | 490 (310-760) | 36 ± 9.8 | 3 | 230 (160-350) | 83 ± 3.2 | 3 |
| PF-06883365 | 250 (200-330) | 50 ± 11 | 3 | 610 (180-2100) | 50 ± 9.8 | 6 | n.t. | | |
| Exenatide | 14 (9.2-23) | 70 ± 9.5 | 4 | 9 (7.4-11) | 75 ± 7.6 | 4 | 0.60 (0.42-0.84) | 125 ± 2.8 | 3 |
| Liraglutide | 34 (19-63) | 96 ± 5.3 | 4 | 20 (15-26) | 99 ± 1.4 | 4 | 1.8 (1.4-2.5) | 117 ± 2.6 | 3 |

[a]Conducted in buffer containing 0.1% BSA. [b]The $EC_{50}$ value is expressed as the geometric mean with 95% CI while the efficacy value is reported as the arithmetic mean ± SD from the number of replicates indicated, each performed in duplicate. For N = 2, the two replicates are listed. Efficacy data are presented relative to the response of 1 µM GLP-1 (100%) for β-Arr assays and relative to the maximal effect of GLP-1 (100%) for receptor internalization.

β-Arr, beta-arrestin; CI, confidence interval; FAP, fluorogen-activated protein; GLP-1, glucagon-like peptide-1; N, number of replicates; n.t., not tested; SD, standard deviation

Table S6.

**Competition binding affinities.**

| Compound | $[^{125}I]$-GLP-1 binding | | $[^{3}H]$-PF-06883365 binding | |
|---|---|---|---|---|
| | $K_i$ (nM)[a] | N | $K_i$ (nM)[a] | N |
| PF-06882961 | 360 (140-920) | 4 | 80 (62-91) | 4 |
| PF-06883365 | n.t. | - | 51 (43.-61) | 16 |
| Exenatide | 0.092 (0.073-0.12) | 17 | 0.053 | 1 |
| Liraglutide | 4.4 (3.4-5.5) | 19 | 6.3 | 1 |

[a]The $K_i$ value is expressed as the geometric mean with 95% CI performed in duplicate. For N = 1, the replicates are listed.
CI, confidence interval; GLP-1, glucagon-like peptide-1; Ki, concentration required to produce half maximum inhibition; N, number of replicates; n.t., not tested

Table S7.

**Preclinical pharmacokinetic parameters of PF-06882961**

| Species[a] | Dose (mg/kg) | $C_{max}$ (ng/mL) | $T_{max}$ (h) | $AUC_{0-\infty}$ (ng·h/mL) | $CL_p$ (mL/min/kg) | $Vd_{ss}$ (L/kg) | $t_{1/2}$ (h) | Oral F (%)[b] |
|---|---|---|---|---|---|---|---|---|
| Rat | 1.0 (IV) | ---- | ---- | 296 ± 39.8 | 57.3 ± 8.68 | 0.86 ± 0.38 | 1.13 ± 0.84 | ---- |
|  | 5.0 (oral)[b] | 141 (153, 128) | 0.5 (0.5, 0.5) | 168 (151, 184) | ---- | ---- | 0.63 (0.46, 0.79) | 11 (10, 12) |
|  | 100 (oral)[b] | 2820 (2580, 3060) | 0.75 (0.5, 1.0) | 11900 (10300, 13500) | ---- | ---- | 2.37 (2.25, 2.49) | 39 (37, 44) |
| Monkey | 1.0 (IV) | ---- | ---- | 1240 (1390, 1080) | 13.8 (12.0, 15.5) | 0.266 (0.268, 0.264) | 1.89 (2.03, 1.74) | ---- |
|  | 5.0 (oral)[b] | 68.7 (86.3, 51.0) | 1.5 (2.0, 1.0) | 303 (363, 242) | ---- | ---- | 6.92 (5.71, 8.11) | 5.0 (4.0, 6.0) |
|  | 100 (oral)[b] | 1150 ± 715 | 3.3 ± 2.5 | 11000 ± 3500 | ---- | ---- | 6.37 | 9.0 |

[a]All experiments involving animals were conducted in our AAALAC-accredited facilities and were reviewed and approved by Pfizer Institutional Animal Care and Use Committee. Pharmacokinetic parameters were calculated from plasma concentration–time data and are reported as mean (± SD for *n* = 3–4 and maximum values for *n* = 2). All pharmacokinetics studies were conducted in males of each species (Wistar rats and cynomolgus monkey). IV doses for PF-06682961 were administered as a solution in [5% polyethylene glycol 400: 95% (12% (w/v) sulfobutyl-β-cyclodextrin in water), (v/v)] or [10% DMSO: 50% polyethylene glycol 400: 40% deionized water, (v/v/v)]. Oral pharmacokinetics studies were conducted in the fasted state using the crystalline-free form of the 2-amino-2-hydroxymethyl-propane-1,3-diol (tris) salt form of PF-06882961. For oral pharmacokinetics studies, PF-06882961-tris salt was formulated in [2% Tween 80: 98% (0.5% (w/v) methyl cellulose A4M in distilled water), (v/v)]. [b]PF-06882961-tris salt.
AUC, area under the curve; $CL_P$, plasma clearance; IV, intravenous; Oral F, oral bioavailability; SD, standard deviation; $Vd_{ss}$, steady-state volume of distribution.

Table S8.

Broad panel screening of PF-06882961.

| Type of Test | Test cells/Tissues | Results |
|---|---|---|
| In vitro pharmacology binding assays, cellular and nuclear receptor functional assays, and enzyme assays – 60+ receptors, ion channels, transporters, enzymes. Single point at 10 µM with dose-response follow up. | Cell lines, cell membranes from brain and endogenous cells, and recombinant proteins from human and rat | Significant inhibition at $Na^+$ channel (60.7%; $IC_{50}$ = 7.6 µM, Ki = 6.8 µM), $Cl^-$ channel (80%; $IC_{50}$ = 3.5 µM, Ki = 2.9 µM), and 5-HT1B receptor (60.5%; $EC_{50}$ = 5.0 µM). |
| Assessment of activity at phosphodiesterase (PDE) subtypes in a dose response (0.1 nM to 200 µM). | Human PDE subtypes (1B1, 2A1, 3A1, 4D3, 5A1, 6 [bovine], 9A1, 10A1, 11A4, 7B, 8B). | PDE3A1 inhibition ($IC_{50}$ = 2342 nM)<br>PDE10A1 inhibition ($IC_{50}$ = 215 nM)<br>For all other PDEs: $IC_{50}$ > 10 µM. |

PDE, phosphodiesterase

Table S9.

Class-B GPCR selectivity of PF-06882961.

| Assay | $IC_{50}$ or $EC_{50}$ (nM) | % Effect | n |
|---|---|---|---|
| hGIPR, cAMP, agonist mode | >10,000 | N.M. | 4 |
| hGIPR, cAMP, antagonist mode | >10,000 | N.A. | 4 |
| hGCGR, cAMP, agonist mode | >20,000 | N.M. | 3 |
| hGCGR, cAMP, antagonist mode | >20,000 | N.A. | 2 |
| hGLP-2R, cAMP, agonist mode | >20,000 | N.M. | 3 |
| hGLP-2R, cAMP, antagonist mode | 3,600 | N.A. | 2 |

Agonist and antagonist in vitro potency of PF-06882961 at the human GIP-R, GLP-2R and GCG-R, as measured using cAMP production assays.

cAMP, cyclic adenosine monophosphate; GCGR, glucagon receptor; GIPR, glucose-dependent insulinotropic peptide receptor; GLP-2R, glucagon-like peptide-2 receptor; GPCR, G-protein coupled receptor; h, human; N.M., non-measurable; N.A., not applicable

Table S10.

**Safety of PF-06882961.** Summary of treatment-emergent adverse events in healthy human volunteers in FIH study (Cohorts 1 and 2) – all causality (treatment-related)

| | Placebo [fasted] | Placebo [solution] | Placebo [fed] | 3 mg | 10 mg | 25 mg [solution] | 30 mg | 100 mg [fasted] | 100 mg [fed] | 300 mg |
|---|---|---|---|---|---|---|---|---|---|---|
| Participants evaluable | 12 | 2 | 2 | 6 | 6 | 6 | 6 | 6 | 6 | 12 |
| Number of AEs | 5 (2) | 1 (0) | 0 | 3 (1) | 1 (0) | 6 (5) | 2 (2) | 0 | 5 (4) | 22 (17) |
| Participants with AEs | 3 (1) | 1 (0) | 0 | 2 (1) | 1 (0) | 3 (2) | 2 (2) | 0 | 3 (2) | 11 (10) |
| Participants with SAEs | 0 | 0 | 0 | 0 | 0 | 0 | 0 | 0 | 0 | 0 |
| Participans with severe AEs | 0 | 0 | 0 | 0 | 0 | 0 | 0 | 0 | 0 | 0 |
| Participants discontinued due to AEs | 0 | 0 | 0 | 0 | 0 | 0 | 0 | 0 | 0 | 0 |

Included all data collected since the first dose of study drug.
Except for the number of AEs, participants were counted only once per treatment in each row.
SAEs were according to the investigator's assessment.
AE, adverse event; FIH, first-in-human; SAE, serious adverse event
All doses were administered in the fasted state, except for 2 participants in placebo who received placebo in the fed state and 6 participants who received 100 mg both in the fasted and fed states.

Table S11.

**Human pharmacokinetics of PF-06882961.** Pharmacokinetic parameters following single oral doses of PF-06882961 to healthy human study participants.

| | Parameter Summary Statistics[a] by PF-06882961 Dose and Treatment | | | | | |
|---|---|---|---|---|---|---|
| Parameter | 3 mg (fasted) | 10 mg (fasted) | 30 mg (fasted) | 100 mg (fasted) | 100 mg (fed) | 300 mg (fasted) |
| N, n | 6, 6 | 6, 5 | 6, 5 | 6, 5 | 6, 6 | 12, 12 |
| $AUC_{inf}$, ng•hr/mL | 41.34 (50) | 88.96 (60) | 327.9 (42) | 1175 (51) | 923.6 (52) | 4864 (45) |
| $C_{max}$, ng/mL | 6.347 (59) | 13.24 (54) | 41.17 (62) | 176.5 (91) | 76.76 (23) | 732.3 (91) |
| $T_{max}$, hr | 2.00 (1.25-4.00) | 6.00 (3.00-6.00) | 4.00 (1.25-6.00) | 6.01 (0.750-6.03) | 6.04 (6.00-12.0) | 4.00 (0.300-12.2) |
| $t_{½}$, hr | 5.135 ± 4.72 | 4.336 ± 1.24 | 6.102 ± 4.34 | 5.968 ± 2.18 | 4.957 ± 0.882 | 5.677 ± 1.53 |

[a]Values are geometric mean (%CV) except $T_{max}$ (median and range) and $t_{½}$ (arithmetic mean ± SD).
AUC, area under the curve; $AUC_{inf}$, AUC from time 0 extrapolated to infinite time; %CV, percent coefficient of variation; $C_{max}$, maximum plasma concentration; hr, hour; N, number of participants in the treatment group and contributing to the summary statistics; n, number of participants with reportable $AUC_{inf}$, $t_{½}$; SD, standard deviation; $t_{1/2}$, half-life; $T_{max}$, time of the first occurrence of $C_{max}$
